# Identification of oxygen-independent pathways for pyridine-nucleotide and Coenzyme-A synthesis in anaerobic fungi by expression of candidate genes in yeast

**DOI:** 10.1101/2020.07.06.189415

**Authors:** Thomas Perli, Aurin M. Vos, Jonna Bouwknegt, Wijb J. C. Dekker, Sanne J. Wiersma, Christiaan Mooiman, Raúl A. Ortiz-Merino, Jean-Marc Daran, Jack T. Pronk

**Affiliations:** Department of Biotechnology, Delft University of Technology, Van der Maasweg 2629 Delft, The Netherlands

## Abstract

Neocallimastigomycetes are rare examples of strictly anaerobic eukaryotes. This study investigates how these anaerobic fungi bypass reactions involved in synthesis of pyridine nucleotide cofactors and coenzyme A that, in canonical fungal pathways, require molecular oxygen. Analysis of Neocallimastigomycete proteomes identified a candidate L-aspartate-decarboxylase (AdcA), and L-aspartate oxidase (NadB) and quinolinate synthase (NadA), constituting putative oxygen-independent bypasses for coenzyme A synthesis and pyridine nucleotide cofactor synthesis, respectively. The corresponding gene sequences indicated acquisition by ancient horizontal gene transfer event involving bacterial donors. To test whether these enzymes suffice to bypass corresponding oxygen-requiring reactions, they were introduced into *fms1Δ* and *bna2Δ Sacharomyces cerevisiae* strains. Expression of *nadA* and *nadB*, and *adcA* from the Neocallimastigomycetes *Piromyces finnis* and *Neocallimastix californiae*, respectively, conferred cofactor prototrophy under aerobic and anaerobic conditions. This study simulates how horizontal gene transfer can drive eukaryotic adaptation to anaerobiosis, and provides a basis for elimination of auxotrophic requirements in anaerobic industrial applications of yeasts and fungi.

## Introduction

Neocallimastigomycetes are obligately anaerobic fungi with specialised metabolic adaptations that allow them to play a key role in the degradation of recalcitrant plant biomass in herbivore guts [1]. Despite complicated cultivation techniques and lack of genetic-modification tools [2], several evolutionary adaptations of these eukaryotes to an anaerobic lifestyle have been inferred from biochemical studies [3-5]. Sequence analysis implicated extensive horizontal gene transfer (HGT) events as a key mechanism in these adaptations [6-8]. For example, instead of sterols, which occur in membranes of virtually all other eukaryotes [9] and whose biosynthesis involve multiple oxygen-dependent reactions [10], Neocallimastigomycetes contain tetrahymanol [3, 6]. This sterol surrogate [11] can be formed from squalene by a squalene:tetrahymanol cyclase (STC), whose structural gene in Neocallimastigomycetes showed evidence of acquisition by HGT from prokaryotes [6, 12]. Expression of an STC gene was recently shown to enable sterol-independent anaerobic growth of the model eukaryote *S. cerevisiae* [13].

Further exploration of oxygen-independent bypasses in Neocallimastigomycetes for intracellular reactions that in other eukaryotes require oxygen is relevant for a fundamental understanding of the requirements for anaerobic growth of eukaryotes. In addition, it may contribute to the elimination of nutritional requirements in industrial anaerobic applications of yeasts and fungi.

Most fungi are capable of *de novo* synthesis of pyridine-nucleotide cofactors (NAD^+^ and NADP^+^) and Coenzyme A (CoA) when grown aerobically. As exemplified by the facultatively anaerobic yeast *S. cerevisiae* [14], canonical fungal pathways for synthesis of these cofactors are oxygen dependent. In *S. cerevisiae*, biosynthesis of CoA involves formation of β-alanine by the oxygen-requiring polyamine oxidase Fms1 [15]. This intermediate is then condensed with pantoate to yield the CoA precursor pantothenate [16, 17] (Fig. 1A). Similarly, the yeast kynurenine pathway for *de novo* synthesis of NAD^+^ involves three oxygen-dependent reactions, catalyzed by indoleamine 2,3-dioxygenase (Bna2; EC 1.13.11.52), kynurenine 3-monooxygenase (Bna4; EC 1.14.13.9), and 3-hydroxyanthranilic-acid dioxygenase (Bna1; EC 1.13.11.6) [14] (Fig. 1B). The Neocallimastigomycete *Neocallimastix patricianum* has been shown to grow in synthetic media lacking precursors for pyridine-nucleotide and CoA synthesis [18]. This observation indicates that at least some anaerobic fungi harbour oxygen-independent pathways for synthesizing these essential cofactors. Genomes of Neocallimastigomycetes lack clear homologs of genes encoding the oxygen-requiring enzymes of the kynurenine pathway. Instead, their genomes were reported to harbour genes encoding an L-aspartate oxidase (NadB) and quinolinate synthase (NadA), two enzymes active in the bacterial pathway for NAD^+^ synthesis [6] (Fig. 1A). Since bacterial and plant aspartate oxidases can, in addition to oxygen, also use fumarate as electron acceptor [19, 20], it is conceivable that NadA and NadB may allow for oxygen-independent NAD^+^ synthesis in anaerobic fungi. No hypothesis has yet been forwarded on how these fungi may bypass the oxygen requirement for the canonical fungal CoA biosynthesis route.

**Fig. 1:**
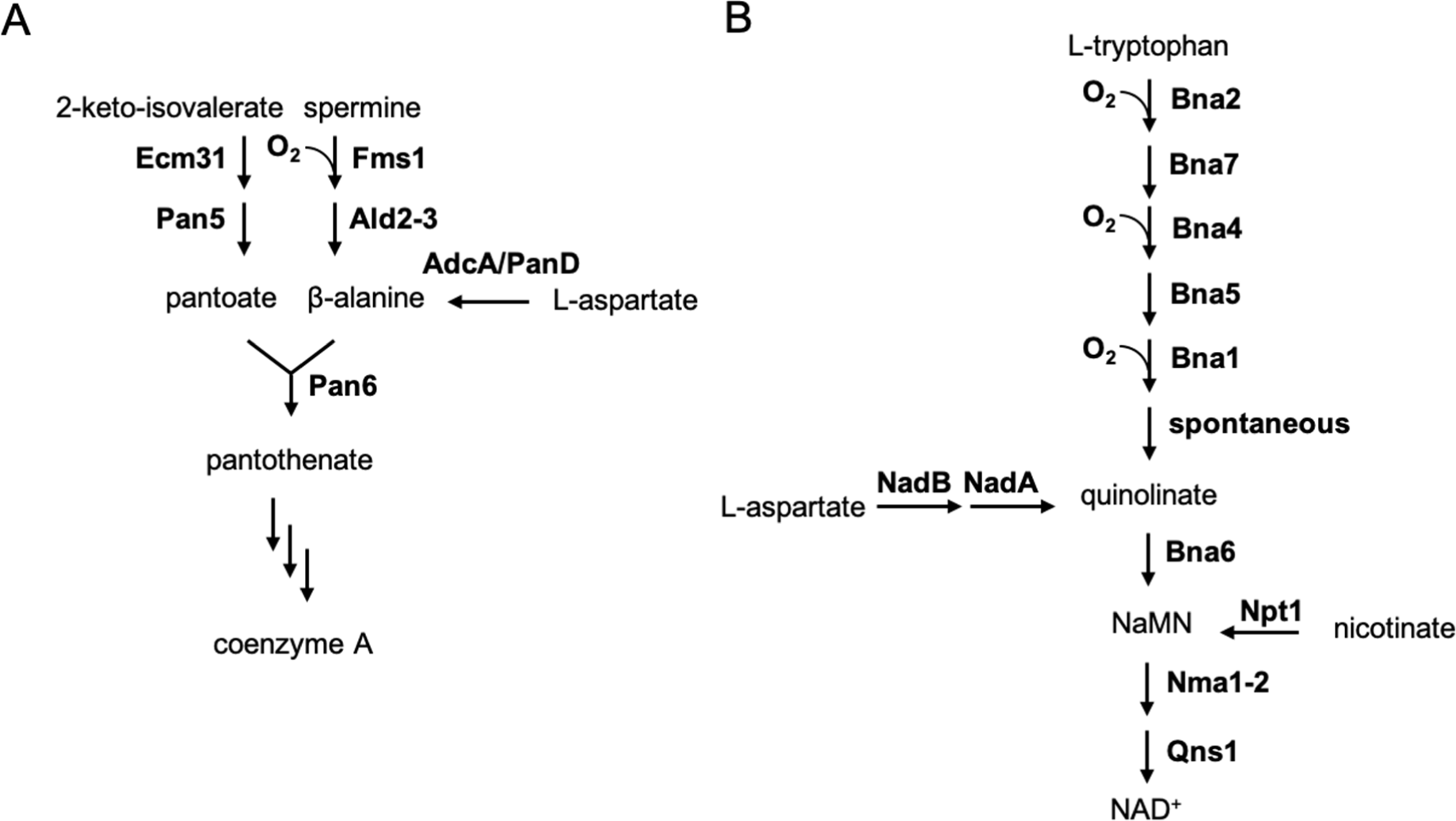
CoA and NAD^+^ biosynthetic pathways in *S. cerevisiae* and oxygen-independent alternatives. CoA synthesis includes the condensation of pantoate and β-alanine. In *S. cerevisiae* β-alanine is formed from spermine in two steps using the oxygen-dependent poly-amine oxidase Fms1 (A). Other organisms, including archaea, bacteria, and insects, can by-pass this oxygen requirement by synthesizing β-alanine from aspartate using L-aspartate decarboxylase (AdcA/PanD). NAD^+^ is synthesized via the kynurenine pathway in 9 reactions starting from tryptophan, 3 of which require oxygen (B). Other organisms that include plants and bacteria are able to bypass this oxygen requirement by synthesizing quinolinate from aspartate using L-aspartate oxidase and quinolinate synthase (NadB and NadA, respectively)

The goals of this study were to identify the pathway responsible for oxygen-independent synthesis of CoA in Neocallimastigomycetes and to investigate a possible role of NadA and NadB in oxygen-independent synthesis of pyridine-nucleotide cofactors. A candidate L-aspartate decarboxylase (Adc) encoding gene was identified by genome analysis of Neocallimastigomycetes and its phylogeny investigated. Candidate Neocallimastigomycete genes for L-aspartate oxidase and quinolinate synthase, previously reported to have been acquired by HGT [6], as well as the candidate Adc gene, were then functionally analysed by expression in *S. cerevisiae* strains devoid of essential steps in the native cofactor synthesis pathways. As controls, previously characterized genes involved in oxygen-independent NAD^+^ biosynthesis by *Arabidopsis thaliana* [21], and a previously characterized Adc encoding gene from the red flour beetle *Tribolium castaneum* (*TcPAND*) [22] were also expressed in the same *S. cerevisiae* strains. The results demonstrate how heterologous expression studies in yeast can provide insight into evolutionary adaptations to anaerobic growth and selective advantages conferred by proposed HGT events in Neocallimastigomycetes. In addition, they identify metabolic engineering strategies for eliminating oxygen requirements for cofactor biosynthesis in anaerobic industrial applications of *S. cerevisiae*.

## Material and Methods

### Strains, media and maintenance

*S. cerevisiae* strains used and constructed in this study (Table 1) were derived from the CEN.PK lineage [23]. Yeast cultures were routinely propagated in YP (10 g L^-1^ Bacto yeast extract [Becton, Dickinson and Co., Sparks, MD], 20 g L^-1^ Bacto peptone [Becton, Dickinson and Co]) or synthetic medium (SM) [24]. YP and SM were autoclaved at 121 °C for 20 min. SM was then supplemented with 1 mL L^-1^ of filter-sterilized vitamin solution (0.05 g L^-1^ D-(+)-biotin, 1.0 g L^-1^ D-calcium pantothenate, 1.0 g L^-1^ nicotinic acid, 25 g L^-1^ myo-inositol, 1.0 g L^-1^ thiamine hydrochloride, 1.0 g L^-1^ pyridoxol hydrochloride, 0.20 g L^-1^ 4-aminobenzoic acid). Where indicated, nicotinic acid or pantothenic acid were omitted from the vitamin solution, yielding SM without nicotinic acid (SMΔnic) and SM without pantothenic acid (SMΔpan), respectively. A concentrated glucose solution was autoclaved separately for 15 min at 110 °C and added to SM and YP to a concentration of 20 g L^-1^ or 50 g L^-1^, yielding SMD and YPD, respectively. SMD with urea or acetamide instead of ammonium sulfate (SMD-urea and SMD-Ac, respectively) were prepared as described previously [25, 26]. For anaerobic growth experiments, sterile media were supplemented with Tween 80 (polyethylene glycol sorbate monooleate, Merck, Darmstadt, Germany) and ergosterol (≥95 % pure, Sigma-Aldrich, St. Louis, MO) as described previously [27]. Yeast strains were grown in 500-mL shake flasks containing 100 mL medium or in 100-mL shake flasks containing 20 mL medium. Shake-flask cultures were incubated at 30 °C and shaken at 200 rpm in an Innova Incubator (Brunswick Scientific, Edison, NJ). Solid media were prepared by adding 15 g L^-1^ Bacto Agar (Becton, Dickinson and Co) and, when indicated, 200 mg L^-1^ G418 (Thermo Scientific, Waltham, MA). After genotyping, engineered strains were restreaked twice to select single clones. Removal of the gRNA carrying plasmid was done as previously described [28]. Stock cultures were prepared by adding glycerol to a final concentration of 33 % (v/v), frozen and stored at −80°C.

**Table 1:** *S. cerevisiae* strains used in this study.

| Name | Relevant genotype | Parental strain | Reference |
| --- | --- | --- | --- |
| CEN.PK113-7D | MATa *URA3* | - | [23] |
| CEN.PK113-5D | MATa *ura3-52* | - | [23] |
| IMX585 | MATa *can1Δ::Spycas9-natNT2 URA3* | CEN.PK113-7D | [28] |
| IMX581 | MATa *ura3-52 can1Δ::Spycas9-natNT2* | CEN.PK113-5D | [28] |
| IMX2292 | MATa *can1Δ::Spycas9-natNT2 URA3 fms1Δ* | IMX585 | [74] |
| IMK877 | MATa *can1Δ::Spycas9-natNT2 URA3 bna2Δ* | IMX585 | This study |
| IMX2301 | MATa *can1Δ::Spycas9-natNT2 URA3 bna2Δ sga1::pTDH3-PfnadA-tENO1 pCCW12-PfnadB-tENO2* | IMK877 | This study |
| IMX2302 | MATa *can1Δ::Spycas9-natNT2 URA3 bna2Δ sga1::pTDH3-AtNADA-tENO1 pCCW12-AtNADB-tENO2* | IMK877 | This study |
| IMX2293 | MATa *ura3-52 can1Δ::Spycas9-natNT2 fms1Δ* | IMX581 | This study |
| IMX2300 | MATa *ura3-52::pTDH3-NcadcA-tENO2 URA3 can1Δ::Spycas9-natNT2 fms1Δ* | IMX2293 | This study |
| IMX2300-1 | MATa *ura3-52::pTDH3-NcadcA-tENO2 URA3 can1Δ::Spycas9-natNT2 fms1Δ* Colony isolate 1 | IMX2300 | This study |
| IMX2305 | MATa *ura3-52::pRPL12b-TcPAND-tTDH1 URA3 can1Δ::Spycas9-natNT2 fms1Δ* | IMX2293 | This study |

*Spy*: *Streptococcus pyogenes*; *Pf*: *Piromyces finnis*; *Nc*: *Neocallimastix californiae*; *At*: *Arabidopsis thaliana*; *Tc*: *Tribolium castaneum*.

### Molecular biology techniques

DNA was PCR amplified with Phusion Hot Start II High Fidelity Polymerase (Thermo Scientific) and desalted or PAGE-purified oligonucleotide primers (Sigma Aldrich) by following manufacturers’ instructions. DreamTaq polymerase (Thermo Scientific) was used for diagnostic PCR. Oligonucleotide primers used in this study are listed in Supplementary Table S1. PCR products were separated by gel electrophoresis using 1 % (w/v) agarose gel (Thermo Scientific) in TAE buffer (Thermo Scientific) at 100 V for 25 min and purified with either GenElute PCR Clean-Up Kit (Sigma Aldrich) or with Zymoclean Gel DNA Recovery Kit (Zymo Research, Irvine, CA). Plasmids were purified from *E. coli* using a Sigma GenElute Plasmid Kit (Sigma Aldrich). Yeast genomic DNA was isolated with the SDS/LiAc protocol [29]. Yeast strains were transformed with the lithium acetate method [30]. Four to eight single colonies were re-streaked three consecutive times on selective media and diagnostic PCR were performed to verify their genotype. *Escherichia coli* XL1-blue was used for chemical transformation [31]. Plasmids were then isolated and verified by either restriction analysis or by diagnostic PCR. Lysogeny Broth (LB; 10 g L^-1^ Bacto Tryptone, 5 g L^-1^, Bacto Yeast Extract with 5 g L^-1^ NaCl) was used to propagate *E. coli* XL1-Blue. LB medium was supplemented with 100 mg L^-1^ ampicillin for selection of transformants. The overnight grown bacterial cultures were stocked by adding sterile glycerol at a final concentration of 33 % (v/v) after which samples were frozen and stored at −80 °C.

### Plasmid construction

Plasmids used and cloned in this study are shown in Table 2. Plasmids carrying two copies of the same gRNA were cloned by Gibson assembly [28, 32]. In brief, an oligo carrying the gene-specific 20 bp target sequence and a homology flank to the plasmid backbone was used to amplify the fragment carrying the 2µm origin of replication sequence by using pROS13 as template. The backbone linear fragment was amplified using primer 6005 and pROS11 as template [33]. The two fragments were then gel purified and assembled *in vitro* using the NEBuilder HiFi DNA Assembly Master Mix (New England BioLabs, Ipswich, MA) following manufacturer’s instructions. Transformants were selected on LB plates supplemented with 100 mg L^-1^ ampicillin or 50 mg L^-1^ kanamycin. Primer 11861 was used to amplify the 2µm fragment containing two identical gRNA sequences for targeting *BNA2*. The PCR product was then cloned in a pROS11 backbone yielding plasmid pUDR315. The coding sequences for *AtNADA, AtNADB, PfnadA, PfnadB*, and *NcadcA* were codon-optimized for expression in *S. cerevisiae* and ordered as synthetic DNA through GeneArt (Thermo Fisher Scientific). The plasmids carrying the expression cassettes for *TcPAND, AtNADA, AtNADB, PfnadA* and *PfnadB* were cloned by Golden Gate assembly using the Yeast Toolkit (YTK) DNA parts [34]. These plasmids were cloned using the pYTK096 integrative backbone that carries long homology arms to the *URA3 locus* and a *URA3* expression cassette allowing for selection on SM lacking uracil. The *TcPAND* coding sequence was amplified using the primer pair 11877/11878 and pCfB-361 as template. Then, the linear *TcPAND* gene and plasmids pUD1096, pUD1097, pUD652, and pUD653 carrying the coding sequence for *AtNADA, AtNADB, PfnadA*, and *PfnadB*, respectively, were combined together with YTK-compatible part plasmids in BsaI (New England BioLabs) golden gate reactions to yield plasmid pUDI168, pUDI245, pUDE931, pUDI243, and pUDI244, respectively. A detailed list of the YTK-compatible parts used for constructing each plasmid can be found in Supplementary Table S2.

**Table 2:**
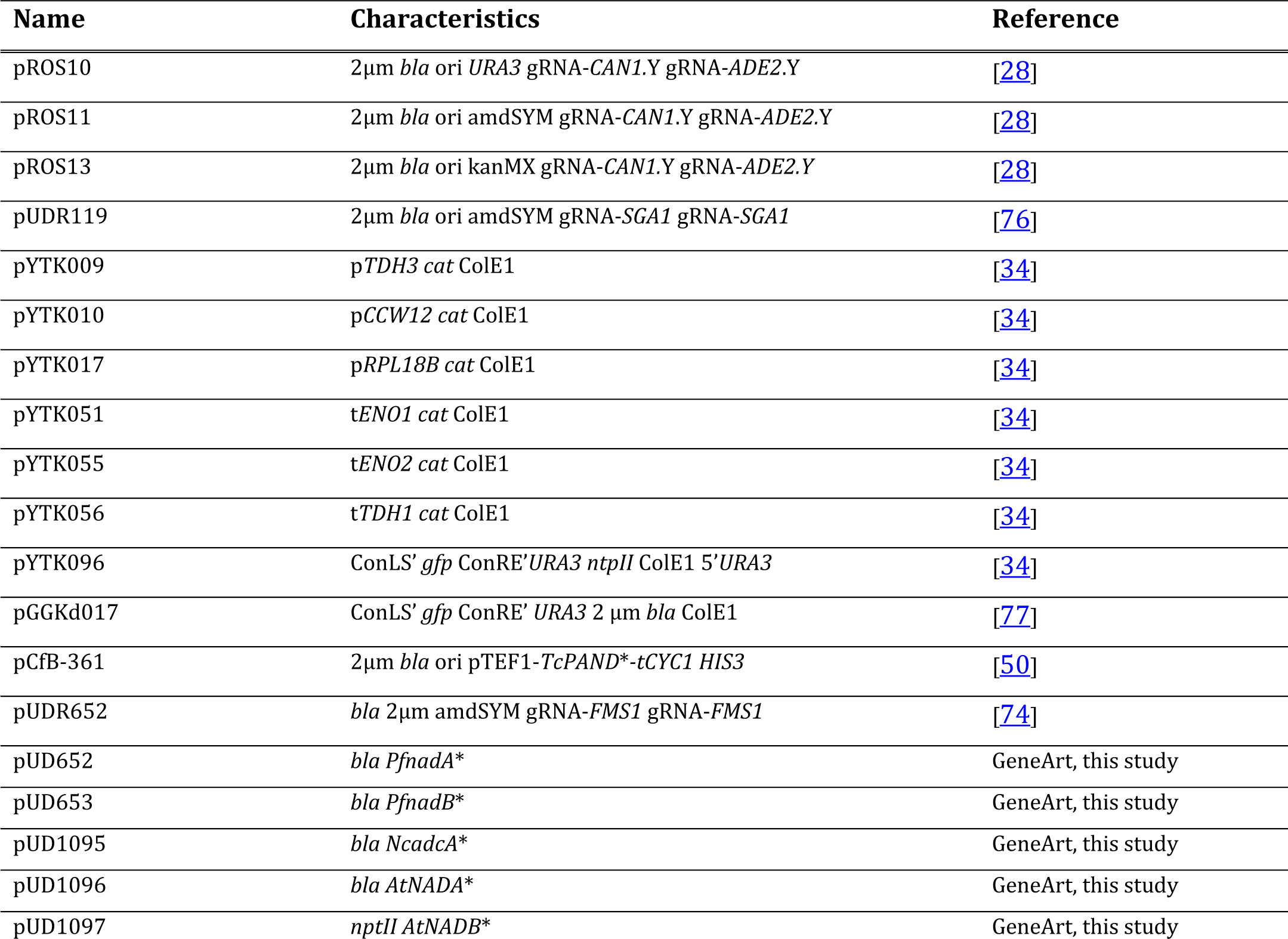
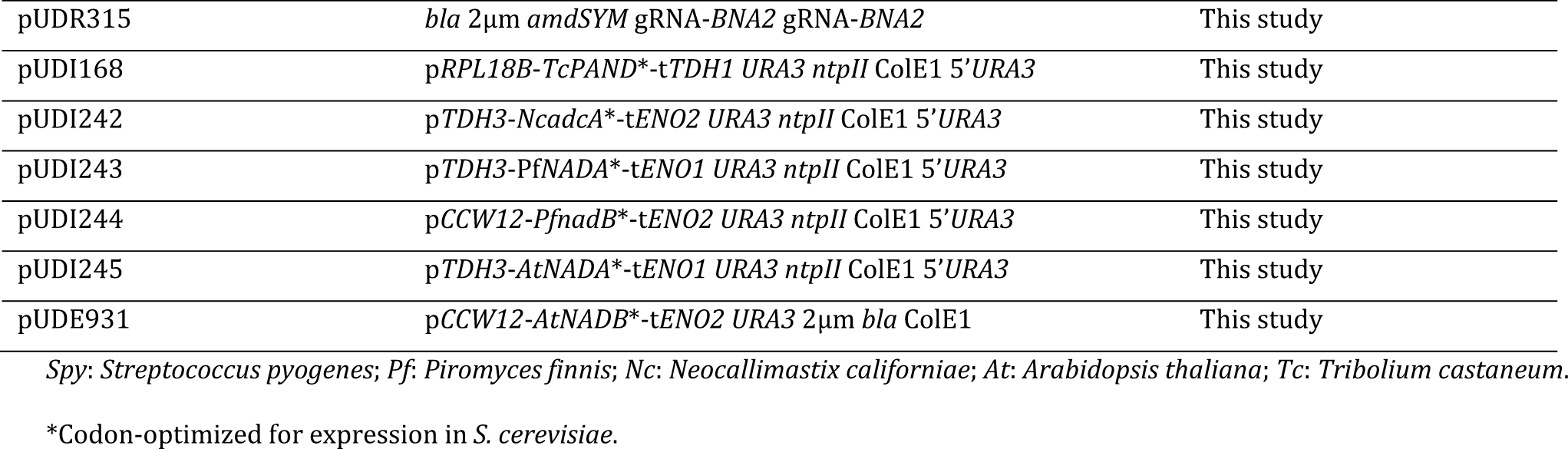
plasmids used in this study.

The plasmid carrying the expression cassette for *NcadcA* was cloned by Gibson assembly. The *pTDH3* promoter, the *NcadcA* coding sequence, the *tENO2* terminator and the pYTK0096 backbone were amplified by PCR using primer pairs 16721/16722, 16723/16724, 16725/16726, and 16727/16728 respectively, using pYTK009, pUD1095, pYTK055, and pYTK096 as template, respectively. Each PCR product was then gel purified and combined in equimolar amounts in a Gibson reaction that yielded pUDI242.

### Strain construction

*S. cerevisiae* strains were transformed using the LiAc/SS-DNA/PEG and CRISPR/Cas9 method [28, 30, 35]. For deletion of the *BNA2* gene, IMX585 (*can1Δ::Spycas9-natNT2*) was transformed with 500 ng of the *BNA2* targeting gRNA plasmid pUDR315 together with 500 ng of the annealed primer pair 11862/11863 as repair dsDNA oligo, yielding strain IMK877. The resulting strain was then used for the integration of the two heterologous *NADB-A* pathways. Expression cassettes for *AtNADA, AtNADB, PfnadA, PfnadB*, were amplified from plasmids pUDI245, pUDE931, pUDI243, pUDI244, respectively, using primer pairs 13123/13124, 13125/10710, 13123/13124, 13125/10710, respectively. Then, 500 ng of each pair of gel purified repair cassettes were co-transformed in IMK877 together with 500 ng the *SGA1* targeting gRNA plasmid, yielding IMX2302 (*sga1::AtNADA AtNADB*) and IMX2301 (*sga1::PfnadA PfnadB*).

For deletion of the *FMS1* gene, IMX581 (*can1Δ::Spycas9-natNT2 ura3-52*) was transformed with 500 ng of the *FMS1* targeting gRNA plasmid pUDR652 together with 500 ng of the annealed primer pair 13527/13528 as repair dsDNA oligo, resulting in IMX2293. Then, 500 ng of plasmids pUDI168 and pUDI242 carrying the expression cassettes for *TcPAND* and *NcadcA*, respectively, were NotI (Thermo Fisher) digested and separately transformed in IMX2293, yielding IMX2305, and IMX2300, respectively. Selection of IMX2305 and IMX2300 was done on SMD agar plate since the integration of each Adc encoding cassette also restored the *URA3* phenotype. In contrast, selection of IMK877 was done on SMD-Ac agar plates while selection of IMX2302, IMX2301, and IMX2293 was done YPD-G418 agar plates. Strains IMK877, IMX2300, IMX2302, and IMX2301 were stocked in SMD, while IMX2305 and IMX2293 were stocked in SMDΔpan and YPD, respectively.

### Aerobic growth studies in shake flasks

For the determination of the specific growth rate of the engineered strains under aerobic conditions, a frozen aliquot was thawed and used to inoculate a 20 mL wake-up culture that was then used to inoculate a pre-culture in a 100 mL flask. The exponentially growing pre-culture was then used to inoculate a third flask to an initial OD_660_ of 0.2. The flasks were then incubated, and growth was monitored using a 7200 Jenway Spectrometer (Jenway, Stone, United Kingdom). Specific growth rates were calculated from at least five time-points in the exponential growth phase of each culture. Wake-up and pre-cultures of IMX2301 and IMX2302 were grown in SMDΔnic. Wake-up and pre-cultures of IMX2300 and IMX2305 were grown in SMDΔpan while wake-up and pre-cultures of IMK877 and IMX2292 were grown in SMD.

### Anaerobic growth studies in shake flasks

Anaerobic shake-flask based experiments were performed in Lab Bactron 300 anaerobic workstation (Sheldon Manufacturing Inc., Cornelius, OR) containing an atmosphere of 85 % N_2_, 10 % CO_2_, and 5 % H_2_. Flat-bottom shake flasks of 50-mL were filled with 40 mL SMD-urea media containing 50 g L^-1^ glucose as carbon source, to ensure depletion of the vitamin/growth factor of interest, and 20 g L^-1^ glucose for the first transfer. Media weer supplemented with vitamins, with and without pantothenic acid or nicotinic acid as indicated, and in all cases supplemented with Tween 80 and ergosterol. Sterile medium was placed inside the anaerobic chamber 24 h prior to inoculation for removal of oxygen. Traces of oxygen were continuously removed with a regularly regenerated Pd catalyst for H_2_-dependent oxygen removal placed inside the anaerobic chamber. Aerobic overnight shake-flask cultures on SMD-urea were used to inoculate the anaerobic shake flask without pantothenic acid or without nicotinic acid at an initial OD_600_ of 0.2. Cultures were cultivated at 30 °C with continuous stirring at 240 rpm on IKA KS 260 Basic orbital shaker platform (Dijkstra Verenigde BV, Lelystad, the Netherlands). Periodic optical density measurements at a wavelength of 600 nm using an Ultrospec 10 cell density meter (Biochrom, Cambridge, United Kingdom) inside the anaerobic environment were used to follow the growth over time. After growth had ceased and the OD_600_ no longer increased the cultures were transferred to SMD-urea with 20 g L^-1^ glucose at an OD_600_ of 0.2 [27].

### Anaerobic bioreactor cultivation

Anaerobic bioreactor batch cultivation was performed in 2-L laboratory bioreactors (Applikon, Schiedam, the Netherlands) with a working volume of 1.2 L. Bioreactors were tested for gas leakage by applying 0.3 bar overpressure while completely submerging them in water before autoclaving. Anaerobic conditions were maintained by continuous sparging of the bioreactor cultures with 500 mL N_2_ min^−1^ (≤0.5 ppm O_2_, HiQ Nitrogen 6.0, Linde Gas Benelux, Schiedam, the Netherlands). Oxygen diffusion was minimized by using Fluran tubing (14 Barrer O2, F-5500-A, Saint-Gobain, Courbevoie, France) and Viton O-rings (Eriks, Alkmaar, the Netherlands). Bioreactor cultures were grown on either SMDΔpan or SMDΔnic with ammonium sulfate as nitrogen source. pH was controlled at 5 using 2 M KOH. The autoclaved mineral salts solution was supplemented with 0.2 g L^−1^ sterile antifoam emulsion C (Sigma-Aldrich). Bioreactors were continuously stirred at 800 rpm and temperature was controlled at 30 °C. Evaporation of water and volatile metabolites was minimized by cooling the outlet gas of bioreactors to 4 °C in a condenser. The outlet gas was then dried with a PermaPure PD-50T-12MPP dryer (Permapure, Lakewood, NJ) prior to analysis. CO_2_ concentrations in the outlet gas were measured with an NGA 2000 Rosemount gas analyser (Emerson, St. Louis, MO). The gas analyser was calibrated with reference gas containing 3.03 % CO_2_ and N6-grade N_2_ (Linde Gas Benelux, Schiedam, The Netherlands). Frozen glycerol stock cultures were used to inoculate aerobic 100 mL shake flask cultures on either SMDΔpan or SMDΔnic. Once the cultures reached OD_660_ > 5, a second 100 mL aerobic shake-flask pre-culture on the same medium was inoculated. When this second pre-culture reached the exponential growth phase, biomass was harvested by centrifugation at 3000 g for 5 min and washed with sterile demineralized water. The resulting cell suspension was used to inoculate anaerobic bioreactors at an OD_660_ of 0.2.

### Analytical methods

Biomass dry weight measurements of the bioreactor batch experiments were performed using pre-weighed nitrocellulose filters (0.45 µm, Gelman Laboratory, Ann Arbor, MI). 10 mL culture samples were filtrated and then the filters were washed with demineralized water prior to drying in a microwave oven (20 min at 360 W) and weight measurement. Metabolite concentrations in culture supernatants were analysed by high-performance liquid chromatography (HPLC). In brief, culture supernatants were loaded on an Agilent 1260 HPLC (Agilent Technologies, Santa Clara, CA) fitted with a Bio-Rad HPX 87 H column (Bio-Rad, Hercules, CA). The flow rate was set at 0.6 mL min^-1^ and 0.5 g L^-1^ H_2_SO_4_ was used as eluent. An Agilent refractive-index detector and an Agilent 1260 VWD detector were used to detect culture metabolites [36]. An evaporation constant of 0.008 divided by the volume in liters, was used to correct HPLC measurements of ethanol in the culture supernatants, taking into account changes in volume caused by sampling [37]. Statistical analysis on product yields was performed by means of an unpaired two-tailed Welch’s t-test.

### Homology and phylogenetic analyses

A set of 51 aminoacid sequences previously used to discriminate between glutamate decarboxylases and L-aspartate decarboxylases [38] was re-used to identify candidate Neocallimastigomycete Adc sequences. These sequences were used as queries against a database containing all 58109 Neocallimastigomycete proteins deposited in Uniprot trembl (Release 2019_02), which represented 5 species (*Neocallimastix californiae, Anaeromyces robustus, Piromyces sp* E2, *Piromyces finnis*, and *Pecoramyces ruminatum*), and extracted according to the NCBI taxid 451455. Sequence homology was analysed using BLASTP 2.6.0+ [39] with 10^−6^ as e-value cut-off resulting in 13 Neocallimastigomycete sequences as shared hits from all 51 queries (Supplementary Table S3). Four of these sequences showing homology to experimentally characterised proteins with L-aspartate-decarboxylase (Adc) activity originated from *N. californiae*, and were checked for RNAseq read coverage and splicing junction support revealing A0A1Y1ZL74 as best candidate (Supplementary Fig. S1).

A0A1Y1ZL74, also referred to as *NcadcA*, was used for a second round of homology search using HMMER 3.2 [40] against a database with a balanced representation of taxa across the 3 domains of life. This database was built from Uniprot Release 2019_02 to include all refseq sequences from Bacteria (taxid 2), Eukarya (taxid 2759), and Archaea (taxid 2157; TrEMBL and Swiss-Prot categories were also included in this case). Selection for hits with more than 60% alignment length and evalue < 10^−6^ resulted in a total of 325 sequences (103 from Bacteria, 101 from Eukaryotes, and 121 from Archaea).

The set of 325 A0A1Y1ZL74 homologous sequences, together with those from Tomita *et. al*. (2015) [38] were aligned with Clustal Omega 1.2.4 [41] and then used to build a maximum likelihood phylogenetic tree with RAxML-NG 0.8.1 [42] using default parameters with the exception of the use of the PROTGTR+FO model and 100 bootstrap replicates. The resulting phylogenetic tree drawn with iTOL [43] is shown in Fig. 2, corresponding alignments and trees are provided in Supplementary Files 1 and 2. Multiple sequence alignment was also performed with Clustal omega 1.2.4 [41] to compare selected aminoacid sequences showing candidate and experimentally characterised Adcs, against bacterial PanDs. These sequences and alignments are shown in Supplementary File 3.

**Fig. 2:**
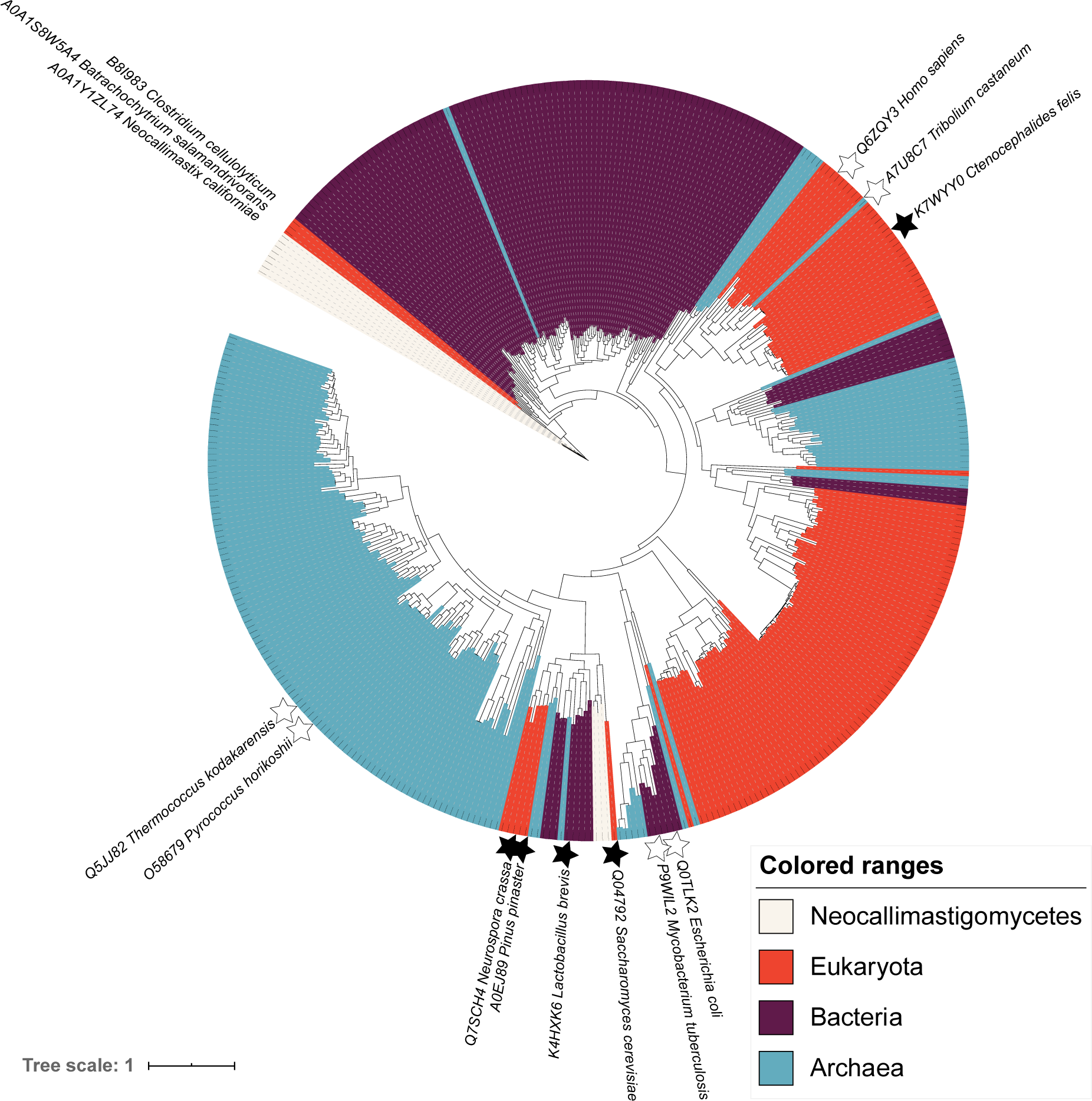
Maximum likelihood phylogenetic tree of aspartate decarboxylase and glutamate decarboxylase sequences. Sequences of proteins with demonstrated enzyme activity are marked with white stars (L-aspartate decarboxylases) or black stars (glutamate decarboxylases). A version of this tree with all sequence identifiers, branch support, distances and bootstrap values, is provided in the Supplementary File 2. An interactive visualisation can be accessed in https://itol.embl.de/shared/rortizmerino

### Whole-genome sequencing and analysis

Genomic DNA of strains IMX2300 and IMX2300-1 was isolated with a Blood & Cell Culture DNA Kit with 100/G Genomics-tips (QIAGEN, Hilden, Germany) according to the manufacturers’ instructions. The Miseq Reagent Kit v3 (Illumina, San Diego, CA), was used to obtain 300 bp reads for paired-end sequencing. Genomic DNA was sheared to an average of 550 bp fragments using an M220 ultrasonicator (Covaris, Wolburn, MA). Libraries were prepared by using a TruSeq DNA PCR-Free Library Preparation kit (Illumina) following manufacturer’s instructions. The samples were quantified by qPCR on a Rotor-Gene Q PCR cycler (QIAGEN) using the Collibri Library quantification kit (Invitrogen Carlsbad, CA). Finally, the library was sequenced using an Illumina MiSeq sequencer (Illumina, San Diego, CA) resulting in a minimum 50-fold read coverage. Sequenced reads were mapped using BWA 0.7.15-r1142-dirty [44] against the CEN.PK113-7D genome [45] containing an extra contig with the relevant integration cassette. Alignments were processed using SAMtools 1.3.1 [46], and sequence variants were called using Pilon 1.18 [47], processed with ReduceVCF 12 (https://github.com/AbeelLab/genometools/blob/master/scala/abeel/genometools/reducevcf/ReduceVCF.scala), and annotated using VCFannotator (http://vcfannotator.sourceforge.net/) against GenBank accession GCA_002571405.2 [48].

### Data availability

DNA sequencing data of the *Saccharomyces cerevisiae* strains IMX2300 and IMX2300-1 were deposited at NCBI (https://www.ncbi.nlm.nih.gov/) under BioProject accession number PRJNA634013. All measurement data and calculations used to prepare Fig. 3-4 and Tables 3-4 of the manuscript are available at the 4TU.Centre for research data repository (https://researchdata.4tu.nl/) under doi: 10.4121/uuid:c3d2326d-9ddb-469a-b889-d05a09be7d97.

**Table 3:**
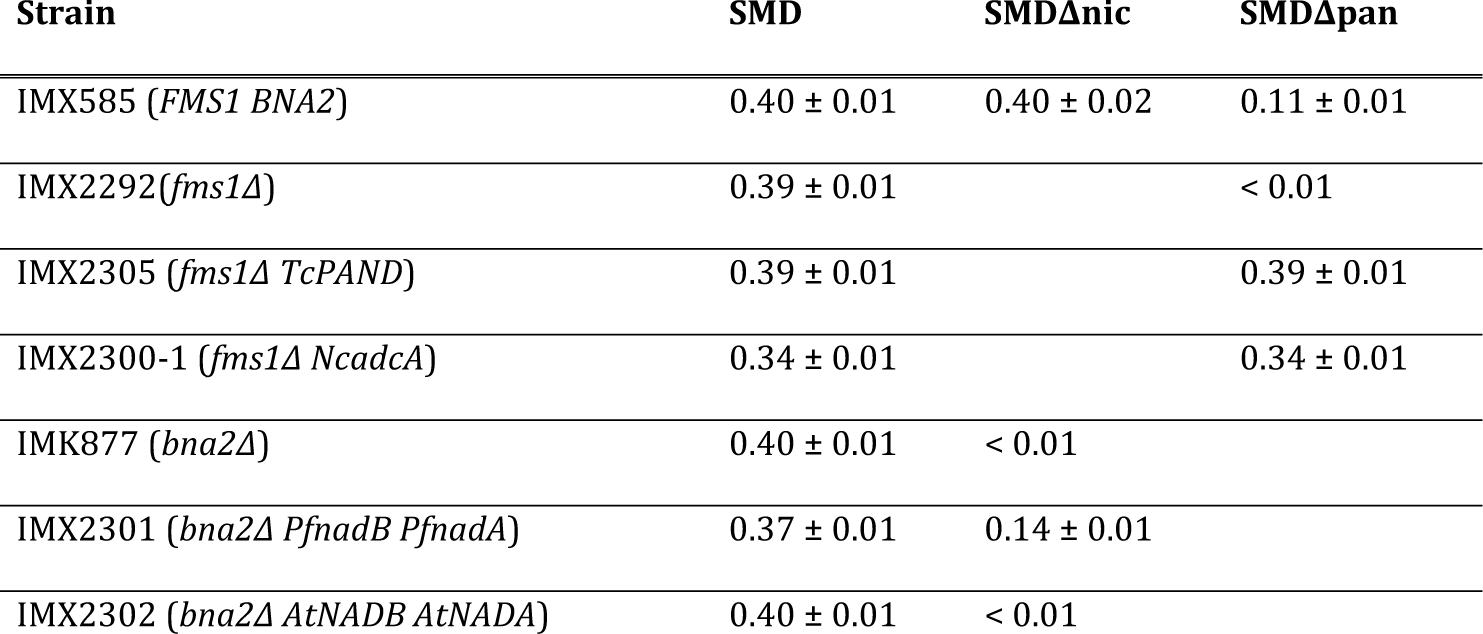
Aerobic characterization of engineered strains. Specific growth rates of *S. cerevisiae* strains grown in SMD, SMDΔnic and SMDΔpan media. The values are average and mean deviation of data from at least two independent cultures of each strain

**Table 4:**
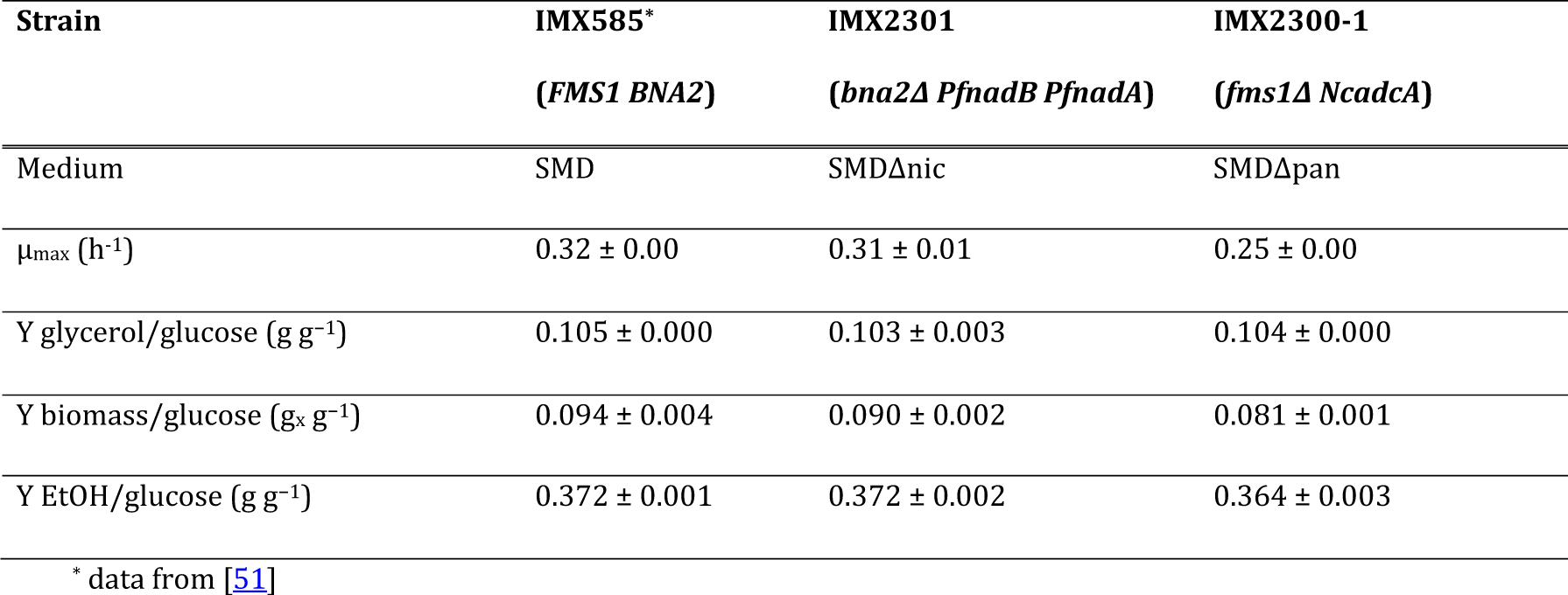
Maximum specific growth rate (µ_max_) and yields of glycerol, biomass and ethanol on glucose in anaerobic bioreactor batch cultures of *S. cerevisiae* strains IMX585, IMX2301 and IMX2300-1. Cultures were grown on SMD, SMDΔnic, or SMDΔpan, respectively, with 20 g L^-1^ glucose as carbon source (pH = 5). Growth rates and yields were calculated from the exponential growth phase. The ethanol yield was corrected for evaporation. Values represent average and mean deviation of data from independent cultures (n = 2). Carbon recovery in all fermentations was between 95 and 100%

**Fig. 3:**
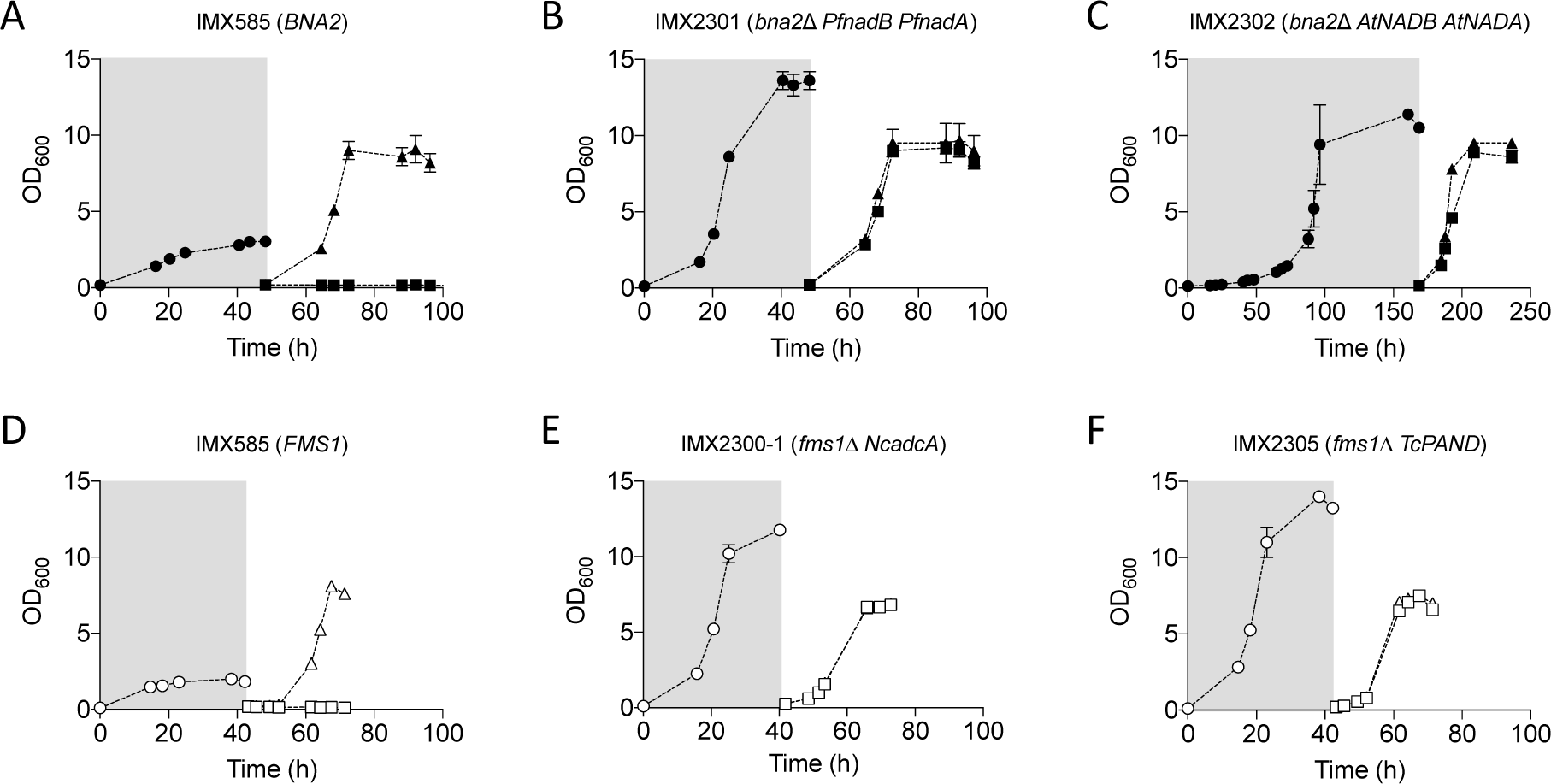
Anaerobic growth of *S. cerevisiae* strains dependent or independent on supplementation of nicotinic acid (NA) or pantothenic acid (PA) in SMD medium containing Tween 80 and ergosterol. Strains IMX585 (A), IMX2301 (*bna2*Δ *PfnadB PfnadA*) (B), and IMX2302 (*bna2*Δ *AtNADB AtNADA*) (C) transferred to medium with 2 % glucose with (▴) or without (▪) nicotinate after a carry-over phase in SMDΔnic containing 4 % glucose (• in grey box). Strains IMX585 (D), IMX2300-1 (*fms1*Δ *NcadcA*) (E), and IMX2305 (*fms1*Δ *TcPAND*)(F) transferred to medium with (△) or without (□) pantothenate after a carry-over phase in SMDΔpan containing 4 % glucose (○ in grey box). Anaerobic condition in the chamber were maintained using a palladium catalyst and a 5 % hydrogen concentration. Error bars represent the mean deviation of independent cultures (n=2)

## Results

### Identification of a candidate oxygen-independent L-aspartate decarboxylase involved in CoA synthesis in anaerobic fungi

Decarboxylation of L-aspartate to β-alanine by L-aspartate decarboxylase (Adc), an enzyme that occurs in all domains of life [38], enables an oxygen-independent alternative for the canonical fungal pathway for CoA synthesis (Fig. 1A). A set of 51 amino acid sequences of Adcs homologs listed by Tomita *et al*. (2015)[38] were used as queries against all Neocallimastigomycete proteins deposited in the TrEMBL section of the UNIPROT database. This search yielded 13 Neocallimastigomycete hits (e-value < 10^−6^, Supplementary Table S3), 4 of which originated from *N. californiae*. Only one of these hits, A0A1Y1ZL74, did not reveal annotation errors upon RNAseq read mapping and showed the highest read coverage (Supplementary Fig. S1) and was selected as best candidate Adc encoding gene.

The sequence A0A1Y1ZL74 (hereafter refered to as *Nc*AdcA) was used for a second round of homology search against a broader set of Adc sequences, with a similar sequence representation of taxa across the 3 domains of life (103 sequences from Bacteria, 101 from Eukarya, and 121 from Archaea). The resulting set of *Nc*AdcA homologs, together with the set definded by Tomita *et al*. (2015) [38], were subjected to multiple sequence alignment. A subsequent phylogenetic tree (Fig. 2) showed that *Nc*Adc sequences are closely related to those of chytrid fungi (A0A0L0HIP1 from *Spizellomyces punctatus*, A0A1S8W5A4 from *Batrachochytrium salamandrivorans* and F4NWP2 from *Batrachochytrium dendrobatidis*) and from the rumen-associated anaerobic bacterium *Clostridium cellulolyticum* (B8I983). Neocallimastigomycete, chytrid and *C. cellulolyticum* Adc homologs were more closely related to each other than to characterised eukaryotic Adc and bacterial PanD sequence*s*. These results indicate that an ancestor of *C. cellulolyticum* donated an Adc-encoding sequence to a common ancestor of chytrids and Neocallimastigomycetes.

Comparison of bacterial PanDs (Q0TLK2 from *E. coli* and P9WIL2 from *Mycobacterium tuberculosis*) against Adcs from other bacteria (B8I983 from *C. cellulolyticum*), and eukaryotes (including A7U8C7 from *Tribolium castaneum*) showed only little sequence homology between *Nc*Adc*s*, known bacterial PanDs, and eukaryotic Adcs (Supplementary File 3). The only conserved region encompassed the full length of PanDs (126-139 amino acids), which represents less than 60 % of the full length of other Adc sequences (*e*.*g. Nc*Adc*A* is 625 amino acids long).

### Neocallimastigomycete *PfnadB, PfnadA* and *NcadcA* genes support aerobic pyridine-nucleotide and CoA synthesis in yeast

Neocallimastigomycetes were previously reported to have acquired an L-aspartate oxidase (*nadB*) and a quinolinate synthase gene (*nadA*) by HGT [6]. Hence, UNIPROT entries A0A1Y1V2P1 and A0A1Y1VAT1 from *Piromyces finnis* were functionally reassigned as NadA and NadB candidates and the corresponding genes were tentatively named *PfnadB* and *PfnadA*. These sequences, together with *NcadcA*, were codon-optimised and tested to bypass the corresponding oxygen-requiring reactions in *S. cerevisiae*.

The *BNA2* and *FMS1* genes of *S. cerevisiae* were deleted by Cas9-mediated genome editing. The inability of strain IMK877 (*bna2Δ*) to synthesize quinolinic acid and of strain IMX2292 (*fms1Δ*) to synthesize β-alanine was evident from their inability to grow on glucose synthetic medium (SMD) lacking nicotinic acid or pantothenate, respectively (Table 3). Strain IMK877 was used for heterologous complementation studies with codon-optimized expression cassettes for *PfnadB* and *PfnadA*, while an expression cassette for *N. californiae NcadcA* (A0A1Y1ZL74) was introduced into strain IMX2292. Congenic strains expressing previously characterized *NADB* and *NADA* genes from *Arabidopsis thaliana* (*At*NadB and *At*NadA, Q94AY1 and Q9FGS4)[21], and a previously characterized gene from *Tribolium castaneum* encoding an aspartate decarboxylase (*Tc*PanD, A7U8C7)[22] were tested in parallel.

Aerobic growth of the engineered *S. cerevisiae* strains was characterized in shake-flask cultures on SMD or on either SMDΔnic or SMDΔpan (Table 1). In contrast to the reference strain IMK877 (*bna2Δ*), *S. cerevisiae* IMX2301 (*bna2Δ PfnadB PfnadA*) grew in SMDΔnic, indicating complementation of the *bna2Δ*-induced nicotinate auxotrophy by *PfnadB* and *PfnadA*. However, the specific growth rate of the engineered strain in these aerobic cultures was approximately 3-fold lower than that of the reference strain IMX585 (*BNA2*, Table 1). Strain IMX2302 (*bna2Δ AtNADB AtNADA*) did not grow in SMDΔnic, suggesting that the plant NadB and/or NadA proteins were either not functionally expressed or not able to complement the nicotinate auxotrophy in these aerobic yeast cultures.

Strain IMX2300 (*fms1Δ NcadcA*) grew in SMDΔpan, indicating complementation of the panthotenate auxotrophy. However, this strain reproducibly showed a lag phase of approximately 48 h upon its first transfer from SMD to SMDΔpan, and grew exponentially thereafter at a rate of 0.34 ± 0.01h^-1^. To explore whether the lag phase of strain IMX2300 reflected selection of a spontaneous mutant, it was subjected to three sequential transfers in SMDΔpan. A single-colony isolate, IMX2300-1 from the adapted population showed a specific growth rate of 0.34 ±0.01 h^-1^ in both SMD and SMDΔpan (Table 1). Whole-genome sequencing of IMX2300-1 did not reveal any mutations in coding DNA sequences that were considered physiologically relevant in this context when compared to the non-adapted strain IMX2300 (Bioproject accession number: PRJNA634013). When both strains were compared to the reference CEN.PK113-7D sequence [45], a total of 16 mutations were found including eight non-synonymous and eight synonymous, with most mutations occurring in either Y’ helicases or Ty elements (Supplementary File 4). These elements resided in highly repetitive chromosomal regions and were therefore prone to biased variant calling when using short-read sequencing technologies. The only non-synonymous mutation found in both IMX2300 and IMX2300-1 involved a leucine to methionine change in amino acid 315 of an Mtm1 homolog predicted to be a high affinity pyridoxal 5’-phosphate (PLP) transporter, involved in delivery of the PLP cofactor to mitochondrial enzymes. Overall, these observations indicate that the lag phase of strain IMX2300 most likely reflected a physiological adaptation or culture heterogeneity rather than a mutational event [49].

The specific growth rate of *S. cerevisiae* IMX2305 (*fms1Δ TcPAND*) on SMDΔpan did not significantly differ from that of the reference strain IMX585 on SMD, and it was almost four-fold higher than the specific growth rate of the reference strain on SMDΔpan. These results are consistent with a previous study on functional expression of *TcPAND* in *S. cerevisiae* [50].

### Expression of Neocallimastigomycete *PfnadB, PfnadA*, and *NcadcA* suffice to enable anaerobic pyridine-nucleotide and CoA synthesis in yeast

To investigate whether expression of heterologous *PfnadB, PfnadA*, and *NcadcA* was sufficient to enable anaerobic growth in the absence of nicotinate and pantothenate, respectively, growth of the engineered *S. cerevisiae* strains on SMD, SMDΔnic and/or SMDΔpan was monitored in an anaerobic chamber (Fig. 3).

Growth experiments on SMDΔnic or SMDΔpan were preceded by a cultivation cycle on the same medium, supplemented with 50 g L^-1^ instead of 20 g L^-1^ of glucose to ensure complete depletion of any surplus cellular contents of pyridine nucleotides, CoA, or relevant intermediates. Indeed, upon a subsequent transfer to SMDΔnic or SMDΔpan, the reference strain IMX585 (*BNA2 FMS1*), expressing the native oxygen-dependent pathways for nicotinate and β-alanine synthesis, showed no growth (Fig. 3 panels A, B and C).

Both engineered strains IMX2301 (*bna2Δ PfnadB PfnadA*) and IMX2302 (*bna2Δ AtNADB AtNADA*) grew anaerobically on SMDΔnic. This provided a marked contrast with the aerobic growth studies on this medium, in which strain IMX2302 did not grow. Strains IMX2305 (*fms1Δ TcPAND*) and the aerobically pre-adapted IMX2300-1 (*fms1Δ NcadcA*) both grew on SMDΔpan under anaerobic conditions (Fig. 3 panels D, E and F).

### Characterization of engineered yeast strains in anaerobic batch bioreactors

The anaerobic chamber experiments did not allow quantitative analysis of growth and product formation. Therefore, growth of the *S. cerevisiae* strains expressing the Neocallimastigomycetes genes, IMX2301 (*bna2Δ PfnadB PfnadA*) and IMX2300-1 (*fms1Δ NcadcA*) was studied in anaerobic bioreactor batch cultures on SMDΔnic or SMDΔpan and compared with growth of *S. cerevisiae* IMX585 (*BNA2 FMS1*) on the same media. The reference strain IMX585, which tipically grows fast and exponentially in anaerobic bioreactors when using complete SMD [51], exhibited extremely slow, linear growth on SMDΔnic and SMDΔpan (Fig. 4). Similar growth kinetics in ‘anaerobic’ bioreactor cultures of *S. cerevisiae* on synthetic medium lacking the anaerobic growth factors Tween 80 and ergosterol were previously attributed to slow leakage of oxygen into laboratory bioreactors [27, 52, 53].

**Fig. 4:**
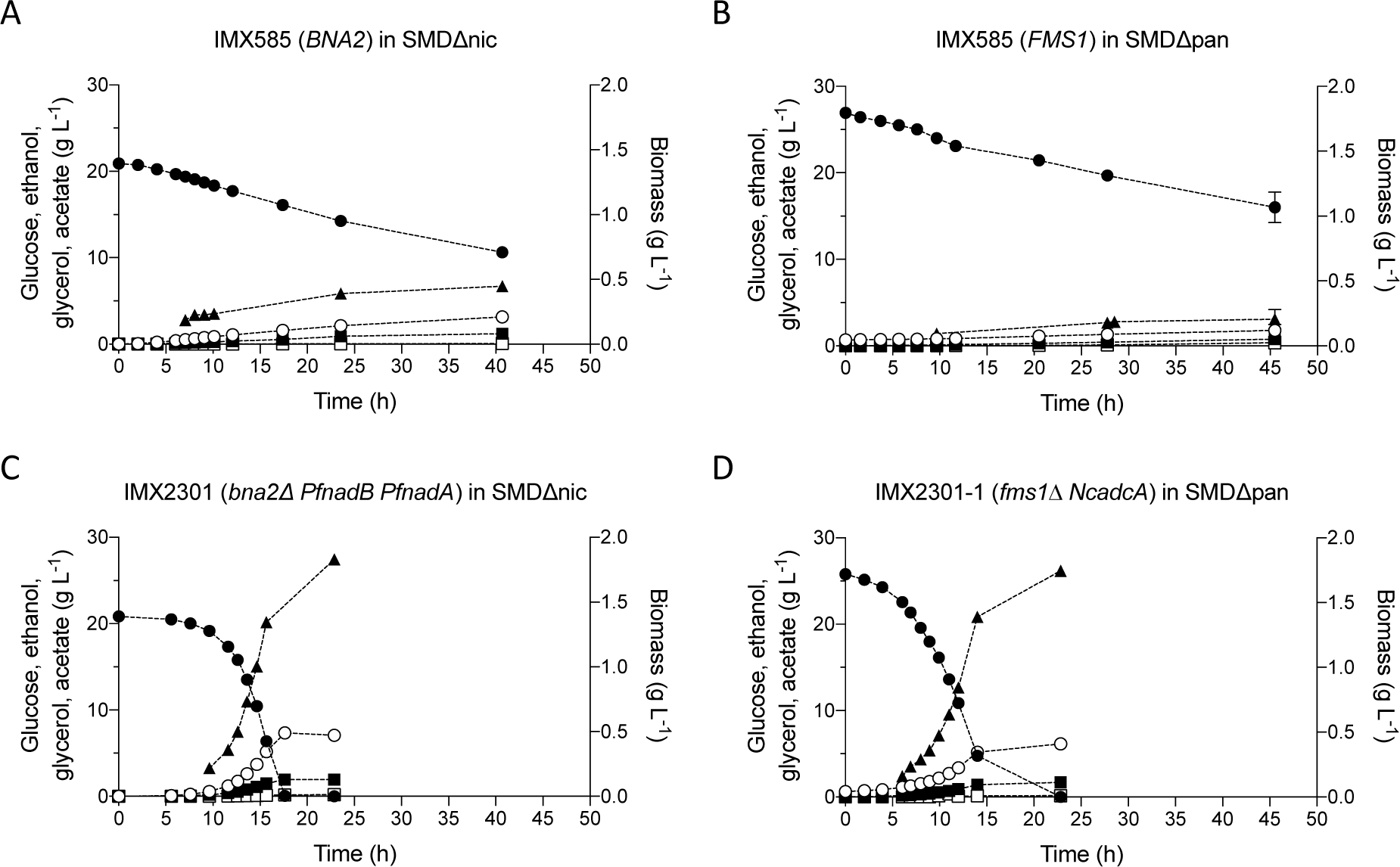
Anaerobic batch cultivation of IMX585 in SMDΔnic (A) and SMDΔpan (B), IMX2301 in SMDΔnic (C) and IMX2300-1 in SMDΔpan (D). All strains were pre-grown in the corresponding medium lacking one vitamin prior to inoculation in the bioreactor to avoid carry-over effects. Values for glucose (•), ethanol (○), glycerol (▪), acetate (□) and biomass (▲) are shown over time. Error bars represent the mean deviation of independent cultures (n=2)

In contrast to the reference strain IMX585, the engineered strains IMX2301 and IMX2300-1 exhibited exponential anaerobic growth on SMDΔnic and SMDΔpan, respectively (Fig. 4; Table 2). The specific growth rate of strain IMX2301 (*bna2Δ PfnadB PfnadA*) on SMDΔnic was not significantly different from that of the reference strain on complete SMD [51], indicating full complementation of the anaerobic nicotinate auxotrophy of *S. cerevisiae*. The specific growth rate of strain IMX2300-1 (*fms1Δ NcadcA*) on SMDΔpan was only 20 % lower than this benchmark (Table 2). Biomass and ethanol yields of strain IMX2301 grown in anaerobic batch cultures on SMDΔnic and strain IMX2300-1 grown on SMDΔpan were not significantly different from those of the reference strain IMX585 grown on complete SMD (*p*-value > 0.05, Table 2).

## Discussion

This study shows how expression of *PfnadB, PfnadA*, and *NcadcA* genes from Neocallimastigomycetes, as well as corresponding orthologs from other species (*AtNADB, AtNADA*, and *TcPAND*), confer oxygen-independent nicotinate and panthotenate prototrophy to the facultatively anaerobic yeast *S. cerevisiae*. These results also provide insights into how acquisition of these genes by HGT conferred selective advantage to Neocallimastigomycete ancestors under anaerobic conditions.

Genomic analyses previously suggested that genomes of Neocallimastigomycetes encode a putative L-aspartate oxidase (NadB) and quinolinate synthase (NadA) as alternatives to the canonical kynurenine pathway found in other fungi [6]. However, functionality of these Neocallimastigomycete proteins in an oxygen-independent pathway for synthesis of quinolinate from L-aspartate [19, 20] had not been demonstrated until now. Neocallimastigomycetes appear to have acquired *nadB* from a proteobacterium while a eukaryotic donor was implicated in the acquisition of *nadA* [6]. Our results demonstrate that expression of *nadB* and *nadA* homologs, either from the Neocallimastigomycete *P. finnis* or from the plant *A. thaliana* [21], suffice to allow anaerobic synthesis of NAD^+^ of *S. cerevisiae*. Due to the involvement of the Bna2 and Bna4 oxygenases in NAD^+^ synthesis by *S. cerevisiae*, nicotinate is an essential growth factor for this yeast under anaerobic conditions [14, 54, 55]. The present study represents the first demonstration of a metabolic engineering strategy to eliminate oxygen requirements for NAD^+^ synthesis in this yeast. A similar strategy was successfully applied to enable oxygen-independent synthesis of pyridine nucleotides in the bacterium *Pseudomonas putida* [56].

Functional expression of heterologous NadA quinolinate synthases in *S. cerevisiae* was observed despite the fact that these enzymes are 4Fe-4S iron-sulfur cluster proteins [57, 58], which are notoriously difficult to functionally express in the yeast cytosol [59-62]. However, earlier studies on functional expression of the 4Fe-4S activating protein of bacterial pyruvate-formate lyase [63, 64] demonstrated that low-levels of expression can occur without modification of the yeast machinery for cytosolic assembly of Fe-S clusters. The inability of *At*NadB and *At*NadA to support NAD^+^ synthesis in aerobic cultures may be due to oxygen-sensitivity of the 4Fe-4S cluster in the *At*NadA quinolinate synthase domain [65]. In contrast to *Pf*NadA, *At*NadA carries an N-terminal SufE domain which, in other organisms, has been demonstrated to allow this oxygen sensitive enzyme to remain active under aerobic conditions by reconstituting its Fe-S cluster [65].

Whereas an alternative to the kynurnine pathway for NAD^+^ synthesis was previously inferred from genome sequence analysis, the pathway by which Neocallimastigomycetes synthesize Coenzyme A had not previously been explored. Six pathways for synthesis of the essential CoA precursor β-alanine are known: (1) decarboxylation of L-aspartate [66], (2) transamination of malonate semialdehyde with L-glutamate as aminodonor [67] or L-alanine [68], (3) by reduction of uracil followed by hydrolysis of the resulting dihydrouracil [69], (4) oxidative cleavage of spermine to 3-aminopropanal followed by oxidation of the aldehyde group [16], (5) 2,3-aminomutase of alanine [70], and (6) addition of ammonia to acryloyl-CoA, followed by hydrolysis of the resulting CoA thioester [70]. Of these pathways, the options (1), (2), (3), (5), and (6) can, in principle, occur in the absence of oxygen. Yeasts and other filamentous fungi typically form β-alanine from spermine (pathway 4), but in some species the use of pathway 3 was also reported [71]. While the aspartate decarboxylation route has not previously been demonstrated in wild-type fungi, functional expression of bacterial and *T. castaneum Tc*PanD was used in metabolic engineering of *S. cerevisiae* to boost supply of β-alanine as a precursor for 3-hydroxypropionate production [22, 50]. Phylogenetic analysis of putative members of the pyridoxal-dependent L-aspartate decarboxylase family encoded by genomes of Neocallimastigomycetes allowed for identification of *NcadcA* which complemented a pantothenate-auxotrophic mutant of *S. cerevisiae*.

Amino acid sequence analysis of the characterised *Nc*AdcA (A0A1Y1ZL74) yielded the highest homology with sequences from chytrid fungi and *Clostridium* bacteria. This observation is in agreement with previous research showing that HGT events played a major role in shaping the genomes of Neocallimastigomycota [4, 6, 7], with Clostridiales as important sequence donors [6]. Phylogenetic analysis of Adc sequences (Fig. 2) are consistent with an earlier report on multiple evolutionary origins and variable evolutionary rates of pyridoxal-5’-phosphate-dependent enzymes, including Adcs and glutamate decarboxylases [72, 73]. A separate clade of Neocallimastigomycete sequences show homology with characterised glutamate decarboxylases (e.g. Q04792 from *S. cerevisiae* and K4HXK6 from *Lactobacillus brevis*; Fig 2. These results further support acquisition of an Adc encoding DNA sequence by HGT rather than by neofunctionalization of a glutamate decarboxylase gene.

Wild-type *S. cerevisiae* strains cannot grow in anaerobic environments unless supplemented with pantothenate. Expression of either *NcadcA* or *TcPAND* in an *fms1Δ S. cerevisiae* strain, which lacks the native oxygen-dependent pantothenate biosynthesis pathway, enabled growth in panthothenate-free medium under aerobic and anaerobic conditions. Although the different specific growth rates of *S. cerevisiae* strains expressing *NcadcA* or *TcPAND* indicate that changing expression levels and/or origin of ADC encoding genes may be required to achieve optimal growth, these results provide a proof-of-principle for a simple metabolic engineering strategy to eliminate oxygen requirements for pantothenate synthesis.

This work contributes to the understanding of how Neocallimastigomycetes adapted to their anaerobic lifestyle by acquiring genes that enable oxygen-independent synthesis of central metabolic cofactors. Experiments with engineered *S. cerevisiae* strains showed that contribution of the heterologous genes to *in vivo* oxygen-independent cofactor synthesis did not require additional mutations in the host genome. These results indicate how acquisition of functional genes by HGT, even if their expression was initially suboptimal, could have conferred an immediate advantage to ancestors of anaerobic fungi living in cofactor-limited anoxic environments. A similar approach was recently applied to study the physiological impact on *S. cerevisiae* of expressing a heterologous gene encoding squalene-tetrahymanol cyclase, which in Neocallimastigomycetes produces the sterol surrogate tetrahymanol [13]. Functional analysis by heterologous expression in *S. cerevisiae* circumvents the current lack of tools for genetic modification of Neocallimastigomycetes [2], and can complement biochemical studies [3-5] and genome sequence analyses [6, 7].

Pantothenate and nicotinate, together with the other compounds belonging to the B-group of water-soluble vitamins, are standard ingredients of chemically defined media for aerobic and anaerobic cultivation of yeasts [48]. *S. cerevisiae* strains have been shown to contain the genetic information required for *de novo* synthesis of these vitamins and, can even be experimentally evolved for complete prototrophy for individual vitamins by prolonged cultivation in single-vitamin depleted media [74, 75]. In large-scale processes, addition of nutritional supplements increases costs, reduces shelf-life of media and increases the risk of contamination during their storage [48]. Therefore, metabolic engineering strategies for enabling oxygen-independent synthesis of NAD^+^ and pantothenate are of particular interest for the development robust yeast strains with minimal nutritional requirements that can be applied in anaerobic biofuels production [48]. Further studies of the unique evolutionary adaptations of Neocallimastigomycetes may well provide additional inspiration for engineering robust fungal cell factories that operate under anaerobic conditions.

## Acknowledgments

We thank Dr. Irina Borodina for providing us the codon-optimized *TcPAND* gene and Sabina Shrestha for constructing strain IMK877. TP and J-MD were supported by the European Union’s Horizon 2020 research and innovation programme under the Marie Sklodowska-Curie action PAcMEN (grant agreement No 722287). AMV, WJCD, JB, SJW, RAO-M, CM and JTP were funded by an Advanced Grant of the European Research Council to JTP (grant # 694633). Authors declare that they have no conflict of interests.

## Author contributions

All authors contributed to the experimental design. TP, AMV, JMD and JTP wrote a first version of the manuscript. All authors critically read this version, provided input and approved the final version. RAO-M, AMV, and TP performed the phylogenetic analysis. TP constructed the *S. cerevisiae* strains and performed the aerobic characterization. AMV, WJCD and TP performed the anaerobic chamber experiments. JB, CM, TP, AMV, and SJW performed and analysed the bioreactor experiments.

## Notes

### Competing Interest Statement

The authors have declared no competing interest.

