## Supplementary material for "Identification of oxygen-independent pathways for pyridine-nucleotide and Coenzyme-A synthesis in anaerobic fungi by expression of candidate genes in yeast"

**Supplementary Fig. S1: Verification of L-aspartate decarboxylases from *Neocallimastix californiae* using RNAseq data.** Illumina libraries were obtained from the Sequence Read Archive using the SRR7140690 run identifier [2]. Reads were mapped using STAR 2.6.1a_08-27 [3] against genome assembly GCA_002104975. Alignments were processed using samtools 1.3.1 [4] and visualized using Artemis [5]. Errors are highlighted inside black dashed lines. **A)** A0A1Y2ADH1, **B)** A0A1Y2B2H7, **C)** A0A1Y2AA19, **D)** A0A1Y1ZL74 ........ 6

**Supplementary File 4: Variant calling results.** Provided as tab-delimited text file showing non-synonymous (NSY) and synonymous (SYN) mutations found after whole-genome sequencing analysis of strains IMX2300 and IMX2300-1. First column shows GenBank identifiers for the corresponding *Saccharomyces cerevisiae* CEN.PK113-7D chromosome. Links are provided to the amino acid sequence deposited in GenBank and to the *Saccharomyces* Genome Database for CEN.PK113-7D proteins with an *S. cerevisiae* S288C. ........ 53

References ........ 55

### Supplementary tables

### Supplementary Table S1: Oligonucleotide primers used in this study

| Primer number | Primer sequence | Product(s) |
|---|---|---|
| 6005 | GATCATTTATCTTTCACTGCGGAGAAG | gRNA pROS plasmid backbone amplification |
| 11861 | TGCGCATGTTTCGGCGTTCGAAACTTCTCCGCAGTGAAAGATAAATGATCCAGAAGAGCATATTCCATTTGTTTTAGAGCTAGAAATAGCAAGTTAAAATAAG | 2μm fragment for *BNA2* gRNA plasmid |
| 11862 | GTCAACGCCGATATGAACAACACTTCCATAACCGGACCACAAGTACTACATAGAACAAAACATTATACATTTTATTAACGCCCCCCCTTTTTTTTTTTTGTTTGATGCAGAAGCCTCGCA | *BNA2* KO repair oligo fwd |
| 11863 | TGCGAGGCTTCTGCATCAAACAAAAAAAAAAAAGGGGGGGCGTTAATAAAATGTATAATGTTTTGTTCTATGTAGTACTTGTGGTCCGGTTATGGAAGTGTTGTTCATATCGGCGTTGAC | *BNA2* KO repair oligo rev |
| 11877 | GCATCGTCTCATCGGTCTCATGCATCGTCTCATCGGTCTCAT | YTKflank_TcPanD_fwd |
| 11878 | ATGCCGTCTCAGGTCTCAGGATTCACAAATCGGAACCCAAT | YTKflank_TcPanD_rev |
| 16721 | CAATTCGTCGCAATACAACGCAGTTCGAGTTTATCATTATCAATACTGC | *pTDH3* amplification_fwd |
| 16722 | ATGGTTTCTTTGTCGACCATTTTGTTTGTTTATGTGTGTTTATTCGA | *pTDH3* amplification_rev |
| 16723 | AACACACATAAACAAACAAAATGGTCGACAAAGAAACCATTAA | *NcPanD* amplification_fwd |
| 16724 | AATTCTTAGTTAAAAGCACTTTACTTGATCAGCTTGTGGTTCA | *NcPanD* amplification_rev |
| 16725 | ACCACAAGCTGATCAAGTAAAGTGCTTTTAACTAAGAATTATTAGTCTTTTCTG | *tENO2* amplification_fwd |
| 16726 | CTGACGAGCAGATTTCCAGCATTTTTCAAACTGCAAATTCAAGAA | *tENO2* amplification_rev |
| 16727 | GAATTTGCAGTTTGAAAAATGCTGGAAATCTGCTCGTCAG | pYTK096 amplification_fwd |
| 16728 | ATAATGATAAACTCGAACTGCGTTGTATTGCGACGAATTG | pYTK096 amplification_rev |
| 13527 | AACAAGAAGTGAGTTAATAAAGGCAAAAACAGTGGTCGTGTGAGAAGTAGAATTTCACCTAGACGTGGAATCTATTTTTTCGAAATTACTTACACTTTTGACGGCTAGAAAAG | *FMS1* KO repair oligo fwd |
| 13528 | CTTTTCTAGCCGTCAAAAGTGTAAGTAATTTCGAAAAAATAGATTCCACGTCTAGGTGAAATTCTACTTCTCACACGACCACTGTTTTTGCCTTTATTAACTCACTTCTTGTT | *FMS1* KO repair oligo rev |
| 13123 | TTTACAATATAGTGATAATCGTGGACTAGAGCAAGATTTCAAATAAGTAACAGCAGCAAACAGTTCGAGTTTATCATTATCAATACTG | *NadA* repair fragment fwd |
| 13124 | ATAGCATAGGTGCAAGGCTCTCGCCGCTTGTCGAGCTATTGGCATGGATGTGCTCCCTAAATACATGGGTGACCAAAAGAGC | *NadA* repair fragment rev |
| 13125 | TTAGGGAGCACATCCATGCCAATAGCTCGACAAGCGGCGAGAGCCTTGCACCTATGCTATCACCCATGAACCACACGG | *NadB* repair fragment fwd |
| 10710 | TATATTTGATGTAAATATCTAGGAAATACACTTGTGTATACTTCTCGCTTTTCTTTTATTATTTTTCAAACTGCAAATTCAAGAAAAAGCCAC | *NadB* repair fragment rev |

Supplementary Table S2: List of DNA parts combined in single BsaI Golden Gate reactions to obtain the transcriptional unit plasmids

| **Plasmid** | **Promoter** | **Gene** | **Terminator** | **Backbone** |
|---|---|---|---|---|
| pUDI168 | pYTK017 | *TcPAND* PCR | pYTK056 | pYTK096 |
| pUDI245 | pYTK009 | pUD1096 | pYTK051 | pYTK096 |
| pUDE931 | pYTK010 | pUD1097 | pYTK055 | pGGKd017 |
| pUDI243 | pYTK009 | pUD652 | pYTK051 | pYTK096 |
| pUDI244 | pYTK010 | pUD653 | pYTK055 | pYTK096 |

**Supplementary Table S3: Neocallimastigomycete BLASTP hits (e-value < $10^{-6}$).** Obtained using the dataset from Tomita et al. [1] as queries against a Neocallimastigomycete-specific amino acid sequence database

| Entry | Entry name | Status | Protein names | Gene names | Organism | Length | Pfam |
|---|---|---|---|---|---|---|---|
| A0A1Y1VIL9 | A0A1Y1VIL9_9FUNG | unreviewed | PLP-dependent transferase | BCR36DRAFT_345242 | *Piromyces finnis* | 600 | PF00282 |
| A0A1Y1WT12 | A0A1Y1WT12_9FUNG | unreviewed | Glutamate decarboxylase (EC 4.1.1.15) | BCR32DRAFT_296089 | *Anaeromyces robustus* | 797 | PF00282 |
| A0A1Y1XDI3 | A0A1Y1XDI3_9FUNG | unreviewed | PLP-dependent transferase | BCR32DRAFT_266694 | *Anaeromyces robustus* | 594 | PF00282 |
| A0A1Y1ZL74 | A0A1Y1ZL74_9FUNG | unreviewed | PLP-dependent transferase | LY90DRAFT_708822 | *Neocallimastix californiae* | 625 | PF00282 |
| A0A1Y2AA19 | A0A1Y2AA19_9FUNG | unreviewed | PLP-dependent transferase (Fragment) | LY90DRAFT_677094 | Neocallimastix *californiae* | 433 | PF00282 |
| A0A1Y2ADH1 | A0A1Y2ADH1_9FUNG | unreviewed | PLP-dependent transferase | LY90DRAFT_463975 | *Neocallimastix californiae* | 577 | PF00282 |
| A0A1Y2AZ59 | A0A1Y2AZ59_9FUNG | unreviewed | Glutamate decarboxylase (EC 4.1.1.15) | LY90DRAFT_628503 | *Neocallimastix californiae* | 481 | PF00282 |
| A0A1Y2B2H7 | A0A1Y2B2H7_9FUNG | unreviewed | PLP-dependent transferase | LY90DRAFT_460911 | *Neocallimastix californiae* | 524 | PF00282 |
| A0A1Y3MYE4 | A0A1Y3MYE4_PIRSE | unreviewed | Uncharacterized protein | PIROE2DRAFT_17257 | *Piromyces sp.* (strain E2) | 450 | PF00282 |
| A0A1Y3NC87 | A0A1Y3NC87_PIRSE | unreviewed | Uncharacterized protein | PIROE2DRAFT_7587 | *Piromyces sp.* (strain E2) | 248 | PF00282 |
| A0A1Y3NC92 | A0A1Y3NC92_PIRSE | unreviewed | Uncharacterized protein | PIROE2DRAFT_7586 | *Piromyces sp.* (strain E2) | 169 | |
| A0A1Y3NFH0 | A0A1Y3NFH0_PIRSE | unreviewed | Aminotran_5 domain-containing protein | PIROE2DRAFT_61071 | *Piromyces sp.* (strain E2) | 196 | PF00266 |
| A0A1Y3NIR5 | A0A1Y3NIR5_PIRSE | unreviewed | Uncharacterized protein | PIROE2DRAFT_58808 | *Piromyces sp.* (strain E2) | 127 | PF00282 |

### Supplementary figures

**Supplementary Fig. S1: Verification of L-aspartate decarboxylases from *Neocallimastix californiae* using RNAseq data.** Illumina libraries were obtained from the Sequence Read Archive using the SRR7140690 run identifier [2]. Reads were mapped using STAR 2.6.1a_08-27 [3] against genome assembly GCA_002104975. Alignments were processed using samtools 1.3.1 [4] and visualized using Artemis [5]. Errors are highlighted inside black dashed lines. **A)** A0A1Y2ADH1, **B)** A0A1Y2B2H7, **C)** A0A1Y2AA19, **D)** A0A1Y1ZL74

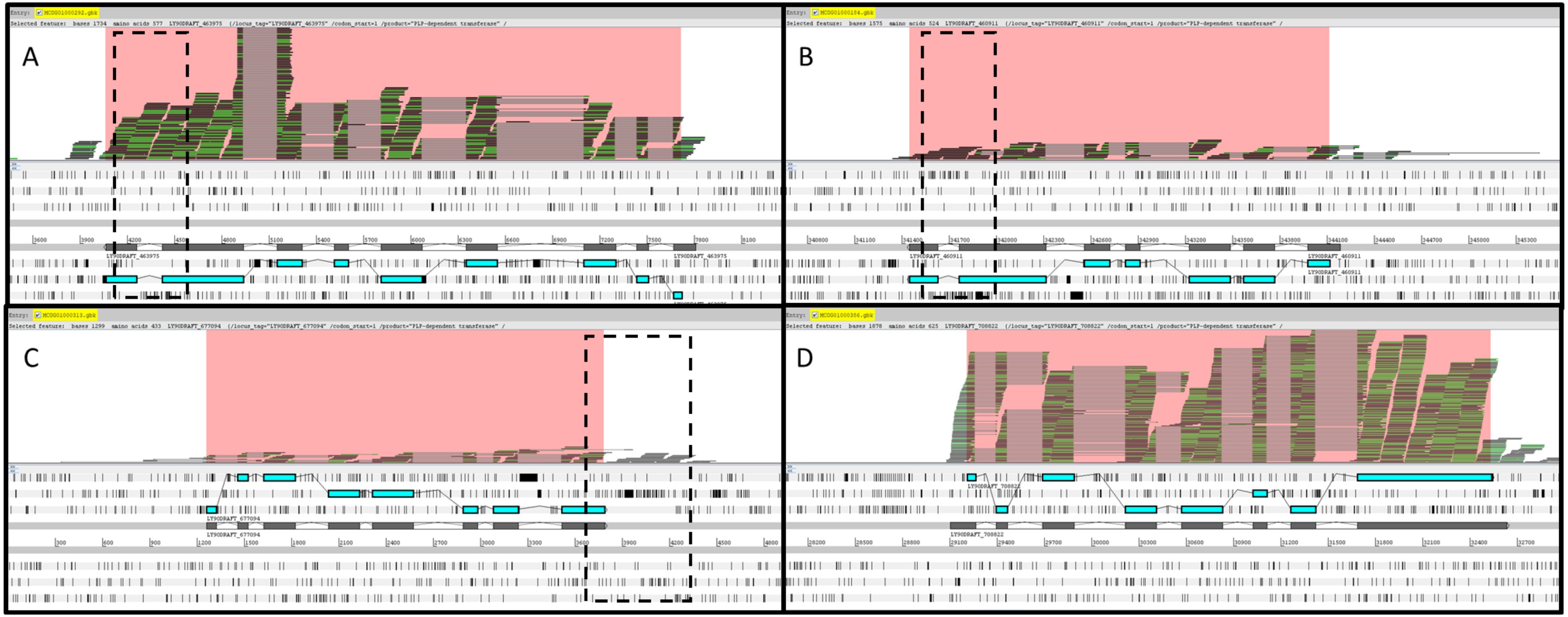

### Supplementary Files

#### **Supplementary File 1: 387 Amino acid sequences used for the phylogenetic analysis shown in Fig. 2.** File provided in FASTA format with UNIPROT identifiers in the headers. Subscripts were added to the headers as follows: A_ for sequences from Archaea, B_ from Bacteria, E_ from Eukaryota, N_ from Neocallimastigomycetes, and T_ from Tomita et al. [1].

```
>B_A8FX14
MTARQAKASEEALLRIFTIPEAPGSTLSVIEQNISQNLMGFLQESVVAVEKPLTEIERDFQEHQIPAAPKFVSDYADEMMKTLVAHSVHTSAPS
FIGHMTSALPYFVLPLSKMMVGLNQNLVKIETSKAFTPLERQVLGMMHHLIYDENETFYNSWMHSANVSLGAFCSGGTVANITALWTARNQLLK
ADGDFKGIAAQGLMKGLRHYGYNDLAILVSERGHYSLGKTADLLGIGRENIIQIPTSNDNRVDVDKMRVTAKALERDNIKVMAIVGVAGTTETG
NIDPLDKLATLAEELDCHFHVDAAWGGASLLSKKYRHLLKGIERADSVTIDAHKQMYVPMGAGMVIFKDPTFANAIKHHAEYILRKGSKDLGSQ
TLEGSRPGMAMLVHACLQIIGRDGYEILINNSLEKARYFAELIKTTDNFELVSEPELCLLTYRYVPESVQKAMQQARTDGDIERLLQFNRLLDG
LTKFVQKRQREQGTSFVSRTRINPEHCHDIDVDLKSVVFRVVLANPLTTNEILQQVLVEQTQIASTDKKFLPQLLELAAH
>B_A3WL97
MWTGKHMNDVDDKSYAEVNQESLYRIFTLPEAPDSTLSRIEKDLSENLLGFLGDHIVAKEKPLQDIERDFEKASIPENPQFVSDHTEFLLEKLV
AHSVHTASPKFIGHMTSALPYFLMPLAKLMVGLNQNLVKIETSKAFTPLERNVIGMLHHLVYQRESSFYNRWMHSAQHSLGAFCSGGTVANMTA
LWVARNQLLKPDGDFAGIAAEGLAAGLAHYNYKRLVVLVSERGHYSLSKAADVLGIGRKNLVAVATDANNRIKIDALRQACQDVREQGGEVLSI
VGVAGTTETGHIDPLADMADIAEQLGIHFHVDAAWGGATLLSNDYRHLLSGIERADSVTIDAHKQMYVPMGAGLVLFKDPAMVKAIEHHAEYII
RQGSKDLGSHTMEGSRPGMAMMVFSAMHVIGRRGYELLINNSLEKARYFSSLIAEQDDFEVITEPELCLLTYRFVPKRVKLALEAANVEQVERL
TPHLNALTRYIQKRQRESGRSFVSRTKLTPEHYYRVPTVVFRVVLANPLTTHDMLREIIAEQREIAEKATSLRGALHTEMRELGLIRDA
>B_A0A1I2AY29
MSERLYEKWFLGDSNQSLEQYYDFMNQAAETLETYFATGKSSFSGNTPSQVRAQLMTNFIDMGPEKGQNTETLFKHIGESILQNAVHVHHPACV
AHLQCPPLIPAMAAEVLIAGMNQSLDSWDQSGAATIVEQEVVSWLCDLYQLPNGDGVFTTGGTQSNYMGLLLAREHISERLWNWNPRQKGMHPE
SHRLRILCSEEAHFTVQQSAAFLGLGEQSVVSISTDENYRLSITELEQKLDGLFQKDMLPFALFATAGTTDFGSIDPLPQMAELAHEYGLWFHV
DAAYGGAVQLSEMYKGKIDGIQEADSITIDFHKQFYQPISSGVFLLKDQKNFQHLKLNADYLNPEEDEAEGIPNLVGKSVQTTRRFDALKLFLS
LQSLGVNRLGSMVDTTIDLARQVAQLLVEDPYIQIEHEPELNAIVFRFSHAQVDDKGIDEINMKIHDELLRTGKAVIARTKVNGKAYLKLTLLN
PNTTIDDIETLLQELKCIAFNLLEQQEGLLV
>B_U4K3X4
MVSENKTADVSFESLLKIFTVPEAPDSTLGHIENKLSQNLNQFLREHIVAEEKPLADIEQQFSNPYLPEQPQFVSEHTQNLLDNLVSQSVHTSA
PSFIGHMTSALPYFLMPLSKIMIALNQNLVKIETSKAFTPLERQVLGMLHGLIYHEKPEFYQQWMHSANHSLGAFCSGGTIANITALWVARNNT
LKADGEFKGVEKEGLFKAMKHYGYEGLAILVSERGHYSLKKAADVLGLGQECLVSVKTDSKNKLCPDDLEQKIALLKSQNIKPFAIVGVAGTTE
TGNIDPLERIAVISQKHGCHFHVDAAWGGATLMSNNYRHLLKGIELADSVTIDAHKQLYIPMGAGMVLFKDPSAMTAIEHHAQYILRKGSKDLG
SHTLEGSRSGMAMLVYSSMHIISRPGYELLIDQSIEKAKYFASLVSEQDDFELVTEPELCLLTYRYVPEVAKKALVNANEDEKGALHEALNNLT
KQIQKKQRETGKTFVSRTRLNPVQWQHRNTIVFRVVLANPLTTHEILQSVLCEQREIAAGFPNLMTQVSELSMEILNKKAS
>B_U4K047
MVYFTSNSQEISSETAARADRASKVEDFFNGKADLTALAAIEETLQHDYEYFLCKLSDVAVSGQSTDFEVPESPLTTEAFCNFLQGLLTCIAPV
FSPEFVGHMTSALPKHAIMMSKILAATNQNVVKVETSNQFTSLEKSIINYFHKLCYQSDYSESSIPSDLSLGNFCSGGTVANLSALWVARNRAL
GANVGEVGLLKALKGCPYDDLAIVVSESGHYSLKKAADLLGIGKENLIAVKTDSHHKIDIADLRKTLSRLRDDNIGVLAIVGVAGATETGVIDP
LNEMADLAREYQCHFHVDAAWGGASLLSDQHRELFSGIEKADSVTIDAHKQLYVPMGAGMVLFKDPHAVQSISHHANYIIRKGSDDLGAHTIEG
SRGAVAAMVAANLALLGKKGMSQLIDNSIEKAKALATHIATLTDFELTSEPTLCILTYRYAPDSLMSSLNLLDAESRYNALEMLDELNKEIQET
QRDTGIGFVSRTKIRVNKEEQKRVVFRVVLANPLTQMSHLTAILDEQIRIASQSEGLTKLNQYIQHSLKRSA
>B_I5B6M9
MSSSFSFFSLPGNSTWTQGDKNIRKTDGQLIADRKSLDRVFIRPYDENSKKTLVKYMEQILFGLHDFLNRHVGVTEEISITELAKNYMDVQISD
HPQKNLGQVIEDIIKDIAPKAVNVASPYFIGHMTSALPFFMVHLKAITAALNQNLIKMETSKVLAVIERQVLAKIHCLIFKQSQAFYRAHVQNT
RTALGAFTSGGTTANITAMWVARNKLFPPKDDFSTIEEDGLFKAMSVHHTDRVVILVSRRAHYSLRKASGILGLGNKNIIAVDVKPDHTIDIDK
LKQTIKDLKQEGRTKIAAIVGIAGATETGTIDPLQQMADICEQEQIHFHVDAAWGGPILLSHTYAHLLSGIERADSVTIDGHKQFYMPMGAGMV
YFKNANALDAIAYHARYVNRKGSVDLGIKTLEGSREAACLILDASLKIMGAKGYALMIDHGIETARAFAEKIEERPEFELVTRPVLNILTYRLV
PMSFRQKLATARGEERKRLNQELDEINIRIQRIQREAGKSFVSRTRLKLVPEDDFMVVVLRSVIMNPYTTQAILDDILDEQEQIYHKF
>B_Q07ZT1
MTSKITRQANASEDSLLRIFTAPEDSTSTLSIIEQKLSQDLAGFLGNSIAALEKPLSEIETDFSAYQIPTEPKFVSDYADEIMQNLVAHSVHTS
APSFIGHMTSALPYFVLPLSKMMVGLNQNLVKIETSKAFTPLERQVLGMMHNLVYQQSDDFYHQWMHSANHSLGAFCSGGTVANITALWIARNR
LLQPRGDFKGVTREGMTKALRFYGYDDLAILVSDRGHYSLGKTADLLGIGRDNIISIPTDEHHKVDVAAMRDAAEKLAQQNIKVMAIVGIAGTT
ETGNVDPLTELAALASELNCHFHVDAAWGGASLLSKKYSHLLKGIELADSVTIDAHKQMYVPMGAGMVLFKDPELAKAIVHHAEYILRVGSKDL
GSQTLEGSRPGMAMLVHACLQIIGRDGYEILINNSLEKARYFGSIIDQHADFELVTAPELCLLTYRYVPKSVQAILTTALANQDSALVASINEL
LDGLTQFIQKHQREQGKSFVSRTRIKPAKYLRQPTTVFRVVLANPLTSHQILHDVLTEQVEIAALDSEYLPQLLAL
>B_F0RIL5
MKKLVEAYSPEDFRKRGHDLINELADHFENTLQNKNNKTIHWNQPKDEHLFWKDYLENGNEENLFSTIISHSIHVHNPKYIGHQVTPTLPITAL
SALVSASLNNGMAVYEMGVAASAIERIVTDYICKKAGYTQAANGILTSGGTLANLTALLTARKAKVPTNVWTDGSTNSLGIMVSEEAHYCVDRA
ARIMGLGDAGIIKIPSTKNFNMNITLLEEYYQKAKDKGIEVFAIIGSAPSTSTGIYDDLEAIATFSKQKNIWFHVDGAHGGAAIFSKTHKHTVK
GIEKADSIVIDGHKMMMMPALTTALLFKDGNTSHATFSQKADYLLEESKEEDWYNIAKRTFECTKTMASIHWYTILKTYGEEIFDEYVTTLYNL
GLQFAEMIIANPNFEIAVQPASNIVCFRYVNPNLSEKETNSLNEKIRQSILEKGEFYIVQTKLRGIHYLRVTIMNPFTTKTHLQQLLERVLEEI
N
>B_Q1LR80
MKARKAGRSDNAREATGNDAEMGEGCPLHWFDADLGAFDDLERRVAEHPADFFAGQDFDPVGACATREASFASVQIPEDPTAPQQHADHLLNDV
FRHVMPVASPTFVGHMTSSLPSFMPSLAKIVAALNQNVVKLETSGALTGLERQVIGMLHHLVFARDDAFYAQWLHNAEHSLGAFCSGGTTANLT
ALWASRNNTLRARDGFAGINQAGLVAALRHYGYAGLAIVVSERGHYSLRKAADILGIGRENLVPVGVDEDGRMRVDLLRDTLRDLQQRNIRTVA
IVGIAGTTETGAVDPLDAIADVAQEIGCHFHVDAAWGGATLLSARERARFAGIERADSVVIDAHKQFYVPMGAGMVLFRDPAWTQDIIQHANYI
VRKGSVDLGRHTLEGSRGSAAVMLYANLHLLGRKGLAKLIDTGIDNAKYFASLIEQQPDFELDSRPQLCILTYRYVPAPVRAALLSASPEQREQ
LLEALDALTINIQEMQRDAGRSFVSRTQLTSSQWGGRPIAVFRVVLANPDTTHEILQSLLEEQRGLAKQSPMLPALMAMVGG
>B_A0A0F4QH65
MGPKRCAVASEESLMRIFTVPEAPSSTLSVIEQEISSNLAGFLNENIAAIEKPLHEIEKDFQSAMIPEEPMFVSDYAQDIMEQLVAHSVHTASP
SFIGHMTSALPHFVLPLSKLMVGLNQNLVKIETSKAFTPLERQVLGMMHHLAYGENDAFYEKWMHSAKTALGAFCSGGTIANISALWIARNRLL
GPDGDFKGLASEGIMAAMLHYGYKGLAVLVSERGHYSLGKAADVLGIGRSNFVAVKTDDNNKVSVEAMRAKAQELSERGIKVMAIVGVAGTTET
GNIDPLPEMAALAQELGCHFHVDAAWGGATLLSDNYRHLLKGIELADSITIDAHKQMYVPMGAGIVLFKDPTATDAIEHHAEYILRKGSKDLGS
```

HTLEGSRPGMAMLVHACLRVIGRKGYEMLIDKGIEKAQYFADLIHADEDFELVSEPELCLLTYRYVPKQIRQAIDRAEEQERIDIYAALNRFTA
SMQKRQRESGRSFVSRTRLTPAQYQHQPTVVFRVVLANPLTSKQMLNEILSEQKGLAQSDPVFKKYLAKYMA
>B_A0A1Y0IE65
MASRKKTANANLESMYRVFTAPEAPQSTLSQIEDYISKNLAGFLQEHIVAVERDLSEVEKDFADSGIPDKPMFVSEQAQFLLDKLVSNSVHTAA
PSFIGHMTSALPYFMLPLSKIMIALNQNLVKTETSKAFTPLERQVLGMLHRLIYQHDDAYYAQWMHDPLHALGSMCSGGTVANLTALWVARNLA
FPEEGGFKGIRREGLTRALLYYGYTGAAVLVSERGHYSLKKAADILGIGHDYIIPIPTNDQNKMDIDALRNECIRLKKDSIRILSIIGIAGTTE
TGNVDPLEDIADVAKEFNAHFHVDAAWGGPTLFSRKFSHLLKGIEKADSVTIDAHKQLYVPMGSGLVLYKNPTSVNAIEHHAQYIIRKGSRDLG
STTLEGSRPGMALLIHSGLKVIGKDGYELLINQGIEKANLFANLIEQHPDFELVTPPELNILTYRYCPAEAQAALKVASQEQTNKINQILSRVT
KSIQKTQRERGKAFVSRTRLNPAVYGRSPCVVFRVVLANPLTTPDILTSILEEQRELSAEPGIATELQHLQTLVSEVRQATT
>B_Q9KSV7
MVSEHKSAQVNFDSLLKIFTVPEGPDSTLTKIDEELSRNLNHFLRKHIVAEEKPLKEIEKDFSNAHIPEQPQFVSDHTQYLLDTLVSHSVHTAS
PSFIGHMTSALPYFLMPLSKIMIALNQNLVKIETSKAFTPLERQVLGMIHRLIYGETDHFYQQWMHSAEHSLGAFCSGGTIANITALWVARNNA
LKAEGDFPGVEKAGLFKAMRHYGHEGLAILVSERGHYSLKKAADVLGIGQEGLVAVKTDAHNRICPHDLEQKITELKANKIKVFAVVGVAGTTE
TGNIDPLRTIAQICQREQIHFHIDAAWGGATLMSNRYRGLLDGVELADSVTIDAHKQLYIPMGAGMVLFKDPNAMRSIEHHAQYILRQGSKDLG
SHTLEGSRSGMAMLVYASMHIISRPGYQLLIDQSIEKARYFADLIDAQTDFELVSQPELCLLTYRYLPEHVRMALEKSQGVQRAQLNELLNELT
KFIQKKQRETGKSFVSRTQLNPHQWDKLATIVFRVVLANPLTTKEILHNVLDEQREIAQQAPKLMRQIEHLTQCILNQ
>B_A3QG03
MTSKQTRQATASEEALMRIFTLPEAPNSTLGKIEKNLSENLMGFLKESIVAVEKPLTEIEKDFQAYQIPTAPSFVSDYAEQMMQTLIAHSVHTS
APSFIGHMTSALPYFVLPLSKMMVGLNQNLVKIETSKAFTPLERQVLGMMHHMVYGQTEEFYQSWMHSASHSLGAFCSGGTVANITALWIARNR
LLKPDGDFKGVAQSGLMRALRHYGYDDLAILVSTRGHYSLGKAADLLGIGRDNIISVPCASDNKVDVAKMREAAEQLAEQNIKVMAIVGVAGTT
ETGNIDPLDELANLAEQLGCHFHVDAAWGGASLLSSKYRHLLAGIERADSVTIDAHKQMYVPMGAGMVLFKDPEFANAIKHHAEYILRKGSKDL
GSQTLEGSRPGMAMLVHACLQIIGRDGYEILINNSLEKARYFGELIAAQDDFQLVSRPELCLLTYRYVPAKVQAQLNEAVAAGDAARVSEINAL
LDGLTKFIQKRQREQGKSFVSRTRIIPANNLEQTSVVFRVVLANPLTSNEILQQVLDEQRDIAKLDDRFLPKLLAL
>B_A0A1S2TUI8
MTASKQRQATASENSLMRIFTVPEAPESTLSRIEQALSEDLAGFLTQNIAALEKPLSEIEKDFQAFEIPHQPRFVSDYTEEMMQNLVAHSVHTA
APSFIGHMTSALPYFVLPLSKMMVGLNQNLVKIETSKAFTPLERQVLGMMHHLIYGQTTDFYHQWMHSANHALGAFCSGGTVANITALWIARNQ
LLKADGEFRGINREGLGAALCHYGWKSLAILVSERGHYSLGKAADLLGIGRDNIVSIATDANNKVDVNAMREAATKLAQQGIRVMAIVGVAGTT
ETGNIDPLPELAELATELNCHFHVDAAWGGASLLSSKYRHLLKGIELADSVTIDAHKQMYVPMGAGMVLFKQPELARAIVHHAEYILRKGSKDL
GSQTLEGSRPGMAMLVHACMQIIGKEGYEILINNSLEKARFFAGLIEQHSEFELVSAPELCLLTYRWCPSWAQTALSRQLAQGNLEKVKEINGI
LDDLTRAIQKKQREQGKSFVSRTRIQPARYHRETVTVFRVVLANPLTTDTILADVLNEQTELAASETGLLQVLKQACN
>B_A0A075P054
MGEAQVSLEHLFRVFTKPEHKDSKLAQIEQHLSDNILDFLSQHVVTKKTSLEEVEKDFANAKVPESPEFVSTHAETLLDKLVAHSVNTYSPTFI
GHMTSALPYFHLPLSKLMVGLNQNLVKIETSKAFTPLERQVLGMLHNLVYDRSEAFYEQHLHSAQHALGAFCSGGTIANITALWVARNKLLGPQ
PGFAGVAKAGLAAAYRFHNINHLGVMCSKRGHYSLSKAVDALGLGREQLIALPAPQQTLDPEKALRAGKRYQEEGNKLLAIVGVGGTTETGHVD
PLDELADVAEQLGCWFHVDAAWGGATLFSSQYRDRLKGIERADSVTIDAHKQMYVPMGAGMALFKDPENANAVRHHAQYILRAGSKDLGATTLE
GSRNGMAMMVYSALHIFGRRGYELLIDRSIQKAKAFADMIDKHPDFELTTSPTLSLLTYRVCPQALQHTLKQVDDSTRDEINEKIDTLVVSVQK
QQREAGKSFVSRTRLEAPDYPAKCITVFRVVLANPLTSHNDLAAILAEQHLLATSTPLWQSLSQLAAQDPA
>B_A0A1Y6G133
MSKTNFAEVNQEALLKIFTIPEAPDSTLGKIERHLSENLMGFLGEHIVAREKPLQEIEQNFQASSVPEQPVYVSDHTEFLLDKLVAHSVHTASP
RFIGHMTSALPYFLLPLAKLMVGLNQNLVKIETSKAFTPLERNVLGMLHHLVYGQDEAFYKRWMHSAEHSLGAMCSGGTVANITALWVARNRLL
KADGDFAGVANDGLAAGLAHYNYKRATVMVSERGHYSLSKAADVLGIGRRNLVAVATDENNRIRIDALREACEKVAAEGGKVMAIIGVAGTTET
GHIDPLDQMADVAKEIGTHFHVDAAWGGATLLSNTHRHLLKGIERADSVTIDAHKQMYVPMGAGMVLFRDPGMVRAIEHHAEYIIRHGSKDLGS
HTMEGSRSGMAMMVFSALHVMGRSGYELLIDGSMEKARYFAKLIDDQNDFEVITHPELCLLTYRFVPTRVKVALKTASAEQVRRLTPHLNALTR
FVQKRQRETGQSFVSRTKLTPAQYGREPTVVFRVVLANPLTTQDMLAEVLEEQRQIARQATSLLGALHTELRELGLTNRD
>B_A1S4V1
MPPAVHKTARRDEMTEKHLRKATASEDNLLRIFTVPEAADSTLGRIEQQLSADLAGFLQDNIAALEQPLSDIEDHFYSVEVPPQPQFVSDYVDD
IMAHLVAQSVHTAAPSFIGHMTSALPYFLLPLSKLMVGLNQNLVKIETSKAFTPMERQVLGMMHHLIYGREKDFYQGFLHSASHALGAFCSGGT
VANITALWIARNQLLKARGNFRGVTREGLHKALKHYGWDDLAILVSERGHYSLGKAADLLGIGRDNIISVPVDAHNKVDIDAMRVAAAQLQARN
IRVMAIVGVAGTTETGNVDPLAEMAALAKEIDCHFHVDAAWGGATLLSEKYRYLLSGIELADSVTIDAHKQMYVPMGAGMVLFKNPSFASAIEH
HAEYILRQGSKDLGSQTLEGSRPGMAMLVHACLNIIGREGYEILINGSLERARFFAGLIEAQTDFELISEPELCLLTYRYVPAKVRLALAQAVA
QHDKDKLDAIGSLLDDLTRAIQKTQREQGKSFVSRTRITPKRYGNDKRTVFRVVLANPLTTEAILAAVLEEQRALASGETEILQALDALC
>B_A0A0C2M9Q6
MTLSKQEMPKKSSFFETEQALNYEIEEQVMQLFASSSHVTSIESQIDEITDKFSQNFLNTIDANTNIDLDSLLRSFSDSQIPVQPASLESYLKY
IDNNIVAHSIHTSSPRFIGHMTSALPCFVRPLAKLMTAMNQNAVKIETAKALSFCEREALAMLHRLIYNLDDNFYAQHIQNNLSTLGILVSGGT
VANITALWCARNTALGPKDDFLGIEKEGLTAALDFYGYKGAAIIGSELMHFSFDKAADLMGIGTHRLMRIPTDCNNRVNIQALRQAVMECRAQN
LLIIAIVGVAGTTDSGGVDSLTEIAEIAQEAHVHFHVDAAWGGPLIFSQQHRHKLAGIERADSVTIDGHKQLYLPMGIGLVFFRDPHIAKAIEK
QASYTMRKGSFDLGKRALEGSRPGMALFLHAGLNLIGLKGYEFLIDEGIRKTQYMADRICSMPEFELLAEPDTNLLLYRYIPEQLRELVFKKQL
TEIDNQLINQFNERLQKIQRQIGRTFISRTTKTTTSFGKEIPIIVMRAVIANPLTTEEDIDAVLNDQIQIASEIEI
>B_A0A0V8JBT8
MESNSQMTTVVNLKTKFDDLFIKEGMSGHDAYLQAIDQTKKVVADVLVQARGPFSGETPQMIQRAVKEMKLATEKGQPLEVVLEDVATLLLHSI
QVNHKACIGHLHCPPLVPALAAEMVISAANQSLDSWDQSSSATYLEQELIDWLCNRLGLGENADGTFTSGGTQSNYMGLLLARDAFCEKQWNWN
VQKKGLPPEARRLRILCSEAAHFTVEKSASQLGLGEDAVILVSTDRHQRMSLTDLKEKLRILKAENLLPYALVGTCGTTDFGSIDPLLHMAKAA
KENGLWFHVDAAYGGALMLSHNDHHRLRGIEQADSITADFHKLFFQPISCGAFLLKERKQFKHISYHADYLNPEQDDEEGLLHLVNKSVQTTRR
FDALKLFMSLRAVGLDRFGEMIDHTIELARKTAQLMKESRFFTVINAEPEINAVVFRYEPSFLKNNFDALNKQIHSELLQGGKALLAKTMHEGE
LYLKFTLLNPRVTMEDMKKLIADIAELGETIHRREGKSS
>B_Q2LV45
MNSLRTDNIPPPRKIETTLEYMHKLFIMPNSSDRFIEFGDLLLDMIHDFFQQKGGIHSEIPLKKLAKIFNNIDIPQNQMLIRDVLQEIKGNIIN
HSVKVANPYYIGHMTSAIPYFMILLEMIAVSLNQNQVKIESAKASTFVEREFLCWMHRLVYQNPSQFYKKNIQNHRFALGNITSDGTVANMTAL
ALAVAKAFPPDGKRFEGIRNEGLPRSLEYYGYRQALVLVSRRGHYSICKIGSILGIGSKGVVFSPVHPYTNKADIDKLWQTIEKIRKEDKRVGK
PSRFLALVGIAGTTETGNIDNLDAMAGVARELNAHYHVDAAWGGGALLMENGRELFTGIDKADSVSLDAHKLLFSPNAMGICLFKNIDDSLKLY
HTSNYIIRKGSVDLGRFTLEGPRPFACLKPWAAMKIIGRRGYELIFLHACGLQDIFITLIKNDPQFELLNTPELFIINYRFVPKELREMLDRLM
LDPRANAEKITQINNVVNEINVELHKKIRVRDTSFVSRTRLESTRYSPRKIVVLRAITINPNTEPHMLEQILVEHRLLGEQIWQDRNLLRKTLH
ICNKLIDV
>B_A0A0U2ZFL7
MSQAKASLAHLYRVFTKPEGKDSRLAQIEAQLSDNLADFLSGHIVTEEIPLEQIEQDFSQAQVPEQPQFVSDHAQHLLDTLVSKSVNTYSPRFV
GHMTSALPYFHLSLAKLLVGLNQNLVKIETSKAFTPLERQVLGMMHRLVFAGNDGFYQQHLHSATSSLGTFCSGGTLANLNALWVARNNALPPV
AGFRGISEDGLIAAMQHHNINQLAVISSARGHYSLAKAMDLLGLGKSQLHKLNCPQQTLDPQQVLQKGQQILQQGGRVMAIVGIAGTTETGHID

PLDELAEVASELNCHFHVDAAWGGATLFSERHAPLLSGIEKADSVTLDAHKQMYVPMGAGMVLFKDPLAGNAVRHHAQYILRQGSKDLGATSVE
GSRNGMAMMLYASLHILGRQGYELLIDESLAKARKFAQLIKAHKDFELVTEPVLCLLTYRLCPGHLREKGRVISDKHNHQLDELNRLIQKQQRE
AGHSFVSRTRLSTESYPEPITVFRVVLANPLTTEQDLQAILAEQSELATKHPLWDRL
>B_A0A1I1P8H3
MNEQQMKIKRNAVASEESLMRIFTAPEAPGSTLNRIEQEISSNLVGFLNENIAAIEKPLHEIEKDFQASQIPEQPTFVSDHANQIMDKLVAHSV
HTAAPSFIGHMTSALPHFVLPLSKLMVGLNQNLVKIETSKAFTPLERQVLGMMHHLVYGQDDNFYQKWMHSAKNSLGAFCSGGTIANISALWIA
RNRLLKADGDFKGITSDGVMAAMLHYGYKGLAVLVSERAHYSLGKAADVLGVGRSNFIAIETDNDNKVNVIAMRSKALELESQGIKVMAIVGVA
GTTETGNIDPLNALADLSEELDCHFHVDAAWGGATLLSNNNRHLLKGVERADSITIDAHKQMYVPMGSGLVLFKNPNASNVIEHHAEYILRKGS
KDLGSHTLEGSRPGMAMLVHACLQIIGREGYEMLIDKGIEKARYFADLIKLSNDFEVVTEPELCILTYRYVPEKIKQLLLAEGKDALDEKTKLE
IYIALNRLTASIQKRQREQGFSFVSRTRLTPSQYNKLPTVVFRVVLANPLTSEPILQDILEEQKGLAISDPIFKKYLNKYL
>B_A0A1G5GHE0
MEEQVSAEKSDGMKQEWVNLIRVFTCPEDEKTRDVLVHHMRQILFELHDFLIQNVGITEAVSLKEISDRCTDTVMNSLPEKRLADVVSGVFEEI
APHAVNVASPYFVGHMTAAIPYFMVHLKAIVAALNQNVVKLETSKMLSVVEKQVVAKVHRIIYQKDEAFYREHVQNTHTSLGSFTEGGTTANLT
ALWVARNRALGPVGAFEGVEVEGLPAAYAAHNIDRCVILVSRRCHYSLRKCGGILGIGNGNIIPVPVDRTNRMRVDRLAEMIEGFKKEGRTRVA
AVVTVAGTTETGNVDPIPEISALCRKEGIYLHADAAWGGPTLMSETYAHLLKGIEQVDSVTIDGHKQFYMPMSTGMVFFKDPTSLDAVAYHANY
IIRKGSVDLGSKTLSGSREANCLILDSAFKIMGSRGYGMLIDHGIELARSFAEEIERRGIFQLVTRPELNILTYRVCPAPFQEEMAQASPERIL
ELNEHFNEVNRALQQEQREAGNSFVSRTVLELETVFPGFPEGKVVVLRSVLMNPLTTMDVLRDILDEQEALYADWLRGTSTRKGRNREDAI
>B_A0A1E7Z8S9
MGEAQVSLKHLFRVFTKPEHKDSKLAQIEQHLSDNILDFLSQHVVTKKTSLEEIEQDFSDAFVPESPEFVSSHAENLLDKLVAHSVNTYSPTFI
GHMTSALPYFHLPLAKLLVGLNQNLVKIETSKAFTPLERQVLGMMHNLVYGQEDSWYQQYLHSARHALGAFGSGGTVANITALWVARNKMLGAD
KGFAGVAKSGLAAAYRHYDINQMGVMCSRRGHYSLSKAVDVLGIGRDNLITLPCPQQTLPPEQALAEGKRYQEEGNKLLALVGIGGTTETGHVD
PLDELADVARELGCWFHVDAAWGGATLFSETYRTRLKGIERADSVTIDAHKQMYVPMGAGMVLFKNPEDANSVRHHAQYILRQGSKDLGATTLE
GSRNGMAMMVYSSLHIFGRRGYELLIDESIKKARKFADMIEAHPDFELVTAPTLSILTYRVCPPVIQEKLKTADEGEKNQINHKIDVLVVNVQK
LQREAGKSFVSRTRLENPDFNSRPITVFRVVMANPLTTEQDLKNILAEQSKIAHARHLWKELEKSESSLTT
>B_K9VN97
MNKLTVKAKNSQDVMQPLKASVTPLEEKLMELFSASEKSFLAEEAMEERISALVEDFLYAKTASTDIDLELIIQEFTENKIPALPADVGNYIDD
LAQTLVAHSIHTSSPRYIGHMTSALPYFVRPIAKLLIAMNQNLVKMETAKVLTPCERQTLAMMHRLIYDFSEDFYNQHIQKNDSTLGIVVTGGT
LANITALWCARNASLGPKNNFSGIEKEGLYAALEFYGYKGMVIIGSSLMHYSFEKAAGLLGIGDRNLLKVPTDSHHRMDLAALRQTVAECRDRN
LHILAIIGVAGSTDSGSIDPLPEIAAIAQSANAHFHVDAAWGGPVLFSERHRHKLAGIQLADSVTIDGHKQMYLPMGIGMVLFRNPQIAKVIEK
NAPYTVRKDSIDLGKRSLEGSRSGASLFLQAALNIIGHQGYEFLIDAGIHKTQHLANLIRENPAFELLLEPEINLLLYRYIPETFREKAAKKQL
TEADNYFINEFNERLQEAQRQAGDTFVSRTTVQNTCYGKTIPIVAFRVVLANPLTTEADISFVLNDQLKIAAKLSLTIANSDRCDEVNYAVY
>B_A0A127VCS8
MNTTLNWVSPDILGTIKLKESDQLAISFLNQHITYKELNEKSNQLANFFLSKGIKKKEVIGVGMSLSIDYVIAMLAIMKAGAVFLPLQITGPAK
RLEQILDIAKPVLILENNREKIIPNSLIFSTSAVRQTEEADIIPWYNFSRLLPELLIDGDDPVYIFFTSGSTGVPKAILGNHKSLAHFIIWESN
TYQLNHQIRVGAMTPTTFDPSLRDILVPLFNGGILVIPNQENIDFSERLRWLIEEKINLIHIVPSVYEKIHDELELLGQHKNKIELSHIFFAGE
PLYAERVSKWKVQLPESTLLINLYGPTETTLAKFHHQLTAQDLLKGGIIPLGKAISDTTMLLFDEKGQPVEAGTIGEIIIHTEFCSLGYLGEEL
NAGKFIDYAGGKAYKTGDLGKLNEEGNLEYHGRIDAQIKINGLRIEISEIESTLLKYQGVKKCAVLYTEYSKTALSAFIISDEQLIKEELRLFL
KDYLPQPMIPAQFITLDNFPYNQNGKIDRVALAQLLTEQTGIPEQIITNEAVAEQFSELEQFIFEIWAEVLKKEKIGLHDNFFEIGGHSLNGLV
IVSKVFKKLGIRFEITSFFNNPTISSFAAVISQLQMQNQDQQQGQNQQQNQHTQQNHDKKQAQTNEEIKLVPTQAFYQVSNAQKLMFLLNEITG
NSSVYNIQGAYTIKGQLNYQILEESFKELISRHEALRTCFVKDDEGGFFQRILPVEAVDFKLGITQIHSAGAGHDFLQTAFSYAFDLTKDLLLR
AGLIEKDGEYILYYCMHHIISDGWSLEVLISDLLHIYKGKLATPNNLLAPLKLQYKDYAAWDNQRNSELAAAEKYWLAQFSGNLPVLNFPLNFS
RPAVQHYAGDSISIDIPQETVQGLRNSCKTQNATYFSGLLTVLYTLLYRYTGQKDIVLGSPVANRDNPELKNQIGLYVNTIAYRAAIKDDDSFN
TLLHTIKNMVIEGNQHKNYPFEKLIENLDIARDLSLNPLFSVMVNFQTPQEIKSTADLPENLSIEKFDFPQIKSKFDFSFYFYDQQGHTKLMLI
YNTGLFTREFMQDLLDNFLNLAKSLLVQPETNLSYIPYLSDKEKKKVLEVFNPTNSPKKAVQSLTRLFGQHVKHHSSNTAFINQDKTYSYGFVE
RLSNQFAHYLYQEYAVGKNDRLIVSLERNQWLPIAILGILKLNAVYIPVEPGAPKLFQERIIEEAGCKHVIDEVFLEKFIAVADHYAAEEPDAI
NYPAGESAAVNPKNDIITIIYTTGSTGKPKGVIIKNSNIVNRLEWMWSEYAFHPDEVCVLKTAISFVDHLWELFGPLLKGIPLVCLHNNDISDI
PHLIGLLNKFKVTRIVLVPSLLKEILAYPEACRAQLSALTYWTSSGEDLPGILVQKFYEIFDSRKYKLLNIYGSTEVTADATCHDTSIDFKQNK
TILALFNTDIDIPGLIENYDHADHIFAADQPDRILSGFKNIEFESVTSPEEYLKYLEQRLLPNIINISSNKYIGHMTGPLPNFSLSLSSLVNRL
NQNLVKIETSLAATMIEIEVIKRFHQAVFNRSDAFYSGLSHKNEGIPGIFTNGGTLSNITALFYALNNCLAPHGSYQGYGNSGLIKGLLHYGYK
DVVILGSRWCHYSVGKALKLFGLGKDSFREIIVDQGNEADIRKKVTQQILDLKAQNVLILSIIGVAGTTESGAVEPLNLLGELAKEQAIHYHVD
AAFGGAYILSEKYSHLLDGINKADSVTLCAHKQLHVPVGSSFCLFADPGFVKNSENNTNYQARKNSYDLGKFTIEGTRNFNSLILHGLFSVFGK
KGLAEVIDYNHELSRTFADLIHQNNAFELFESPKLNIVLYRYVPFSLRNKTVLTDQDLLLINDLNEQIQKTQFERGNSFVSYTLVRNYDRDSLQ
HVFFRTVFSNPNTTQKVLKEILQEQLDIAAELDHTLGINTEQQNYNSNFTVHTNTSIGKPIANVKVYILDDHLQLLPVGLEGIIYIGGENITDG
YLGDEELTGDKFIENPFKPGEILFRTDDVGKWTVEGEIIYIGRRDFQYKVRGNRINTNSIEETIAQLSGIDQALVIFNDQKLTAFVSAATIPDL
SVLRLMLKEILPAYMIPNEFVWIKDFPKLTNGKINRKALHAEIGTVLTENVARILPENDLEQQVYDLWKLIFNQKELSVRADFFAIGGHSLSLM
KLIQYYHKAFHVQVKMQDLFLTTTIEGHAGLIRNLDKNPMAPIEKVAESDTYPVSDGQRRIWTLSQTADSSRAYHMSGAMTIHKALNTAKFNAA
IREVIKRHESLRTNFIWTEKGEVRQIIRPDELLDFELEIINVADQTQEEIKDALIAFNNKPFKLADDLLLRIRLFQVATDHFIFSYCFHHIISD
LRSMKIFARDLWNQYQYLNENLESTITPLKLQYKDYSAWLQKAKHRIQIPASVYYKELLSDFELLNLPADYQRPLLKTYTGNTYTYEIPAALCT
LVRRKSLSYGVTPFTVLVAAINGLFYRYTQQEDILIGMVTEGRGHPDIDQQIGFFVNTLPLRTQLDPKKGFGHLCKEIQHLVLQNFNFQAYALD
ELIDELQVARDPSRSPLFDVVVTYENYEQELNFSEMEALNLEKKTAQFDITFGFVENKDFISSKIEFNTSLYAHSTIQRMATHLAEFLAAALAK
EEDALYTLDYLTPEEKNYE
>B_K9Z2H8
MSSSSQARPFDRYFLTAQTESIESYQEAIALTQELIVNNILQEDKPYFGLNPSLLQESFRDFCPKNFTQQDYPQIEPELAEIIRHGAMVTHPAC
VAHLHCPPLIPAIAAELIIGSLNQSMDSWDQSPSATILEQQLTRWLCDLFGYSNSADGTFTSGGTQSNLMGLLLARDNYAKTHLKWHIQRQGLP
PEAQQFRIFCSDVAHFTIRQGAAILGLGENGVIPIETDENFQLKPEILSAKLRTLQQDNLRPIAIVATAGTTDFGSIDPLPELAKIARDHGLWF
HVDAAYGGALKLSQNHGHKLKGIELADSITVDFHKLFYQPISCGAFLLKNQANFGLIKLHADYLNPESNEAQGIPDLVTKSIQTTRRFDALKLW
LSLKTLGVETFGEMIDSTIELAGAIALIIAEDAELELANIPTINAVVFRYQPSQGTATEIDRINEQIPKKLMLEGKGIIAQTQVKGRNYLKFTL
LNPLTTLKDLQKLLIEIKSLGQTLHTTDINHELKKNC
>B_A0A1E5CWM7
MVSEQKTVDVNFESLLRIFTVPEGPDSTLTQIEEKLSQNLNQFLREHIVAEEKPLLEIEKDFSNAHIPEQPEFVSEHTEHLLDSLVAHSVHTAS
PSFIGHMTSALPYFLMPLSKIMIALNQNLVKIETSKAFTPLERQVLGMLHRLIYQQDDKFYSRWMHSSNHSLGAFCSGGTIANITALWVARNNA
LRAEGSFKGVEKEGLFRAMKHYGYEGLAILVSERGHYSLKKAADVLGIGQDSLIAIKTDHHNRICPRALDEKMSELKANKILPFAAIGVAGTTE
TGNIDPLKEMAAICRKHDCHFHVDAAWGGATLMSNSHRSLLDGIEEADSVTIDAHKQLYIPMGAGMVLFKDPDAMMAIEHHAQYILRKGSKDLG
SHTLEGSRSGMAMLVYASMHIIGRPGYELLINESIEKAQYFAELIQQQQDFELVSQPELCLLTYRYLPQDVKQALFIADKGSKKKLNNLINELT
KFVQKKQRETGKSFVSRTQLNPEQWDELNTIVFRVVLANPLTTHDILQSVLVEQRQIASLAPKLMAKIQQLTQEILA
>B_A0A0S2KBH9

MSESKKLASASLESLYRIFTVPEAPDSTLGRIDQSISQNLAGFLQEHIVASERDLSDIEKDFADPQIPEQPTFVSDQIQFVLDKLVAQSVHISA
PSFVGHMASALPYFMLPLSRIMMALNQNLVKVETSKAFTPMERQVVGMIHHLVYGRDKDFYTQWMHDRAHALGSFCSGGTIANLTALWAARNNV
LAPTEGFAGLGQEGLVAGMQHYGYKGLAILVSERGHYSLGKAADILGIGRRSLIAIETDANNKIRTDLLAKKCDELAAANIKVMAIVGIAGASE
TGSIDPLDQMADIAQERGIHFHVDSAWGGPTLFSNTYRPLLKGIERADSVTLDAHKQLYVPMGAGICVFKEPSTLSSVEHHAEYIIRKGSKDLG
SHTVEGSRPGMAILVHSGLHIIGRQGYELLIDQGISRARHFADMIRADKDFELISEPELNILTYRYVPQDIQTALESAPASLRTEVNEQLNLVT
KRIQKTQRGLGRSFVSRTRLNPHHHNRNAVIVFRAVLANPLTTNQILQDMLAEQKEIAATPPIQALLKQVRNLFCGC
>B_A0A1G9K0B9
MNRAILTHEEIPKSMIENRENPIAIKKEEFKKLGYQLIDSIADFIDSIEERPVTTTKTPSELQKTLGDAPLPTNGTPAGELLTKTTDLLFNNSL
FNGHPKFLGYITSSAAPLGALADLLASSVNANVGAHILSPIATEIEKQTVKWLAEFIGVSPNYGGILVSGGNMANFTAFLAARTAKAPKSIKED
GISNASEKLTVYCSKSTHTWIEKAAILFGLGTKSIRWIPTTSSNQLDEKILEETIKEDIKNGCKPLMVVGTAGDVSTGVVDNLAAISNLCKKYD
LWFHIDGAYGIPAAVIPELQRMFEGVSEADSIALDPHKWLYSPLEAGCTLVKDPQHLIDTFSSHPEYYNFSSVEGEVAQNFYEFGLQNSRGFRA
LKVWLTLQQVGRNGYIKLINEDIELSKLLLLQAENHEELEAIAQNLSITTFRYIPKHYKSSAGQKEDYLNALNEALLNDLQMGGEVFLSNAIVN
EKYCLRACFVNFRTSQKDIGEIIAIVVKEGRKTHERLSQTKD
>B_A0A068QSK3
MTCATQAKNDDEADKMALHTSCLDEQAWQALLDQFTPHPGEEGLLELETNIADDPVTYCRRELGHPIGELTELEKVFVSVDIPEFPLNTDNYAR
YLSEDILNHVVPVSSPTFVGHMTSALPKHLPALGKVLTALNQNLVKLETSHILTALERQVLGMMHKLVYNRDEDFYQQWLHSGEHALATFCSGG
TLANLTAMWACRNQLMPADGDFAGLAREGLARGLLHYGYRGLAILVSEQGHYSLKKTVDVLGLGQDALVKVETDREGRICIDALLAQLQTLRQR
NIKPMAIVGIAGSTETGAIDPLNRLADIAEQAQCHFHVDAAWGGASLMSERYRHLFAGIERADTVTIDAHKQLYVPMGSGMVLFRQPTLTNTIA
QHANYIVRKGSKDLGRHTLEGSRSAMSLMLHSNLYLLGRRGLARLIEAGIEKARQFADLIHQQEDFELISEPQLCLLTYRYVPPKVLQALQESS
APVRQQLHEWLNALNQDIQTAQWRAGKSFVSRTCLKPVQWDRQPTTVLRVVLANPLTTLEILENMLEEQRQLARQSPFWQTLQAWAVIS
>B_A0A068QSK8
MYPEVRRVYMSEKMSEKQEWQALLDQFISHAGEEGLFELETNIADDPVTYCRRELGHPIGELTELEKVFVSVDIPEFPLNVDSYARYLSEDILN
HVVPVSSPAFVGHMTSALPKHLPALGKVLTALNQNLVKLETSHILTALERQVLGMMHKLVYHRDEDFYQQWLHNGEHALATFCSGGTLANLTAM
WACRNQLMPADGDFAGLAREGLARGLLHYGYRGLAILVSEQGHYSLKKTVDVLGLGQDALVKVETDREGRICIDALLAQLQTLRQRNIKPMAIV
GIAGSTETGAIDPLNRLADIAEQEQCYFHVDAAWGGASLMSERYRHLFAGIERADTVTIDAHKQMYVPMGSGMVLFRQPTLTNAIAQHANYIVR
KGSKDLGRHTLEGSRSAMSLMLHSNFYLLGRRGLASLIEASIEKAQQFADLIHQQEDFELISEPQLCLLTYRYVPPTVLSALREGSASVRQQLH
ELLNALNQSIQTSQWTAGKSFVSRTHLKPVQWDRQPTTVLRVVLANPLTTLEILENMLEEQRQLARQSPFWQTLQDLVGD
>B_A0A1H2Q464
MTGNKKSAHASLEAMYRVFTVPEAPESTLSRIDQNISRNLAGFLQEHIVAVERDLSEVEKDFSDPVIPEKPIFVSEQTQFLLDKLVANSVHTAS
PSFIGHMTSALPYFMLPLSKIMIALNQNLVKTETSKAFTPMERQVLGMIHRLVYNEDDSYYRKWMHDPRHALGAMCSGGTVANLTALWVARNRA
FPAEGSFRGIHQEGLFRALRYYGHEGAAILVSRRGHYSLRKAADVLGLGRESLVAVDTDDNNRIQPDALRDKCLELQKQKIKVMAICGIAGTTE
TGNVDPLDAIADIAREFGAHFHVDAAWGGPTLFSRRHNHLLRGIEQADSVTFDAHKQLYVPMGAGLVVFRDPALASAVEHHAQYIIRQGSRDLG
STTLEGSRPGMSMLIHSGLKILAREGYEILIDQGIEKARRFADMIEQRPDFELVTAPELNILTYRYAPADAQHALTLADPLQSERINTSLNRIT
KFIQKTQRERGKAFVSRTRLEPACYHHFPCLVFRVVLANPLTTEDILTDILNEQEELATDSGLSDEMTILRQLTEAVLTATGAESRQA
>B_A0A1G7ZM35
MESEQKTVDVSFENLLKIFTVPEGPDSTLTKIEAELSQNLNQFLREHIVADEKPLKEIEKDFSDAAIPEQPSFVSDHTQHLLDTLVAHSVHTAA
PSFIGHMTSALPYFLMPLSKIMIALNQNLVKIETSKAFTPLERQVLGMLHRLIYQRNDAFYQQWMHSAEHSLGAFCSGGTIANITALWVARNNA
LKAQGSFQGVEKVGLYKAMQHYGYQGLAVLVSKRGHYSLKKAADVLGLGQDELVAVETDHNNRLCVKDLAQKMASLKAQNIKVIAVIGVAGTTE
TGNIDPLSDIASLCAEHGCHFHVDAAWGGATLMSNRYRYLLDGIEKADSVTIDAHKQLYIPMGAGMVLFKDPDAMRSVEHHAQYILRKGSKDLG
SHTLEGSRSGMAMLVYAAMNIISRPGYELLINQSIEKAKHFAALIKAQPDFELISQPELCLLTYRYVPTATKQALEVANQAQQQALNDKLNELT
QFIQKTQRETGRSFVSRTQLNPQHLNQLDTIVFRVVLANPLTHNEILESVLSEQREIAKQAPNLSKQLQSLTNDILEK
>B_Q39V49
MPKNRDAARASLENLYRIFTVPEAPDSTLGAIDQAIAGDVAGFLQTHIVAIERPLEEIEADFSSFSIPEEPTYVSEYTEFVKENLVAHSVHTAS
PAFVGHMTSALPYFMLPLARLMTALNQNVVKVETSKAFTPMERQVLAMLHHLVYGRNDDFYPQWIHNSQHALGAFCSGGTLANVTALWVARNRL
FAPDGEFRGIAQEGLARALKHRGADGIAVLVSERGHYSLGKAADLLGIGRDDLIKIKTDANNRIDLKALREECRRLQDRNTLPLALVGIAGTTE
TGNVDPLEAMADLAQELGCHFHVDAAWGGPTLFSDRHRHLLRGIERADSVTIDGHKQLYVPMGAGMVVFKDPTALSAIEHHANYILRHGSKDLG
SHTLEGSRPGKAMLVHAGFSIIGRKGYELLIDMGIERARTFADMIQRHPDFELISEPELNILTYRYCPPAIQQALTDATAQQRAAINGLLDQVC
QLLQKYQREAGKTFVSRTRLHVARHDMELTVLRVVLANPLTTDEILEAVLAEQCEIVRLPEIQALLRQAEELCPGLAKAV
>B_A0A1I4S397
MTDSRLAALLQQHFSIDALERAPAPLQHALRLLGDWLGASRAQPYPNTPFAELAALFGDQPLPAEGMPVEAFLEQLDATVLANTAQLNHPKYIG
HMTQALPWISVVAEALTAALNQNQVKIETAYASTLIEKQVLGWLHRQVYRGGEALYAEAMAAPNIALGNMVGGGTMGNLTALAVALEHQLPGYR
KDGLLAALERSGHRGLAVIGSARSHYSLKKALATLGLGENALRVVPVARDNRIDLAALEDEVAALRAAGVKLLAVVGIAGATETGSIDPLAQLA
AVARREGAWFHVDAAWGGALLLAEQFRPLYEGIALADSVVLDGHKLLWVPMPQSMVLFRDSASLNLLKHNANYILRDNSGDLGQTSLEGSRRFD
ALKLWTSFKVLGVSGYRTLLEQAATLTGQMRALLAEQADFELVTDSDTFILTYRYLPQRLRQQMERLLADGRLEQAAALNQRLNALNSELQTRQ
KELGHSFVSRTVLESTAYPGQTTVLRVVLTNMTTTEEHLREILAEQADIGEGLLEQMGL
>B_A0A1H4BYW3
MDGNRNNSIEINKEEFRKIGHQLIDDISDFLSSIDKKPVTIKESPSQLQSILGNTSLPEKGTPATELISRATDLLLNHSLLNGHPKFLGYITSS
AAPIGALADLLAASVNPNVGAHILSPMATEIEKQTIQWLAEFIGVSPNYGGILVSGGNMANFTAFLAARTAKAPKSIKEDGLSNISQKLTIYCS
KTTHTWVDKAAILFGLGTKSIRWIQTDSSNKMDNKVLEETIKEDIENGFKPIMVIGTAGDVSTGVVDNLKGISTICKDYDLWFHIDGAYGAPAT
IIPKYKNLFDGLSEADSIALDPHKWLYSPLEAGCTLVKNPQHLIDTYSSHPEYYNFSKDENEIAQNFYEYGLQNSRGFRALKVWLSLQQIGKSG
YEKLIGEDIELSELLFDLAKKNPELEAVSQNLSITTFRYIPFNSKDDNDYLNKLNEELLNALQTGGELFLSNAIVNEMYCLRACVVNFRTTKKD
IKEIIDIIIREGRKTNLKLQQRKVN
>B_A0A1W1H5C0
MESPDQLKPIDISKHSKLVADWDSLQRIFIRPEDEDCKATLLKYMEQILFGLHEFLNSHVGVTEEISLLNLTETYKETLISSEPEKKLADVISD
IISSIAPRAVNVASPYFVGHMTAAIPFFMVHLKAIVAALNQNVIKLETSKVLSVLEKQVLAKIHRLIFDLDESFYQKHVQSTDTSLGSFTTGGT
TANLTALWVARNHFFSPCDGFEGIETEGMAGALNHHGFDKAIYLVSKRCHFSLKKAGGILGVGNKHVIPVDVNAEYRMDVSKLSKMIKSFSKSA
GTSGRKGKTAIMAVVGVAGATETGIVDPLYEIADVCREHSIHFHVDAAWGGPVLLSEAYAGKLAGIERADSVTIDGHKQFFMPMTCGMVFFRDP
KIMDRIAYYSNYVNRQGSVDLGIKTLEGSREANSLILDSALKIMGTKGYALMIDHGIETTRAFADEIKKRQMFELVTEPELNILTYRLVPPHIG
KLREKAYDRKEIERLDGLLDDINIMVQRKQREAGRSFVSRTRVNCAPCRENDHEVVVLRAVIMNPMTTIRVLSEVLDEQEKIYKSISHEEA
>B_Q4BVC3
MISEFYQFYLAAMNDFSEQIAALSPSEAMNSEQRSQPTRENQIKAEKISTEQRILQYFKPEEIQRGTFTQPDISLDALAQQFKRSELPTYPITV
DDYYRQLFQEVLPYAIDTGSPTYIGHMTSALPDFLHDMSKLISKLNQNLVKIETSKSLLFLEREAIAMLHRLVYNFSEEFYTENIQQKNRNLGI
VTTGGTTANISALLCARNAGLLSKENSTELLKESLYKVLSKKGYEDMVIIGSRLMHYSLNKAASILGLGTDNIIFVDNNSEGKLNCKQLEEKIQ
ECQENRLFILALIGIAGTTETGQIDPLWKLGEIAEKYNIHFHVDGSWGVPTIFSDKHKGKLKGIEKADSVTICGHKQLYLPQGISICLFKDPQL
LNYAATTARYQAQQDTYDLGRFTIEGSRSALSLCLHGALHIIGKKGYEMLIEQGIYQAQYFARKIDKLKSFELILEPTLNIVNYRYIPDYLRDK
LRKKMLTNAEIERLNQINQEIQQEQFEQGKTFVSKTILFDPKYSKKIIVFRTVLSNPNTTATDLDNVLEDQLRIVDKIQRTKGG
>B_A0A1E7Q4B7

MTEPIQRQATASEESLIRIFTIPEAPDSTLSIIEQNLSQNLAGFLRNSIVALEKPLWQIERDFQAHQIPEMPEFVSDYAEQLMQKLVAHSVHTA
SPSFIGHMTSALPYFVLPLSKMMVGLNQNLVKIETSKAFTPMERQVLGMMHHLVYGEADGFYKKWMHSANHSLGAFCSGGTVANITALWIARNR
LLKADGEFRGIAREGLFRAMQHYNYQGLAILVSERGHYSLGKAADILGIGRDALIAIKTDANNKIDVTELANTMAELTARNIRVMAVVGVAGTT
ETGNVDPLHQLADLAEQYQCHFHVDAAWGGATLFSERYRHLLAGIERADSVTIDAHKQMYVPMGAGMVVFKHPSDAHAIAHHAEYILRKGSKDL
GSQTLEGSRPGMAMLVHACLQIIGSAGYQILIDRSLEKARYFAKLIMDNSDFELITEPELCILTYRYVPASVQKAMAKANPAQLQRFNELLNGM
TKFIQKRQREEGKSFVSRTRLTPAKYQRQETIVFRVVLANPLTTEQILHDVLLEQDHIAQLDKEFYPKLLQLAKKLAA
>B_K6ZXH3
MAKAEVSLEHLFRVFTKPEHSDSTLAKIERHLSDNISDFLSQHVVTKKTSLEEIEKDFSSASVPDSPEFVSEHAESLLNKLVAHSVNTYSPTFI
GHMTSALPYFHLTLAKLLVGLNQNLVKIETSKAFTPLERQTLGMMHNLIYQQDDKFYEDYLHSADHALGSFCSGGTVANITALWIARNQLLGAD
GKFPGVAKAGLAAAYRHYGYENIGVMSSNRGHYSLSKTVDVLGIGRDNMLTVQSPTQKLDPQQALKLGKAYQESGNKLLAIIGIAGTTETGHVD
PLDELADVAKELNCWFHVDAAWGGATLFSERYRSKLAGIERADSVSIDAHKQMYVPMGAGMALFKDPKHSNAVRHHAQYVLREGSKDLGATTLE
GSRNGMAMMVYSSLHILGRRGYELLINKSIELAFEFAEMINVHPEFELTTQPTLSLLTYRLRPFSLSQLAGMPASDITNINNRLNSLTVSVQKQ
QREAGKSFVSRTRIEIEKYNGEAITVFRVVLANPLTTKEHLSDILQEQLTIANNNRIWNELSVNPTSKIA
>B_A0A1M4ZHC6
MTNKRHAPIEIDNEEFRKAGHKLIDTIADFLGTIQQKPVTTGETSKALQALLGMASLPADGTSPEHIIDKASALLLDHSLFNGHPKFFGYITSS
PAPIGALADLLAAIVNPNVGAQVLSPMATQIELQTIQWLSEFIGVPSTYGGILVSGGNMANFTAFVTARTIKAPKKIKSGGIAEEDKLVVYCSK
ATHTWIEKAAVLFGLGSNAIHWVETSADNKMKVEALERMIQKDLQDGCRPIMVIGNAGDVSTGAVDDLQDLSAICKKYELWFHIDGAYGIPAAV
LPELKDMFAGVDGADSIALDPHKWLYSPLEAGCTLVKDPNHLIETYSSHPVYYNFSSTVERTHNFYEYGFQNSRGFRALKVWMSLLQVGRNGYE
QMIREDIELSKLMYALVDAHAELEAMTQSLSIATFRYVPANVKTIDNKEDYLNRLNESLLNELQQGGQVFLSNAVIGEKYCLRACIVNFRTSRE
DIEQVVQIIVAAGKKVHKALMEKPAINSQGA
>B_A0A0B5FTX7
MPRSKESARANLENLYRIFTVPEAPDSTLGRIDQAISDNVTGFLQKHIVALENSITDIEKSFDDPKIPEEPTFVSDYTEFIKEKLVAQSVHTAA
PGFIGHMTSALPYFMLPLSRIMIALNQNLVKVETSKAFTPMERQVLAMLHRLIYSESDAFYHRWIHDSRYALGAFCSGGTIANITALWAARNQL
CSPSGNFRGITNEGFFRAMQHLGCQGLAVLVSRRRHYSLGKAADLLGIGRENLVLVETDDRNRINMAALREQCRRIQDKGIRPLSLIGIAGTTE
TGNVDPLEEMADLAEEIGCHFHVDAAWGGPTLFSEKYRSLLKGIERADSVTIDAHKQLYVPMGAGMVLFKDPAALSSIEHHAAYIIRHGSKDLG
SHTLEGSRPGKAMLVHAGLSIIGRKGYELLIDQGIERARRFAEMVQEHPDFELVTAPELNILTYRYNPASVQEALAHADAETVREVNELLDKMT
KRIQKRQREAGKTFVSRTSLTPERYRGQTIIVFRVVLANPLTTEDILASVLEEQCEWARHPEVSPLLDQLQEMTAAA
>B_A0A1G6DFW1
MSDNKFAEVSQEALLKIFTIPEAPDSTLGRIEKHLSENLMGFLGERIVAREKPLHEIEQKFNASQIPEEPRYVSDHTEFLLNNLVAHSVNTASP
RFIGHMTSALPYFLLPLAKLMVGLNQNLVKIETSKAFTPLERNVLGMLHHLVYQQDDSFYQHWMHSAQHSLGAMCSGGTVANMTALWVARNLLL
KPDEQFAGIAAEGLAAGLAHYGYRRLTVMVSERGHYSLGKAADVLGIGRRNLIAIPADEHNRIRVDELRKACAEVVADGGHVLAIVGVAGTTET
GHIDPLNELADVAEEVGAFFHVDAAWGGATLLSKNHRHLLAGIERADSVTIDAHKQMYVPMGAGMVLFKDPSMVRAIEHHAEYIIRHGSKDLGA
HTLEGSRPGMAMLIYSALHVMGRRGFELLIDQSIARAQMFASLIAEQDDFEVITEPELCILTYRFVPTRVKQLIQDATPEQVRRITPHLNALTR
FIQKRQRETGRSFVSRTKLTPSQYYREPTVVFRVVLANPLTTKEMLVEVLQEQREIAAQAMSLRGALHTELRELGLLRDAALR
>B_Q8EG41
MTQKLPRQATASEESLMRIFTVPEDADSTLSIIEQKLSEDLAGFLGDSIAALEKPLSEIETDFQTFEIPNQPRFVSDYTDEIMQNLVAHSVHTA
APSFIGHMTSALPYFVLPLSKMMVGLNQNLVKIETSKAFTPLERQVLGMMHHLIYAQHDDFYRNWMHSANHSLGAFCSGGTVANITALWIARNQ
LLKADGDFKGVTREGLIKALRHYDYDDLAILVSERGHYSLGKAVDLLGIGRDNIISIPTDADNKVDVTQMRKIAVELAHKRIKVMAIVGVAGTT
ETGNIDPLKQLAALASELNCHFHVDAAWGGASLLSNKYRHLLDGVELADSVTIDAHKQMYVPMGAGMVLFKNPEFAHAIAHHAEYILRRGSKDL
GSQTLEGSRPGMAMLVHACLQIIGRDGYEILINNSLEKARYFAEQIDAHPDFELVTAPELCLLTYRYVPASVQAAMQVAIEQGDKAKLERFNEQ
LDGLTQFIQKHQREQGKSFVSRTRIQPARYFRQPTVVFRVVLANPLTSHEILNQVLIEQGEIATLDKEFLPALLAMVAE
>B_G4QDQ4
MAKAEVSLEHLFRVFTKPEHSDSTLAKIERHLSDNISDFLSQHVVTKKTSLEEIEKDFFSAAVPDSPEFVSEHAESLLTKLVAHSVNTYSPTFI
GHMTSALPYFHLTLAKLLVGLNQNLVKIETSKAFTPLERQTLGMMHNLIYQQDDKFYEDYLHSADHSLGSFCSGGTVANITALWVARNQLFSAD
GKFPGVAKAGLAAAYRHYGYDNIGVMSSNRGHYSLSKTVDVLGIGRNNMLTVHSPTQKLDPHQALRLGKAYQESGNKLLAIVGVAGTTETGHVD
PLDELADVAKELDCWFHVDAAWGGATLFSERYRGKLAGIEHADSVTIDAHKQMYVPMGAGMALFKDPKHSNAVRHHAQYILREGSKDLGATTLE
GSRNGMALMVYSSLHILGRRGYELLINKSIELALEFAAMINAHPDFELTTQPTLSLLTYRLRPQTLSGLENMPVSDIANINKKLDSLTVSVQKQ
QREAGKSFVSRTRIEIEKYNGEVITVFRVVLANPLTTKEHLSDILEEQIEIASNNRIWNSLSTNLSTKIA
>B_B3E5L1
MPTKARANLETLYRIFTVPEAPDSTLGAVDQAITADVAGFLQNHIVAMERPLEEIEASFSSVTIPEEPTYVSDYTEFVKENLVAQSVHTASPGF
IGHMTSALPYFMLPLTRLMTALNQNTVKVETSKAFTPLERQVLAMLHHLIYRCPDEFYPSWIHNSQAALGAFCSGGTIANTTALWVARNRFFAP
DGAFRGIAQEGLARALKHRGVDGIAVLVSERGHYSLGKAADLLGVGRDHLVKVKTAEDNRIDLKALRQECQRLQDQNIRPLALVGIGGTTETGN
IDPLEAMADLARELDCHFHVDAAWGGPTLFSDRHRHLLAGIERADSVTIDAHKQLYVPMGAGMVLFRDPTAVSAIEHHAAYILRHGSKDLGSHT
LEGSRPGKALLVHAGLSIMGRKGYELLIDLGIERARTFAGMIRQHPDFELTSEPELNILTYRYCPAAIQHLLATAPQAEQARINGLLDQVCQLL
QKHQRESGKTFVSRTRLRMSRYGEEITVLRSVLANPLTTDEILTSVLSEQCEIVRQPEIQALLQQVWR
>B_A0A0M4DHR6
MPKHRDVARANLENLYRIFTVPEAPDSTLGEIDEAISRDVAGFLQTHIVALERSLEDIEADFFNTAIPEEPTFVSEYTEFVKENLVAQSVHTAA
PGFIGHMTSALPYFMLPLSRIMTALNQNLVKVETSKAFTPMERQVLAMLHRLVYRGGDTFYARWIHDSRHALGAFCSGGTIANVTALWVARNRL
FAPAGDFRGIAQEGLVRSLKHLGCDGLAVLVSRRGHYSLGKAADLLGLGRDNLVLIDTDDDNRIDLKLLRAEFLRLQEENIRPLALVGIAGTTE
TGNVDPLAAMADLAAEFGCHFHVDAAWGGPTLFSDRHRSLLSGIERADSVTIDAHKQLYVPMGAGMVVFKDPTALSAIEHHAAYILRHGSKDLG
SHTLEGSRPGKAMLVHAGLSIIGRKGYELLIDLGIARAKTFAAMIRQHPDFELTSVPELNILTYRYCPQAVQQKLVLATSLQRTHLNALLDQVN
LLLQKEEREAGKTFVSRTRLRLAPYGQELTVLRVVLANPLTTDEILASVLEEQCRIVQHPEIRELMLQVDALCAEIEDGPTAPACAGAR
>B_A0A1K1QNC9
MKTTAAIGIPKLFSCQFEGKLLDAFPFEQGYTPASAEAYIDFVKQELAPQIVNVAHPRYIGHMTGPVPPLMHELHGCMALFNQNLVKLETSGIA
THIEREVIGELHRIFFQKGEEFYKYYTPDPDHCLGNITSGGTLSNITALSYALSKKLSNPEDPSASINNIGLLKSMVNKGYTDIVVLGTAHSHY
SMNKAMRLLGLGYQSFIKIDPAILKSAEGRGMLSGMMENYKKAGTLVLALIGVAGTTEAGIIDPLEMMAEVAAEHDVHFHVDAAFGGAFMFSDK
LADKLQGITQADSITLCGHKQLYLPMGISVCLFKSPELAGYSEINSHYQARKGGIDLGKYTIEGSRPFSAFIIHGVLRLIGKEGYAEILESNHD
RARYFGELVAAHGSFELYNTPELNIVLYRYVPTWLRTRVKKGTLSKTEQQELNRLNVIIQKRQFDKGSSFVSYTEIPDASGSGERVVWLRAVLM
NPYTSPQDLQEILQEQETIAAAVIL
>B_Q15NV7
MAVAEVSLEHLFRVFTMPEGKDSKLAQIEQHLSDNLADFLSQHVVTKVTSLEQIEQSFAEFNVPEHPEFVSEHAANLLEKLVANSVNTYSPTFI
GHMTSALPYFHLSLAKLLVGLNQNLVKIETSKAFTPLERQVLGMMHELVYAQKSDFYNTYLHNANHALGAFCSGGTIANVTALWVARNKALAAD
GVFKGVAREGLAAGLAHYGYKKMAILSSRRGHYSLSKSVDILGLGREQLITLDCPTQRLSPEKALAFGKEYAEQGNKILSIVGVAGTTETGHVD
PLDELADVAQELGCHFHVDAAWGGATLFSSTHRKILKGIERADSVTIDAHKQMYVPMGAGMVLFKDPNDSNAVRHHANYILRAGSKDLGATTLE
GSRNGMAMMVYASLHIFGRQGYELLIDQSIEKAKVFARMIQSHPDFELITHPTLSLLTYRVNPQEVQQQVKADFSLNVKFNEKLDKLTVYVQKQ
QREAGKSFVSRTRLQTQTYGDQTMTVFRVVLANPLTTENDLQNILDEQIAIASKSQNWAELTEKVI
>B_A0A1W2CRD4

MEKQYIKKDLVADWESLQRIFIRPEDDASRETLIKYMEQILFGLHDFLNKNVGVTEAISLKALTDLYKETVISPHPEKKLADVISDIIQGIAPR
AVNVASPYFVGHMTAAIPFFMVHLKAIVAALNQNVIKLETSKVLSVLEKQVLAKIHRLIYKNSEIFYHTHVQSTDTTLGSFTTGGTTANLTALW
VARNLLLKPFDRFQGVEAQGMAEALKAYSLDKAVILASPRGHFSLKKAGGILGIGHKNVIPVDVDAHHRVDVLKLNRRIVDLKKNGKIGIMAVV
GIAGATETGIVDPLDAIADVCEDHGLHFHVDAAWGGPVMLSERYAFKLKGIERGDSVTIDGHKQFFMPMTCGMVYFRDPLAMDAIAYHANYVNR
EGSVDLGIKTLEGSREANSLILDSSLKIMGTRGYGLMVDHGIETAQGFSRAIHARELFEMVTEPELNILTYRILPPKIKEQILSASPGEKEKLE
DRVNQLNIKVQREQREAGKSFVSRTRIRQGTKDTEGQVVLRAVIMNPMTDMDILNEVLDEQEEIAKKILQSPEPF
>B_A0A1I3QNC8
MTGKKPSAQASIEAMYRVFTVPEAPDSTLSRIDQNISRNLAGFLQEHIVAVERDLSDVEKNFSDSAIPEKPVFVSEQTQFLLDKLVADSVHTAS
PAFIGHMTSALPYFMLPLSKIMIALNQNLVKIETSKAFTPLERQVLGMIHRLVYQQDGAFYRKWMHDPSHALGAVCSGGTVANLTALWVARNRA
FPAEGSFRGLHQEGLFRALKYYGYEGAAILVSDRGHYSLRKAADVLGLGRDALVPVPTDDENRIQIDALRDKCLELQRQKIKVMAICGVAGTTE
TGNVDNLDAMADIAREFRAHFHVDAAWGGPTLFSRTHQHLLRGIEQADSVTFDAHKQLYVPMGVGLVVFKTPGLASAVEHHAQYIIRKGSKDLG
STTLEGSRPGMSMLIHSGLRIIGREGYEILIDEGINKARTFAEMITEDADFELVTKPELNILTYRYCPAVVQQALAVANKRQAERINTSLNRIT
KYLQKTQRERGKAFVSRTRLEPARYTHFPCIVFRVVLANPLTTPDILMDILDEQKELARESGIAEEMAIVKELAETVLKEKETSEG
>B_A0A0J8JNG0
MDINHTKIATADMESLHRIFTVAEAPDSTLGKIEKDISENLAGFLNEHIVAREIPLSEIERDFNHAHLPEEPTFVSEHTKFLLDKLVAHSVHTA
APSFIGHMTSALPYFMLPLSKIMIALNQNLVKIETSKAFTPLERQVIGMMHHLVYAESDAFYQDFMHSESDSLGAFCSGGTVANISALWVARNN
ELKASGDFKGVNNSGLYRGLKHYGYDGLAILVSTRGHYSLAKAADILGLGRDDVVAVPVDGEHKVLVDEMKLKANQLKQQNIKVMAIVGVAGTT
ETGNVDPLNDLADLAAELDCHFHVDAAWGGATLFSEKHKSLLKGIERADSVTIDAHKQMYVPMGAGLVLFKSPRLASEIEHHAEYVLRKGSKDL
GSHTLEGSRPGMAMLVYSALNVLGRKGYELLIDRSIEQAAYFANLIQNNADFELVSQPELCILTYRFVPKQIKQLLKQATPKQALLINEHLDDL
TQFIQKSQREAGKSFVSRTRINCARYDNQVITVFRVVLANPLTTQQILQEMLEEQVCLANKSQIALPELIRILES
>B_A0A1H2SAR8
MMKTIHDSSLDEWFLNRTSTSQTAYKQAVGAAVNTVQTQFSSLEKPMSGVQADFLSSLLREDTICPEEGLHLDTVLYEVGQKVLGHSVAVHHPA
CVAHLHCPPLIPALAGEVMVSATNQSMDSWDQSPAATLLEERMIQWLSRTFGLSPQGDGVFTSGGTQSNFMGLLLARDHYAWTRFGWNVQKRGL
PPEAREMRILCSEAAHFTVKQSAALLGLGEDAVVTVKTDERHRIHLDDLDQQLSQLYGDGLHPFAIVATAGTTDFGSIDPLPELVSRAHALGLW
LHVDAAYGGALILSNSYKERLQGIEAAHSITVDFHKLFYQPISCGAFLVNDVKHFNRIKQNADYLNSTEDEELGIPNLVGKSIQTTRRFDALKL
YISLRTLGRAQFASMIEKTINTATATAQMIAKDQRLEVINPVPELNAVVFRFIPEMNHPDMPYSKWQDQINHEIRHALLASGDVVLANTKVNGN
VCLKLTLLNPRTTLEDTHAILDRVKKTGKRRELQKGEWNQHASNIGSASYHAELL
>B_A1AMJ0
MPKKRDAARANLETLYRIFTVPEAPDSTLGAIDQAIADDVIGFLQTHIVAVEHDLEEIEAYYSSPVIPEEPTYVSEYTEFVKRNLVAQSVHTAA
PGFVGHMTSALPYFMLPLARLMTALNQNLVKVETSKAFTPMERQVLAMLHRLVYRRPDDFYPPWIHNSQHALGAFCSGGTIANVTALWVARNRL
FAPSRTFGGIAREGLFRALKHHGCDGIAVLVSERGHYSFGKATDLLGFGRDNLIKVKTDGHNRIDLKLLREEYRRLRDRNILPLALVGIAGTTE
TGNVDPLEAMADFAQEIGCHFHVDAAWGGPTLFSDRYRSLLAGIERADSVTIDGHKQLYVPMGAGMVVFKDPTAQSAIEHHANYILRRGSKDLG
SHTLEGSRPGMAMLVHAGFCIIGRKGYELLIDRGIERARTFAEMVRSHPDFELTSEPELNILTYRFSPAKVQQALARATDEQRPYINALLDQVN
QQLQKHQREAGKTFVSRTRLCVGPRQEEFTVLRAVLANPLTTDEIMEAILAEQCEIARTPEIQALLRQIERRCAGGERDEPLQLQDYPGELDEL
DES
>B_A0A0H4P273
MNFNFQKWFLHPDGKSEKTYRILMDEIMTLICEQTKKAKKPFSGTSHLEIEEKVKEALHIPIAGQEVAKVMAEIQDVIVGDSLWISHPSAMAHL
HCPPLLPSIAAETMIGALNQSMDSWDQSPSATYVEEALVKFFTQKIGYSHEADGVFTSGGTQSNYMGLLLARNKACETYFKVNAHQKGLPYEAN
KLRILCSEHAHFTVQQSAAQLGLGANAVVTVSTDDQQKLSIDDARDKLKKLRREGLIPFMIVATAGTTDFGSIDSISETARLAKDEGLWLHVDA
AYGGALLFSHQYRSLLDGLRLADSITIDFHKLYFQSISCGAFFVKNNQNFHRIAYHADYLNPEEDQEKGMIHLVEKSVQTTRRFDALKLWMTFK
LLGTDLLGQMIDYTIDLAKETASLMKKDPCFEVPVYPEMNAILFRYLPSRTEDRKYVDEMNRKIQQALYENGELIIAKTKQNGKLYLKCTMLNP
LNTIDHMKQHIERMKRLGEKIEKEQGEKRHEYSIHYEPHYDAKLN
>B_A0A1Z3HL75
MQLFNPSAQSQLVEQAIEEKLAVLAQAFLHENLSTTDIDFEHLMERFSDSELPRAPFEFEQYFPSLAEDIVTHAVHTSSPRFIGHMTSALPYFV
RPLAKLVTALNQNAVKVETAKSLSPCERQSLGRMHRLIYQYSTDFYKQHIHNHQSTLGILVSGGTVANITALWCARNNALGARGKFAGVEQEGL
SAALSFYGYKGATIIGSNLMHYSFDKAADLLGIGSQNLIKIPVDDRNHIDLKLLHQAVLHCQSQRKLILAIIGIAGTTDAGGIDPLHDIASIAR
SAKTHFHVDAAWGDPLLFSRQHRHKLAGIEQADSVTIDGHKQMYLPMGIVMVFFRNPHLAKAIEKSASYTIRNGSLDLGKRSLEGSRPGMALFL
HAALNLIGVEGYEFLIDEGLRKTQYMAERIRTMPEFELIAKPDINLLVYQYIPEPFRERATHKQLIKSDYCVISRFNERLQKVQRQVGRTFISR
TVKSIRHFDQEVTIVALRAVLANPLTSETDIDAVLDDQLQIAARLEIETSLEGPVRVV
>B_Q2SN93
MSKTKKIAKASLETMYRVFTIPEAPNSTLGRIDQKISQNLAGFLQDHIVAVEKDLSEMEKDFAESRIPEDPVFVSEQTQFLLDKLVSQSVHTAS
PSFIGHMTSALPYFMLPLSKIMIALNQNLVKTETSKAFTPLERQVIGMMHRLVYDRDDAYYHEWMHNSSVALGSMCSGGTVANITALWVARNLC
FPADSVFKGVRREGMFKALKHYGYEGAAVLVSKRGHYSLSKSADLLGLGSDNIIAIPTGANNKIDLQALRATCEKLRDANVRVISLVGIAGTTE
TGNIDPLEDMAAIAKEFNTYFHVDAAWGGPTLFSNNYKHLLKGIELADSVTMDAHKQLYVPMGAGLVVFKTPSTSNAIEHHAQYIIRQGSRDLG
SKTLEGSRPGMAMLIQSGLKIIGRTGYEILIDLGIEKAKTFAAMIDQADDFELVSEPELNILTYRYHPVWVRQAFEFADEERRQAINDCLSRIT
KGIQKTQRARGKAFVSRTRLNPAAYDGQACVVFRVVLANPLTTVEILQDILEEQRAIAAEDILSERMDELRNLCAIPTAKVAAS
>B_A0A1H3XGS2
MSKRQATASEEALWRIFTVPEAPDSTLSKIEQNISQNLAGFLRESIVAVEKPLWQIERDFQDFQIPLAPQFVSDYADAMMEKLVAHSVHTASPS
FIGHMTSALPYFVLPLSKMMVGLNQNLVKIETSKAFTPLERQVLGMMHHLVYQQDAGFYTKWMHSANHALGAFCSGGTIANISALWIARNRLLG
PDGDFRGIAREGLFRAMKHYGYDGLAVLVSERGHYSLGKATDVLGLGRDDLIAVKTDGNNKVDMAEMRRICAELQAKNIRVLAVIGVAGTTETG
NIDPLHQLADLAQELQCCFHVDAAWGGATLLSEKYRHLLSGIERADSVTIDAHKQMYVPMGAGMVVFKDPSSAHAIEHHAEYILRKGSKDLGSQ
TLEGSRPGMAMLVHACLQVIGRDGYEILINRSLEKARYFAGLVKAHSDFELITEPELCLLTYRYVPAEVQVALHKASPEQREKINELLNSLTKF
IQKRQREQGKSFVSRTRLTPEQYQRQEMIVFRVVLANPLTTQDILHDILQEQTEIARQSRQVYPKLMQWVKELAA
>B_A0A0S2K4U2
MGPKRCAVASDETLMRIFTVPEAPDSTLSKIELEISNNLVGFLNENIAAIEKPLHEVEKDFQSAAIPEQPMFVSDYAQDIMEQLVAHSVHTASP
SFIGHMTSALPHFVLPLSKLMVGLNQNLVKIETSKAFTPLERQVLGMMHHLAYGADESFYGKWMHSAKTSLGAFCSGGTVANITALWIARNRLL
QPQGDFKGINSQGLVAAMMHYGYKGLAVLVSERGHYSLGKAADVLGIGRDNFIAIKTSDDNKVDVQAMREKALELEAQGIKVMALVGVAGTTET
GNIDPLSDMADLAQELNCHFHVDAAWGGATLLSANARPLLAGIERADSITIDAHKQMYVPMGAGMVLFKDPHATDAIEHHAEYILRKGSKDLGS
HTLEGSRPGMAMLVHACLRVIGRQGYEMLIDKGIAKAKYFAELIKQEEDFELVSEPELCLLTYRYVPKAIKQAIANANEDEKIDIYAALNRFTA
SMQKRQREAGRSFVSRTRLTPVQYDNQPTVVFRVVLANPLTSEAMLKEILDEQKQLAKTDPIFKKYLAKYI
>B_A0A0C5WRF6
MAVVDHKKADATQESLHRIFTVPEAPESTLGRIEKEISENLNVFLQTHIAAREKPLAEIEKDFSSADIPESPSFVSDHTHFLLDKLVAQSVHTS
APTFIGHMTSALPYFLMPLSKIMIGLNQNLVKIETSKAFTPLERQVLGMLHNLIYRENDAFYQQWMHSANHSLGAFCSGGTIANITALWVARNS
ALKPDGDFKGVAQEGLFKAMKHYGYEDLVVLVSERGHYSLKKAADVLGLGRDCLIPIKTDGHNRVRVDDMRAKLEELKQNQVKAFAIVGVAGTT
ETGNIDPLEELADLAQEYGCHFHVDAAWGGATLMSNTYRPLLKGIERADSVTIDAHKQLYVPMGAGMVIFKNPALMTAIEHHAEYILRKGSKDL
GSHTLEGSRSGMAMLLYASLNIISRPGYEMLINTSIEKAQYFASLINQDSDFELISEPELCLLTYRYAPARTQEALKLADPEQREELLAALDDM
TKFIQKRQRESGKSFVSRTRITPQAWGRRLTTVFRVVLANPLTTEQILKDVLEEQKDIAKQSLISFPKINTLTESILNQQVTL

>B_A0A0C5WU90
MSLLECRLERELLSTEEESAIAQVGKARELYQLSFSGRLGHDLPRHGNKEKINSGLKLSTSRHRFDRVDIGNQPVAPFEYLDFLETVVFPHTVN
TFSTKFLGHMTAPLPEFITEVAGVISRMNQNQVKVETSNVLTLLERQVLGAVHHKIYRKQAGFYESYMQCPESCLGVVTGGGTLANITSLSYAL
NSAFRADGNFAGLTKEGLVSALNHYGFKDVVVIGSKRMHYSVDKAAKLLGLGERNVIRLETDSHGRVDLAQMRSNLEECRKGKIHVLALIGIAG
ATETGSVDPLEEIGALAAEFGVHFHADAAWGGAYTLSSNHRELVKGIEMADTVTICAHKQLYIPMGTSLCIFKSPGFASHSENNTAYQCKKGSY
DLGRYTIEGSRPASILMLHALLNLWGDAGMGHVFDTTLSLTQNFTGKIKSSENFLLVQEPALNIVVYRYIPVTLREKVLTKEKLTYAEIELINE
INKKIQSEQFINGNSFVSSTTLVNGEEEVVVFRAVFCNPLTRENDLDALLADQAKIARDIEIA
>B_G2PQR2
MHLKENILNQFISPQGTNKELVEKKLQEITTQLLDFLSNAANKPTYPKFQYFGTQHFEIPTEPNSDNQIGLRLQELFDFNMNPANPKYIGHMDS
IPTLWSIIGDYVASAMNNNQLSLEMSPILTQLEYSITRQFATLFGFPNSAGGVMLSGGSLSNLQALIVARNEKLNLNNGNISALQKEPVIFTSE
HSHSSIQKIGMILGIGADNVIKIKADENSKMDVVHLEQQVEEQKKLGKIPFSVVATAGTTVSGNIDPLDDINRIAKENNLWFHIDAIYGGAVIF
SEKYKHLMNGIDNADSISFNPQKWLYVAKTCSMVLFSDFQNMIENFRISAPYMKEQEDFINLGEINIQGTKYAEIVKLWLSLLGLGKKGIQELI
DFSFEMTEKFISEIKKREYLKLVSKPELNLICFRGEPNYIQETEFDEWNKNLQNHLINETDFFISLPRYKDDLWLRTVLLNPFLTEEYIDSLFK
EIDIFDEKYKNQT
>B_A8H648
MTARKATASEEALLRIFTVPEAPDSTLSVIEQNISQNLMGFLQESVVAVEKPLSEVELDFQQYHIPAAPQFVSDYADNMMQTLVAHSVHTSAPS
FIGHMTSALPYFVLPLSKMMVGLNQNLVKIETSKAFTPLERQVLGMMHHLIYNQDETFYQSWMHSANVSLGAFCSGGTVANITALWTARNQLLK
ADGDFKGIAKQGLLKGLRHYGYDDLAILVSERGHYSLAKTADLLGIGRENIIQVPTSDDNKVDVVKMREIAEQLDKDNIKVMAIVGVAGTTETG
NIDPLNALADLAAELNCHFHVDAAWGGASLLSNKYRHLLAGIERADSVTIDAHKQMYVPMGAGMVIFKDPTFANAIKHHAEYILRQGSKDLGSQ
TLEGSRPGMAMLVHACLKVIGREGYEILINNSLEKARYFADLITAEADFELVSKPELCLLTYRYVPQSVQIAMAKAREIGDTATLAQFNGLLDG
LTKFVQKTQREQGTSFVSRTRINPESHQLMDLKAVVFRVVLANPLTSHDILQQVLAEQAQIAKSETHFLPQLLTLAQS
>B_B9M3A1
MLKNREAARANLENLYRIFTVPEAPDSTLGAIDQAIAADVTGFLQTHIVAIERDLEAIEADFASAIIPEEPTYVSEYTEFVKENLVAQSVHTAA
PGFVGHMTSAIPYFMLPLARLMTALNQNLVKVETSKAFTPMERQVLAMLHHLVYRRDADFYPAWIHNSQHALGAFCSGGTIANVTALWVARNRL
FAPSADFGGLAQEGLGRALQHRGAEGIAVLVSERGHYSFGKAADLLGLGRDNLIKVQTDAHNRVDLKLLREEVRRLQDRNILPLALVGIAGTTE
TGNVDPLEALADLAGELGCHFHVDAAWGGPTLFSDRFRSLLSGIERADSVTIDAHKQLYVPMGAGMVVFKDPTALSAIEHHANYILRHGSKDLG
SHTLEGSRPGMAMLVHAGLSIIGRKGYELLMDMGIERARTFAAMIRQYPDFELTSEPELNILTYRYCPLNVQKALAAAPAEQRAGINALLDQVC
QLLQKHQREAGKTFVSRTRLRVARHDEELTVLRVVLANPLTTDEILAAVLAEQCEIVQQPEIQGLLQQVEEICTGLTAKAADQSPN
>B_E3I3F0
MASEKLTEISQHPSPSEDAGADVADYLLGSDLSKLRGWVEARSASFLRSADAGERSPIRRVREKFRSCEVPDTAMALDDYLSLLDQDVLPHCSS
LASPRYLGHMTSPIPGFLPELGRLVQTLNQNVVKMETSGSLTFVERQVLGMLHKETYGFDTAFYDRRVQDRDSTLGLFASGGTIANLTALRAAK
QRASVHAGGGARMAVIGSELMHYSFAKGADLMGLELRRVPVDEQNRMLPAALEREIEACEAEGVTVAALIAIAGTTEFGSVDPLASICRLGQAR
NIHVHVDAAWGGGFLLSPRNRHILAGIELADTVTIDGHKQLMVPLGCGMLFFRDPEVSKTIMHHAPYAVRPNSEDQGRFTLEGTRPATAIYVHA
ALHLIGKSVYDALFSASLERTRIMARHIEGMPEFELTSRPDMNILTYRYIPEAMRGTTPGPADNHRISRFNVALQRAQRDKGDSFVSRTFRPVS
RHGDEPLALLRAVLLNPRTTEDDILFLLRDQVGIARELENSPAFHE
>B_A0A1I0GM92
MTIEKQQAKATEATLNRIFTIPESPETTLGKIERQISENLGGFLNEHIVALEKPLTEIEQDFSGANIPEQPSFVSDHTQHLLDKLVAQSVHTAS
PSFIGHMTSALPYFVLPLSKIMVGLNQNLVKIETSKAFTPLERQVLGMMHNLVYKQADDFYDAHMHSRTHALGAFCSGGTIANLTALWVARNNA
FKAQGSFKGIAAEGLHKALAYYGYDDAVILVSERGHYSIKKSADVLGLGRDNVIAVATDSEQKIDIEQLRKCCESLSKQNKKIMAIVGIAGTTE
TGKVDPLAEMAEVARQYQAHFHVDAAWGGATLLSNNYRHLLAGIDEADSVTIDAHKQMYIPMGAGIVVFKDAEAANIIKHHAEYILRKGSKDLG
SHTLEGSRPGMALLLYASLHIISRPGYELLIDKGIEKAHYFAQLIAQHDDFELMTAPELCLLTYRYVPAWLQQKLKNKNTPLATEELINAKLDR
LTKVIQKQQRESGQSFVSRTRINMERYNNRLVVVFRVVLANPLTSKEVLQEILNEQVAIAEQHQHLLADFT
>B_Q1K434
MTDNAQANLKNLYRIFTIPEAPDSTLGSIDQAITDNLAGFLQEHIVALERDLEEIERDFVDTAIPEQPTFVSDYTEFVKKKLVAQSVHTASPGF
IGHMTSAMPYFMLPLSRIMIALNQNLVKVETSKVFTPLERQVLAMLNRLIYNHDDAFYQRWIHDSQSALGAFCSGGTIANITALWTARNQLCGP
DGDFQGIAQEGLFAALKHLNCEGLVVLASRRGHYSLGKAVDLLGLGRDNLIAVDTDDNSRVDMRQLRNHCYRVQNEGKRVMALVGIAGTTETGN
VDPLEEMADLAEELGCHFHVDAAWGGPTLFSETHRWRLRGIERADSVTIDAHKQLYVPMGAGMVLFKDPAALSAIEHHAAYILRHGSKDLGSHT
LEGSRPGKALLVHAGLSIIGRRGYELLIDQGIARAQGFADRIVAHDDFELVTPPELNILTYRYNPSWLQRVMAQAPEETVKEINILLDQITQMI
QKEQREAGKTFVSRTRLEPARYHGETITVFRVVLANPLTSDDILDAVLQEQCELARQESLRDCFDQLRQIAAGCRVERLAL
>B_K8W5U2
MKETFFIKTLFSPSAPYFQEGIDLCIDAFSQQKDEVNTFNKDVLGNTFSSLTIPEEPASIKNYLSDILETIIPACSHLASPTYLGHMTSPLPSF
IPEIGRLIQTLNQNMVKMETSRGLTLLERQLLKLLHQEIFNCNDDFYADINQDFSKTTGIFTSGGTMANLTAMWVATRQARQNSESRLAVIGSE
LLHYSFDKAAALLGFELYRLPVNQQQKVYVDHIEQQIQACTKQGITVAAIVGIAGTTDFGSIDPLFEIAELGRRYGIHVHIDAAWGGAFILSHK
HKFLLAGIELADTVTIDGHKQMLMPIGTGMLFFKKPELSRVLIHTAPYAVRKTSLDQGRFTLEGTRPANMLYLHACLHLIGKQGYGELFGTAMD
NIKNMASYIKNHPAFELLTEPTINILSYRFIPKRFRNKALTAEDNQQISEFNCLLQKIQRSRGKSFVSRSERKFACHGNKALVFLRVVLLNPCV
EKEHILFMLQDQLDIADDIEQNYDQLKSK
>B_A0A139WW23
MTLSKQEMPNQNYLCETEQTLHYEIEEQVMQLFASSSQTTSIEDKIDNIIHDLSQEFLSTVNANTDIDINSLLGKFCESKIPLEPANFESYLQD
LGKNVITHSIHTSSPQFIGHMTSALPCFVRPLAKLMTAMNQNAVKIETAKALSFCERESLAMLHRLIYNFSDNFYVQHIQNNSSTLGILVSGGT
VANITALWCARNAALGPKDGFGGIEKEGLAAALDFYGYKGAVIIGSELMHFSFDKAADLMGISTHGLIRVPTDCNNRVDIQALRQAVIECRAQN
QLIIGIVGVAGTTDSGGVDSLSEIADIAQEAHVHFHVDAAWGGPLIFSEQHRHKLAGIERADSVTIDGHKQLYLPMGIGMVFMRDPHMAEAIEK
QASYTMRKGSFDLGKRALEGSRPGMALFLHAGLNLLGLKGYEFLIDEGIRKTQYMADRICMMPEFQLLAEPDTNLLLYRYIPEPLRELVAKKQL
TEIDNQLIDQFNERLQKIQRQIGRTFISRTTKTTVSFGKEIPIIALRAVIANPLTTEEDIDAVLNDQIQIASQFEISNFLEAIEKSV
>B_N6WX51
MTGKKKTAHASLEAMYRVFTVPEAPDSTLSRIDQDISRNLAGFLQEHIVAIERDLGEVVRDFNESAVPEKPIFVSEQTQFLLDKLVANSVHTAS
PSFVGHMTSALPYFMLPLSKIMIALNQNLVKTETSKAFTPLERQVLGMIHRLVYNEDSAYYRKWMHDPRYALGAMCSGGTIANLTALWVARNQA
FPAEGSFRGLHEEGLFRALRYYGYEGAAILVSRRGHYSLRKAADVLGLGRNALIPIDTDDENRIQTDALRDKCLELQKQKVKILAICGIAGTTE
TGNVDPLDAMADIAREFGAHFHVDAAWGGPTLFSRTYGPLLKGIEQADSVTFDAHKQLYVPMGAGLVVFRNPELVGAIEHHAQYIIRKGSRDLG
STTLEGSRPGMAMLIHSGLKILAREGYELLIDQGIEKAKLFAEMIDRREDFELVTRPELNILTYRYCPAATQDALSRADTLQSERINASLNRIT
KYIQKTQREHGKAFVSRTRLEPAKHHHFPCVVFRVVLANPLTTPEILRDILDEQTQLAKESELQDEIEMLENLTQSVPPLTHTREAL
>B_K9U459
MDLQHFLNHNPESLAQYAQTVQQTQKLIQDCLSQQERVFSGLSPATLSEFLTQKILPEQGIPTQQVLQEAVEAIFSHSIAVHHPDCVAHLHCPV
LIPSLAAEMLISAFNQSMDSWDQSGAATLLEQNLINFMCQLYGYDANADGTLTSGGTQSNFMALMLARDYALQQRGFTSQLHGLPNLAGRFRIL
CSEVAHFSVQQSAAILGLGMNAVVRVKVDKNYRLCSQHLVQCLEDIDRQDLIPICIVATAGTTDFGSIDPIAEMTAIAHEHNTWLHVDAAYGGA
LMLSDRHRQKLAGIHQADSITIDFHKLFYQPIPCSLFLLKDKSRFELMRLNVAYLNPEHNEDEGIPDLVTKSIQTTRRFDAVKPYIAFRALGRE
FFAGVVDRGIEITQQIAAHIEQDPQLELAVSPEMSTVVFRYRSTNAVNLGIKQALLQTGKAVIGQTEIQGKAYLKFTLINPLVTFDRTVALVEK
IKRLGTMLSESPHFLQPAIEAGVGNQKSKVKS

>B_K0NEB9
MVKIRNKQYPLVADWNTLNRVFIRPEDEIGRKTLVKYMEQILFGLHDFLNKHVGVTEEISLIKLAEEYTDTKINAHPQKKLADVIKDIINEIAP
RAVNVASPYFIGHMTSAIPFFMVHLKAITAALNQNVIKLETSKVLSVLEKQILAKMHRMIFNFDDHFYLEHVQNTETSLGNFTTGGTTANLTAL
WVARNKCFPSKGNFESIEKNGIFEAMKTHDLEKAVVLVSKRGHYSLRKAGGVLGLGNQNVIPIDVDANRTIDIIKLKSKIKELQANKKTKIVAI
IGIAGATETGIIDPLPKLADICQKHGIHFHVDAAWGGPVLLSEKYCHLLKGIERADSVAIDCHKQFYMPMTSGMVYFKDPTAMDHIVYHSSYVN
RPGSVDLGIKTLEGSREANSLILDSALKIMGSKGYALMIDHGIETAREFAKMISQRSLFQLVTKPQLNILTYRLVPLDIQKKLETADKEERQKL
NYILDDINIKIQRIQREAGKSFVSRTRLKNSPDDYNNNVVLRSVIMNPMTNIDIINEVLDEQESIFHKLVNE
>B_A0A1I4JUD5
MATNAHVAPSEIAKNYDPYFLHESEESIALYQQLMQQTQMALVHNFSKSKKPYSGFSHRELKALLNKTFSETVPEKPRNMSELINDIGEVILKH
SIAVTNPSCIAHLHCPPLSIALAAEVMISGTNQSMDSWDQSSAATVLEQKVIHWLSQMFLQTEQADGVFTSGGTQSNFMGLLLARDRYLYKQLN
WNAQEQGLPPEANRLRIFCSKDAHFTVRQSAFLLGLGEQAVIPLETDHQHRICLSDLDHRLKEVKEENLLPFALVATAGTTDFGSIDPLNELAN
RVQAHQMWFHIDAAYGGALILSDRHKEKLNGIHLGDSITVDFHKLFYQPISCGAFLVKDQSNFNFIKLNADYLNPEDDEETGIPNLVTKSIQTT
RRFDSLKLFMSLQALGRKTFADMIDYTIDLAHDTAGLIEQDIELELMNQPELNAIVFRYANHSYDENVVNKINTIIRSKLLSQGHAILARTRVN
GCIYLKLTLLNPRTTLKDIQSILSQVKKIGYQCYQNGGV
>B_A8ZVT2
MPRRTDRRDKIGRNGKPMRNTANRNTATTQAGSRPLIADWNALMRVFIRPEDEACRTTLIKYMEQILFGLHEFLAQHVGITEDVSLKALARQFK
STAIAENPEKRLAEVIQELIREISPYGVNVASPYFVGHMTSALPFFMVHLKTIAAALNQNLVKLETSKIFSLLEKQVLAKIHRLIYRFPEQFYC
DHVQNPETTLGSFVEGGTTANITALWVARNTFFAPRNDFAGIEADGSAAAFAEYGVKRAVILVSRRGHYSLRKAGGILGIGNKNIVAIDVDKEH
RLDIGCLEQTIADIRKQGDTAIVAVVGIAGATETGAVDPLRKIGELCAREKLFFHVDAAWGGPTILSDTHRHLLDGIELADSVTIDGHKQFYMP
MSCGMVYFKNPHIMDAVAYHANYVNRHGSVDLGIRSLSGSREANSLVLDSALKIMGKKGYALLIEHGIDTARKLAEEIERRDLFEVVTWPQLNI
LTYRVCPAGLKEIMQTGTDEERIEAVHALNHINKTVQRIQREAGKSFVSRTTLMRRHYEKGIVVLRCVIMNPLTTMAVLNDILDEQESIFRRYF
PSPPPVYPS
>B_W0PGU5
MNVVLNSGADSATDSINEAQTPQALIDALFLANTTVIRNRVDESCRTFFEASDAIDEMGIEGIRSKFMQCDAPTQGTTAEAYLDMFDNEVLPHC
SHLASPRYLGHMTSPLPQFLPEIGRLVQTLNQNVVKIETSRGLAFLERQVIGMLHREIFGADRQFYENHLQARSSALGIFTSGGTLANISALWT
AMRRAGTTGTAENSSPHADKPVIIGSALMHYSFDKGAQLLGAELHKVPVNDHCQIDLDALREAIDKYQQAGRKIACLVAIAGTTDYGSIDALDK
IGEIGRQINAHVHVDAAWGGGLILSESNKHLLDGIAQADTVTIDGHKQLMLPLGCGMLFFRDPALSQLTMHHAPYAVRATSFDQGRFTLEGSRP
ATALYLHAALHIIGKNGYDALFSESLARARYMAQAIEARPEFELMTAPVMNILTYRYIPEKYRNQHIDDAGNIEINRFNIALQKWQRQKGDSFV
SRTFRAINQYDNQGLTLLRAVLLNPLTTFEDIDFLLNDQLQIAHLLENEMAGAPATH
>B_Q487K9
MKATKRIAEATQESLHRIFTIAEAPDSTLGRLEQEMSQNLVGFLNNHIVASKNALTDIEQDFINARIPEQPEFVSDHMHHLLDKLVAQSVHTSS
PSFIGHMTSALPSFILPLSKLMVGLNQNLVKVETSKAFTPLERQVLGMMHNLVYQHDDVFYKNWMHSAEHSLGAFCSGGTVANITALWVARNKL
LKADGDFRGVAREGLHRAMRHYGYQDLAILVSDRGHYSLKKSADILGIGQENVIAIPTDEHNKIDCQKLADKCQQLAAQNIKVLAIVGVAGTTE
TGNIDPLDKIAEIAQQNQCHFHVDAAWGGATLLSNKYRPLLKGIEQADSVTIDAHKQMYVPMGAGLVIFKDPASVSAIEHHAEYILRKGSKDLG
SHTLEGSRPGMAMLVYSSLHIISRPGYEMLINQAIEKAEYFADIIHQHDDFELITRPELCLLTYRYAPKSVQALLARNDDEANKSVNMLLGKLT
KFIQKRQREDGRSFVSRTRIEVSRYGGEKVIVFRVVLANPLTTKEILQGILQEQCQLAQESEQFLPELLQAAK
>B_Q12LG2
MMAQLMTSITPRKAQASEDSMLRIFTVPEDAESTLSIIEQKLSEDLAGFLGNSIAALEKPLSEIETDFQAFQIPTAPRFVSDYTDEIMQNLVAH
SVHTAAPSFIGHMTSALPYFVLPLSKMMVGLNQNLVKIETSKAFTPLERQVLGMMHHLVFSQDGDFYQQWMHSANHSLGAFCSGGTVANITALW
IARNRLLKADGNFNGVNREGMLKALRHYGFDDIAILVSERGHYSLGKAVDLLGIGRDNIISIATTADNKVDVEAMRLAAKALASKNIKVMAVVG
VAGTTETGNIDPLTELAALAQELDCHFHVDAAWGGASLLSKKYRHLLKGIELADSVTIDAHKQMYVPMGAGMVLFKDPELANAIVHHAEYILRV
GSKDLGSQTLEGSRPGMAMIVHACLQIIGLEGYEILINNSLEKARYFSELIKKSPDFELITEPELCILTYRYVPSTVQLAMSDACQRQDWHALA
QFNECLDGLTQFIQKHQREQGKSFVSRTRIQPARYNRQHTTVFRVVLANPLTGLEILQGVLAEQQEIAALDNEFLPQLLALASA
>B_A0A1I6JRS3
MTGKKKSAQASIEAMYRVFTVPEAPESTLSRIDQNISGNLAGFLQEHIVAIERDLSDVEKNFSDSAVPEKPIFVSEQAQFLLDKLVANSVHTAS
PSFIGHMTSALPYFMLPLSKIMIALNQNLVKTETSKAFTPMERQVLGMIHRLVYQQDGAYYRKWMHDPRHALGAMCSGGTVANLTALWVARNRA
FPAEGSFRGLHQEGLFRALKYYGHDGAAILVSRRGHYSLRKAADVLGLGRDSLIPIDTDDENRIQIDALRDKCLELQRQKVKIMAICGIAGTTE
TGNVDPLDAMADVAREFGAHFHVDAAWGGPTLFSRQYGHLMRGIEKADSVTFDAHKQLYVPMGVGLVVFRDPTMASAVEHHAQYIIRKGSRDLG
STTLEGSRPGMAMLIHSGLKILAREGYEILINQGIEKAKTFAAMISDDPDFELVTEPELNILTYRYCPERVQQALAAADPLQAERINMSLNRTT
KFIQKAQRERGKAFVSRTRLEPARYSKFPCIVFRVVLANPLTTKEILQDILEEQKELAKDPRLADEMEILHQHAEAALADAPAKRA
>B_Q6LS17
MAVDNRKADATLESLHRIFTVPEAPESTLGLIEQKISQNLNEFLGTHIAAREKPLIEIEKDFSSAEIPENPSFVSDHTQFLLTKLVAQSVHTSA
PSFIGHMTSALPYFLMPLSKIMIGLNQNLVKIETSKAFTPLERQVLGMMHNLIFNEETSFYQQWMHSADHSLGAFCSGGTIANITALWVARNNV
LKPDGEFKGVANEGLFRAMKHYGYDDLAILVSERGHYSLKKAADVLGIGRDCLIPIKTDDNNRIRVDDLTATLQSLKEKNIKPFAIIGVAGTTE
TGNIDPLDELADIAQAHDCHFHVDAAWGGATLMSNKYRSLLKGIERADSITIDAHKQLYVPMGAGMVIFKNPALMTSIEHHAEYILRKGSKDLG
SHTLEGSRSGMAMLLFASLNIISRQGYEMLINNSIDKAKHFANMVKQDPDFELVTEPELCLLTYRYVPAETQQALTIANAEDKHLLHDALNDLT
KFIQKRQRESGKSFVSRTRLTPEKWDHKITTVFRVVLANPLTTDNILQDVLIEQKELAKESVHSLPRLFTLTKRILANN
>B_A0A0F4P522
MGPKRCAVASEESLMRIFTVPEAPDSTLSRIEQEISSNLAGFLNENIAAIEKPLHEVEKDFQSAVIPEVPMFVSDYAQDIMEQLVAHSVHTAAP
SFIGHMTSALPHFVLPLSKLMVGLNQNLVKIETSKAFTPLERQVLGMMHHLVYGSDDAFYDKWMHSAKTSLGAFCSGGTIANITALWIARNRLL
KADGEFKGIGASGIVAAMLHYGYRGLAVLVSERGHYSLGKAADVLGIGRDNFISIKTAGNNKVDVDAMRAKAHELEAQGIKVLALVGVAGTTET
GNIDPLEEMADLAEELNCHFHVDAAWGGATLLSNNNRHLLKGVERADSITIDAHKQMYVPMGAGMVLFKDPAASDAIEHHAEYILRKGSKDLGS
HTLEGSRPGMAMLVHACLRVIGRQGYEMLIDKGIEKARYFAELIKQTPDFELISEPELCLLTYRYVPEQIKAAIAQADEQEKIEIYAALNRFTA
SMQKRQREAGRSFVSRTRLTPEQYDRQPTVVFRVVLANPLTSESLLKEIMDEQRQLAMSDPVFKKYLAKYIK
>B_A0A254TI57
MEELLKTYFSAQDLDENAASLQQALQLIREWLFVDKRHAYPPDDFESLKKAFNTPAMPQDGSAFPAVLDALQQQVFAHSVPVNHPKYIGHMTQA
LPWVSVLVESFIATLNQNQVKIETAYSSTLIEKQVLGWMHRQIYRFDDAFYRDSFESNESTLGNMVNGGTMGNLTALSVALESQMPGLRKKGFL
TLLREKDYTGFAVIASARVHYSVKKSLATLGLGEDALCVIPTDENNRIRLDVLEERIAELKQRKVKILALIGVAGTTETGSIDPLEELGRIAQR
EDIWFHVDAAWGGALLLAEKYRDMFHGVEMADSVVIDGHKLFWVTMAQGMVLFKSDKSLNHLLHTANYIIRKSSGDLGRTSLEGSRRFDALKLW
FSFKLFGLKGYEALLDRCQALAANMRELVMDDPDFELITCSDTFILTYRYAPRELQARLKDSVRTGDRENAVRCNETLNRLNTALQDEQKRRGL
SFVSRTLLESTPYPGQTTVLRVVLTNMLTTQEDLKEILAEQKEIGAGL
>B_A0A1M5I3T2
MGEAQVSLEHLFRVFTKPEHKDSKLAQIEQHLSDNILDFLSQHVVTKKTSLEEVEQDFADAAMPESPEFVSTHAENLLEKLVAHSVNTYSPTFI
GHMTSALPYFHLSLSKLLVGLNQNLVKIETSKAFTPLERQVLGMMHNLVYEQSEPFYEQYLHSARHALGAFCSGGTVANITALWVARNKLLGPD
SGFSGVARAGLASALRHYNINNLGVLCSKRGHYSLSKAVDLLGIGREGLLTLPCPDQVLDPEKALQAGKAYQEQGNRLLAIVGIGGTTETGHVD
PLDELADVAKELGCWFHVDAAWGGATLFSQSYRYIMKGIERADSVTIDAHKQMYVPMGAGMTLFKNPEHANAVRHHAQYILREGSKDLGATTLE

GSRNGMAMMVYSSLHILGRRGYELLINQSIDKAKQFAKMIDEHPDFELVTSPTLSLLTYRVCPQSVQQTLLALPAGKAQRLNGKVDRLVVNVQK
LQREAGKSFVSRTRLESPTYGAYPITVFRVVLANPLTTVADLKAILQEQHDIAMRNRVWQDLTLSGEPLANVVNN
>B_A0A1H3YWG2
MLIWYQIMTDTALANLENLYRIFTVPEAPDSTLGAVDQAISENVADFLQKHIVALERSLDDIENDFTASKIPEEPSFVSDYAEFVKQKLVAQSV
HTAAPGFVGHMTSALPYFMLPLSRIMTALNQNLVKVETSKAFTPMERQVLAMMHHLIYGQTEGFYNQWIHDSQSALGAFCSGGTIANTTALWVA
RNHLCAPDGEFSGIAKAGLFKSLQHLGCDGLAILVSRRAHYSFGKAVDLLGIGRDNLVLVDIDYDSRIDMQALRAEYLKLQQQNIRPLAVVGIA
GTTETGHVDPLNELADFAAEIGCHFHVDAAWGGPTLFSDRHRHLLAGIERADSVTIDAHKQLYVPMGAGMVLFKDPLALSAIEHHANYILRHGS
KDLGSHTLEGSRPGMAMLVHAGLSIIGRKGYELLIDLGIERARHFADLINQHPNFELITAPELNILTYRYNPAWLQDKMGQLDKDQQQKVNCLL
DLVVQEVQKRQREAGKTFVSRTRLLSALYPDGQISVFRSVLANPLTTDEILSVVLQEQCAIVKETEVQNLMTEIEQICG
>B_A0A244CNG4
MEAKRYAVASEESLMRIFTVPEAPGSTLARIEQEISSNLAGFLNENIAAIEKPLHEIEKDFQLANIPEEPTFVSDHAQQIMDQLVAHSVHTAAP
SFIGHMTSALPHFVLPLSKLMVGLNQNLVKIETSKAFTPLERQVLGMMHHLVYGQNDGFYQKWMHSAKNALGAFCSGGTVANISALWIARNRLL
RPDGEFKGIASDGLMAAMMHYGYNGLAVLVSERGHYSLGKAADVLGIGRKNFIGIATDSNNKVDVAAMREKAHELEAQGIKVMAIVGVAGTTET
GNIDPLHEMADLSEQLGCHFHVDAAWGGATLLSNNYRHLLAGVERADSITIDAHKQMYVPMGAGIVLFKDPEASNVIEHHAEYILRKGSKDLGS
HTLEGSRPGMAMLVHACLSVIGRKGYEMLIDKGIKKAQYFAELINQDPDFELISEPELCLLTYRYVPAKIQQILRSADEETRMDIYIALNRFTA
SMQKRQREAGRSFVSRTRLTPMQYQQLPTVVFRVVLANPLTSETILKNILSEQKELAQSDPIFKKYLQKYLA
>B_A0A0C4Y9E7
MRWFDADRGAFESLEQWIAEHPADFFASERFDPVGACATREAVFASVELPETPTSPQAHADHLLHDVFRHVMPVASPTFVGHMTSSLPSFMPSL
AKVVAALNQNVVKLETSGALTGLERQVIGMLHKLVFAQDSAFYGRWLHDADHALGAICSGGTVANLTALWASRNKLLGACDGFAGIHRAGMVAA
LRHYGHDGLAIVVSERGHYSLRKAADVLGIGRDNLVPVAVDSDGRMRIDLLRDTLRDLQRRNIRPMAIVGIAGTTETGAVDPLDAIADVAQEAG
CHFHVDAAWGGATLLSERERWRFAGIERADSVVIDAHKQFYVPMGAGMVLFRSPAWTQEIIQHANYIVRKGSVDLGRHTLEGSRGAAAVMLYAN
LHLLGRKGLAQLIDRSIDNARYFASLIARQPDFELNSRPQLCILTYRHVPETVHAALATACAERRDKILDALDALTISIQEMQRDAGRSFVSRT
QLMSAQWGGRPIAVFRVVLANPDTTHAILQDILDEQRALAAASPCIAPLMALVAAPDAA
>B_Q5E6F9
MVTDNKTADASFESLLRIFTVPEAPDSTLGIIEKELSQNLNQFLREHIVAEEKPLTEIEKDFTDSSMPESPTYVSEHTEHLLDTLVSQSVHTSA
PSFIGHMTSALPYFLMPLSKIMIALNQNLVKIETSKAFTPLERQVLGMLHRLIFGQKDSFYQHWMHSADHSLGAFCSGGTIANITALWVARNRL
LKPEGDFEGIAKQGLFAALMHYKCNGLAIFVSERGHYSLKKAADVLGIGQDGVIAVKTDNNNRVCLDDLELKIAQAKAKNIKPLAIVGVAGTTE
TGSIDPLRELANVAQREGCHFHVDAAWGGATLMSNTYRHLLDGIDLADSVTIDAHKQLYVPMGAGMVIFKDPELMSSIQHHAEYILRKGSKDLG
RHTLEGSRSGMAMLLYSCFNVISRPGYELLINQSIEKAHYFADLIQQQDDFELITEPELCLLTYRYVPSNVKAALAIATDEQKIEIYEHLDNLT
KYIQKTQRETGKSFVSRTRLTPEAYQHQPTIVFRVVLANPLTTKEILQNVLIEQREIASSSEISLPLLNQIVGNILH
>B_A0A0F7JX37
MTPGKASDQKHAVANLSNLYRIFTVPESPDSTLGRIDQEISQNLNGFLRNHIVATEQDLTIIEQGFSSPRIPEQPTFVSDYTEFLLDTLVAQSV
HTAAPGFVGHMTSALPNFMLPLSRIMTALNQNLVKIETSKAFTPMERQVLGMLHALIYGKDDAYYQHWMHDPECALGSFCSGGTLANITALWVA
RNNLFPADDGFQGIAKEGLYRAFRHYGCEGAVVLISQRGHYSFGKAADLLGIGRDSIIAIPTDADNRIDIGELQATCDRLKEQGIRIISLVGIA
GTTETGNVDPLDKLADIAEQQGCHFHVDAAWGGPTLFSETYKPLLKGIERADSVTIDAHKQLYVPMGAGMVLFKNPASLSSIEHHAEYILRKGS
KDLGSHTLEGSRPGTAMLVHSALNIIGRKGYELLIDKGIGIARTFAAMIDADADFELITEPELNLLTYRYVPAPLQALLPKLPAGELAAVNREL
DALTMQIQKVQRAAGQSFVSRTRLSRPQYDNDTITVFRVVLANPLTTPEILREMLAEQKQIATEDPEAAAIVAGLLESVAED
>B_B7VPR7
MVTEQKTADVSFDSLLRIFTVPEGPDSTLTQIEDKLSRNLNQFLREHIVAEEKPLREIEKDFSNAHIPEQPEFVSEHTEHLLDSLVSHSVHTSS
PSFIGHMTSALPYFLMPLSKIMIALNQNLVKIETSKAFTPLERQVLGMLHRLIYQDSDQFYSRWMHSANHSLGAFCSGGTIANITALWVARNNA
LKAQGSFKGVEKEGLFKAMKHYGYEGLAILVSERGHYSLKKAADVLGIGQEGLVSVKTDNDNRICTDDLRLKIEQLKQNKIKPFAVIGVAGTTE
TGNIDPLRDIAEVCAESDCHFHVDAAWGGATLMSNNHRHLLDGIELADSVTIDAHKQLYIPMGAGMVLFKKPDAMTAIEHHAQYILRKGSKDLG
SHTLEGSRSGMAMLVYASMHIISRPGYELLIDQSINKARYFADLIKNQNDFELVSEPELCLLTYRYVPESVKAALLKAKAPERVELNELLNELT
KFIQKKQRETGKSFVSRTRLNPEIWAHQPIIVFRVVLANPLTGKDILSSVLEEQREISKLAPNLMSKITKLVKLINA
>B_D4ZAE7
MSMTPRRATASEEALLRIFTVPEAPDSTLSVIEKNISQNLMGFLQESVVAVEKPLSEIELDFQQHQIPSAPQFVSDYADEMMKTLVAHSVHTSS
PSFIGHMTSALPYFVLPLSKMMVGLNQNLVKIETSKAFTPLERQVLGMMHHLIYSEDDKFYKNWMHSANYSLGAFCSGGTIANITALWTARNQL
LKADGDFKGVSAQGLMKGLRHYGYDDLAILVSERGHYSLAKTADLLGIGRDNIIQVPTSSDNKVDVDKMRAMAKQLDLDNIKVMAIVGVAGTTE
TGNIDPLDELATLAVELNCHFHVDAAWGGASLLSNKYRHLLKGIERADSVTIDAHKQMYVPMGAGMVIFKDPAFANAIKHHAEYILRKGSKDLG
SQTLEGSRPGMAMLVHACLQIIGRDGYEILINNSLEKARYFAELIHGQDDFQLVSEPELCLLTYRYVPKSVQEAMQETRESGNTEKLIEFNSLL
DGLTKFVQKRQREQGTSFVSRTRINPESRASLNIQSVVFRVVLANPLTTREILQQVLAEQIEIAQQDNEFLPQLLALASN
>B_Q87QB3
MVKEQKTADVSFESLLKIFTVPEGPDSTLTKIDESLSRNLNQFLREHIVAEEKPLREIEKDFSSAQIPEQPEFVSDHTEHLLDTLVSHSVHTSA
PSFIGHMTSALPYFLMPLSKIMIALNQNLVKIETSKAFTPLERQVLGMLHRLIYGQNDKFYRKWMHSANHSLGAFCSGGTIANITALWVARNKA
LKADGAFNGVEKEGLFKAMKHYGYEGLAVLVSERGHYSLKKAADVLGLGQEGLVAVKTDANNRIVVDDLKTKIAELKEQNIKPIAVIGVAGTTE
TGSVDPLSQIAQVCQEHNCHFHVDAAWGGATLMSNHYRHLLEGVELADSVTIDAHKQLYIPMGAGMVLFKDPDAMKSIEHHAQYILRKGSKDLG
SHTLEGSRSGMAMLVYAAMHIISRPGYELLIDQSIEKARYFADLIKQQDDFELVSEPELCLLTYRYLPPLIREALDKAEGTQKEKLNELINQLT
QFIQKRQRETGKSFVSRTRLNPDQWQRMNTIVFRVVLANPLTTRDILSSVLDEQREIAKQAPSLTAKIEEMASDILGA
>B_A0A1G6Z257
MTEQAKANLENLYRIFTVPEAPDSTLGRIDQALTADLAGFLQQHIVATGCSLEEIERDFSSTQIPEEPTFVSDYTEFVKNKLVAQSVHTASPGF
VGHMTSAMPYFMLPLSRILIALNQNTVKVETSKAFTPLERQVLAMLHRLVFAQDDAFYQRYIHDSRCALGAFCSGGTIANTTALWLARNRLCKP
DGMFRGIAQEGLLRALQHLRCEGLVVLASRRAHYSLAKGVDLLGLGRDNLIPVETDENNRIDLRCLRQHCQRLQDENKRVLALVGIAGTTETGT
VDPLDALADLAAELECHFHVDAAWGGPTLFSQTHRGLLKGIERADSVTIDAHKQLYAPMGAGMVLFRDPRAVSAIEHHAAYILRPGSKDLGTHT
LEGSRPGKALLVHAGLSIIGRKGYELLIDQGIERARHFADCIRAHPDFELVTAPELNILTYRYNPAWSQQAMAQASQEQRRVANVLLDQLTVTI
QKAQREAGNTFVSRTRLEPAQYQGDSITVFRVVLANPLTTDALLEAVLAEQCGWARRFAELLAPLQALQAGWRQA
>B_W0V410
MTDTRMDQLLQQHFSIDTLKQAPAPLHKALRMLGGWLAAQRSQPYPSTPHAELLPQFTEIDPPAGGMPVADFLDLLDSKVLTHTAQLNHPMYIG
HMTQALPWVSVLAEAFTAALNQNQVKIETAYVSTLIEKQLIGWLHRQVYRRGDAEYAASMRDTGHALGNVVNGGTMGNLTALAVALEHQLPGTR
KQGLFATMTQSGYRGLAVIGSARSHYSLKKSLATLGLGEAALHLVPVGRDNRIDIAALQEKIAELKQAQIKIVAMVGIAGTTETGAIDPLAAMA
EIAEREQIWFHVDAAWGGALLLAEQFRPLYDGIARADSVVIDGHKLLWVPMAQSMVLFKDRNSLNQLKHNANYILRSNSGDLGQTSLEGSRRFD
ALKLWTSFKIFGVDGYTALLQQAATLTGQMRELLADQQDFELMTDSDTFILTYRYIPQQWREQLAGLLAHGNMEAALLLNQKVNELNAKLQNRQ
KAQGTSFVSRTVLESTAYPGQTTVLRVVLSNVRTRPRHLRAILAEQRQLGTLLLAEPGKQLNT
>B_F4KV58
MTQQNNPQKAAPLEGLSLLQHVYDPATFRQQGHALIDLLADHLEAVQHQQDPTVMPYQNPEESYTYWQQELLAPLLNDPLPLFEAVIKRSVKVH
HPHYLGHQVAVTAPAAALAGLVSTVLNQGMALYEMGMVSVPMERLVTELLAQKIGYDSSSNGILTSGGTLANLTALLTARSMKAPSNVWTEGHQ
ERLAIMVSEEAHYCVDRAARIMGLGSAGIIKLPTDERFKMRTDLLEQYLTQAKSKGLHVFAIVGSCCSTSTGSHDDLVAIADFAERHNIWFHAD
GAHGGAAVFSQKYRHLLNGMERADSVVVDFHKVLMTPALATALVFKKGSDGFNTFQQRAQYLWNSSEADWYNPGKRTFECTKYMMSLKIFVLLR

```
LYGEAGFAAAVERLYDLGQKFAAMISTRPDFELAMPPECNIVCFRIVKNGVADLNAYNLQCRQKLLEKGHFYIVQTTLRDVVYLRISIMNPLTT
ENDFVLLLDELTAIQHES
>B_K9TK94
MLQIHPVNSSIETWNPSIEGTILSLFDPSPQAENWEKNVDQQILKIAYQFLRQIEASSDIELPDLMAQFQEFHLPDDASSFDTYLESLTKTLIP
HSIHTSSPRFIGHMTSALPHFMRPLAKLMTALNQNLVKTETAKAFTPYERQALGILHRSLYQFPDEFYDYHLQNSHSTLGMVVSGGTSANFTAI
WCARNSSLGPKGDFKGIEKEGLAAALEFYGYQGQVIIGSSLMHYSFEKAADLLGIGSQGLIKIPASRHNRIDLDCLRETVAECRAQKQHIIAIV
GIAGTTDSGGIDPIEEMAAIAQAAGVHFHVDAAWGGPLIFSQQHRHKLAGIEQADSVTIDAHKQLYAPMGIGMVMFQNPQLAKAIEKHACYTVR
EGSADLGQRSLEGSRPAMSLFLHAGLHAIGLKGYEFLIDEGIRKTQYMAEQVRLSPEFELLAEPEINLLIYRYIPEQLREFVAKGELTETQNQA
IDLVNEQLQKAQRHAGKTFIARTITQTTRYGPMAIVALRAALANPLTTEADIDAVLDDQIALATQLNLGGSQF
>B_A0A1M6AT35
MGKIFAMKKNVLENVYDPSVFRTIGHELIDLLADHLEKVQSDREHPVLPYKDPEVVLDYWKKDFSSDTGVMDMFGNILSQSINVHHPRYIGHQV
AVPAFISSLSGLISDVMSNGTGVYEMGMAANAIERIVTDLVAQRIGYDQNASGLLTSGGSLANLTALLAARKAKAPSSVWEKGHQEKLAILVSE
EAHYCIDRAARILGLGEEGIVKVPVNGDYSIDTSKVAGCLEEAQSKGLHVIAIIGCACSTATGSYDDLDYLGDFADKHNLWFHVDGAHGAGVIF
SEKYKHMVKGIAKADSVVLDFHKMLMTPALNTALIFKNSIDSYRTFEQKAQYLWDSQHSQEWYNSGKRTFECTKLMMSIKVYAILKAYGEEAFT
QNVNTLYDLAETFSEMIDKQESFELAHHPQANIVNFRYAPEQVDIDLNTLNNNIRQALIQSGKFYVVQTTINEEKYLRTTMMNPLTTKDDLIAL
IEEVHHLGTTFVKEKVSNSF
>B_A0A0D8CQ70
MTTGKRKAKATQESLHRIFTIPEAPESTLGLIEKEISENLAGFLGNHIVATEKPLTDIEQDFACSQIPEEPEFVSDHMHHLLDKLISQSVHTSS
PSFIGHMTSALPYFILPLSKLMVGLNQNLVKIETSKAFTPLERQVLGMIHRLVYRDEEDFYQRWMHSANHSLGAFCSGGTVANLTALWVARNNL
LKPDGDFKGIAREGLFKALKHYDCDGLAILVSERGHYSLKKSADVLGIGQESVIAIPTDEDNRIDCQLLRQKCRELTENNIKILSIVGVAGTTE
TGNIDPLNEMADIAGEYGCHFHVDAAWGGATLLSEKYRPLLNGIERADSVTIDAHKQMYVPMGAGLVVFKNPSSVTAIEHHAEYILRKGSKDLG
SHTLEGSRPGMAMLVYASLHIISRPGYEMLINQGIEKAAYFAGIINQHPEFELISEPELCLLTYRYVPPGIQELLRNGSQEQQQEINGLLGKLT
KFIQKRQRENGKSFVSRTRIEVSRYGGEKILVFRVVLANPLTSKEILHDILQEQTELAQESENFYPQLKAML
>B_A0A0C1NBK7
MTLSKKETTPNYCLEVESALKYGIEEKVMQMFTPLSHATSIGKKMDVRVATLAENFLEVNNTSTDIDIDSLVEKFADSNVPIEPSNFDSYFEYL
ANNVVSHSVQTSSPKFIGHMTSALPYFVQPLAKLMTAMNQNSVKIETAKALSFYERQALAMMHRLIYQFGDRFYAQHAQNSESTLGILVSGGTT
ANITALWCARNAFLGPKDGFLGVETEGLTAALDFYGYKGAVIIGSELMHYSFDKAADLLGIGTRGLVKIPANREGRVDISALRQAVAQCRLQNQ
YIIAIVGIAGTTDSGGVDALSDIAEIAQQNNIHFHVDAAWGGPLIFSKQHGHKLAGIERADSVTIDGHKQLYLPMGIGMLFLRDPHLAKAIEKQ
ASYTMRKGSFDLGKRALEGSRPGMALFLHAGLNLIGSQGYEFLIDEGIRKTQYMADRIRSMPEFELLAEPDTNLLIYRYIPEPLRGLVAKKQLT
EIDNLLINEFNENLQKNQRQTGHTFISRTTKIISEFDKKIPVIALRAVIANPLTTEDDIDIVLNEQIEIALKLQFQFL
>B_A0KNW7
MKLTELFALPDPAFGQAMTDLLETFHHSDDSQAPYQRDQFTHQLDQSRMPATGIAMDEYLARLATLVPGASHLTSPRYMGHMTAPLPAFTAELS
RLVVMLNQNPMKMESSRLLSFLEREVLAKLHRLIYRADDLFYDAQMHAKDAALGVMTSGGTIANVTALWLARNRACGDNLFALYEQGYRGAVIL
GSRLMHYSFDKGMDLLGLGARSVWRLPTDDQNRLDMAALEQALAQCKEQKLKVLALVGVAGSTDFGSIDPLPELADIAEREQIHFHVDAAWGGP
TLFSPRYQNLLAGIERADTVTLDGHKQLLVPLGTGMLLCRQPDLMMAVKREAPYAIRASSFDQGRFTLEGTRPANALYLDAAFQLFGQQGYAEL
IEANYDRARLMASLIDQDPAFELMSAPVMNLLSYRCIPPHLQGKVLDEAANEQVNAFNVALQKAQRAEGHSFVSRTQRAVGRYGPQPLTLLRAV
LLNPLIEERHIRDLLADQQRLGAEIARQLFG
>B_B8I983
MELTFNPQNFSPEETQSGETKKEILDLFSVTDSSLLKARNETDRRVSEVINEFIGGSEVSTDVRLNDLLTNFCKSEIPDKPLNVDEYIAFFNKN
IIRHSTRTSSPRYIGHMTSALPYFIKPLAGLIVTLNQNVVKAETAKVLTPYERQVIAMLHRAIYQFDSDFYNHHIQNRESVLGILTSGGTIANL
TALWCARNSALGPCEGFNGVEKEGMDAALKFYGYKDAVIIGSAMMHYSFEKAADLLGIGSNNSIKIPVDKNNHIDLSELRKTVEECHRKRRLII
AIIANAGTTDCGAIDPIERVAEIAYKEGCHFHVDAAWGGPLLFSDKYRNRLKGIELADSVTIDGHKQLYLPMGLGMIFMRNPQAAKSIEKNSNY
IIRRGSIDLGRRSLEGSRPAMSIYLHAALNIIAKGGYEFLINEGIKKAEYMAGLIKSMREFELLTEPDMNMLLYRFIPERLRGKAADHMLDDAD
NEVINKYNEQLQKLQRNGGYSFVSRTSFSSELYQNKSLVALRAVLANPLTCEANINEVINDQLNIAKRLNL
>B_A0A1E8FJP1
MGEAQVSLKHLFRVFTKPEHKDSKLAQIEEHLSDNILDFLSQHVVTKKTSLEEVEQDFADAKMPETPEFVSTHAENLLEKLVAHSVNTYSPTFI
GHMTSALPYFHLSLAKLLVGLNQNLVKIETSKAFTPLERQVLGMMHNLIYDQDEDFYGKYLHSARHALGAFCSGGTVANITALWVARNKTLGPQ
GSFPGVARAGLASAYRHYDINNLGLLCSRRGHYSLSKAVDLLGIGRENLITLPCPNQTLDPALALKQGREYQEQGNKLLALVGIGGTTETGHID
PLDELADVAKELGCWFHVDAAWGGATLFSSDYRGILKGIEKADSVTIDAHKQMYVPMGAGMALFKNPEDANAVRHHAQYILREGSKDLGATTLE
GSRNGMAMMVYSSLHILGRKGYELLINQSIDKAKKFAAMIDAHPDFELVTSPTLSILTYRVCPQVIQQQLKSLPAAQSQRLNSKVDRLVVNVQK
LQREAGKSFVSRTRLESPIYGPYPITVFRVVLANPMTTKRDLKAILEEQYEIAMRNRVWQELLASVPQET
LDQA
>B_C0QAM8
MTVTHNSKKVGQMNSNDRKNPSLIADWKTLERIFIRPEDEACRKTLVKYMEQILFGLHDFLNSNVGVTQEISLLELTNLYRDTLISPDPEKKLE
NVISDIIHKIAPRAVNVASPYFVGHMTAAIPFFMVHLKTIVAALNQNVIKLETSKVLSVIEKQVLAKIHRLIYRRDEAFYQTHVQSVDTSLGAF
TTGGTTANLMAMWVARNHFLGAGVEQQGMAAAMQAKGVERAVILVSKRGHYSLKKAGGILGLGNSNVLAMDVDQHQRMDMKKLEVTIRNLKKDG
KTGIIAVVGIAGATETGTVDPLGEMADLCLGQGIHFHVDAAWGGPVLLSERYAPLLKGIERADSVSMDGHKQFYMPMTSGMVYFRDPGAMDQIA
YYSNYVNREGSVDLGIKSLEGSREANALILDSSLKIMGSRGYALMIDHGIETAKAFAAEIEQRVMFEVVTPPELNILTYRMVPPDVCRRLEHAD
PKGREILNGYLDELNIYIQRTQREAGKSFVSRTRICPVPGSEGGRVVLRSVIMNPMTTNAVLREILDEQE
RIGQGFCDPIPQIL
>B_Q74CG6
MPKNRDAARASLENLYRIFTVPEAPDSTLGAIDQAISGDVTGFLQTHIVAIERDLEDIEANFSSFSIPEEPTYVSEYTEFVKENLVAHSVHTAS
PAFVGHMTSALPYFMLPLARLMTALNQNVVKVETSKAFTPMERQVLAMLHHLVYRRDEDFYPSWIHNSRHALGAFCSGGTIANITALWVARNRL
FAPNGAFRGIAQEGLARALKHRGADGIAVLVSERGHYSLGKATDLLGIGRDDLVKVKTDANNRIDLKALREECRRFQDRNTLPLALVGIAGTTE
TGNVDPLEAMADLAQELGCHFHVDAAWGGPTLFSDRHRSLLKGIERADSVTIDGHKQLYVPMGAGMVVFKDPTALSAIEHHANYILRHGSKDLG
SHTLEGSRPGKAMLVHAGFSIIGRKGYELLIDMGIERARTFADMIKQHPDFELISEPELNILTYRYCPAAVQQTLHDVTDRERADINALLDLVC
QLLQKFQREAGKTFVSRTRLHVARHDRELTVLRVVLANPLTTDEILESVLAEQCELVQLPEIQAVLQQVEELCTGLAKAASW
>E_U3I574
SMEPEEYRRRGKEMVDYICQYLSNVRERRVIPDVQPGYMRAQLPDSAPMDPDSWDNIFGDIEKIIMPGVVHWQSPHMHAYFPALTSWPSLLGDM
LADAINCLGFTWASSPACTELEMNVMDWLAKMLGLPDKFLHHHPDSVGGGVLQSTVSESTLVALLAARKNKILEMKLSEPDTDESLLNSRLIAY
ASDQAHSSVEKAGLISLVKMKFLPVDENFSLRGETLKKAIAEDRKKGLVPIFVCATLGTTGVCAFDNLSELGPICDAEGLWLHIDAAYAGTAFV
CPEFRLFLDGIEYADSFTFNPSKWMMVHFDCTGFWVKDKYKLHQTFSVNPVYLRHPNSGAAVDFMHWQIPLSRRFRSLKLWFVIRSFGVKKLQA
HIRHGTETAKFFESLVESDPLFEIPAKRHLGLVVFRLKGPNWLTEKLLKELSSSGKLFLVPATIRDKFIIRFTVTSQFTTREDILQDWNIIQHT
AAQIISQHYGLHHISSGDEARIPNMVMEHNSDVISNASQLYVEEEKYKTPSRKIVVQPKKVAVSYNTCVISQQVKGQGDPLDDCFPEDAQDVTK
HKLTSFLFSYLSVQGKKKTARSLSCNSVPVTGLLEQCNPKAAATDKKESHANARILSRLPEEVMMLKKSAFKKLIKFYSVPNFPECSIQCGLQL
PCCPLQAIV
>E_A0A2B4SYD9
```

MLQSRKRTTQQTAGSGGKRRRMSIGVPMYSHHRNMLSKHQDGSHEREKPWAALGAWFMGPKAENGNVFHELVTQTIDSQINFRRHVYFPCDPPY
VTDELRDTHAFKASMNKLKTEMEELQNKMMNSVPFYSSRYKGHVNWDVAMPAHLGYICALLYNQNNCAAESSTVTTSFELEVGTDLCVMMGYDP
DKSMGHLVTGGTVANIEALWAARNVKFFPLGLRRALCKEEKLANAKHYEVFFPQRGEKGELISGTEWELLNLDTSSILIMPEEVQKLAGLENPS
ELMRLLSDYLYESIGAAEFARHHPLIENTCVIVPSTAHISFTKAVTILGLGKSSLVPVAVDENSRMDAVVLKDILEKHLNRQAPVMAVVAVMGT
TEESSIDPLTDILKIRKDFSKKGLDFSIHADGAWGGYFCSMLRDQPKNFYLKAPEDSGFVPQLYLSSYVHDQLSALDQCDTITIDPHKSGFCPY
PAGALCYKDRKMNTFLQITNSVVYYHGDVALGNIGIEGSKPGAAAASVLLANRVIGLHRNGYGRILSECTFTAKMLYCLWMTLPEEDDQFIIET
TKPLPSSWKGMSKKKQKEFIRERITGKNNEELAKDDEAMEYLKEVGPDTLVSCFTVNLKGNKSVDVCNSLNMAIFQGLSHSSGERTAHRIPMIV
TSSSMLHHKHSSALKNFKKRLGVKDPKSHHDFVSTGVVNDKHQVIVYYAGNFNNISKQYGTVATLQFNSDSQAKEYKSKQDSLMATSTEPSPIV
FRSKKSTLHDVFFGESEYGDKAKAFDLYIGLPSRGSKPFMSARLKVVDVPQYEHFDDDEYPEFLSYFMYGDKKDAFLFHIPTKNLDFLQIVKLD
GIPKGVGSEGIKDLLLKKGIEVNLPEISGSWPEGHDDVKDPLKNHKYEIKFVGIDGEEVASKVRIKRKVWFDGTKLKNCANTNKTGKKRSNVEK
ERGHSPNIEEFIIDIRDRSCPCDPVSPTKALISIPIEVSPFLTLTSAVGIQLESQFQFQRPIQMSAQVRNFLPNRLITKKGVLHMKKPGSSVSR
CVGADPCGLLPPYSSPELPLAPPPVPYAGFPPPPVPYGPQGFSQPASMAPNGYGMHSINAMAQDAAAAMAAKLKLLGGTHPEKKFVPHLPPMGR
PQTGKSSYNPPPTLPEPQAAAGFPPALPEQLPYQQSFGYNQQGMPPQTAGFDSQPAPGFPQPVVPAIQGSFANLYLYRLSASSFSPGPGPRICP
SVAIAHRTSRGTQLVNNDVPQVPFPAEMNQAPAFNDPGQQMAPNAANMLQSRKRTAHRAAGPGGKRQRMSKGAPMYSHHRNMFSKHQDGSHKRE
KPWAALGAWFMGPKAENGNVFHELVTQTIDSQINFRRHVYFPCDPPYVTDELRDTHAFKASMNKLKTEMEELQNKMMNSVPFYSSRYKGHVNWD
VAMPAHLGYICALLYNQNNCAAEASTVTTSFELEVGTDLCVMMGYDPKKSMGHLVTGGTVANIEALWAARKVKFFPLGLRRALCKEEKLANAKH
YEVFFPQRGTKGELISGTEWELLNLDTSSILTMPEEVQKLAGLENSSDFMELLSDHLYESVGAAEFARRHPLIEKTCVIVSSTAHISFTKAVTI
LGLGKNSLVPVAVDENSRMDAGVLKDILEKYLDTQVPVMAVVAVMGTTEESSIDPLTDILKIRKDFSKKGLDFSIHADGAWGGYFCSMLRDQPK
NFYLKAPEDYGFVPQLYLSSYVHDQLSALDQCDTITIDPHKSGFCPYPAGALCYKEKKMNTFLQITNSVVYYHGDMTLGDIGIEGSKPGAAAAS
VLLANRVIGLHRNGYGRILSECTFTAKMLYCLWITLPEEDDQFMIETTKPLPSSWKGMSEKKQKEFIRERIIAKTPEDSGFVPQLYLSSYVHDQ
LSALDQCDTITIDPHKSGFCPYPAGALCYKDRKMNTFLQITNSVVYYHGDVALGNIGIEGSKPGAAAASVLLANRVIGLHRNGYGRILSECTFT
AKMLYCLWMTLPEEDDQFIIETTKPLPSSWKGMSKKKQKEFIRERITGKNNEELAKDDEAMEYLKEVGPDTLVSCFTVNLKGNKSVDVCNSLNM
AIFQGLSHSSGERTAHRIPMIVTSSSMLHHKHSSALKNFKKRLGLDPKGDSSVKFLITTCMDPWATSVEFLDDLASIMRNSILCAIGRVKDPKS
HHDFVSTGVVNDEHQVIVYYAGNFNNISKQYGTVATLQFNLDSQAKEYTSKQDSVMATSTEPNPIVFRSKKSTLYDVFFGESEYGDKAEVFDLF
IGLPSRGSKPFMTANLKVVDVPQYEHFDDDEYPEFLSYFMYGDKKDAFLFHIPTKNPDFLQIVKLDGIPKGVGSEGSKDLLLKKGIEVNLPEIS
GSWPEDHDDVKDPLKNQKYEIKFVGIDGEEVASKVKIESNVWFDGTNLNK
>E_G1X4X9
MDSSQFRQAAHAAIDQIVDYYDNIRDRRVLSDVEPGYLRQLLPQGIPETGEKWEDIQKDIEAKIMPGMTHWQSPNFLAFFPSNSSFPGILGDMY
SAAFSCAAFNWQCSPAVTELETIVLDNVAKLINLPEEYHSTSEGGGVIHGTASEAIVTVIVAARDRYIARSKERWAEEGLSEDEIEDKVCTLRG
RMVALGSDQAHSSTKKGAIIAGVRFQTIETKIGDYALNGELVKQKIEDLESKGLVPFYITVTLGTTPTCATDDFASISATLSTYHTTHPTTPKI
WAHIDAAYAGAALVLPEYSHIPSIFPTFADSFDFNMHKWLLTNFDCSCLYVKRRRDLIDALSITPAYLRNEYSDRGLVTDYRDWQIPLGRRFRS
LKAWFVTRTFGVEGLRAHVRNGIAGGEAFTQLLEADKERYELVSKPAFALNVFRVNPPPKLAKEVENDKKEFERRCNEVTRKVGDRVNKEGKIF
ITQTVLGKGEEAITAIRVVGGAPAVQVQDLRNAFAIITEVVDRVWEEEVREHEAGAREQVELQAV
>E_I1S8G3
MALSSNSTFGNLLVPAAHDDDENQIRTLFTQVVDLGIDFRASDTIFSEETEASPSRISFDKIPESGLSYEELIQQFASVASKSSNWGSPNFLGF
SDAANNVAGLAAALLIPLLNQNIANQEICSPEATFIEMEVVHWLRETLGYPVPETYTKASGIGGVLTLGGCLSNTIALLAAREKCFPGSGIQGI
PVLPTKIRVLVPSVTEHYSIRSAMAWLSLGEKNVIRVPVDSKFHMDREALKQIIDRERECGNIIMASKILEDKNVWFHVDACHGSQLAFSEKHR
YKLRGIGKADSITIDSHKTMLVPYNCSLILFRDPSTHVALSTNSDLILNTQWSLGRVTPFIGSKAFDALKIWSTIKLFGRRRLGQIIDERLELT
SAIQDEVTQRPNLILLNKTDINSCMMVYIPFQIQEYCIEKIIPLSDADLEKINLMNRRIKDTVRKEGMYYIHGFPLQSCPHERFITPDKQVFVL
RTMNGNPLSTIENAKGLLDRIERLGQLFFSEAGYRCIQAGPSLNHLRRAEDKLSHKIRNLFGNQDFVAVVYGSSALPRNAILSDIDLMVFIRSV
EPGQREDMVTAFRSIMNQESILVDDEVPLSRKLSIPFELAARAAESGPVLDDAGEIRSICKTPEYLSSNKMLQRLIFGVLTTPNRIIAANANGA
SSFETLKSNAGKTLVGLIRQLHHASFSTTDEFVKLAISDGVRSGEQYLGYKDRSEVVEKLSQIFDKAYGSEYGRPVDPDSS
>E_A0A0D2IJV6
MDQHPSLTEIIATLSKILTTPTLLTAQPKSISESKSRPPILPRDIPNFNPLKLELTSLALNFEDRDSDTLAALSRHLTTDILPHLNLASLSPNY
YGFVTGGATPAALLGDFLTSIYDQNVQVHLPRETIATTLEVATLNLLVQLFRLPEAEWLIGGSGPGGGTFTTGATASNVLGLALGREYALRKAL
ERKSKMNGTALSCGEWGIAEMMMKAGVRKIQILSTLPHSGTAKAASLVGIGRRNIISISADHDPLQIDLGRLEQEARKEHVLSILAISTGEVNT
GRFATNSSALMSRLRSICDELGIWIHVDGAFGLFGRIFSPDDLEYREIVQGVQGLELADSITGDCHKLLNVPYDSGVFFTRHKGLSEDVFRNGN
AAYLTGAMGDGDVIQSPLHIGIENSRRFRALPVYSTLRAYGRDGYLDMLRRQVRLARRVSKWLLKDDRFEVLPGGADTDEVLAKTFIVVLFRVR
DEEVSKDFVKNVNATGRIYISGTVWDGKPAARIAVSNWQVDVERDGGLIEDVLNQITGGRS
>E_A0A091VHQ0
GKEMVDYICQYLSNVRERRVTPDVQPGYMRAQLPDSAPMDPDSWDNIFGDIEKIIMPGVVHWQSPHMHAYFPALTSWPSLLGDMLADAINCLGF
TWASSPACTELEMNVMDWLAKMLGLPDKFLHHHPDSVGGGVLQSTVSESTLVALLAARKNKILEMKVSEPDTDESSLNSRLVAYASDQAHSSVE
KAGLISLVKMKFLPVDENFSLRGETLKKAIAEDRRKGLVPVFVCATLGTTGVCAFDNLSELGPVCDAEGLWLHIDAAYAGTAFVCPEFRLFLDG
IEYADSFTFNPSKWMMVHFDCTGFWVKDKYKLHQTFSVNPVYLRHPNSGAAVDFMHWQIPLSRRFRSLKLWFVIRSFGVKKLQAHVRHGTETAK
FFESLVKSDPLFEIPAKRHLGLVVFRLKGPNWLTEKLLKELSSSGRLFLIPATIHDKFIIRFTVTSQFTTREDILQDWNIIQHTAAQIVSQNYG
LHCINSGDGARIPNMIVKPSSDAISSASQLYLDGGKYKTPSRKIVVQPKKLEASASMCVISQQVKGQGDTPDDCFPEDVQNVTKHKLTSFLFSY
LSVQGKKKTARSLSCNSVPMTGGLEQCNPKAAATDKESHANARILSRLPEEVMMLKKSAFKKLIKFYSVPSFPECSIQCGLQLPCCPLQAIV
>E_B4Q567
MLASENFPTHHFKESIFKPYSTTSGDDLASVTPLTATAALVASTPSPADSTSAVAFEQASKMLATAANNNNNNNNNNITSTKDDLSSFVASHPA
AEFEGFIRACVDEIIKLAVFQGTNRSSKVVEWHEPAELRQLFDFQLREQGESQDKLRELLRETIRFSVKTGHPYFINQLYSGVDPYALVGQWLT
DALNPSVYTYEVAPLFTLMEEQVLAEMRRIVGFPNGGQGDGIFCPGGSIANGYAISCARYRHSPESKKNGLFNAKPLIIFTSEDAHYSVEKLAM
FMGFGSEHVRKIATNEVGKMRLSDLEEQVKQCLENGWQPLMVSATAGTTVLGAFDDLAGISELCKKYNMWMHVDAAWGGGALMSKKYRHLLNGI
ERADSVTWNPHKLLAASQQCSTFLTRHQQVLAQCHSTNATYLFQKDKFYDTSFDTGDKHIQCGRRADVFKFWFMWKAKGTQGLEAHVEKVFRMA
EFFTAKVRERPGFELVLESPKCTNISFWYVPPGLREMERNREFYDRLHKVAPKVKEGMIKKGSMMITYQPLRQLPNFFRLVLQNSCLEESDMVY
FLDEIESLAQNL
>E_A0A087YBQ9
EAMATSEPRATEEEQDPNAANLRPQSGTYEYAWMHGCTRKLGMKICGFLQKNNSLEERGRLAGQKNLLSCDNSDRDSRYRRTETDFSNLFARDL
LPAKNGEEPTIQFLLEVVEILTNYIKKTFDRSTKVLDFHHPHQLLEGMEGFNLELSDQPESLEQILVDCRDTLKYGIRTGHPRFFNQLSSGLDI
IGLAGEWLTSTANTNMFTYEIAPVFVLMEQLTLKKMREMIGWPGGEGDGLFSPVTGGAISNMYSVMIARYKYFPEVKTKGMSAAPRLVLFTSEH
SHYSIKKAGAALGFGTDNVILLSTDERGRVIPADLEAKIIDAKQKGYVPLFVNATAGSTVYGAFDPINEIADICEKYNLWLHVDGAWGGGLLMS
RKHRHKLNGVERANSVTWNPHKMMGVPLQCSAILVREKGILAGCNSMCAGYLFQPDKQYDVTYDTGDKAIQCGRHVDIFKFWLMWKAKGTIGFE
QHIDKCLDLSQYLYNKIKNREGFQMVFDGVPQHTNVCFWYIPPSLRGMPDSDERREKLHRVAPKIKAMMMESGTTMVGYQPQGNKVNFFRMVVS
NPAATQSDIDFLIDEIERMGQDL
>E_A0A090MC16
XXLRFLDEIATHMVEYYRNSARAADPVSTYNSPLALHEKFKEVGIPLAIGTGEEPVSMSLLTTAMNTVIQNSARTSHPMFMNQLYAGVDPIALA
GEWASSAMNSNVHTYEVAPVFTEIERSMLAKVATLWLGENADGTTPNHDGLFVPGGSIANLYSIILARERVCPEAKKTGMPPGYVAFCSEQSHY
SYKKCAHMVGLGMDNMIKVDCGPNGAMLPEALERAVAEAIAAGKKPFYVGATAGTTVLGAYDPYDALADLCERNNMWLHVDGAWGGAAILSKRH

KHLMKGAERADSFCWNPHKLLGIPLQCSIVLSKHAGSFMAANSYKADYLFQPDKLDSEADLGDRTIQCGRKSDALKLWLAWKYRGDAGWERLVD
HSFALAKFVETEVVSDQTGAWALAAPAQCANVGFWYVPKRLRPFNKETATPKQMKELAKVAPKLKDRMQRAGLAMIGFQPVPAFGLQNFFRLVL
PNPRHNSESKLRELMHQMDKLGEDL
>E_A0A2B7Z7I8
MSIQSLQSPEDKEFPLQIWQTALSPWTSPLLPAPATLSKVRSSLITPLPTTGLGFSETKRHIINDIAPGFNGSSLSANYYGFVTGGVTPAALLA
DNIVSAYDQNVHVHIPDHSIATDVEDRALTFLLDLFDLDHKYWAHKTLTTGATGSNVLGLALGREFVLRAAVERKAGADLKVMKSVGEHGMAEV
LIAAGLKGLQVVSTYPHSSLGKAAGVLGIGRANVKSVCADGGGESSPPLRFDFQILEKELARSDMASIVAVSCGEVNTGHFATGGLKEFRKIRQ
LCDKYGAWLHIDGAFGLFGRILKSGGEFDRIRKGCQGLELADSITGDAHKLLNVPYDCGFFFSRHANLAEEVCRNPNAAYLSAGTGGGGIPAPC
NNGLENSRRLRALPVYATLVAYGKDGYRDMLERQIRLARSVVGWLFEHPAYAVLPHNPEKESLLQDTFMIVLFRAKDEDLNRVLVNKINATSKM
YVSGTSWDGKPACRIAISNWRVNEEQDLEMITSVLGEIAQ
>E_F6NX32
MAASAPSSSSSGGVPDPNSTNLQPPSSIAHGCTRKLGMKICGFLQKNNNVDDKGRIVGLFNDQQPRSILTRDNERDSRFRRTETDFSNLYARVN
LILTDLLPAKNGEEYTMQFLLEVVEILTNYVRKTFDRSTKVLDFHHPHQLLEGMEGFNLELCDQPESLEQILVDCRDTLKYGVRTGHPRFFNQL
SSGLDIIGLAGEWLTSTANTNMFTYEIAPVFVLMEQLTLKKMREIVGWPNGEGDGIFSPGGAISNMYSVMVARYKHYPEIKIKGMAAAPRLVLF
TSEHSHYSIKKASAVLGFGTENLILLRTDERGRVIPADLEAKVIDAKQKGFVPMFVNATAGSTVYGAFDPINEIADICEKYNMWLHVDGAWGGG
LLMSRKHKHKLSGIERANSVTWNPHKMMGVPLQCSAILVREKGLLQGCNSMCAGYLFQPDKQYDVTYDTGDKAIQCGRHVDIFKFWLMWKSKGT
TGFEKHIDRCLELSEYLYHKIKNREGYEMVFQGEPQHTNVCFWYIPPSLRLLPDGEEKRHRLHKVAPKIKALMMECGTTMVGYQPQGEKVNFFR
MVVSNPAVTRSDIDFLIDEIERLGQDL
>E_E7FDZ2
TLEVMAASAPSSSSSGGVPDPNSTNLQPPSSNYDWSGVAHGCTRKLGMKICGFLQKNNNVDDKGRIVGLFNDQQPRSILTRDNERDSRFRRTET
DFSNLYARDLLPAKNGEEYTMQFLLEVVEILTNYVRKTFDRSTKVLDFHHPHQLLEGMEGFNLELCDQPESLEQILVDCRDTLKYGVRTGHPRF
FNQLSSGLDIIGLAGEWLTSTANTNMFTYEIAPVFVLMEQLTLKKMREIVGWPNGEGDGIFSPGGAISNMYSVMVARYKHYPEIKIKGMAAAPR
LVLFTSEHSHYSIKKASAVLGFGTENLILLRTDERGRVIPADLEAKVIDAKQKGFVPMFVNATAGSTVYGAFDPINEIADICEKYNMWLHVDGA
WGGGLLMSRKHKHKLSGIERANSVTWNPHKMMGVPLQCSAILVREKGLLQGCNSMCAGYLFQPDKQYDVTYDTGDKAIQCGRHVDIFKFWLMWK
SKGTTGFEKHIDRCLELSEYLYHKIKNREGYEMVFQGEPQHTNVCFWYIPPSLRLLPDGEEKRHRLHKVAPKIKALMMECGTTMVGYQPQGEKV
NFFRMVVSNPAVTRSDIDFLIDEIERLGQDL
>E_A0A0V1H7S9
MDAEEFRKWGKKMIDFVADYWIHLPSRTPMSDVKPGYLRSLLPEEAPTDPDSWENIFSDIETVILQGTTHWHHPLFFAYFPTGNSYPSILGDIL
SAGIGCIGFTWNSSPSCTELEMVMMDWLAKLLNLPEYFLYSHSGPGAGMIQGTASECVLFSMLAAKNKTCKKYETENKQHHICEKNLIAYCSDQ
AHSSVERAAMLAHVQIRKVPSDENYRMTRVALQAVIENDINAGFIPFFVCATLGTTNSCAFDCLTEIGLLCKEKEIWLHVDAAYAGSAFICPEY
RYLLDGIEYADTFNFNPHKALMINFDCSAMWFKNVLEIENAYYVNPQYLKHEHQNMMPDFRNWQIPLGRRFRSLKLWLTFRALGVRFLQENIRK
MCRLAKEFADLVVQDERFELVAPVILGLVCFRLKDTNEVNEKLYQLINNQRRIHVVSSVLRNVFVLRISISSALTESADISFAWKVISASATKL
LTS
>E_A0A2H5NKN4
MESGGLKPMDAEQLRENAHKMVDFIADYYKSIENFPVLSQVQPGYLHNLIPDSAPHHPESLQNVLDDIQEKILPGVTHWQSPNYFAYYPSNSSV
AGFLGEMLSAGLNIVGFSWITSPAATELEMIVLDWLAKLLKLPEDFLSSGQGGGVIQGTASEAVLVVLLAARDNALKRVGKNSLEKLVVYASDQ
THSALQKACQIGGIHPQNFRVLKTDSSTNYSLSPDSLAEAISRDLTIGLIPFFLCATVGTTSSTAVDPLLALGNIAKSNGMWFHVDAAYAGSAC
ICPEYRQYIDGVEEADSFNMNAHKWFLTNFDCSALWVKDRNTLIQSLSTNPEFLKNKASQANMVVDYKDWQIPLGRRFRSLKLWMVLRLYGLEN
LQGYIRNHIQLSKHFEGLVAQDLRFEVVTPRIFSLVCFRLLPPHNDEDHGNKLNHKLLDDINSTGKIFISHTVLSGKYILRFAVGAPLTEWRHV
NAAWEVMQDKASALLARLSIE
>E_A0A099Z5N3
GIEMVDYICQYLSSVRERRVTPDVQPGYMRAQLPDSAPMDPDSWDNIFGDIEKIIMPGVVHWQSPHMHAYFPALTSWPSLLGDMLADAINCLGF
TWASSPACTELEMNVMDWLAKMLGLPDKFLHHHPNSVGGGVLQSTVSESTLIALLAARKNKILEMKVSEPDTDESSLNSRLIAYASDQAHSSVE
KAGLISLVKMKFLPVDENFSLRGETLMKAIEEDRNRGLVPVFVCATLGTTGVCAFDNLSELGPICDAEGLWLHIDAAYAGTAFVCPEFRLFLDG
IEYADSFTFNPSKWMMVHFDCTGFWVKDKYKLHQTFNVSPVYLRHPNSGAAVDFMHWQIPLSRRFRSLKLWFVIRSFGVKKLQDHVRHGTETAK
FFESLVKSDPLFEIPAKRHLGLVVFRLKGPNWLTEELLKDLSSSGRLFLIPATIHDKFIIRFTVTSQFTTREDILRDWNIIQQTAARIVSCRYT
LHRVRSCDGARAPGMIVKPSSDATASAPQCYPGVGEHKTPSRKIVVQPKKLAVSPSSCVINQQVEDQGDPLDDCFSEDAQEVTKQKLSSFLFSY
LSVQGRKKTARSLSCNSVPMTGAPEQCNPQAGATDKKESHTNAKILSRLPEEVMMLKKSAFKKLIKFYSVPSFPECSVQCGLHLPCCPLQAIV
>E_T1KV43
MSHTQETEKISWLADLYQRTNKNWFGLNHSSEASSTSNNVSESKKDNVADDTLIRKVAEMILDEFHLIGDSDNVKNTNTKLVNFTQPADLEKIL
KLEITKDGLTIKQLEDFCRQVIKYSVKTTHPHFYNQLYGGVDQFGVAGAWLTDALNTSQATYEIAPVFTLLERKIIEYCGSKCGWDLDKIDGIF
SPGGSISNMYGMILARYNKFSDTKENGVRGLPRLVAFGSDSSHYSLSKSSIWMGHGTNSVVKVKTDDRGKMIPSELEKEIENTIKDGAVPFVVI
ATAGTTVLGAFDPIREVAAICAKYNIWLHVDAALGGTFLLSLKHRTLLDGIELADSVAWNLHKLAGAPLQCSLFLTRHCQLLQHCNSLNAEYIF
QSDKYYDADYDLGDKSIQCGRKVDSLKAWLTLASRGEDEWEHLVDNIIQMNRYITDKLAHRPNFKLVLPQFEGSTVSFWYIPEKLRNQTHIDPA
TLHKVCPAIKSRMMSSGSLMIGYQPLTCKNLPNFFRLSLTCIPPATMEDMDFIVDEIERLGKDLEF
>E_A0A1S8W5A4
MDGPKCKQEPATNTDTTIVSTTTVNTTAVSTTAATTAATTTTSTDGTGRIVAATPMSRLAMVDADIDEGFHDDDYPEEFPFWYGGASGEQHMIH
YGSTDDAVDETSAASSVASKTTPLSASAHKAVFPLDSFLQSSSTIDLRIRSNDAVGHPTSFHDHDSVLHYFVPTTQSEKLLDKYISGVIEAFLH
EPSPIYTTSTHIPHDPQSAFGGLSIPDGGRPSGSDLEAYLNHLKTNVIDRSTRTASSRMIGHMTTALPFFHRPLARLLAALNQNVVKIETASTF
TNLERQTLGMLHKAFYDLPDTFYQRHAYAPDYSLGVMASGGTIANITALWIARNKALAPNSSNGCRGIDKEGVVSAMGKYGYKRAVIIGSALMH
YSFKKAADLLGLGEEGLCLIPTDAHFCMRVDLLKSKVDELISEGALIIAIVGIAGTTETGSIDPLFDIYSIAHRHNIHFHVDAAWGGPLIFSPE
HRCKLNGISQADSITVDGHKQLYTPMGLGILLLRSPSLALYIRKTASYVIRNDSPDLGKYTLEGSRPANALYLHASLSLLGKHGLGILVTRSVT
IVRQTAVRLDSHPSRCFQILHQPMSNLLLYRYVPSALRDAIADGSYISTKDDEAWISEATRRIQIRQASCVSVGSVHPHSLTSPNPTGGIDPPA
PFGQQQQPLPGFVSRTRVWFRGHHVDALRVVVANPLTTWDDIEGVISDQLHIGAEIEAEMQREQMVKRLQSWISTPCSSVSDTMLSEPEAAKAT
LSFVSATDLHNGSHGDPGWWPGWPFDL
>E_A0A0V1MFM8
MDAEEFRKWGKKMIDFVADYWINLPSRTPMSDVKPGYLRSLLPEEAPTDPDSWENIFSDIETVILQGTTHWHHPLFFAYFPTGNSYPSILGDIL
SAGIGCIGFTWNSSPACTELETVMMDWLAKLLNLPEYFLYSHSGPGAGMIQGTASECVLFSMLAAKNKTCKKYETENKQHHICEKNLIAYCSDQ
AHSSVERAAMLAHVQIRKAPSDKNYRMTRVALQAVIENDINAGFIPFFVCATLGTTNSCAFDCLTEIGLLCKEKEIWLHVDAAYAGSAFICPEY
RYLLDGIEYADTFNFNPHKALMINFDCSAMWFKNVLEIENAYYVNPQYLKHEHQNMMPDFRNWQIPLGRRFRSLKLWLTFRALGVGFLQENIRK
MCRLAKEFADLVVQDERFELVAPVILGLVCFRLKDTNEVNEKLYQLINNQRRIHVVSSVLRDVFVLRISISSALTESADISFAWKVISASATKL
LTS
>E_A0A200PVP3
MAPNPTNQSNSILTLNKLELRENGFKSMDEAEKLRENVLKPMDSEQLRENAHKMVDFIADYYKNIETFPVLSQVEPGYLEKLLPDSAPNHPESL
QNVLDDVQTKILPGVTHWQSPDYYAYFPSNSSTAGFLGEMLSAGLNIVGFSWVTSPAATELEVIVLDWLAKMLKLPEHLLSSGQGGGVIQGTAS
EAILVALLAARDKVLRKVGKNSLPKLVAYASDQAHASMLKACQIAGIHPENIRLVTTDSSTNYALSPDVLGEAISKDIAAGLIPFFLSSTVGTT
SSTAVDPLYALAKIAKGNEMWFHIDAAYAGSACICPEYRHYFDGVEEADSFNMNAHKWFLTNFDCSPLWVKDRSALIQSLSTKPEYLRNKASEA

NMVVDFKDWQIPLGRRFRSLKLWMVLRLYGLENLQCYIRNHIKLARHFEELVASDSRFEIVVPCKFSLVCFRLLPAHNDQDCGNKLNQDLLDAV
NSTGKIYISHTVLSGKYILRLVVGAPLTEERHINAAWKVFQDEATVLLGGV
>E_G1TK30
MTAALLTDFILMADSKPLPCADGDPVAMEALLREVFSIVVDEAVLKGTSASEKVCEWKEPEELKQLLDLELRGQGERREQLLERCRAVIRYSVK
TGHPRFFNQLFSGLDPHALAERIITESLNTSQYTYEIAPVFVLMEEEVLSKLRALVGWTSGDGVFCPGGSISNMYALNLARYHRYPECKQRGLR
ALPPLALFASEECHYSIKKGAAFLGLGTDSVRVVKADERGKMVPEDLERQIRLAEAEGAVPFLVSATSGTTVLGAFDPLEAIADVCQRHGLWLH
VDAAWGGSVLLSQTHKHLLDGISRADSVTWNPHKLLGAGLQCSALLLRDTSNLLKRCHGSQASYLFQQDKFYDVALDTGDKVVQCGRHVDCLKL
WLMWKAQGGQGLELRVDRAFALARYLLEEVKKREGFELVMEPEFLNVCFWFVPPSLRGQRSPNYSQELAQVAPALKERMVKEGSMMIGYQPHGA
RVNFFRVVVANPALTRADVDFLLDELERLGRDL
>E_D8SSC2
MDPQEFRAQAHKMVDFIADYYRDVESLPVRSQVTPGYLRSSLPSNAPEEPQSFDTVLDDVKSMIVPGVTHWQNPNFFGFFPSNSSTAGMLGEFL
SGGFNVDGSEWATSPAATELEMLVLNWLGKLLNLPDEFLFNRSGNGGGVIHASASEAVLVALLAARGRAISENKAKGLEEQEILSKLLVYTSDQ
THPCLHKACVIVGLPKSNLVILPTLATDDYALSLPILKSAVRNGVTKGFIPFFLGATVGTTSSSAIDPLPALADIAKEYGMWFHVDAAYAGNAC
ICPEFRHFLNGVENAHSFNLSANKWLLTNIDCSILWLKRYEFLNLLFFIYTISFQLKTSSIQSRVVNFKDWQVAQGRRFRQLWFVMRLYGALGL
RNHIRTHINHAKHFEILVREDSRFEILAPCRFGLVCFRLKPSVKHEDNGWKLNSSLLEAINSGGKIFMTHTVLSGVYTLRMSIGGTQTKRENVD
DAWKIIQEEAQNLLDQEII
>E_D8R3Z6
MGEANIGPKPIDAEEFRKHAHEMVDFIADYYRDIESFPVRSQVSQPGYLKTLLPPAAPEDPEALEEVFADIQSKIIPGVTHWQSPNFFGYYPSN
SSTAGLLGEMLSAGLNIVGFSWITSPAATELEIIVLDWLAKLLKLPDEFLFGGNGGGVIQGTASEAVSVVLLAARTRAISENKRKGLSEAEILS
KLAVYTSDQTHSCLQKGCAIAGIPLENLVIVPTDSSTNYAVSPAAMRQALEDGVKQGLLPFFLCGTVGTTSSSAVDPLSALGDIAKDFGMWFHV
DAAYAGSACICPEFRHHLDGVEKADSFNMNAHKWLLTNFDCSALWVKESSHLVSALSTTPEFLRNKASDLNQVVDYKDWQIPLGRRFRSLKLWF
VMRMNGASGLRSYIRNHVRLAKRFEGFVREDPRFQLLVPRTFGLICFRLKPESDDPDNGRTLNSTLLEAVNSSGRMFITHTVLSGVYTLRMAIG
GPLTQDKHVDAAWKLIQEEATTLLVKGPSHILANNLRLSPILANNLRLSPILANNRI
>E_A0A1U8A7E6
MEGGMKPMDAEQLRENAHKMVDFIADYYKSIESFPVLSQVEPGYLRKIIPDSAPNQPESLQNVLDDIQAKIIPGVTHWQSPNFFGYFPSNSSIA
GFLGEMLSGGLNIVGFSWVTSPAATELETIVLDWLAKMLKLPEDFLSSGQGGGVIQGTASEAVLVVLLAARDKALRMFGRNSIGKLVVYASDQT
HSALRKACQIGGIHAENCRLLKTDSSTNYALSPEVLTEAISQDTASGLVPFFLCATVGTTSSTAVDPLLALGKIAKDNKMWFHIDAAYAGSACI
CPEYRHYLNGVEEADSFNMNAHKWLLTNFDCSVLWVKDRSALIQSLSTNPEYLKNKASEGNMVVDYKDWQIPLGRRFRSLKLWMMLRLYGQENL
LNYLRNHIELAKYFEELVSVDQRFEIVVPRTFSLVCFRLLPPQGDEDCINKMNRDLLEVVNSTGKIFLSHTILSGKYILRLAVGAPLTEGRHIS
AAWKVLQDEATTLLEQYMK
>E_A0A1U8A5K9
MEGGMKPMDAEQLRENAHKMVDFIADYYKSIESFPVLSQVEPGYLRKIIPDSAPNQPESLQNVLDDIQAKIIPGVTHWQSPNFFGYFPSNSSIA
GFLGEMLSGGLNIVGFSWVTSPAATELETIVLDWLAKMLKLPEDFLSSGQGGGVIQGTASEAVLVVLLAARDKALRMFGRNSIGKLVVYASDQT
HSALRKACQIGGIHAENCRLLKTDSSTNYALSPEVLTEAISQDTASGLVPFFLCATVGTTSSTAVDPLLALGKIAKDNKMWFHIDAAYAGSACI
CPEYRHYLNGVEEADSFNMNAHKWLLTNFDCSVLWVKDRSALIQSLSTNPEYLKNKASEGNMVVDYKDWQIPLGRRFRSLKLWMMLRLYGQENL
LNYLRNHIELAKYFEELVSVDQRFEIVVPRTFSLVCFRLLPPQGDEDCINKMNRDLLEVVNSTGKIFLSHTILSGKYILRLAVGAPLTEGRHIS
AAWKVLQDEATTLLEQYMKT
>E_A0A0D1XU64
MDQTMSLTDVVSVLQEIMQTPGLPSSEDAVQQGQETSSVSTKKSTRPILPINFPPLDTLALVDSTQPTEHDNSLQSLKSHLIEHILPYLNSQSL
SPNYYGFVVGGVTPAALLGDFIASIYDQNVFMHLPNDTLCTAIEVQALNALALDFFRLSPREAWEVGGPGSGGATFTTGATGSNILGLALGREF
VLRKALETKGVALADASVGEYGLFEVMDLAGVKKIQVLSTLPHSTIAKAASVVGIGRRNVVSIATHDDPLQIDLEKLKAEVEKKDTLNILAISA
GEVNTGRFATYSGANMKEIRHICSQNHVWIHVDGAFGLFGRLFTNDDPDYSHVARSLEGLDLADSITGDCHKLLNVPYDCGIFFTRHKSLSEHV
FRNGNAAYLTSGLGPDDGIQSPGNIGLENSRRFRALPVYATVKAYGRQSHVAMLKRQINLARRITEWLLRDDRFQVLPGGLDKADIINKTYIIV
LFRATDPEVNKSLVKMVNADGRIYVSGTVWDGEPAARIAVSNWRVDVDRDATLVQQVLNSVARSR
>E_A0A0Q3PWB0
MDPEEYRRRGKEMVDYICQYLSNVRERRVTPDVQPGYMRAQLPDSAPVDPDSWDNIFGDIEKIIMPGVVHWQSPHMHAYFPALTSWPSLLGDML
ADAINCLGFTWASSPACTELEMNVMDWLAKMLGLPDKFLHHHPDSVGGGVLQSTVSESTLVALLAARXNRILEMKVSEPDTDESSLNTRLIAYA
SDQAHSSVEKAGLISLVKMKFLPVDENFSLRGETLKKAIAEDRKKGLVPVFVCATLGTTGVCAFDSLSELGPICDAEGLWLHIDAAYAGTAFVC
PEFRLFLNGIEYADSFTFNPSKWMMVHFDCTGFWVKDKYKLHQTFSVNPVYLRHPNSGAAVDFMHWQIPLSRRFRSLKLWFVIRSFGVKKLQDH
VRHGTETAKFFESLVKSDPLFEIPAKRHLGLVVFRIKGPNWLTEKLLKELSSSGRVFLVPATIXDKFIIRFTVTSQFTTREDILQDWNIIQHTA
ARIVSQNCGLHCISSGDGTGIPNIIAEPSSXVISNASQLSPDGGKYKTPSRQIVVQPKKLAVSPSTCVISQEVKGQGDPLXDCFPEDAQDVAKH
KLTSFLFSYLSVQGKKKTARSLSCNSVPVTGGLEPCNPKAAAADKESHANTRILSRLPEEVMIFKKSAFKKLIKFYSVPSFPECSIQCGLQLPC
CPLQAIV
>E_W6Z9E5
MEVQSQEIFNQLAVEIAEIHVHPPPENVLPSGDTLSSARSKLQTHLPTKGVGLEESIRHLRQDLVPAFNASSRSPNYYGFVTGGVNQAAALADN
LVTAFDQNVQVHLPNETIATDVEDRALSLLCELLNFEPSQWPHRIFTTGATAANVLGLACGREYVIAEASAHRTDAENSVGEIGIVEAMRRAGI
DDIQILTTVPHSSLSKAASILGLGRASVKCLGCSDAPHKFDMQLLKKSLELPGAASIVVVSASEVNTGVFATSGPEEMQELRKLCDMHGAWIHA
DGAFGLFGRILSSPAHSSIIEACADLELADSITGDGHKLLNVPYDCGFFLSRHRAMAERVFQNPNAAYLASGNGPDTIMSPLNIGLENSRRFRA
LPVYASLVAYGRDGYRDMLERQIQLARGIAQHILESSQYELLPQEAAPHEDILSGIFIIVLFRAKDEELNKQLVDKIKATRKIYVSGTSWEGRP
ACRFAISNWMTDVERDLPVIKQVLRDIA
>E_I3KQZ2
MATSEPRAAGGDQDPNSANLRPPTTTNEYAWMHGCTRKLGMKICGFLQKNNSLEEKGRLAGQKNLLSCDNSDRDARFRRTETDFSNLFARDLLP
AKNGEEPTMQFLLEVVDILTNYVKKTFDRSTKVLDFHHPHQLLEGIEGFNLELSDQPESLEQILVDCRDTLKYGVRTGHPRFFNQLSSGLDIIG
LAGEWLTSTANTNMFTYEIAPVFVLMEQLTLKKMREMIGWPNGEGDGLFSPGGAISNMYSVMIARYKYFPEVKTKGMSAAPRLVLFTSEHSHYS
IKKAGAALGFGTDNVILLSTDERGRVIPADLEAKILDAKQKGYVPLFVNATAGSTVYGAFDPISEIADICEKYNLWLHVDGAWGGGLLMSRKHR
HKLNGIERANSVTWNPHKMMGVPLQCSAILVREKGILAGCNSMCAGYLFQPDKQYDVTYDTGDKAIQCGRHVDIFKFWLMWKAKGTAGFEQHID
KCLDLSQYLYNKIKNREGYEMVFDGVPQHTNVCFWYIPPSLRGMPDGDERREKLHRVAPKIKAMMMESGTTMVGYQPQGNKVNFFRMVVSNPSA
TQSDIDFLIDEIERLGHDL
>E_A0A319ELX8
MDLKDIESTSQEQGRLHQKLWEMTQTWQPDSVIPSASDLSRARASLPKSLTENGAGFENTTQHILNDLVPAFNRSSISPNYYGFITGGITPAAL
FADNLVSAYDQNVQVHMTEHSISTDVEHNALGLLADLLRLDRSQWHNGTFTTGATGSNIQGIACGREFVLRAAAKKKGISIDSVGEYGLFELIH
AAGLSGVQVLTTLPHSSLTKAAGILGIGRANVKSVCRDDHYLQFDMEKVEAELARSDKASIIAVSCGEVNTGHFATSTLSEMENLRRLCDKYGA
WIHVDGAFGIFGRVLEGPEFSTISKGCEGMELADSIAGDGHKLLNVPYDCGFFLCRHAGEANNVFQNANAAYLNGAQSGGPSIPSPLNIGMENS
RRFRALPAYASLVAYGRTGYRQMLERQIRLARMLTEWLYDHPKYTALPKLDNKEALLDQTYISVLFSAKDEELNSNLPKRINETSKMFVSGSAW
QGRPACRIAISNWRVVEDRDFAVVTAVLDGVAGGRS
>E_A0A2I0MD22

MEPEEYRRRGXXXVDYICQYLSNVRERRVTPDVQPGYMRAQLPDSAPMDPDSWDNIFGDIEKIIMPGVVHWQSPHMHAYFPALTSWPSLLGDML
ADAINCLGFTWASSPACTELEMNVMDWMAKMLGLPDKFLHHHPDSVGGGVLQSTVSESTLVALLAARKNKILEMKISEPDTDESSLNSRLVAYA
SDQAHSSVEKAGLIALVKMKFLPVDENFSLRGETLKKAIAEDRKKGLVPVFVCATLGTTGVCAFDNLSELGPICDAEGLWLHIDAAYAGTAFVC
PEFRLFLDGIEYADSFTFNPSKWMMVHFDCTGFWVKDKYKLHQTFSVNPVYLRHPNSGAAVDFMHWQIPLSRRFRSLKLWFVIRSFGVKKLQAH
VRHGTETAKFFESLVRSDPLFEIPAKRHLGLVVFRLKGPNWLTEKLLKELSSSGKLFLIPATIHDKFIIRFTVTSQFTTREDILQDWNIIQHTA
AQIVSQNYELHCINSGAGARIPNMIVKPSSDAISNASQLYLDEGKHKIPSRKTEVQPKKLAESPSMCVISQQVKGQGDPLDDCFPEDVQDVTKN
KLTSFLFSYLSVQGKKKTARSLSCNSVPMTGSLEQCNPKAAATDKKESRANAKILSRLPEEVMMFKKSAFKKLIKFYSVPSFPECSIQCGLQLP
CCPLQAIV
>E_F4NWP2
MQTTAHISTAEASLHINHKDTDEGFEDDYPEEFPFWYSFPSGEQHIDRHQQKLLFESSSASKAVFPLDLFLKQRSSVDLRIRNDSTVAFDDNHQ
SFHNQDSVLHYFVPTTKSEKLLDQYISGVIEAFLDEPAPIYATAVSRTHSLTGKFGGLSIPDGNRSSVADLESYLNHLKLNVIDKSTRTGSSKM
IGHMTTALPFFHRPLARLLAALNQNVVKIETASTFTNLERQTLAMLHKAFFGSSDEFYTRYAYAPEYALGVMASGGTIANITALWIARNKALAV
NAANGCRGIDKEGYISAMLKYGYKRAVIIGSTLMHYSFKKAADLLGLGEEGLVLIPVDDAFRMRIDVLKAKVEKCVAENTLVIAIVGISGTTET
GSIDPLLDIACIAHKYHIHFHVDAAWGGPLIFSPEHSSKLAGISQADTITVDGHKQLYTPMGLGILLAKCPSLVTFIRKTAGYVIRHDSPDLGK
FTLEGSRPANVLYLHASLNLLGKQGLGTLMTRSVTVVKQMAVRLNLHPSQSFQTLHEPMSNVLLYRYIPSDLRESIADGTYVPNPEDEDRVSEV
TKRLQIYQASCIPSDAVFPEVSTHATDESTNGEAGLHSQQNQLQGGFVSRTRVLFKGFHVNALRVVVANPLTTWRDVEGVISDQLKMGAMIEEE
MKREQMVKRVRDMYPMHRPNAVVPDSAKSCTALAACDGTDTCKPDDKIEWWPGWPFDL
>E_F6ZTZ7
MDGNNSDEEPRLGIEPETFRHAATNMVDYIINYHRDIHKRQTFPDVEPGFMQARLPKEAPDYPESWQEVFSDIETVVMDGMTHWQSPGFFSYYP
ATTSYPSMLADMLCNGISCVRFSWASSPSATELETVMMDWLAKAIGLPECFIHGGHGPGGGVIQGSASESTLMALMAARNKTIRQELSRDKSLR
THDIVARMVAYSSQCTHSCMDRAGVFALVEVRKLPVGKDGVMRGSVLKEAVMKDKDDGRIPMFVCASIGTTPCCTFDDLEEIGKICEEQEIWCH
VDAAYAGAALICPEFRYICKGVERVTSFNFNPHKWLMVQIDCSAMWVRNSDDLINSAEVNPLFLHHKAQDSAIDYRHWQIPLGRPFRSLKLWFV
LRMVGIEGLRSNIRRGVQEAKHLERLIRSDERFEILFPVTLGLVCFKFKHPGLLLEEENSLNERLYQKIHNDKRILLVLAMVNGVYFIRVCTGS
THCSIAQVNKCWNVIKEMAEQL
>E_A9SPU0
MGRLAPARSVVTKPLDGEEFRAMGHQMVDFIADYFRDLETYPVQSQVQPGYLKKLLPESAPQDQDSLEDIFYDMHSKIFPGITHWQSPSFFAYY
PSQTSTASILGEMLSASLSVVGFSWITSPAATELEIIVMDWLAKMLQLPSEFLSTGNGGGVIQGTACEAILVVMLAARKRAIARAAAEQGISEA
EALGKLTVYTSDQAHACVNKASQLAGIATKNLRLIHADASTNYAVCADKVAKCVAADKAAGLIPFFLVGVIGTTSSAAVDPLSDLGDIAQEHSL
WYHIDGAYAGNVCICPEYRPLLNGVEKADSFDMNLHKWFLTNFDCSCLWVKDRSPLLAALTTNPEYLRNKQSEANAVVDFKDWQIPLSRRFRAL
KLWMVLRMHGSDFLQTYLRSHCEQAKHFETLVRADSRFELMSQRIFSLVCFRVKPAAGDKDNGYTLNKKLVEALNTGGDIMLTHTTLEGVYTIR
FAIGAARTEMRHIVAAWKEIQRQTSKLLKC
>E_A9SPU5
MGRLAPARSVVTKPLDGEEFRAMGHQMVDFIADYFRDLETYPVQSQVQPGYLKKLLPESAPQDQDSLEDIFYDMHSKIFPGITHWQSPSFFAYY
PSQTSTASILGEMLSASLSVVGFSWITSPAATELEIIVMDWLAKMLQLPSEFLSTGNGGGVIQGTACEAILVVMLAARKRAIARAAAEQGISEA
EALGKLTVYTSDQAHACVNKASQLAGIATKNLRLIHADASTNYAVCADKVAKCVAADKAAGLIPFFLVGVIGTTSSAAVDPLSDLGDIAQEHSL
WYHIDGAYAGNVCICPEYRPLLNGVEKADSFDMNLHKWFLTNFDCSCLWVKDRSPLLAALTTNPEYLRNKQSEANAVVDFKDWQIPLSRRFRAL
KLWMVLRMHGSDFLQTYLRSHCEQAKHFETLVRADSRFELMSQRIFSLVCFRVKPAAGDKGNGYTLNKKLVEALNTGGDIMLTHTTLEGVYTIR
FAIGGARTEMRHIDAAWEEIQRQTSKLLKC
>E_K1X4T2
MDSKQFKEAATSAIDEIVNYYETIEDRRVVSNVEPGYLKKLLPDGPPQDGESWGDIQKDIESKIVPGLTHWQSPNFMAFFPASSSFPGMLGELY
SAAFTAPAFNWICSPAVTELETIVLDWLAKLLNLPDCYLSTSHGGGVIQGSASEAIVTSMVAARDKYLRETTSHLSGAELEDAIAYKRSKIVAL
GSEAAHSSTQKAAQIAGVRYRSIPVSKDTNFALTGAGLEEMLKQCKAQGLEPFYLTTTLGTTATCAVDDFGSIAATLAKYAPPNVTGEIWVHVD
AAYAGAALVCPEYQHLTASLEHFHSFDMNMHKWLLTNFDASCLFVRKRKDLIDALSIMPSYLRNEFSESGLVTDYRDWQIPLGRRFRSLKIWFV
MRTYGVNGLQAHIRKHVKLGEMFADLLRTREDLFKIVTGPTFALTVFTIVPKIAGKEEQDAITKAVYELINKRGEIYITSSVVAGEYVIRVVSA
NPMAEEKFLKKAFDILVDTAEELRDGRPSRRGVNGAIVNGKGEGVGEVAVLNGNGAAAV
>E_C3Y5S8
MNDNVVIIYWTISSTHLVVKDFKDRLVSAGVNNGHPTFFGFINAGGGTYPASLGAFVPAALATYSGHAGNISSGGSLATLTALAVARDSRELKA
ADFHRCVVYCSEFTHYAVQKGLRAVGMREAILRNTPVDKALKMTAAALERQINEDKEAGLLPFLVVATVGTTLTGSVDPVNDIADVCDRHQLWL
HVDAAYGGFFALCDEVKQLFVGVERSDSIVVNPHKGLFTSGGVGVLVVKDGEKLQQCCSLEQTFHFFKGHSIFSAENVSPSELSFELTRPFRGA
QMWLPLKVFGVGVFRTALEEKLLLARYFHRKLKETGEFELPLEPELSVVVFRATAPPGVDINNFNQQLLDDLISDGKIFITPAVFSDQYYLRVC
VLCFKTHIEHIDMCFSLIAMAGANHNLRQRLADLKAKTSVLEVQEVDRDAMTKKTVQLINAYDDNVTKLTYGPIKMLDKGSLETRDDISENPVD
FDSVMQDLNDRLVRAGVIPGHPMFLGFIPSGHGTYPAALGGFLPAAFTTYSVVHLESPAAVQMENRLIKWVADLFGYPAGHAGNISSGGSLATL
TALAVARDSRELKATDFHRCVVYCSEFTHYAVQKGLRAVGMREAILRNTPVDKALKMTAAALQRQINEDKKAGLLPFLVVATVGTTLTGSVDPV
NDIADVCDRHQLWLHVDAAYGGFFALCDEMKQLFCGVERSDSIVVNPHKGLFLPSGVGVLVVKDGKKLQQCCFSEQTSSVFKGMQFFSADHLSP
CELSFELTRSFRGAQMWLPLKMFGVGVFRAALEEKLLLARYFYGKLKETGEFELPLEPELSVVVFRATAPPGADINNFNQQLLNDLVSDGKIFM
TPAELYGQFYLRVCVLCFRTHIEHIELCLALIREKRGHLWGSFQSNA
>E_F7VTW6
MAAANNTPSLPDHMQPAYVDRILKPLESRISELLDEFCPSSQGAERLNDIKKLVVNYVVLQILTQTFDPNLCVTPRPKDLAHAEAWVENYCHDS
SIHLGGEPPQITGLAGHGKEFYSTLTHIIADIVPALNSQALSSRYYGFVTGGVHPVAQAADNVVTALDQNVQVHLPTTHSISTVVEHHALSMLR
SLLALDGFQGKTFTTGATASNIMGLACGREAVISARLPDYAKTGGVGELGLVGACMAAGVREVQILTSKGHSSLYKAASVVGLGRAVVKIWGFP
DTPWVLDLKAVERELKREGVASLIVVSAGEVNTGQFGTSGNAMKELRKLADRYKAWIHVDGAFGIFARALPKVDRFAKLRERTAGLELADSIAA
DGHKLLNVPYDNGIFLCSKPEVMSQVFQNPNAAYLAPVASTSRSGTDTEIQSPLHVGLENSRRFRALPVYALLLHLGRENMGEMLARMVDLARE
IAAFIRDSDKYELLPDETGADIECTHIIVLFKAKNPDLNEVLVEKINETRRMYVSATKWNGENAVRIAVGSWKVDVEEDFKAVKSVLNSL
>E_H9GNM5
TSRRSTKVLDFHHPHQLLEGMEGFNLELSDNPESLEQILVDCRDTLKYGVRTGHPRFFNQLSTGLDIIGLAGEWLTSTANTNMFTYEIAPVFVL
MEQITLRKMREIIGWTDKDGDGIFSPGGAISNMYSIMAARYKYYPEVKTKGMAAVPKLVLFTSEHSHYSIKKAGAALGFGTENVILIKCNERGK
IIPADLEAKILDAKQKGYAPLYVNATAGTTVYGAFDPIHEIADICEKYNLWLHVDAAWGGGLLMSSKHRHKLNGIERANSVTWNPHKMMGVLLQ
CSAILLREKGILQGCNQMCAGYLFQQDKQYDITYDTGDKAIQCGRHVDIFKFWLMWKAKGTVGFETQINKCLELSEYLYNKIKNREEFEMVFKG
EPEHTNVCFWYIPPSLRGMPDSEERREKLHRVAPKIKALMMESGTTMVGYQPHADKVNFFRMVVSNPAATKSDIDFLVEEIERLGQDL
>E_A0A364MZ99
MDARSQETFEQLAATVATIHSQPSPEHVLPSGATLSSARSKLQPHLPTQGVGLEESIRHLREDLAPGFNASSRSANYYGFVTGGTNQAAALADN
LVTAYDQNVQVHLPNETIATGLEDRALSLLCELLNFEPGQWPHRIFTTGATAANVLGLACGREYVIAEASAHRTDAENSVGEVGIVDAMRTAGI
DDIQVLTTVPHSSLSKAASILGLGRASVKCLGCSDVPHKFDMNLLKKALEQPGAASIVVISASEVNTGVFATSGLEEMQELRKLCDMYGAWIHI
DAAFGLFGRVLSSPAYSSIIQACDGLELADSITGDGHKLLNVPYDCGFFLSRHRNMAERVFQNPNAAYLASGNGSDSIMSPLNVGLENSRRFRA
LPVYASLVAYGRDGYRDMLERQIQLSRGIAQYILQSSEYELLPQKDASHEEILDGIYIIVIFRAKDEALNKQLVTKIKATRKIYVSGTSWEGKP
ACRFAVSNWQTNVERDLPIIKQVLQGIL
>E_Q0U7M3

MESARSEYQELLRSAIAGLQLEPSPTHVLPNQKAVEHARKTLANQLREDGLGLSETIRHLQEDLKPAFNASSRSPNYYGFVTGGVTPAASIADN
FVTAYDQNVSVHLPDESIATDVEDHALSLLCELLNFQPDQWPHRIFTTGATASNVVGLACGREYVIAEASAHRTHTENSVGEYGIMEAMHRAGL
DNIQILTTVPHSSLSKAASVLGMGRASIKHVGREDAPHRFDFKLLKSLMEQPSSASIVAVSASEVNTGLFATSVLEEMEELRRLCDMHGAWIHV
DGAFGILGRLLDSPRYQTITQACAGLELADSITGDGHKLLNVPYDCGFFLSRHRHIAERVFQNPNAAYLATSNGPDSIMSPLNVGLENSRRFRA
LPVYASLIAYGKSGYRDMLERQIDLSRGIAKFILESETYELLPESDKSMEERLSTIYIIVLFRAKDGDLNKNLVDKIKATKKIYVSGTAWDGKP
ACRFAVSNWQTDVTRDLPIIEEVLQSVAE
>E_A0A369RT79
MDMSLDGLSDPQFQQMYDDEFQSSLNKNAHIDDFIHAMVQIMVKYMKRCHDRHEKVVEFRQPEELNKLISFELKDTPEDLATVIDYCRETLNYC
VKTGHPRFFNQLYGGMDIVGLCGQWLSATANTSMYTYEVAPIFVLMERAVLRHMGQLVGFSDGDGIFAPGGSLSNMYAISLARHKKFPTSKMEG
LFSLPRMAILTSKHVRSKANQKKFLKPLSSDGAYFMGFGLNNVVMINCDAKGRMLASDLENQIIHLQSQGIAPILVNATSGTTVFGAFDPLDEI
ADICQKYDLWLHVDAAWGGAIILSAEKRHLMKGIHRIDSISWNPHKFMGCPFQCSAFLTKKKGLLEECHGIPASYLFQKDKMTYDISYDTGNKS
IQCGRHVDIMKLWLMWKAKGDQGFTKKLHHAYEISNYLTEKIRNRDGFELVVEPVYVSICFWYIPTAIRNLSDGDVKRRKLSQVAPQIKAGMTK
RGSMLVGYQPMDDKVNFFRMIILNSATTFDDMDFILDEIESLGERIEFVQNDRLYRLVSRQVGDMDNSFQELVDQKFQHLYDDEIQKNFHQNPH
LEDFVHAMVNILIKHMKNCHDRSQKVIEFKEPGELKQLINFELREKPESMAAVMEYCRQTLHYCTKSGHPRFYNQLHSGIDVVSVCGEWLAATV
STNMATYEIAPTFLLMEEAVLRHMTQLIGFHDGDGIFAPGGGGSLSNMIAINLARYRKFPASKLKGLFSLPRMAVLTSNHSHYSFQKGSHFMGL
GQENAVIVNCDSEGRMSICDLEDKIVHLLSQDIVPIMVTATCGTTVYGAFDPVDEIANLCQRYDIWFHVDASWGGAALFSDRKRHLMKGVHRAD
SVTWNAHKFMGCPFLCSVLLTKTKGILQECNEIVAPYLFQQDKMTYDVSYDTGNKTIQCSRRIDIMKLWLMWKAKGDEGFTKKVNHACELANYL
IEKIRNREGFKLVHQPMYLNVCFWYIPKALRDMADDEIKRAKLSKVAPQIKAGMTKRGSMMVGFQPVDDKVNFFRMILINYNTTLEDMDFILDE
IETLGEAVKV
>E_Q24062
MLASENFPTHHFKESIFKPYSTTSGDDLASVSPLTATAALVASTSSPADSTSTVAFEQAS
KMLANAANNNNNNNNNITSTKDDLSSFVASHPAAEFEGFIRACVDEIIKLAVFQGTNRSS
KVVEWHEPAELRQLFDFQLREQGESQDKLRELLRETIRFSVKTGHPYFINQLYSGVDPYA
LVGQWLTDALNPSVYTYEVAPLFTLMEEQVLAEMRRIVGFPNGGQGDGIFCPGGSIANGY
AISCARYRHSPESKKNGLFNAKPLIIFTSEDAHYSVEKLAMFMGFGSDHVRKIATNEVGK
MRLSDLEKQVKLCLENGWQPLMVSATAGTTVLGAFDDLAGISEVCKKYNMWMHVDAAWGG
GALMSKKYRHLLNGIERADSVTWNPHKLLAASQQCSTFLTRHQQVLAQCHSTNATYLFQK
DKFYDTSFDTGDKHIQCGRRADVFKFWFMWKAKGTQGLEAHVEKVFRMAEFFTAKVRERP
GFELVLESPECTNISFWYVPPGLREMERNREFYDRLHKVAPKVKEGMIKKGSMMITYQPL
RQLPNFFRLVLQNSCLEESDMVYFLDEIESLAQNL
>E_A0A2V1BNR7
MDSKQFKEAAASAIDEIVSYYDTIEDRRVVSNVEPGYLKKILPEGPPQDGESWADIQKDI
ESKIMPGLTHWQSPNFMAFFPASSSFPGMLGELYSAAFTAPAFNWICSPAVTELETIVLD
WLAKLLNLPDCYLSTSHGGGVIQGSASEAIVTVMVAARDKYLRETTAHLSGLELEDAVAY
KRSKIVALGSEAAHSSTQKAAQIAGVRYRSIPVSKENGFALTGSALAQVLDECKAQGLEP
FFLTTTLGTTATCAVDDFEAIATTLAKYAPPNIPGEIWVHVDAAYAGAALVCPEYHHLTA
QFEHFHSFDMNMHKWLLTNFDASCLYVRKRKDLIDALSIMPSYLRNEFSESGLVTDYRDW
QIPLGRRFRSLKIWFVLRTYGVNGLQAHIRKHVKLGEVFAGLLETRKDLFEIVTGPNFAL
TVFTVVPKGEGKGKEEKDAVTREVYELVNKRGEIYITSSVVGGEYVIRVVSANPLAEEKF
IRKAFGILVETAEEVRSGKGGLKGAGHVVDLNVNGVGEVVELSGNGVAK
>E_A0A177DW43
MEVTDNELFQRLSAAIAKIHSGPPPADVLPSTTTLNDARSKLQPHLPSQGVGLEESIRHL
QEDLAPAFNGSSRSPNYYGFVTGGTTPAALLADNLVSAYDQNVQVHLPNETIATDVEDRA
LSLLCELLDFEPEQWPHRIFTTGATSANILGLACGREYVIAEASAHRTDAEISVGELGIV
EAMRSAGIDDIQILTTVPHSSLGKAAGILGLGRTSVKCLGQSEAPHKFDMQLLKKSLEQP
GAASIVAVSASEVNTGVFATSGLEEMEEIRKLCDMYGAWIHVDGAFGLFGRLLTSPTYSS
IAQACGGLNLADSITGDAHKLLNVPYDCGFFLSRHRNIAERVFQNPNAAYLASGNDANAI
MSPLNIGLENSRRFRALPVYANLVAYGREGYRTMLERQVDLSRGIARYILESDGYELLPK
SSACHEDILKGIYIIVLFRAVDEELNKQLVNKIKATRKIYVSGTAWDGKPACRFAVSNWM
TDVKRDLPIIKQVLQDVVLGKESK
>E_A0A091WHQ1
GKEMVDYICQYLSNVRERPVTPDVQPGYMRAQLPDSAPMDPDSWDNIFGDIEKIIMPGVV
HWQSPHMHAYFPALTSWPSLLGDMLADAINCLGFTWASSPACTELEMNVMDWLAKMLGLP
EKFLHHHPDSVGGGVLQSTVSESTLVALLAARKNKILEMKLSEPDADESSLNSRLIAYAS
DQAHSSVEKAGLISLVKVKFLPVDENFSLRGEALKKAIAEDRSKGLVPVFVCATLGTTGV
CAFDNLSELGPICDGEGLWLHIDAAYAGTAFVCPEFRLFLDGIEYADSFTFNPSKWMMVH
FDCTGFWVKDKYKLHQTFSVNPVYLRHPNSGAAVDFMHWQIPLSRRFRSLKLWFVIRSFG
VKKLQAHVRHGTETAKFFESLVKSDPLFEIPAKRHLGLVVFRLKGPNWLTEKLLKELSTS
GRLFLIPATIHDKFIIRFTVTSQFTTREDILQDWNIIQHTAAQIVSQNYGLHCMSSGDEA
RIPNMIVKPSSDAISSASQLCLGGGKYKTPSRKIVVQPKKVAVSPSMCVISQQVKGQGDP
LDDCFPEDVRDVTKHKLTSFLFSYLSVQGKKKTARSLSCTSVPMTGSLEQCNPKAAATDK
KESHANARILSRLPEEVMMFKKSAFKKLIKFYSVPSFPECSIQCGVQLPCCPLQAIV
>E_A0A0N0BFM6
MPANEETLSVSSDQIRNNFLGKACDNGLQNNSVRVSSNDDEENCQNCCLESIINKDSKEN
CNYKSLPVREIHEKFMKSFVDLLLEEAVFEGTSRKNRVVEWMEPAALQSAIDLKLSDQGS
SHKKLVTLARNVIKYSVKTGHPRFINQLYSSVDPYGLLGQWLTDALNPSVYTYEVSPVFS
LMEEEILYEMRNIVGWKDGRGEGIFCPGGSIANGYAINLARHYRFPELKELGLSSAGRLI
VFTSRDSHYSVKKLTAFLGLGTSNVYEVKTDSRGKMCVTDLEAQIKKALDEGAVPLIVSA
TAGTTVLGAFDPLKKIALICKKYNLWYHVDAAWGGGTLMSKKYRHLLAGVELADSVTWNP
HKLLAAPQQCSTLLLRHEGLLQAAHGSKANYLFQPDKFYDTSFDSGDKHIQCGRRADVLK
FWFMWKAKGTRGLEKHVDRVFELARYFTNYIRHREGFKLILEPECTNVCFWYVPPSKRHL
QTDELSKALQKIGPAVKERMMKKGSMLITYQPLRELPNFFRLVLQNSGLTEADMRFFAEE
IERLAVDL
>E_R7Z469
MQRVYAAELGLDRLLELSTGADLLPSKEVLQHARSSLIPAISKEGLGLSNTVKHLKEDVV
PALNLASQLPNYYGFITGGVTPVAALADNIVTAYDQNVQVHLPEETIATEVEDRALRMLC
ELLDFGPTEWPHKTFTTGATASNVLGLACGREFVIAEAARRTKKGPVNVGDLGLAKALRL

CGLDDIRILTTVPHSSLGKAASITGIGRDSITLVGLPNAPHKFDMTKLQRILQERTSGYI
VAISCAEVNTGLFATSSAAEMLELRELCDKYGAWIHVDGAFGLLVRVLEDHQYLNIRAGC
DGIQFADSITGDAHKLLNVPYDCGFFLGKHRALAENVFRNSNAAYLNVGATDTSFIASPL
NIGIENSRRFRALPVYASLTAYGRRGYRDMLERQVKLARGIARFILRHSEYDLLPNLEVD
DERRLDSIYIIVLFRAKDDSLNKDLVKHVNATKKIYVSGTAWTGAPACRFAVSNWQVDVD
RDLQLVTDVLDRVALAASHA
>E_A0A1D5PSZ5
MQKGLCYNKREEWKEMVDYICQYLSNVRERRVTPDVQPGYMRAQLPDSAPMDPDSWDNIF
GDIEKIIMPGVVHWQSPHMHAYFPALTSWPSLLGDMLADAINCLGFTWASSPACTELEMN
VMDWLAKMLGLPDKFLHHHPDSVGGGVLQSTVSESTLVALLAARKNKILEMKLSEPDADE
SSLNSRLIAYASDQAHSSVEKAGLISLVKMKFLPVDENFSLRGETLKKAIAEDRKKGLVP
IFVCATLGTTGVCAFDSLSELGPICGAEGLWLHIDAAYAGTAFLCPEFRLFLDGIEYADS
FAFNPSKWMMVHFDCTGFWVKDKYKLHQTFSVNPVYLRHPNSGAAVDFMHWQIPLSRRFR
SLKLWFVIRSFGVKKLQAHVRHGTETAKFFESLVRSDPLFEIPAKRHLGLVVFRLKGPNW
LTEKLLKELSSSGRLFLIPATIHDKFIIRFTVTSQFTTREDILQDWNIIQRTAAQIISQH
NELHRVSSGDEAKIPNMISEPSSSVISHASQLYLEGKYKIPSRKIVGQPKKLAASPSTCV
ISQQVQGQGDLDDCFPEDVQDVTKHKLTSFLFSYLSVQGKKKTARSLSCNSVPMTDLLEQ
CNPKTTDSDQKEPHANGRILSRLPEEVMMLKKSAFKKLIKFYSVPSFSECSIQCGLQLPC
CPLQAIV
>E_F1NXM1
MEPEEYRRRGKEMVDYICQYLSNVRERRVTPDVQPGYMRAQLPDSAPMDPDSWDNIFGDI
EKIIMPGVVHWQSPHMHAYFPALTSWPSLLGDMLADAINCLGFTWASSPACTELEMNVMD
WLAKMLGLPDKFLHHHPDSVGGGVLQSTVSESTLVALLAARKNKILEMKLSEPDADESSL
NSRLIAYASDQAHSSVEKAGLISLVKMKFLPVDENFSLRGETLKKAIAEDRKKGLVPIFV
CATLGTTGVCAFDSLSELGPICGAEGLWLHIDAAYAGTAFLCPEFRLFLDGIEYADSFAF
NPSKWMMVHFDCTGFWVKDKYKLHQTFSVNPVYLRHPNSGAAVDFMHWQIPLSRRFRSLK
LWFVIRSFGVKKLQAHVRHGTETAKFFESLVRSDPLFEIPAKRHLGLVVFRLKGPNWLTE
KLLKELSSSGRLFLIPATIHDKFIIRFTVTSQFTTREDILQDWNIIQRTAAQIISQHNEL
HRVSSGDEAKIPNMISEPSSSVISHASQLYLEGKYKIPSRKIVGQPKKLAASPSTCVISQ
QVQGQGDLDDCFPEDVQDVTKHKLTSFLFSYLSVQGKKKTARSLSCNSVPMTDLLEQCNP
KTTDSDQKEPHANGRILSRLPEEVMMLKKSAFKKLIKFYSVPSFSECSIQCGLQLPCCPL
QAIV
>E_U4TX00
MADFCPVPNPDVHGDFLQKVVDLLFKNVVFTTRRDKVLLWKTPEELQGEFDFTLPQAGDS
QDKLLQIMKNTVKFSVKTGHPYFINQLFSGLDPYGLAGQWLTDALNASVYTYEVAPVFTL
MEQHVIKEVCKMVGPQWSDGIFCPGGSFGNGTAMNLARFHKFPQAKLKGCNILPRMVVLA
SEECHYSTYKFAAFLGMGEDNVWPLKTDDVGQIVPEKVEEAIQAVLLEGAVPLMVVATLG
TTVRGAFDPIHAIATICKKYEIWLHVDAAWGGGLLFSQKHRAKLNGIEAANSIVINPHKL
LAVPQQCSMLLVTHPDILHRCHSRGAEYLFQKDKYYDASYDLGDRYLQCGRKCDVFKFWI
MWRAKGSCGFAKHIDTLMDLAEYFEQQVNARPDFVLVSKRQYVNVCFWYLPRYLRGKLNI
SDYNAQLQKVAPQIKAVMVKHGSVMLNYQPLKTLPNFFRFVSQNSALTKKDADFILDHIA
EIGDALFP
>E_U4UGN6
MDCDEFRCRGKEMIEYICSYLDNIEERRVTPNIEPGYLRKLIPENAPEDPENWDSIMADV
ESKIMPGVTHWQHPRFHAYFPSGNSFPSILGDMLSDAIGCIGFSWAASPACTELETVVLD
WLGKAIGLPDQFLALKEGSRGGGVIQTSASECVLVSMLAARAQALKRLKQQHPFVEEGLL
LSKLMAYCSKEAHSCVEKAAMICFVKLRILEPDETSSLRGKTLLLAMEEDETMGLIPFFV
STTLGTTSCCSFDNLPEIGVISHKFPCVWLHVDAAYAGNAFICPELKYLLKGIEHADSFN
TNPNKWLLTNFDCSTMWVRDRIRLTSALVVDPLYLQHGYSDATIDYRHWGVPLSRRFRSL
KLWFVIRSFGISGLQKYIRLHIRLAKRFEAHVLKDRRFEICNEVKLGLVCFRLRGQDKLN
EKLLSNINASGKIHMVPANVNEKYVIRFCVVSHNATEQDIDHAWHVISQFADDLLEMQNA
DKEQDEVYELLERKKKETLAQKRSFFVRMVSDPKIYNPSIAKALPSTRRHLTQPSAESPE
NHITIQPPTDNRERNCLKMDVEEFRTRGKEMVDYICKYMKNLPNQRVTPHLEPGYLKELI
PSEAPVEPENWNDIMEDVDKKIMPGITHWQHPRFHAYFPSGNSYPSILADMLSNAIGCIG
FSWAASPACTELETIVMDWFGKAIGLPHDFITSNKGSTGGGVIQTSASECVLISMLAARN
QAIQYLKKNLIDPNEIEDSAFLPKLVGYCSKEAHSCVEKAAKILLVKVRILEPDEKGSLD
KQTLEDAICKDKENGLFPFFVSTILGSTSSCSFDNLQVIGPICKAEPCIWLHVDAAYAGN
AFICPELKPYLKGIEYADSFNTNPNKWLLVNFDCSCLWVRCRVKLTSALVVDPLYLQHAN
SNESIDYRHWGIPLSRRFRSLKLWFVIRKYGLVGLQKYIRNHITLAKHFETLVSKDNRFE
VLNDVRLGLVCFRLVANDQVNQELLANINASGKLHMIPSMVKNKYIIRFCINAEDAKQED
VEDAWRIITEHASEILEPTNNKKKEFTKPPLTRQMSKRFAFTRSVSKELYKRCRSRSNLL
DGATPILVADSDEEDVEVQQGNNCEAVDFLASSLDDNVFLDEYKKDF
>E_G7XA98
MDLRDIKGTSQEQGQLHQKLWELNQTYTSGNVLPSASDLSRARASLPESLPAEGTGFESA
TQHILNDLVPAFNRSSISPNYYGFVTGGVTPAALFADNLVTAYDQNVQVHLAEHSISTDV
EATALGLLADLLKLDRRSWNNGTFTTGATASNVHGLACGREFVLRAAAKKSGIDIDSVGE
YGLFEIIQATGISGIQILSTMPHSSLAKAAGILGIGRANVKSICRENHYLQIDFERLEAE
LKASKASIVAVSCGEVNTGHFATSSLVEMKRLRSLCDKYGAWLHVDGAFGIFGRILNGAE
FSAISKGCEGMELADSIAGDAHKFLNVPYDCGFVLCRHPGEAEKVFQNANAAYLSGGQSG
ALPIPSPLNIGMENSRRFRALPVYASLLAYGQIGYQEMLQKQVRLSRMIAGWIFDHPKYN
ALPDLSSKDELLDHTYIIVLFSAKDDELNSNLTKEINATSKLFVSGTSWQGRPACRIAIS
NWRVEEERDFALVTDVLDTVAGRV
>E_B2W8T8
MEAPNQASFDKLATAIASIHFQSPPEDVLPSAATLSSARSKLQTHLPTQGIGLEESIRHV
QEDLAPAFNASSRSPNYYGFVTGGTTPAAALADNLVTAYDQNVQVHLPNETIATDLEDRA
LSLLCELLNFDAAQWPHRIFTTGATAANVLGLACGREFVIAEASAHRTDAENSIGEVGIV
EAMRKAGVDEIQILTTVPHSSLAKAAGILGLGRTSVKCLGRSDAPHKFDIQLLKKSLERP

HAASIVAISASEVNTGAFATSSLEEMQDIRKLCDMYGAWIHVDGAFGLFGRILSSSVHSS
IIQACTGLELADSITGDGHKLLNVPYDCGFFLSRHRNIAQHVFQNPNAAYLASGNSADSI
MSPLNVGLENSRRFRALPVYASLVAYGRDGYRDMLERQIRLSRGIAEHILESKDYELLPR
SDASREDLLSGIYIIVLFRARDEELNKQLVDRIKATRKIYVSGTSWEGKPACRFAVSNWM
TDVIRDLPIVKQVLRDVALEKNGR
>E_A0A1L9V1T0
MDLKNIEATSQEQGRLHQKLWELSQTYTPGNVLPSASDLSKARASLPKSLPAEGAGFESV
TQHILNDLVPAFNRSSISPNYYGFVTGGVTPAALFADNLVTAYDQNVQVHLAEHSISTDV
EATALGLLADLLKLDRRHWNNGTFTTGATASNVYGLACGREFVLRAAAKKRGIAIESVGE
YGLFELIQATGISGIQILSTMPHSSLAKAAGILGIGRANVKSICRDDHYLQIDLVRLEAE
LKANKASIVAVSCGEVNTGHFATSSLAEMQSLRSLCDKYGAWLHVDGAFGIFGRILEDAE
FSTISKGCEGMELADSIAGDAHKFLNVPYDCGFVLCRHPGEAEKVFQNANAAYLSGGQSG
ALSVPSPLNIGMENSRRFRALPVYASLVAYGQVGYQEMLQKQIRLSRMIAGWIFDHPKYN
ALPELASKNKLLDQTYIIVLFSAKDDELNRNLTKEINATSKLFVSGTSWQGRPACRIAIS
NWRVEEERDFALVTDVLDKVAGSV
>E_A0A0V0YNL2
MDISENRMIEENQKFMENKFYNDILPLNRSGWLTTEMFIKSVVDLLLQFIQETNNPNTKV
INFHHPTELIAKLDLRIPINPTNLQKVLEDCKEVLKYQVRTGHPRFFNQLSTGLDLISMI
GEWLTATVNTNMFTYEISPVFVLMEKEIIETMCEIVGWPSGKRDGIFSPGGAISNLYAVN
AARHYMFPRCKAIGMVETPNLAMFTSEDSHYSIRGAAALVGIGVDNCFPIPVDEKGKMIP
SKLEEEVILAKKNGYVPFFVCAVGGTTVYGAFDPINEIANICKKYRMWLHVDAAWGGGIL
LSKKHRHLANGIERADSVTWNPHKLMGALLQCSACLIRHEGLLFQCNQMCADYLFQQDKP
YDVSYDSGDKAIQCGRHNDVFKLWIMWRAKGMNGFEQQVNRLMDLANYFTEKIKKTPGYE
LIMENPEFLNICFWYVPKNVRHLESTEKKARLDKVAPKIKAKMMSSGSTMVGYQPDKDKP
NFFRMIISNPATTYEDLDFFIEEIIRLGESL
>E_A0A0V0YMZ9
MDISENRMIEENQKFMENKFYNDILPLNRSGWLTTEMFIKSVVDLLLQFIQETNNPNTKV
INFHHPTELIAKLDLRIPINPTNLQKVLEDCKEVLKYQVRTGHPRFFNQLSTGLDLISMI
GEWLTATVNTNMFTYEISPVFVLMEKEIIETMCEIVGWPSGKRDGIFSPGGAISNLYAVN
AARHYMFPRCKAIGMVETPNLAMFTSEDSHYSIRGAAALVGIGVDNCFPIPVDEKGKMIP
SKLEEEVILAKKNGYVPFFVCAVGGTTVYGAFDPINEIANICKKYRMWLHVDFRMFQAAW
GGGILLSKKHRHLANGIERADSVTWNPHKLMGALLQCSACLIRHEGLLFQCNQMCADYLF
QQDKPYDVSYDSGDKAIQCGRHNDVFKLWIMWRAKGMNGFEQQVNRLMDLANYFTEKIKK
TPGYELIMENPEFLNICFWYVPKNVRHLESTEKKARLDKVAPKIKAKMMSSGSTMVGYQP
DKDKPNFFRMIISNPATTYEDLDFFIEEIIRLGESL
>E_H0V9C5
MADSEPLPSLDGDPKAAEAWLRDVFEIIMDEAIHKGTRTSEKVCEWKEPEELKQLLSLEL
QTHGEAQKQILEHCRAVIHYSVKTGHPRFFNQLFSGLDPHALAGRIITESLNTSQYTYEI
APVFVLMEEEVLKKLRALVGWSSGDGVFCPGGSISNMYAMNLARYQRYPDCKQKGLRALP
PLAIFASKECHYSVNKGAAFLGFGTDSVRVVESDERGKMIPGDLERQIKLAEAEGAVPFL
VSATSGTTVLGAFDPLNIIADVCQRHGLWLHVDAAWGGSVLLSQTHRHLLDGIQRANSVA
WNPHKLLGAGLQCSVLLLQDTSNLLKRCHGSQASYLFQQDKFYDVALDTGDKVVQCGRHV
DCLKLWLLWKAQGGQGLERRIDRAFALAQYLVEEIKKRKGFELVIEPEFVNVCFWFVPPS
LRGKQESPDYSQRLSQVAPVLKERMVKKGSMMIGYQPHGTRGNFFRMIVANPMLTRADID
FLLDELERLGQDL
>E_H0V2I4
MASSTPSSSATSSNAGADPNTTNLRPTTYDTWCGVAHGCTRKLGLKICGFLQRTNSLEEK
SRLVSAFKERQSSKNLFSRENSDRDTHFRRAETDFSNLFARDLLPAKNGEEQTVQFLLEV
VDILLTYIRKTFDRSTKVLDFHHPHQLLEGMEGFNLELSDHPESLEQILVDCRDTLKYGV
RTGHPRFFNQLSSGLDIIGLAGEWLTSTANTNMFTYEIAPVFVLMEQITLKKMREIVGWS
DKDGDGIFSPGGAISNMYSIMAARYKYFPEVKTKGMAAVPKLVLFTSEHSHYSIKKAGAA
LGFGTDNVILIKCNERGKIIPADLEAKVLEAKQKGYVPLYVNATAGTTVYGAFDPIQEIA
DICEKYNLWLHVDAAWGGGLLMSRKHRHKLSGIERANSVTWNPHKMMGVLLQCSAILVKE
KGILQGCNQMCAGYLFQPDKQYDVSYDTGDKAIQCGRHVDIFKFWLMWKAKGTVGFENQI
NKCLELAEYLYAKIKNREEFEMVFDGEPEHTNVCFWYIPQSLRGVPDSLERREKLHRVAP
KIKALMMESGTTMVGYQPQGDKANFFRMVISNPAATQSDIDFLIEEIERLGQDL
>E_A0A1J9RHD0
MSIQPLQSREDKVFPSQIWHTAISPWTSSPLPSTTTLSHVRSSLISLLPNAGLGFSETKR
HIVNDIATGFNGSSLSANYYGFVTGGVTPAALLADNIVSAYDQNVQVHLPDHSVATDVED
RALTFLLDLFDLDHKAWTHKVMTTGATGSNVLGLALGREFIFRAAVERKQGATRKVVKSV
GEHGMAEVLAAAGLKGLQVISTYPHSSIGKAAGILGIGRANVKSICAAGDGRVPLKFDFE
ILEKELARSDMASIVAVSCGEVNTGHFATGGLEEFRKIRQLCDKYDAWLHVDGAFGMFGR
ILKGGGEFDRIWKGCQGLELADSITGDGHKLLNVPYDCGFFLSRHADLAEDVCRNPNAAY
LSAGGGGIPPPCNNGVENSRRLRALPVYATLVAYGRNGYRDMLERQIRLARSVVGWLFEH
PAYAVLPHNPVKESLLQDTFVIVLFRAKDEDLNRVLVSKINAASKIYVSGTSWEGKSACR
IAISNWRVNEEKDFEVITSVLRQIAQ
>E_A0A179G349
MASHELQRLIQREFVSHDNISSVLETLYGAIEMGVQFKAREKVIEHLPLDTLAGKIMQPI
PETGQNLSETLQEFQETVLRHSTNFGSKNFMAFPDCGNSVSALAGAFLMNCMNQNLINSK
HCAPAASMVEINVIQWLRELVGYTVKRDVNNIFDVGGTVVTGGVLANATAMLMARERAFP
GTKDKGTSFDPSRVRVILPQYVEHYSIRASLGWLGLGEENVIRVRTKDFRIDLSDLEHLL
EDNAGNYHFMAIVAYAGDSRSMTIDNFAAVHELAKKHNIWFHIDACHGLQYAFSDALKPR
LGDIHLGDSITIDPHKILFLPYNLSAVLVKDPESFRAISGTSDLIMKENYAFGQMTPFIG
SKAFWSLKLWFTWKTLGRKNIGDLIERRHNLAAYLTEQLQQCRDFLVLNHEVNINSVMFM
YKPSHFQPDSVALDDYVQRLNELNKEIQNTLFCEGDFYVHTFSIPDLGNVVGTGERMLQP
LRYMCGNPLTAETDIDALISRVRKLGQTVEHKYSYHEVNQLRARLA
>E_A0A0A0AJ64

VCQYLSNVRERRVTPDVQPGYMRAQLPDSAPMDPDNWDNIFGDIEKIIMPGVVHWQSPHM
HAYFPALTSWPSLLGDMLADAINCLGFTWASSPACTELEMNVMDWLAKMLGLPDKFLHHH
PDSVGGGVLQSTVSESTLVALLAARKNKILEMKHSEPDTDESSLNSRLIAYASDQAHSSV
EKAGLISLVKMKFLPVDENFSLRGETLKKAIAEDRNKGLVPVFVCATLGTTGVCAFDNLS
ELGPICDAEGLWLHIDAAYAGTAFVCPEFRLFLDGIEYADSFTFNPSKWMMVHFDCTGFW
VKDKYKLHQTFSVNPVYLRHPNSGAAVDFMHWQIPLSRRFRSLKLWFVIRSFGVKKLQAH
VRHGTETAKFFESLVKSDPLFEIPAKRHLGLVVFRLKGPNWLTEKLLKELSSSGRLFLIP
ATIHDKFIIRFTVTSQFTTREDILQDWNIIQQTAAQIVSQNYGLCCISSRDGARIPNMIV
KPSSDALSNASQLYLDGGKYKTPSRKIVVQPKKLAASPGTCVISQQVRGQEGPVDDCFPE
DVQHVTKHKLTSFLFSYLSVQGKKKTARSLSCNSVPMTGSLEQCNPKAAATDKKESHANT
RILSRLPEEVMMLKKSAFKKLIKFYSVPSFPECSIQCGLQLPCCPLQAIV
>E_A0A124BXG1
MDLRDIKGTSQEQGQLHQKLWELNQTYTPGNVLPSASDLSRARASLPESLPAEGTGFESA
TQHILNDLVPAFNRSSISSNYYGFVTGGVTPAALFADNLVTAYDQNVQVHLAEHSISTDV
EATALGLLSDLLKLDRRSWNNGTFTTGATASNVHGLACGREFVLRAAAKKSGIDIESVGE
YGLFEVIQATGISGIQILSTMPHSSLAKAAGILGIGRANVKSICRDNHYLQIDFERLEAE
LKANKASIVAVSCGEVNTGHFATSSLAEMKSLRSLCDKYGAWLHVDGAFGIFGRILNGAE
FSAISKGCEGMELADSIAGDAHKFLNVPYDCGFVLCRHPGEAEKVFQNANAAYLSGGQSG
AFSIPSPLNIGMENSRRFRALPVYASLLAYGQVGYQEMLQKQVRLSRMIAGWIFDHPKYN
ALPELASKDELLDQTYIIVLFSAKDDELNGNLTKEINATSKLFVSGTSWQGRPACRIAIS
NWRVEEERDFALVTDVLDNVAGRV
>E_A0A137PEN3
MDHEEFRQAGYKIIDEIVEYYKTTSQLKPTTDVQPGYLPPLLPKSAPEDPENFDQIHKDF
KDLVVPGLTHWQSSNFFGYFPCTSSYPSMLAEMYSNMFNSQGFDWICNPAASELEGVVMN
WLGQLLGLDDSFLSDKSETKGCGSIQATASEGTLVSMIAARNKVLKKLTDVPSFNDNEEF
RDTKKLVVYTSNQSHSSFVKAAKILNIKIHTLKTDENGSVQPQYLKDQIAQDLGQGLIPF
YVGLCLGTTGVGAVDNILDCLPVAQEHDLWVHVDAAWAGAYLMLPKYQIYNGFKGVDSIV
FNPHKLLLTNFDCCALWTKNRFDLIEALSADASYYTNFATQSGKVIDYKNLQVPLGRRFR
SLKLWFVLRNYGAKGLRGLLQNNLDLAQYLVDLINNDGIFEITHPTTFSLVCFRMKPQIE
SESPDLTNRRNKWIYETISKEGNFLIGHTNFSSKYIFRISIGNQHNTQESVTEVYEKLKS
LAMEVDQQVE
>E_A0A1R3RWV0
MKLKDLKGTSQEQGRLHQKLWELTQTYKVGGVLPSASDLSRARASLPESLTDEGAGFEST
AQHILNDLVPAFNRSSISPNYYGFVTGGITPAALFADNLVSAYDQNVQVHLAEHSISTDV
EATALGLLADLLRLDQHHWSNGTFTTGGTASNIHGLACGREFVLRAVAKRKGIDVDSVGE
YGLFELIHATGLSGIQVLTTFPHSSLTKAAGILGIGRANVKSMCRDDHYLQFDLERLEAE
LARPDKASIVAISCGEVNTGHFATSSLAEMESLRRLCDKYGAWLHADGAFGIFGRVLEGT
EFSTITKGCEGLELADSIAGDAHKLLNVPYDCGFFLCRHSGEAEKVFQNANAAYLTGGQS
GAPSIPSPLNIGMENSRRFRALPAYASLVAYGRAGYREMLQKQIRLSRMIAGWIFDHPKY
NALPELASKDALLDQTYIIVLLSAKDEELNDNLTKRINATSKMWVSGSSWQGRPACRIAI
SNWRVEEERDFALVTDVLDSVAGGRS
>E_H0ZB55
GKEMVDYICQYLSNVRERRVTPDVQPGYMRAQLPDSAPMDPDSWDNIFGDIEKIIMPGVV
HWQSPHMHAYFPALTSWPSLLGDMLADAINCLGFTWASSPACTELEMNVMDWLAKMLGLP
DKFLHHHPDSVGGGVLQSTVSESTLVALLAARKNKILEMQVSEPDTDESSLNSRLVAYAS
DQAHSSVEKAGLISLVKIKFLPVDENFSLRGETLKKAIAEDRKKGLVPVFVCATLGTTGV
CAFDNLSELGPVCDAEGLWLHIDAAYAGTAFVCPEFRLFLDGIEYADSFTFNPSKWMMVH
FDCTGFWVKDKYKLHQTFSVNPVYLRHANSGAAIDFMHWQIPLSRRFRSLKLWFVLRSFG
VKKLQAHVRQGTETAKFFESLVKSDPLFEIPAKRHLGLVVFRLKGPNWLTEKLLKELSSS
GRLFLIPATIHDKFIIRFTVTSQFTTREDILQDWSIIQHTAAQIISQNYGFHYINSGIPT
TVVQPTSDAISNVPQLYLEGGKYKTPSRKTVVQPKKLSVSPSHQQVKGQEDPLDDCFPED
VQNVTKHKLTSFLFSYLSVQGKKKTGRSLSCTSVPMTGNLEQCNPKAAATDKKESRANAR
VLSRLPEDMMMFKKGAFKKLIKFYSVPSFPECSIQCGLQLPCCPLQAIV
>E_A0A319APA8
MDLRDIKKTSQEQGQLHQKLWELNQTYTPGNVLPSASDLSRARASLPKALPAEGAGFESA
TQHILNDLVPAFNRSSISPNYYGFVTGGVTPVALFADNLVTAYDQNVQVHLADHSISTDV
EATALGLLADLLNLDRHRWSNGTFTTGATASNVHGLACGREFVLRAAAKKRGISIESVGE
YGLFELIQATGLSGIQILSTMPHSSLAKAAGILGIGRANVKSICRDDHYLRVDLVRLEAE
LKTNKASIVAVSCGEVNTGHFATSSLAEMESLRSLCDKYGAWLHVDGAFGIFGRILKDAE
FSAISKGCEGMELADSIAGDAHKFLNVPYDCGFVLCRHPSEAEKVFQNANAAYLSGGQSG
AASIPSPLNIGMENSRRFRALPVYASLVAYGQVGYQNMLQKQVRLSRMIAGWIYDHPKYN
ALPELANKDELLNQTYIIVLFSAKDDELNNSLTKEINATSKLFVSGTSWQGRPACRIAIS
NWRVEEERDFALVTDVLDNVAGKL
>E_A0A016U7N4
MTRCFQVSQAIFHKVGDRYSGVKEYVSLGDPLEVPLWVALPICILFVFLVSVSIFFCFKK
PKEKHFEEKKGAAKSAKIVKESREVEDSAPKESETDEGNQSASRKGTDSRKAETGKESTG
SLGDSEPVKKDEKEKKAEAASEKNEPYSPCVLRSRSQQEMIAEPVTSKEFEVYLHQLARF
AVDYYDNPCIYNVTPEVTPGFLYNNLPKTGPVHPESFDAIFNDIKTKIMPGLTHWQHPNF
FGYYPIGRCFPDMLADFITSALAVIGFSWDSCPALTEMEHAMINWVGRSFGLPENFLFQD
SPDSSQGGGTLTESGSDAIFCAVLAARQWKINQIVEEQQRTRGMKYDTIHDIGKRLVVYS
SKDAHSCIEKACKLAMVRYRPIQPTEENQWGITGEQIEEQIKKDLNRDLIPCFINCTLGT
TSTASCDKLTSICPVAKQYGTWLHVDAAYAGSTFIDPKYREVAEGIENAHTINVNLSKFL
LHSATLSIIWTREQKIYKDAFAITPIYLKPSAHGSASDQRDWGLHLSRRFKALKVWFILR
LCGVEGLRRYVARMCEMASYFESLVDQHPNLQIFTPRNFGLFTFQYIEPNFTKDEKNIHT
LRLLRFFNESHKIFLTHAQVAGNHVIRVSMSYERTTKETIDSAFNVMKTITEDYKKRKGD
PTLLKGPPEAPVESDSLVTDAEFFVQAKTPAKSLQSKREPGKSAPKDNDSVPSTAPKPTV
PQAPIPMQPERSSNTTAQQKLPSIAGNVGVASGRSAVGTSHSSQSNPTINSQVQPQPHK

>E_A0A251V0E5
MNPLDPVEFRRNGHMVIDFLADYYQNIQNYPVKSQAKPGFLLESLPDCAPLHPESIETII
NDVQRDIIPGITHWQSPNFFAYFASSGSTASFLGEMLINGFNVVGFNWESSPAATELEMI
VMEWLLKLLQLPNFDEFIKFFSFSNGGGGVLHGTTCEAFICTLHAARESILDHIGRENAR
KLAVYCSDQTHFSFQKSAKIVGINPKNIRQVSTSRSSNFKLSPRLLDEMIKKDIESGLVP
VYLCVTVGTTSMAMVDPVGPLSDVSSAYNMWVHVDAAYAGTACICPEFRHFLNGVEGASS
FSFNAHKWMLTNQSCCCLWVKDKSALTKSLSTNSEYLKNQATESGQVVDYKDWQIVLSRR
FQALKLWMVLRSYGANGLREVIRKHVNLAKDFQMLVSADKRFEIIVPRYFSMVCFRVSPE
VIGQCYDTEHEANEFNQKLLQSVNATGCVYMTHSIVGGVYFIRFVVGATLTEDRHVIMAW
KLICDQATSMLCTPTPKCI
>E_A0A0V0W3Q4
MDAEEFRKWGKKMIDFVADYWINLPSRTPMSDVKPGYLRSLLPEEAPMDPDSWENIFSDI
ETVILQGTTHWHHPLFFAYYPTGNSYPAILGDILSAGIGCIGFTWNSSPACTELEMVMMD
WLSKLLKLPEYFLYSHSGPGAGMIQGTASECVLFSMLAAKNKTCKKYESENKQHHICEKD
LIAYCSDQNSFREDASLDDGTAHSSVERAAMLAHVQIRKVPSDENYRMTRVALQAVIEND
INAGFIPFFVCATLGTTNSCAFDCLTEIGLLCKEKEIWLHIDAAYAGSAFICPEYRHLLD
GIEYADTFNFNPHKALMINFDCSAMFKNVLEIENAYYVNPQYLKHEHQNMIPDFRNWQIP
LGRRFRSLKLWLTFRALGVGFLQENIRKMCRLAKEFADFVVKDERFELVAPVILGLVCFR
LKDTNEVNEKLYQLINNQRRIHVVSSVLKNVFVLRISISSALTETADIYFAWKVISASAT
KLLASY
>E_A0A1L7XGT1
MDSKQFKEAATSAIDEIVNYYDTIADRRVVSSVEPGYLKKLLPNGPPQDGESWADIQKDI
ETKIMPGLTHWQSPNFMAFFPASSSYPGMLGELYSAAFTAPAFNWICSPAVTELETVVLD
WLAKLLNLPDCYLSTSHGGGVIQGSASEAIVTAMVAARDKYLRETTAHLSGLELEDAIAH
KRSRIVALGSDSAHSSTQKAAQIAGVRYRSIPVSKETDFALTGAALEAVLKECKAQGLEP
FYLTTTLGTTATCAVDDFASVASTLAKYAPPNKPGEIWVHVDAAYAGAALVCPEFHHLTA
PFEHFHSFDMNMHKWLLTNFDASCLFVRKRKDLIDSLSIMPSYLRNEFSESGLVTDYRDW
QIPLGRRFRSLKIWFVLRTYGVNGLQAHIRKHIKLGETFAGLIRTRQDLFEITTKPAFAL
TVFNVVPKVADKKAQDRITKDVYELVNKRGEIYITSSIVGGVYVIRVVSANPLAEEEYLR
KAFKILVETAEEVRDGKPIEDTLNGTILDIKGVGEAAVVKEKENSAAK
>E_W6Y6T7
MEVQSQEPFNQLAVEIAKIHVQPPPEDVLPSGDTLSSARSKLQTHLPAKGVGLEESIRHL
RQDLVPAFNASSRSPNYYGFVTGGVNQAAALADNLVTAFDQNVQVHLPNETIATDVEDRA
LYLLCELLNFEPSQWPHRIFTTGATAANVLGLACGREYVIAEASAHRTDAENSVGEVGIV
EAMRRAGIDDIQILTTVPHSSLSKAASILGLGRASVKCLGRSDAPHKFDMQLLKKSLELP
GAASIVVVSASEVNTGVFATSGLEEMQELRKLCDMHGAWIHADGAFGLFGRILSSPAHSS
IIEACAGLELADSITGDGHKLLNVPYDCGFFLSRHRAMAERVFQNPNAAYLASGNGPDTI
MSPLNIGLENSRRFRALPVYASLVAYGRDGYRDMLGRQIQLARGIARYILQSSQYELLPQ
KEASHEDILSGIFIIVLFRAKNEELNKQLVDKIKATRKIYVSGTSWEGRPACRFAISNWM
TDVDRDLPVIKQVLQDIA
>E_A0A2U9BKG5
MGQCETETECVKKKMKKYRVDEEVYNFTILDEEDYQTRDGGGMLRQSFRQVFDVQVRAFG
ENLFWFTETAVIGRPLCSCCDQQTQRAEEQAAPGEEEEEELEARDSRNTAPSGPAEIGGR
THAPLLVEPLPTHEVKDVGQQTEGEDEDLKAPLELLMEFLRAAMDRDFWLARKLCQLILI
YEPDNPEASEFLPLIQKKLLEVELFPDGRTHGGLMATSEPRTTGGDQDPNSANLIPPSTT
NEYAWMHGCTRKLGMKICGFLQKNNSMEEKGRLAGHKNLLAGDNSDRDARFRHTETDFSN
LFARDLLPAKNGEEPTIQFLLEVVDILTNYVKKTFDRSTKVLDFHHPHQLLEGMEGFNLE
LSDQPESLEQILVDCRDTLKYGVRTGHPRFFNQLSSGLDIIGLAGEWLTSTANTNMFTYE
IAPVFVLMEQLTLKKMREMIGWPSGEGDGIFSPGGAISNMYSVMIARYKYFPEVKTKGMS
AAPRLVLFTSEHSHYSIKKAGAALGFGTENVILLSTDERGRVIPADLEAKIIDAKQKGYV
PLFVNATAGSTVYGAFDPINEIADICEKYNLWLHVDGAWGGGLLMSRKHRHKLSGVERAN
SVTWNPHKMMGVPLQCSAILVREKGILAGCNSMCAGYLFQPDKQYDVTYDTGDKAIQCGR
HVDIFKFWLMWKAKGTIGFEQHIDKCLDLSQYLYNKIKNREGYQMVFDGVPQHTNVCFWY
IPPSLRGMPDGDERREKLHMVAPKVKAMMMESGTTMVGYQPQGNNVNFFRMVVSNPAVTQ
SDIDFLIDEIERLGQDL
>E_A0A091FVI7
GKEMVDYICQYLSNVRERRVTPDVQPGYMRSQLPDSAPMDPDSWDNIFGDIEKIIMPGVV
HWQSPHMHAYFPALTSWPSLLGDMLADAINCLGFTWASSPACTELEMNVMDWLAKMLGLP
DKFLHHHPNSVGGGVLQSTVSESTLVALLAARKNKILEMKLSETDADESSLNSRLIAYAS
DQAHSSVEKAGLISLVKMKFLPVDENFSLRGETLKKAIAEDRKKGLVPVFVCATLGTTGV
CAFDNLSELGPVCDAEGLWLHIDAAYAGTAFVCPEFRLFLDGIEYADSFTFNPSKWMMVH
FDCTGFWVKDKYKLHQTFSVNPVYLRHPNSGAAVDFMHWQIPLSRRFRSLKLWFVIRSFG
VKKLQAHVRHGTEIAKFFESLVKSDPLFEIPAKRHLGLVVFRLKGPNSLTEKLLKELSRS
GRLFLIPATIHDKFIIRFTVTSQFTTREDILQDWNIIQHTAAQIVSQNYGLHCTNSGDEA
RIPNMIVKPGSDAISSASQLYLDGGKHKTPSRKIVVQHKKLAACPSTCVISQQVKGQGDP
LDDCFPEDVPDVTKHKLNSFLFSYLSVQGKKKTARSLSCNSVPMTGGLEQCNPKAAATDK
KESHANARILSRLPEEVMKLKKSAFKKLIKFYSVPSFSECSIQCGLQLPCCPLQAIV
>E_A0A1L9N0S7
MDLRDIKGTSQEQGQLHQKLWELNQTYTPGNVLPSASDLSRARASLPEFLPAEGTGFESA
TQHILNDLVPAFNRSSISPNYYGFVTGGVTPAALFADNLVTAYDQNVQVHLAEHSISTDV
EATALGLLSDLLKLDRRSWNNGTFTTGATASNVHGLACGREFVLRAAAKKSGIDIESVGE
YGLFEVIQATGLSGIQILSTMPHSSLAKAAGILGIGRANVKSICRDNHYLQIDFGRLEAE
LKANKASIVAVSCGEVNTGHFATSSLAEMKSLRSLCDKYGAWLHVDGAFGIFGRILNGAE
FSAISKGCEGMELADSIAGDAHKFLNVPYDCGFVLCRQPGEAEKVFQNANAAYLSGGQSG
ALSIPSPLNIGMENSRRFRALPVYASLLAYGQVGYQEMLQKQVRLSRMIAGWIFDHPKYN
ALPELASKDELLDQTYIIVLFSAKDDELNGNLTKEINATYKLFVSGTSWQGRPACRIAIS
NWRVEEERDFALVTDVLDNVAGRV

```
>E_A0A218ZBM9
MAISLSLSYAKLQSIIGSKSNDVLPSLSALGHAEKSLPSSLPEAGLGEPATEAHLFSEIT
LGLSGQKTSSNYYSFVTGGVLPIAEIADNIVTAFDNSCQVHLPDQSISTTVEDRALSMLT
ELLNLGDGWNGRTFTTGATGANVLGLACGREAVVKARQKRAGESGGVGELGLLGACMAAG
IKEIQVLTAMGHSSLYKAASIVGLGRASIKDIALHKGQPWKLDVDALENKLKASDEGVAS
IVVLSMGEVNTGRFATDGLATMARIRQICNEWGAWLHVDGAFGAFARSLPPTAEFSSLIN
SAAGIELADSIAGDGHKMLNVPYDCGFFFTRSSDTLPAIFHNPNAAYLSGGSSAIASPLN
IGLENSRRFRALPVYAVLLAYGREGFAEMFARQVRLARGIAEFLSGSEDFELLPSQPSSD
GNYADVHIIVIFRARDEAVNAELVKRVNGTNKIYVSGTKWDGKPACRIAVSTWRVDVERD
LDLVKEVLRAASKPNV
>E_A0A218Z6N7
MDSKQFKEAATLAIGKIVNYYETIEDRRVVSNVEPGYLQKLLPDGLPQDGEPWGDIQKDI
ESKIMPGLTHWQSPNFMAFFPASSSFPGMLGELYSAAFTAPAFNWICSPAVTELETIVLD
WLAKLLNLPDCYLSTSHGGGVIQGSASEAVVTSVVAARDKYLRETTAHLSGLELEDAIAY
KRSRIVALGSEAAHSSTQKAAQIAGVRYRSIPVLKENNFALTGPELEGMLAECKAQGLEP
FYLTTTLGTTATCAVDDFESIAATLRKYAPPDIPGEIWVHVDAAYAGAALVCPEYQHLTA
AFEHFHSFDMNMHKWLLTNFDASCLFVRKRKDLIDALSIMPSYLRNEFSESGLVTDYRDW
QIPLGRRFRSLKIWFVMRTYGVNGLQAHVRKHIKFGETFASLLETRKDLFEIVAGPNFAL
TVFAILPKIQGKKEQDSITKEVYELINKRGEIYITSSVVASKYVIRVVSANPMAEERFLR
KAFDILVDTAEEVRDGKPSKRSINGVVSGRGEGAGEIAALDGPSR
>E_A0A091JKB2
GKEMVDYICQYLSNVRERRVTPDVQPGYMRAQLPDSAPMDPDSWDNIFGDIEKIIMPGVV
HWQSPHMHAYFPALTSWPSLLGDMLADAINCLGFTWASSPACTELEMNVMDWLAKMLGLP
DKFLHHHPDSVGGGVLQSTVSESTLVALLAARKNKILEMKLCEPDTDESSLNSRLIAYAS
DQAHSSVEKAGLISLVKMKFLPVDENFSLRGETLKKAIAEDRKKGLVPVFVCATLGTTGV
CAFDNLSELGPVCDAEGLWLHIDAAYAGTAFVCPEFRFFLDGIEYADSFTFNPSKWMMVH
FDCTGFWVKDKYKLHQTFSVNPVYLRHPNSGAAVDFMHWQIPLSRRFRSLKLWFVIRSFG
VKKLQAHVRHGTETAKFFESLVKSDPLFEIPAKRHLGLVVFRLKGPNWLTEKLLKELSSS
GRLFLIPATIHDKFIIRFTVTSQFTTREDILEDWSIIQHTAAQIVSQNYGLHCINSGDGA
RIPNMIVKPSSDAIGSVPQLYLDGGKYKTPSRKIVVQPKKLAASPSTCVISQQVKGQGDP
LDDCFPEDGPNVTKHKLTSFLFSYLSVQGKKKTAHSLSCNSVPTTGGLEQCNPKAAATDK
ESHANARILSRLPEEVMMLKKSAFKKLIKFYSVPSFPERSIQCGLQLPCCPLQAIV
>E_A0A0C3I054
MDSKQFKAAATSAIDEIVNYYDTIENRPVLSSVEPGYLKRILPDGPPQDGEPWADIQKDI
ETKIMPGLTHWQSPNFMAFFPASSTFPGMLGELYSAAFTAPAFNWICSPAVTELETVVLD
WLAKLFNLPECYLSTTHGGGVIQGSASEAVVTVMVAARDKYLRETTAHLSGLELENALAH
KRSTMVALGSEAAHSSTQKAAQIAGVRYRSVPVSKENNFSMTGADLEKVLEQCKAQGLEP
FFLTATLGSTSTCAVDDFGSMVPTLAKHAPLNQPGEIWVHVDAAYAGAALICPEFQHLTT
SFEHFHSFNTNMHKWLLTNFDASCLYVKKRKDLIDALSILPSYLRNEFSESGLVTDYRDW
QIPLGRRFRSLKIWFVLRTYGVNGLQAHIRKHVHLGEIFAGLIKTRGDLFEIVTGPTFAL
TVFNIVPKVANKAEQDRLTKEVYELVNKRGEIYITSSVVSGVYVIRVVSANPLAEEKNVR
RAFEILVETAEELRDGKLSSMTTISGNGVLNGSKDAN
>E_A0A151WUY7
FVQKEISFSLDNEPTSDEQIEEVIRQIIRRSMRTSSPYFHNEFFAGFDEYGLIDSCLTEV
LNTNISQSFFSLLEILMKICPVFTLMEKEVIQVSLKLMGYPIRGEGIMTPSSNISIMYAM
MLAKKNVLDQVKTKELYGNVPLVKTDKFSRMDMKDLTKVLKKVKMQQKIPFFMNATAGTP
VFGAIDPLRKIYDNCLKWKMWLHVDVRIGALMFSKCFQHKLRSIERYLFFILIDIVRARW
NPHNMLGISIPISVFFNLVVIIVLFQHLIIKCPKVDAIKFWLMWKARGTSGKVMSMSLDD
NPASTREIETVIRQTIQFSVKTSNPHFHNQLYAGVDEYGLIGSWLTDVLNTSQYTYEVAP
VFTLMEREVIQKSLELVGYPLMPEADGIMCPGGSISNMYGMLLARHKILPCIKKSGLFSM
ESPLACFTSEDSHYSILKSANWLGLGTDQVYKVKTDEFGRMKVSDLKRLLIKARNDGKQP
FFVNATAGTTVLGAIDPLPEIAAVCRSESLWLHVDACLGGTLLFSEKYRYRSDSVSWNLH
KMLGAPLQCSLFLVKGKNTLYEVNCAQAKYLFQQDKFYDVSWDTGDKSVQCGRKVDAMKF
WLMWKARGKIGLTRSVEQAMSCAEYFLKRIKETAGFRLVQPHYQCCNICFWYIPPTMRNE
NETPNWWEKLYCVTVEIKKRLILEGSLMISYMPLPQKEIGNFFRMVVNCQPPPTKSSMDY
VINQIEKVAVDL
>E_F5HAE7
MASTPLPDHMQPTYVDEILKPLESRISELLDEFCPANQGTERLNDVKKLVVNYTVLQILT
QTFDPNLCVTPRPKDLAQAEAWVEHYCRDSFIYPGDEPPKNSGFAGDGKEFYSTLITHIL
ADIVPALNSQALSSRYYGFVTGGVHPVAQAADNVVTALDQNVQVHMPSTHSISTVVEHHA
LSMLRSLLDLDGFHGKTFTTGATASNIMGLACGREAVISARLPEYAKKSGGVGELGLLAA
CMAAGVKEVQVLTSKGHSSLYKAASVVGLGRAAVKDLGFADAPWVLDLKAVERELQREDV
ANIIVVSAGEVNTGQFGTTGETMKELRQLADRYKAWIHIDGAFGIFARALPKTERFAKLF
ERTAGLELADSIAADGHKLLNVPYDNGIFFCSSPEVMSHVFQNPNAAYLAPVASTSSSGT
TTDIQSPLHVGLENSRRFRALPVYALLVHLGRGNMGEMLARMVDLARRIAAFIRGSDKYD
LLPDETADIDCTHVIVLFKAKNPDLNEVLVEKINETRRMYVSPTKWNGENAVRIAVGSWR
VDVEEDLAAVKLVLNSL
>E_A0A319ETC9
MELKDLKGTAQEQGRLHQELWELTQTYKPGGVLPSASDMSRARASLPETLTDEGAGFENT
TQHILNDLVPAFNRSSINPNYYGFITGGITPAALFADNLVSAYDQNVQVHLAEHSISTDV
EATALGLLADLLRLDRQHWSNGTFTTGATASNIHGLACGREYVLRAAAKKKGIAVESVGE
YGLFELIQATGLSGVQVLTTLPHSSLTKAAGILGIGRANVKSMCRDDHYLQFDLERLEAE
LARSDKASIVAISCGEVNTGHFATSSLAEMENLRRLCDKYGAWLHADGAFGIFGRVLEGA
EFSTISKGCEGMELVDSIAGDAHKLLNVPYDCGFFLCRHSGEAEKVFQNANAAYLTGGQS
GAPSIPSPLNIGMENSRRFRALPAYASLVAYGRTGYREMLQKQIRLSRMIAGWIFDHPKY
NALPEVASKDALLDQTYIIVLLSAKDDELNNNLTKRINATSRMWVSGSSWQGRPACRIAI
SNWRVEEERDFALVTEVLDSVAGGRS
```

>E_M2TJG6
MEVQSQEIFNQLAVEIAKIHVQPPPEDVLPSGDTLSSARSKLQTHLPTKGVGLEESIRHL
RQDLVPAFNASSRSPNYYGFVTGGTNQAAVLADNLVTAFDQNVQVHLPNETIATDVEDRA
LSLLCELLNFEPSQWPHRIFTTGATAANVLGLACGREYVIAEASAHRTDAENSVGEVGIV
EAMRRAGIDDIQILTTVPHSSLSKAASILGLGRASVKCLGRSDAPHKFDMQLLKKSLELP
GAASIVVVSASEVNTGVFATSGLEEMQEIRKLCDMHGAWIHADGAFGLFGRILSSPAHSS
IIEACTGLELADSITGDGHKLLNVPYDCGFFLSRHRAMAERVFQNPNAAYLVSGNGPDTI
MSPLNIGLENSRRFRALPVYASLVAYGRDGYRDMLERQIQLARGIAQYIQESSQYELLPQ
KEASHEDILSGIFIIVLFRAKDEEVNKQLVDKIKATRKIYVSGTSWEGRPACRFAISNWM
TDMNRDLPVIKKVLQDIA
>E_A0A0L7RAZ6
MPANEEILSAPLQQSLDNFLGKACVNGCVSSNDDEDDCQKCCLGSNFKQNGKEECQYKSF
PVREVHEKFMRSFIDLLLEDAVFKGTARKNRVVEWMEPSALHSAIDLNLQDQGVSHEELF
SLARNVIKYSVKTGHPRFVNQLYSSVDPYGLLGQWLTDALNPSVYTYEVSPVFSLMEEEI
LREMRKIVGWKDGRGEGIFCPGGSIANGYAINLARHYRFPQLKELGMTSAGRLIVFTSQD
AHYSVKKLSAFLGIGTSNVCEVKTDNKGKMCMVDLEAQIKRVLEDGATPLMVSATAGTTV
LGAYDPLRDTAAICKKYNMWFHVDAAWGGGALMSKKYRHLLDGAELADSITWNPHKLLAA
PQQCSTLLLRHEGLLQAAHGLKASYLFQPDKFYDTSFDSGDKHMQCGRKADVLKFWFMWK
AKGTRGLEKHVDRVFELSRYFTDYIRHREGFKLMLEPECTNVCFWYVPPSKRHLQGEELS
KVFQKIGPAVKERMVKKGSMLITYQPQRELPNFFRLVLQNSGLTEADMRFFAEEIERLAS
DL
>E_A0A0L0HIP1
MPAAKLPDPCNSLGGLTPKSVLPAGAAADPRPLTPESEDIDEGYDEDFPDDFPFWYGGAS
GVQRIGATRKASVDAAPCPENEHEGREEEHDGGEYEHGEPRERRSSYHDHDSVLHYFVAT
SQSENKLDRYLSNIIEAFLNEPAPIYATQPDTPESMRARFVQSEVPLRSKAAGQIGVASA
EEYLRTVKSNVIDRATRVSSPKMIGHMTTALPYFHRPLARLLAAMNQNVVKLETASTMTY
LERQTVAMLHREFYNNPSSFYEAYMHSPEHSLGVFCSGGTIANITAMWAARNRALPSDPQ
NGFKGVEKEGLFKALKHYGYEGAVIVGSRLMHYSFKKAADLLGLGDEGLVTIDSDDAFRM
RMDELEAKLSELQAKKYLIIAVVGIAGTTETGSIDPLDRIALQCQRQGIHFHVDAAWGGP
LIFSAEHRPKLHGIHHADTITVDGHKQLYTPMGLGLLLFQHPTTAHAIRKTANYVIRHDS
PDLGKFTLEGSRPAISLHLHASLHLLGRDGLECLVTRSATLARQMSVRIDTHPSYAFQGL
HLPETNIFLFRYLPKELRVKVHTGEPLTEEEDASVSDCTRRVQVALASAGTEPSEATQEP
SPPTGFVSRTRVRFHGRDVDAFRVVIANPLTSWDDVEDVLRNMLIIGAKVEAEIANERRP
LVAQHVPTVQKVDGNVKEDISRWSELRVPQKGEKMWVGWPFDM
>E_R0IYN0
MQAPNQETFDQLVAEIAKIHFQPPPQHVLPSGDALKDARSKLQPHLPAQGVGLEESIRHL
REDLAPSFNASSRSPNYYGFVTGGANQAATLADNMVTAYDQNVQVHLPNETIATDVEDRA
LSLLCELLNLEPAQWPHRIFTTGATTANVLGLACGREYVVAEASAHRTDVENSVGEVGIV
EAMRRAGIDDIQVLTTVPHSSLSKAAGILGLGRASVKCLGRSEAPHIFDMQLLKKSLEQP
GAASIVAISASEVNTGVFATSNLEEMQELRKLCDMYGAWIHVDGAFGLFGRILSSPAYSS
IIQACAGLELADSITGDGHKLLNVPYDCGFFFSRHRNMAERVFQNPNAAYLASSNGPDSI
MSPLNIGLENSRRFRALPVYASLVAYGREGYQDMLERQIQLSRGIAQYILESNKYELLPQ
KDAPQEEIFSGIFIIVLFRAKDEELNKQLVDKIKATRKIYVSGTSWEGRAACRFAVSNWM
AQVERDLPIIKQVLEELV
>E_B4HXA1
MLASENFPTHHFKESIFKPYSTTSGDDLASVTPLTATAALVASTPSPADSTSAVAFEQAS
KMLATAANNNNNNNNNNITSTKDDLSSFVASHPAAEFEGFIRACVDEIIKLAVFQGTNRS
SKVVEWHEPAELRQLFDFQLREQGESQDKLRELLRETIRFSVKTGHPYFINQLYSGVDPY
ALVGQWLTDALNPSVYTYEVAPLFTLMEEQVLAEMRRIVGFPNGGQGDGIFCPGGSIANG
YAISCARYRHSPESKKNGLFNAKPLIIFTSEDAHYSVEKLAMFMGFGSEHVRKIATNEVG
KMRLSDLEEQVKQCLENGWQPLMVSATAGTTVLGAFDDLAGISELCKKYNMWMHVDAAWG
GGALMSKKYRHLLNGIERADSVTWNPHKLLAASQQCSTFLTRHQQVLAQCHSTNATYLFQ
KDKFYDTSFDTGDKHIQCGRRADVFKFWFMWKAKGTQGLEAHVEKVFRMAEFFTAKVRER
PGFELVLESPECTNISFWYVPPGLREMERNREFYDRLHKVAPKVKEGMIKKGSMMITYQP
LRQLPNFFRLVLQNSCLEESDMVYFLDEIESLAQNL
>E_A0A091F3Y5
GKKMVDYICQYLSSVRERRVTPDVQPGYMRAQLPDSAPMDPDSWDNIFGDIEKIIMPGVV
HWQSPHMHAYFPALTSWPSLLGDMLADAINCLGFTWASSPACTELEMNVMDWLAKMLGLP
DKFLHHHPDSVGGGVLQSTVSESTLVALLAARKNKILEMKVSEPDTDESSLNSRLIAYAS
DQAHSSVEKAGLISLVKIKFLPVDENFSLRGETLKKAIAEDRKKGLVPVFVCATLGTTGV
CAFDNLSELGPVCDAEGLWLHIDAAYAGTAFVCPEFRLFLDGIEYADSFTFNPSKWMMVH
FDCTGFWVKDKYKLHQTFSVNPVYLRHANSGAAIDFMHWQIPLSRRFRSLKLWFVLRSFG
VKKLQAHVRHGTETAKFFESLVKSDPLFEIPAKRHLGLVVFRLKGPNWLTEKLLKELSSS
GRLFLIPATIHDKFIIRFTVTSQFTTREDILQDWSIIQHTAAQIISQNYGLHCINSGDGT
GIPTTVVQPSSDAISNVPQLYLDGGKYKTPSRKTVVQPKKSSVSPSTRVISQQVKGQGHP
LDNCFPEDVQDVTKHKLTSFLFSYLSVQGKKKTARSLSCTSVPMTGNLEQCNPKAAAADK
ESRANARVLSRLPEDMMMFKKGAFKKLIKFYSVPSFPECSIQCGLQLPCCPLQAIV
>E_A0A093HSM0
GKEMVDYICQYLSSVRERRVIPDVQPGYLRAQLPDSAPMDPDSWDNIFGDIEKIIMPGVV
HWQSPHMHAYFPALTSWPSLLGDMLADAINCLGFTWASSPACTELEMNVMDWLAKMLGLP
DKFLHHHPNSVGGGVLQSTVSESTLVALLAARKNKILEMKVSEPDTDESSLNSRLIAYTS
DQAHSSVEKAGLISLVKMKFLPVDENFSLRGETLMKAIEEDRKRGLVPVFVCATLGTTGV
CAFDNLSELGPICDAEGLWLHIDAAYAGTAFVCPEFRLFLDGIEYADSFTFNPSKWMMVH
FDCTGFWVKDKYKLHQTFSVNPVYLRHPNSGAAVDFMHWQIPLSRRFRSLKLWFVIRSFG
VKKLQAHVRHGTETAKFFESLVKSDPLFEIPAKRHLGLVVFRLKGPNWLTEELLKELSSS
GRLFLIPATIHDKFIIRFTVTSQFTTREDILQDWTIIQHTAAQIVSRHYALHRINSDEGA
RIPNMIVKPSSNTSSKASQFYLDDGEHKTPSRKIVVQPKKLAVSPNTCVINQQVEGQGDP

LDDCFPEDVQDVTKHKLTSFLFSYLSVQGKKKTARSLSCNSVPMTGGLEQCNPKAGDTDK
KESHTNTRILSRLPEEVMMLKKSAFKKLIKFYSVPSFPECSIQCGLQLPCCPLQAIV
>E_A0A087QWQ6
GKEMVDYICQYLSNVRERRVTPDVQPGYMRAQLPDSAPMDPDSWDNIFGDIEKIIMPGVV
HWQSPHMHAYFPALTSWPSLLGDMLADAINCLGFTWASSPACTELEMNVMDWLAKMLGLP
DKFLHHHPDSVGGGVLQSTVSESTLVALLAARKNKILEMKLSEPDTDESSLNSRLIAYAS
DQAHSSVEKAGLISLVKMKFLPVDENFSLRGETLKKAVAEDRKKGLVPVFVCATLGTTGV
CAFDNLAELGPICDAEGLWLHIDAAYAGTAFVCPEFRLFLDGIEYADSFTFNPSKWMMVH
FDCTGFWVKDKYKLHQTFSVNPVYLRHPNSGAAVDFMHWQIPLSRRFRSLKLWFVIRSFG
VKKLQAHVRHGTETAKFFESLVKSDPLFEIPAKRHLGLVVFRLKGPNWLTEKLLKELSSS
GRLFLIPATIHDKFIIRFTVTSQFTTREDILQDWNIIQHTAAQIVSQNYGLHCINSGDGA
RIPNMIVKPRSDAISSASQLYLDGGKYKTPPRKIVVQPKKSAASPSTCVISQQMKGQGDP
LDDCFPEDVQGVTKHKLTSFLFSYLSVQGKKKTARSLSCNSVPVTGGLEQCNPKAAATDK
ESHANARILSRLPEEVMMLKKSAFKKLIKFYSVPSFPECSIQCGLQLPCCPLQAIV
>E_A0A1V4JEJ7
MEPEEYRRRGKEMVDYICQYLSNVRERRVTPDVQPGYMRAQLPDSAPMDPDSWDNIFGDI
EKIIMPGVVHWQSPHMHAYFPALTSWPSLLGDMLADAINCLGFTWASSPACTELEMNVMD
WLAKMLGLPDKFLHHHPDSVGGGVLQSTVSESTLVALLAARKNKILEMKISEPDTDESSL
NSRLIAYASDQAHSSVEKAGLIALVKMKFLPVDKNFSLRGETLKKAIAEDRKKGLVPVFV
CATLGTTGVCAFDNLSELGPVCDAEGLWLHIDAAYAGTAFVCPEFRLFLDGIEYADSFTF
NPSKWMMVHFDCTGFWVKDKYKLHQTFSVNPVYLRHPNSGAAVDFMHWQIPLSRRFRSLK
LWFVIRSFGVKKLQAHVRHGTETAKFFESLVRSDPLFEIPAKRHLGLVVFRLKGPNWLTE
KLLKELSSSGRLFLIPATVHDKFIIRFTVTSQFTTREDVLQDWNIIRHAAAQIVTQNYGL
RCINSGAGARIPNMIVKPSSDAISNASQLYLDEGKHKIPSRKTEVQPKKLAESPSMCVIS
QQVKGQGDPLDDCFPEDVQDVTKNKLTSFLFSYLSVQGKKKTARSLSCNSVPMTGSLEQC
NPKAAATDKKESRANAKILSRLPEEVMMFKKSAFKKLIKFYSVPSFPECSIQCGLQLPCC
PLQAIV
>E_A0A1J9PGB1
MSIQPLQTPEDNEFPIQIWQTAISPWTSPVIPAPNTLAQVRSSLITSLPTTGLGFSEVKR
HIVQDITPGFNGSSISANYYGFVTGGATPAALLADHIVSAYDQNVQVHIPDHSVVTDLED
RALTFLLDLFHLDHKCWTHKTLTTGATASNVLGLALGREFVLRGAVERKAGADPEGIRKS
VGEYGMAEVLLEAGLKGLQVLSTYPHSSLGKAAGILGIGRANVRSVCTAGGGESQRPLQF
DYQVLERELGRSDMASIVAVSSGEVNTGHFATQGLEEFRKLRQLCDKYGAWLHVDGAFGL
FGRILKSGGEFDSILKGCQGLELADSIAGDGHKLLNVPYDCGIFFSRHANLAEDVCRNPN
AAYLNAATGANGIPAPCNNGLENSRRFRALPVYATLVAYGKDGYRDMLARQIRLARSVAG
WLFEHPAYEVLPHNPEKEGLLQDTFIIVLFRAKDENVNRVLVNKINATSKIYVSGTNWAG
KPACRIAISNWRANEEQDFEMITSVLGAIAQ
>E_H3CR31
MATSEPRATDAEQEPNSDNLRAPSTTNEYAWMHGCTRKLGMKICGFLQKNNSLDEKSRLA
GQKNMLACNNSERDARFRRTETDFSNLFARDLLPAKNGEEPTIQFLLEMVDILTNYIKKT
FDRSTKVLDFHHPHQLLEGMEGFNLELSDQPESLEQILVDCRDTLKYGVRTGHPRFFNQL
SSGLDIIGLAGEWLTSTANTNMFTYEIAPVFVLMEQLTLKKMREMIGWPNGEGDGLFSPG
GAISNMYSVMIARYKYFPEVKTKGMSAAPRLVLFTSEHSHYSIKKAGAALGFGTENVILL
STDERGRVIPADLEAKIIDAKQKGYVPLFVNATAGSTVYGAFDPINEIADICEKYNLWLH
VDGAWGGGLLMSRKHRHKLSGVERANSVTWNPHKMMGVPLQCSAILVREKGILAGCNSMC
AGYLFQQDKQYDVTYDTGDKAIQCGRHVDIFKFWLMWKAKGTIGFEQHIDKCLDLSQYLY
NKIKNREGYEMVFDGVPQHTNVCFWYIPPSLRGMPDGDERREKLHRVAPKIKAMMMESGT
TMVGYQPQANKVNFFRMVVSNPAVTQSDIDFLIDEIERLGHDL
>E_G1N6G5
SMEPEEYRRRGKEMVDYICQYLSNVRERRVTPDVQPGYMRAQLPDSAPMDPDSWDNIFGD
IEKIIMPGVVHWQSPHMHAYFPALTSWPSLLGDMLADAINCLGFTWASSPACTELEMNVM
DWLAKMLGLPDKFLHHHPDSVGGGVLQSTVSESTLVALLAARKNKILEMKLSEPDADESS
LNSRLIAYASDQAHSSVEKAGLISLVKMKFLPVDENFSLRGETLKKAIAEDRKKGLVPIF
VCATLGTTGVCAFDSLSELGPICGAEGLWLHIDAAYAGTAFLCPEFRLFLDGIEYADSFT
FNPSKWMMVHFDCTGFWVKDKYKLHQTFSVNPVYLRHPNSGAAVDFMHWQIPLSRRFRSL
KLWFVIRSFGVKKLQAHVRHGTETAKFFESLVRSDPLFEIPAKRHLGLVVFRLKGPNCLT
EKLLRELSSSGRLFLIPATIHDKFIIRFTVTSQFTTREDILQDWNIIQHTAAQIISQHNE
LHHISSGDEAKIPNTISESSSSAISHASQLYIEEGKYKIPSRKIVGQRKKLEMSPSTCVI
SQQVEGQADLDDCFPEDVQDVTKHKLTSFLFSYLSVQGKKKTARSLSCNSVPMTDLLEQC
NPKATDHDKKEPHANARILSRLPEEVMMLKKSAFKKLIKFYSVPSFPECSIQCGLQLPCC
PLQAIV
>E_A0A2T3AUC5
MDSKQFKEAATSAIDEIVNYYDTIENRRVVSNVEPGYLRKILPSGPPEEGEAWADIQADI
ETKIMPGLTHWQSPNFMAFFPASSSFPGMLGELYSAAFTAPAFNWICSPAVTELETIVLD
WLAKLLNLPECYLSTTHGGGVIQGSASEAIVTVMVAARDKYLRETTSHLSGLELEDSIAH
KRSKLVALGSEAAHSATQKAAQIAGVRFRSIPVSKESNFALTGEALEETLKQCKAEGLEP
FFLTTTLGTTATCAVDDFAAVASVLAKHAPPNVPGEIWVHVDAAYAGAALVCPEYQHLTA
SFEHFHSFDMNMHKWLLTNFDASCLFVRRRKDLIDALSVMPSYLRNEFSDSGLVTDYRDW
QIPLGRRFRSLKIWFVLRTYGVKGLQAHIRKHIKLGEVFADLIKTRPDLFTILSGPAFAL
TVFTIVPKTAGKEAQDQISKEVYELVNKRGEIFITSSVIAGIYALRVVSSNPLAEEQFLR
KAFRILVETTEEVRDGKVITKENSI
>E_A0A138ZZL4
MTNTTNANGAPTAPSQSPSGAIANPFPFPTTNFAPGGPQDIETFRKNAKEMVDFMCDYYK
TVGDHSTLSRVQPGYLAPLLPKKAPEDPESFESILADVQQHIIPGVTHWQSPNFFAFFPS
NNSFPSLLGDMLSGMINCIGFNWMTSPACTELEMIAMDWLAKLVGLPDAYLNTSGVGGGV
IQMSASDASVVAMIAARQRKLRQVKTEMEAQGKGDEFNEQDVIKRMVFYGSDEAHSCHKK
GAMVLKLNFKTVAADDSYAVKGATLKKAIETDVAAGLLPTFFVGTIGSTSTGATDHLSEI

GPVCRDTGVWLHVDAAWAGSAFVCPEHRELMRGMVDETTGEVYVDSYDFNPHKWLLVNFD
CSAMWLKERADLVDALSIMPAYLRNKASASGSVIDYRDWQVPLGRRFRSLKLWFVMRSFG
AKGLREHITKCVNLTRYFETLVTADDRFELIVPRTLALACFRIKPSVLSAHPGLDVNAVN
KQIADGVNESSNFLITASEVKGKPEVADPATGKVAMVYFLRVTIGGWSEEENVRGVWDVI
RGQADKVLAELR
>E_A0A093GDE1
GKEMVDYICQYLSNVRERRVTPDVQPGYMRAQLPDSAPVDPESWDNIFGDIEKIIMPGVV
HWQSPHMHAYFPALTSWPSLLGDMLADAINCLGFTWASSPACTELEMNVMDWLAKMLGLP
DKFLHHHPDSVGGGVLQSTVSESTLVALLAARKNKILEMKLSEPDTDESSLNSRLIAYAS
DQAHSSVEKAGLISLVKMKFLPVDENFSLRGETLKKAIAEDRQKGLVPVFVCATLGTTGV
CAFDNLSELGPICDAEGLWLHIDAAYAGTAFVCPEFRLFLDGIEYADSFTFNPSKWMMVH
FDCTGFWVKDKHKLHQTFSVNPVYLRHPNSGDAVDFMHWQIPLSRRFRSLKLWFVIRSFG
VKKLQAHVRHGTETAKFFESLVKSDPLFEVPAKRHLGLVVFRLKGPNWLTEKLLKELSSS
GRLFLIPATIHDKFIIRFTVTSQFTTREDILQDWNIIQQTAAQIIGQNYGLHCINSGDGA
RIPNMVLESSSDAISNASQLYLDGGKYKTPPRKIVVQPKKSAVSPSKCVISQQVKGQGDP
LDDCFPEDVQDVTKHKLTSFLFSYLSVQGKKKTARSLSCNSVPMTGSLDQCTPKVTAADR
ESHANARILSRLPEEVMMFKKSAFKKLIKFYSVPSFPECSIQCGLQLPCCPLQAIV
>E_A0A0D2EPM7
MDQRKSLQDVVSQLQKILSTPGLPTSSTVPDKASSRPALILPQDTASLDDLQSKADSSSP
LANHTGDGSDSIAQLASHLRAAILPYLNLASLSPNYYGFVTGGATPAALLGDFLASIYDQ
NVHVHLPNETISTTLEVVTLNLLTQFFRLPKDWSLGPSSSGGGVFTTGATASNILGLALG
REYVLRQALLRKGIPGSADASCGEYGLAELLLRVGASKIQVLSTLPHSSIAKAASVLGIG
RRNVVSIATAHSSSDGDDDSLRIDLERLRTEAQRTDVLNILAVSVGEVNTGRFATDSLDA
MRQLREICDEYGLWMHVDGAFGLFGRVLPVEDADFEELVNGVEGLELADSITGDCHKLLN
VPYDCGVFFTRHKNLSEDVFGNPGAAYLKAAAGDGIQSPLNIGLENSRRFRALPVYCTLM
AYGRDGYVEMLKRQIGLARRVTQWLMRDGRFEVLPRGGGQKEILAKTYIVVLFRLRNDEK
NRDFVRKVNATGRIYMSGTVWEGKPAGRIAVSNWQADVERDGHLIESVLDEVAGH
>E_U3K9Z6
SMEPEEYRQRGKEMVDYICQYLSNVRERRVTPDVQPGYMRAQLPDSAPMDPDSWDNIFGD
IEKIIMPGVVHWQSPHMHAYFPALTSWPSLLGDMLADAINCLGFTWASSPACTELEMNVM
DWLAKMLGLPDKFLHHHPDSVGGGVLQSTVSESTLVALLAARKNKILEMKVSEPDADESS
LNSRLIAYASDQAHSSVEKAGLISLVKIKFLPVDENFSLRGETLKKAIAEDRKKGLVPVF
VCATLGTTGVCAFDNLSELGPVCDAEGLWLHIDAAYAGTAFVCPEFRLFLDGIEYADSFT
FNPSKWMMVHFDCTGFWVKDKFKLHQTFSVNPVYLRHANSGAAIDFMHWQIPLSRRFRSL
KLWFVLRSFGVKKLQAHVRQGTETAKFFESLVKSDPLFEIPAKRHLGLVVFRLKGPNWLT
EKLLKELSSSGRLFLIPATIRDKFIIRFTVTSQFTTREDILQDWSIIQHTAAQIIRQNYG
LHYISSGDGAGIPTTIVQPTSDAISSVPQLYLDGGKYKTPSRKTVAQPKKLSVSPSTCVI
SQQVKGQEDPLDDCFPEDAQDVTKHKLTSFLFSYLSVQGKKKTARSLSCTSVPMTGNLEQ
CNPKAAATDKKESRANARVLSRLPEDMMIFKKGAFKKLIKFYSVPSFPECSIQCGLQLPC
CPLQAIV
>E_A0A2B7X0N5
MSIQPLQSREDKEFPSQVWQTAISPWTSSPLPTPTTLSHVRSSLISALPSAGLGFSETKR
HVINDITPGFNGNSLAANYYGFVTGGVTPAALLADNIVSAYDQNVQVHLPDHSVATDVED
RALSFLLDLFDLDHKKWTHKTLTTGATGSNVLGLALGREFILRAAVERKYGAAREVVKSV
GEHGMAEVLVAAGLKGLQVLSTYPHSSIGKAAGILGIGRANVKSIGAAGASGVSLKFDFD
ILEKELARSDMASIVAVSSGEVNTGHFATEGFEELREIRQLCDKYGAWLHIDGAFGMFGR
ILKSGGEFDRVWKACQGLELADSITGDGHKLLNVPYDCGFFFSRHADLAEDVCRNPNAAY
LSAGGGGIPAPCNNGIENSRRLRALPVYATLVAYGKDGYRDMLERQIRLARSVVGWLFEH
PAYTVLPYNPVKESLLQDTFIIVLFRAKDEDLNRVLVSKINAASTMYVSGTIWDGKPACR
IAISNWRVNEEKDFEVITSVLREIAQ
>E_A0A370TBI7
MDSKQFKDAATSAIDEIINYYDTIQDRRVVSNVEPGYLKKLLPDGPPEEGESWADIQKDI
ETKIMPGLTHWQSPNFMAFFPASSSFPGMLGELYSAAFTAPAFNWICSPAVTELETVVLD
WLAKLLNLPDCYLSTSHGGGVIQGSASEAIVTVMVAARDKYLRETTSHLSGLELEDAIAH
TRSKLVALGSEMAHSSTQKAAQIAGVRFRSVPTTMDDEFAMTGAGLEEVLKQCKADGLQP
FYLTTTLGTTATCAVDDFGSIASTLSKHAPPDAPGEIWVHVDAAYAGAALVCPEYQHLTT
SFEHFHSFDMNMHKWLLTNFDASCLYVKKRKDLIDALSIMPSYLRNEFSESGLVTDYRDW
QIPLGRRFRSLKIWFVLRTYGINGLQAHIRNHIKLGELFAGLLKSRQDLFEILTGPSFAL
TVFRAVLPAGSKAEQNELTKEVYELVNRRGEIYLTSGVVAGIYAIRVVSANPKAEEKYLR
KAFDILVATTEEIRDSKSSNGSVNGAVVHGKGEGVGEEVAQHSNGATK
>E_H3AXV6
MATSAPSSSGDSPDPNTTSLRPTTYDTWCGVAHGCTKKLGLKICGFLQRNNSLDDKSRIV
SSLKERQSATNVLPCENSERHTRFRRAETDFSNLFARDLLPAKNGEEQTMQFLLEVVDIL
LNYIKKTFDRSTKVLDFHHPHQLLEGMEGFNIELSDNPESLEQILVDCRDTLKYGVRTGH
PRFFNQLSSGLDIIGLAGEWLTSTANTNMFTYEIAPVFVLMEQITLKKMREIIGWPNGDG
DGIFSPGGAISNMYSVMAARYKYFPDVKSKGMAAVPKLVLFTSEHSHYSIKKAGAALGIG
TDNVILIKCNERGKIIPADLEAKILEAKQKGHVPIYVSATAGSTVFGAFDPVQEIADICE
KYNLWLHVDAAWGGGILMSRKHRHKLNGIERANSVTWNPHKMMGVLLQCSAILVREKGIL
QGCNQMCAGYLFQQDKQYDVSYDTGDKAIQCGRHVDIFKFWLMWKAKGTVGFENQINKCL
ELSEYLYTKIKRRDGFEMVFDGEPEHTNVCFWYVPPSLRGMPDCEERREKLHKVAPKIKA
QMMESGTTMVGYQPQGDKVNFFRMVVSNPASTRSDIDFLIEEIERLGQDL
>A_F2KP03
MFNVLSELEKFRAEDIPYSRVLSSMCTTPLPIALKAHELFIETNLGDPGIFAGTWKLEQK
LIKMLGELLHNPNAKGYICSGGTEANIQAIRAARNVIRRERKIDRPNIVVPESAHFSFEK
IGDILGVEVRRAKLDEEFKVDVASVESLVDENTVGIAGIAGTTELGQIDPIDELSKLALQ
LGVPLHVDAAFGGFVIPFMNKPYPFDFELEGVTSITIDPHKMGMATIPAGGILFRDEKFL
NALIVETPYLTSRYQYTLTGTRPGTGVASAYAVLKHLGYKGMKQIVDECMRMTALLVEEM

```
TSLGFEPVIEPVMNVVCFKTEKAEKIKEELYRRRWVISTIKNPRAIRLVVMPHVTEEVVK
GFISELKSVLRSV
>A_Q5V1B4
MLQRAEPQDFERVLSSMCTVPHPSAREAAERFLATNPGDPGTYETIAGLEREAVEYLGDI
TGLSDPAGYVASGGTEANLQAIRIARNRADTDDPNVVAPVHAHFSFTKAADVLGVELRTA
PAADYRVNMAAMAELVDEDTVCVVGVAGSTEYGYVDPIPAIADLAETVDALCHVDAAWGG
FYLPFTDHDWHFGHADIDTMTIDPHKVGQAAVPAGGLLARDRTLLDELAVETPYLESTDQ
LTLTGTRSGAGVASAVAAMESLWPAGYRQQYETSMANADWLADQLSARGHDVVGPELPLV
AADLSMPMTDELRDRGWRVSKTGAGEMRVVCMPHVTRSMLRSFVADLDWY
>A_A0A1H1C830
MSADELFLGSDGGDAAYREAMGRAVDAVLDSFADGDDPYSGASPDALAAQFDDPVVPEDG
RGLDAAIDEVSERVLAHSVGTSNPRCAAHLQCPPMIPGLVAEALLTATNQSLDSFDQAPA
ATVLEERVVGALCDLFGLPSGADGVFTSGGTQSNFQALLLARDRHCDRRFGRNVQADGLP
ADAESLRILCSEEAHFTGKQAAHHLGLGERAVVTVPTDDDRRMDPDALDSTLAELDDRDA
DPFALVGTAGTTDFGSVDPLAALADRAAERDLWFHVDAAYGGALAVSDDHGHLIDGIERA
DSVAVDFHKLFYQPISCGAFLLRDGDDFEWMARNAAYLNPEEHDEAGVPNLVAKSVQTTR
RFDALKPYVAFRALGRSGLATLVDRTLELADEAAELVAAADDFELLAEPTLNAVVFRYRP
REGMDDAAVSRLNAAVRSELLRDGRAVVARTDVGGVTSLKFTLLNPTATLDDVAAMLDAV
RDCGSDVAAERGVAA
>A_U6EEL6
MEDKGIPKEQVYQMLRKYKEKDLTHSSGRILGSMCTCPHPVGIKAYTMFLESNLGDPGLF
PGTKAMEDEVITMLGQLLGKEDVYGHIITGGTEANLMAMRAARNLKNVENPEIIVPKSAH
FSFKKAADMLCLDLKMADLDEDYRMDISSVESLISDNTVAIVGVAGTTELGKIDPIEDLS
RICQEQDIHLHVDAAFGGYIIPFLKESGYDLPEFDFRLPGVSSITIDPHKMGMAPIPTGG
ILFRERKHLEAMAIETPYLTEDLQSTVVGTRTGASTAATWALLKHLGREGYQEIATSCMD
VTHKLAEGIEEAGFELVTEPELNIVPFRSRNMSVEELAQQLEKRGWAVSLATYPRSIRII
VMPHLKIEHINDFLKDLKTITGA
>A_U6EDB6
MLSGKKDLKKLDKSQREHNTATTAYGTRYFEKSIPKYEIPLDGMPADAAYQLIHDELNLD
GNPVLNLASFVTTWMEPQADKLIMESVDKNYVDADEYPQTQKIEERVVNILARLFNSPDD
CHSIGTATIGSSEAIMLALLAHKWTWRENRKAEGKPTDKPNIVMGADVHTVWEKFAKYFD
VELKLIPIKDDLYTITAEDVAKEVDENTIAVGAVIGTTFTGQMDPIKEINDVLMDIKKEK
GWNIPIHVDGASGGFIAPFIHPDMEWDFRLEQVRSINVSGHKYGLVYPGVGWVVFKDKSD
VPEDLVFNINYLGGSMSNYSLNFSKASNTIIAQYYNLIRLGFNGYKAVIDNMIENTRYMA
DKLEETGRFEVLNPKILFPLVTVKLKDSDFTVFQLSDKLREHGWIVPAYTLPANADDIAV
LRMVIKENFGRDMVNAFIDDIKTCCKNLESEEEKIKKEDPSLLY
>A_D7EBV8
MNEKGISNQKLTDLLDTAKSNDVGYERVLSAMCTYPHDVAVQAHTKFIEANMGDPGLFPG
TYSLEKEVINMMGQLLHCSSVHGYITTGGTESNIQALRTMVNSSNVANPNVIVPESAHFS
FDKIANILGIEVKKAELDSKFKVDIGSVKSLIDSNTIGLVGIAGSTEFGQIDPINSLSDI
ALENNLYLHVDAAFGGFVIPFLETSYHFDFVLDGVTSIALDPHKMGFSTIPSGGILFRNR
EDLNHLQTHTPYLTISTQSSLTGTRSGASVAATYAVMSYLGKEGYRQIVKQCMDLTNDLV
EGAKKIGINPLIEPVMNVVTLDVQDPDTLRARLRDEFGWYVSITRNPRALRLVLMPHLTH
KNLDLFLQDLEKLVKIG
>A_D5VUB3
MGRVIEELKRFRELDIKYSEGRIFGSMCSSIHPLAKEIVSLFLETNLGDPGLFKGTKLLE
EKAVKLLGEILKNKEPYGFIVSGGTEGNLLAMRVVKKMKGRTIILPKTAHFSFEKAKEMM
DLNLVYAPLTKGYEIDVRFVKDYVEDYKVDGIVGIAGTTEFGTIDNIEKLSEIAKENDIY
LHVDAAFGGFVIPFLPKEYRRKEINYTFDFSLNVDSITIDPHKMLLCPIPAGGIIFKNSS
YKRYLEVDAPYLTETKQATILGTRPGFGAACTYGLLRYFGEEGLKKLVKEVMDRTFYFKE
RLEREGFKLLLEPILNIIAIEDENHIETCKKLKEMGYYPSVCFNAKALRIVVMPHIREEH
IDNFIEVLKEVKRD
>A_Q2FSD2
MDAEGLSTDELFCFLQAKRNEDFSYSHILSSMCTTPHPVAVQAHNLFMETNLGDPGLFPG
TATLEDRLIRWFADLYHEPSAGGCTTSGGTESNIQVLRFCKKTKNVKEPNIIVPASAHFS
FEKACGMMDIEMRVVPVDEQYRMKTDAAGELIDKNTCCIVGVAGTTEYGMTDPIPALGKL
AEQEGVHLHVDAAFGGYVLPFLDDAPPFDFSVPGVGSIAVDPHKMGLSTIPSGVLMVRDE
RVFCNLLVETPYLTTKQAYSLTGTRPGASVAAAYAVMAYLGRKGMKALVTGCMENTRRMI
EGMEAFGVHRKVTPDVNVATFEHVSVPSPWVVSYTRKGDLRIVCMPHVTRDVVEAFLSDF
GESYVSHIS
>A_L0JVZ3
MQAEPQAFDRVLSSMCTKPHPDARDAAERFLATNPGDPATYQRVAELEDEAVALLGEVAG
LDDPAGYVASGGTEANIQAVRIARERTDSPRPNVVVPESCHFSFRKAADVLEVELRVVPT
DDDHRADLTAVRASVDSDTALVAGVAGTTEYGRVDPIPELGEIADSVGATLHVDAAWGGF
ALPFTDYEWHFGHAPVDTMAIDPHKMGQAAVPAGGLLARSESLLDELAVDTPYLESTSQA
TLTGTRSGAGVASAVAAMRTLWPEGYREQYARSQRNAEWLADALEKRGYDVVDPTLPLVA
ATVPRPTFEALRAEGWRLSRTATGELRVVCMPHVTREMLASFVADLDRLEVRASVPVVGD
D
>A_K4MEZ9
MCTAPHRIAVKAHMQFIESNMGDFGLFRGTHEMEKEVIRMTGNMLHCPFTEGYLTTGGTE
SNIQAVRSMRNLHERKHSGSRLNVVVPISAHFSFDKVSDILDIDVRKAPLDSDLKVSIKA
MKSLIDVNTVGLVALAGSTEFGQVDPIGKISELALGKDLPLHIDAAFGGFVLPFLAQEHV
FDFSLPGVTSIAVDPHKMGLSTIPSGILLFKEFKHLRCLKAHTPYLTVDSQYTMTGTRSG
AAVAATFAVMKFLGKEGYTETVSKCMEMTRYLLRKAAEIGVEPVIDPVINVVALKVPEPA
VVRATLSREYNWHVSITQDPKALRLVIMPHMSCEMIDMFMADLTKVLGSCT
>A_I7LLL5
MREVGCPEEELFSFLSLSRQEDLGYQNILSSMCTPPHPVAARAHAMFLETNLGDPGLFPG
```

TAALERLLVRRLGALMHLPEAGGYATSGGTESNIQAFRIAKKRKRTRSPNVVVPESGHFS
FQKACDILGLEIRTVPLDAEFRMDVDAVDGLVDNNTIALVGVAGTTEYGVVDPITRLSEI
ALDREVFLHIDAAFGGMVVPFLDRPIPFDFRLPGVNSISIDPHKMGMSTIPAGCLLVRDP
EYFSSLNVDTPYLTVKQEYTLAGTRPGASVAAAVAVLEYLGMDGMRAVVAGCMENARRLI
EGMETLGYPRAVTPDVNVATFSCDRAPAGWRVSRTRAGDMRIICMPHVTRDVVEAFLGDM
SDLDA
>A_Q0W498
MRERGLGEEEIFAELCEARSRDVPYGRVLSSMCTNPHPIAVKAHQEFVNTNLGDPKLFPG
TADIEHRCIGLIGDLLHLPAATGYISTGGTESNIQALRTAIQMKHTDRRRANIVVPESAH
YSFEKASQMLGIAIRRAPLDDLLRADPSEMAALIDKNTIALVAVAGTTEFGQIDPIEEIG
RLAQEHDLYLHVDAAFGGFVIPFMDRPAKFDFEIPGVQSITIDPHKMGLSTIPSGGLLYR
SESLMKVLEINAQYLTSMVQTSLAGTRSGASAASAYAVLQYLGRAGYREIVATCMENTRI
LREQLEDMGMEPIIEPVLNIVTARAKDPVGLRKKLAEKNWYVSTTVHPCALRMVVMPHVT
ADVIEAFTADLKKVI
>A_E4NVT4
MSFQEAEGNESAPDGTATDAVDGQTLAERMFIGTPAGNDAYIAAIDRCRDAVLSAVGEAD
RPYSGSTYAEHRERLTQDTIPESGRKIDEVIEDLATDVLEESVYPSDEACSAHLQCPPMV
PSLAAEVVLSALNQSMDSFDQAPAATVLEEQVIDDLTSLFGLGDGADGVFTSGGTQSNLQ
GLLLAREHYVAEVFDRSVRTSGLPPTAEKMRILTSEDAHFTVAQAAAQLGLGEDAVVTVP
TGDAHQMDPDELADELARLKQANCHPFALVGTAGTTDFGSVDPLNDLADLADEHDLWFHV
DAALGGALALSETHAGKLDGIERADSLTVDFHKLLYQPISCGVFLLSDGDKFELMGRNAA
YLNPKSDQVSNLVSKSLQTTRRFDALKPYVAFRTLGREGMAALVDRTVTLADRAASLIRS
DPGFELSCPPTINIVTFRYTPERDHPARSPDEWADRINRRARDRLLASGCGVVARTEVDG
CVHLKLTLMNPRTTVADIRELLLTLKGYAAQAETKAISNHGIESDDRLDFAPVCGSADEE
SR
>A_A0A1F2P723
MMEDGICFEDVLAELEELKRRDLGYTRILSSMCTHPHPIAKIAHNLFLEANLGDPGLFPG
TKAIEEEVISMIAALLGDRDATGYITTGGTESNIQAIRAFRNASGKDGENIVVPASAHFS
FDKIGDLLRVEVRKAPLDGEFRVDPAAVEDLIDDRTIGLVGIAGTTEYGQIDPIEDLAKI
ALENDLFLHVDAAFGGFVIPFLDSHPPFDLSVDGVSSITIDPHKMGMSTIPAGGILFKDK
RLLELLSTPTPYLTSKAQFSLTGTRSGAAAAATYAVLRYLGFKGFKRVVDRCIRMTKMLK
DGASSFGVVPPVEPVMNVLTLEVKDLRQVIRELERSGWRVSVTREGFMRLVIMPHLTEPI
LQEFLDDLKDATGN
>A_D2RU46
MTGQRSAVDRDRPPAADADPTPPPTAATAFLGGPDGNAAYADAIERARDVLLESFATSAG
PYAGTDHETLRERIADLQVVPDDGSSIEDTLETVADDVLADSVRVHDPGCVAHLHCPPTV
PALAAELLLSGTNQSMDSFDQAPAASVLEERVVDACCDLFDYPTGADGVFTGGGTESNFL
GLLLARDWYCERRFDRDVQTEGLGPEAASDLRLLCSDAAHFTAEQAAHHLGLGEDAVVSV
PTDDDRRIDLEALDSTFERLEADGRHPFAIVATAGTTDFGSIDPLAALADRAADRDLWLH
VDAAYGGACAISDRLRPKLEGIDRADSIAVDFHKLFYQPIGCGAFLLRDGDRYRHLERNA
AYLNPERDDAAGVPNLVSKSTRTTRRFDALKPFVTFNALGRTGVADCVEYVCELADAVAD
EIRADPALELCCDPELSAVVFRYRPETDSESGSLPTAAVDRVNRAIRDELLADGEVILAR
TEVDGTAALKLTLLNPKTTLSDLRDVLEAVVDRGEALETDREVIDSA
>A_D2RX89
MQSEPQAFDRVLSSMCTEPHPVARDAAERFLATNPGDPGTYPSVSALEEEAIAMLGSIAG
LEEPTGYIASGGTEANIQAVRIARDRAESQRPNVVMPESAHFSFQKAADILGVELRIVPT
DDNFRADLEAVRASVDEATALVIGVAGTTEYGRVDPIPELGEIARSVGAMLHVDAAWGGF
VLPFTDYEWNFEHAPVDTMAIDPHKMGQAAVPAGGLLARSDDLLNELAVDTPYLESTSQA
TLTGTRSGAGVASAVAAMEELWPEGYKRQYVRSQNNAKWLADALEKRGYDVVDPTLPLVA
ADVPRSTFDALRAKGWRISRTATGELRIVCMPHVTREMLASFIGDLDRLEVRASVPVASD
D
>A_A0A075LW21
MIPEDGLSEEEVMGELEKRLSLDLTFDSGKILGSMCTYPHPLAQKIVRKYMDRNLGDPGL
HVGSQKIEREAIQMLGELLHLKKAHGNIVSGGTEANILAVRAFRNVSDVEEPELILPKSA
HFSFLKASDLLKVKLVWAKLNKDYSVNVNDVESKISDNTIGIVGIAGTTGLGVVDDIPSL
SDLALDYGIPLHVDAAFGGFVIPFAKVLGYELPDFDFKLKGVQSITIDPHKMGMAPIPAG
GILFRKKKFVDSISIPAPYLAGGKVSHPMITGTRPGASAIAVWALLKHLGFNGYKEVVRE
AMENALWFAEQIRALKGIYLIREPMLNIVSFGAKNLKTIERALKDRGWGISAHRGYIRIV
MMPHVRREHLEAFLEDLREILATL
>A_I3R9H1
MSDAESLHDTRPDMEGLFLGDDENDREAYQEAIEQACDLVLGEFLDNATPYSGVTPDELA
DTLAQFEMLPSEGDGLETALDRTGPVLRNSVGVSDPHCIAHLQCPPMVPALAAEVLLTAA
NQSMDSWDQSPAATHLEEQFVTELCGLFGYDEAESDGVFTSGGTQSNFVGLLLARNRILL
EEYGVDVQQAGLPPEARDLRILCSEAAHFTAKQAASHLGLGENAVVTVPTDNEYRISVEA
FDEAIVDIRANGNRPFAIVATAGTTDFGSIDPLEPLAERARKYDCWFHVDAAWGGALALS
DEHADKLAGIEAADSIAVDFHKMFYQPISCGAVLVRDESSYDLIDRNAAYLNPERDDEAG
VPNLVSKSVQTTRRFDALKPFVTMQTVGREGLASLMEYTINLADEAVELIERDSDLHVIH
ESELNVVLFRYVPESVPIEMTREEWIGKLNEAIRDSLLEDGEAVVARTTVDGINCLKLTL
LNPRTTREDIRSLLQAITARGTEFETNPDEIRQ
>A_I3R6X8
MNRSNSPETGFIDPSGANAEAIRDLTEDVLDQLLGQLGAAEKRSPLPDESTAPTGTIPES
PRSQTDLLDDLETIAAGSMNPAHPGYIGHMDTMPTTVSVLGDLVASAVNNNMLSVEMSPV
FSELEVQLTETIASEFGLGPNAGGVLASGGSLANLHALSVARNQAFDVHDDGLAGLDGEP
VLFASDVAHTSLQKAAMLLGLGTDAVVAVETNANSRMKPSALNQAVEQAERDGRVPFCVV
ATAGTTTTGNIDPLPAVRDVVDEHDLWFHVDAAYGGALVFSEAERDRLDGIEGADSVTFN
PQKWCYVAKTCAMALFADLDILQEDFRVGAPYMRGDDAIPNLGELSVQGTRRAEVLKLWL
TFQHLGREGLGQLIDESYRLAAVIRDRVADQDALELASEPEMNIVCFRAAPDWCPPDERD

ALNGRLQRYLLSRQDVFVSLPTYRDTRWLRVVLLNPFTDKTTLDRLFDGIDLFLEAERP
>A_I3R7R5
MSEGFDSLDPDPEEFRELGYRAVDMMAEHFANIRAVDTFPETTPPEVAEEFDDPLPRDGE
DPEAVLAEWNERVYPNATHQGSPRWYGYVMGSGTPIGALADALAASVNMNAGAWMGGPSA
TEIERQCLQWLAEMIGYPADCGGVLTSGGTMANHAALYTALQSATEFETRESGLRSIDRP
GRFTLYESAHEGHSSAERVAEMIGVGSDAIRSVPCDEDLRMDPAALDDMLTADVENGRIP
FCVIAYVGSINVSTVDPLAEIADVCADHGVWMHADGACGAVGAILPEKEHLYEGIERADS
VTLDPHKWLSVPYSCGCVLFRDPDAQTQAFSMHAEYLDFTEEEAYHGTNLGFLGPEMSRP
FRALKLWMSLKHRGVEGYRQLLRQNCRCAEHLHDRVVAADDFEVLQEPNLFIYSFRYLPT
DLRDAVADADQREAINDYVDWLNQRITDELRLTGEAFVTTTEIRDDTAIRLSICSHRTTP
ADIDTTFEALREHGEHIDAEGRKRVDSEFADN
>A_A0A1G8UBX3
MQQAQQAGLDGMQPQDFDRVLTSMCTEPHPAAREAAERFLATNPGDPGTYQTVANLEDRA
VELLGAVTGLDEPAGYITSGGTEANIQAVRIARNRAETDDPNVVAPKSAHFSFRKAAELL
GVELRTAPTTDHRANVDAMNELVDEDTVALVGVAGSTEYGVVDPIPEIAELATDAGALCH
VDAAWGGFYLPFTDFEWQFDHAEIDTLTIDPHKVGQAAVPSGGLLARSESVFDPLAIDTP
YLETTSQVTLTGTRSGAGVASAVAAMEALWPEGYREQYRVSMDNAEWLADQLRARGHDVF
GPVLPLVTADLSVPMTDELRSRGWRVSKTGSDEMRIVCMPHVTRSMLRSFVADLDWY
>A_A0A2V3JKE4
MNTHGMDEKTIGSALHDLKSRDTPYERVLSSMCTYPHPVAVAAHQQFIETNLGDPGLFAG
TAEIEHEVVRMMGTLFGNPDAHGYVTTGGTESNIQAIHAIKTARKIREPNIIVPASAHFS
FDKVADILGIEVLKADLDTEFRADIQAVEDLIDGNTIGIVGIAGNTEFGQIDPIRELSDL
ALSKNLFLHVDAAFGGFVIPFLTEKYEFDFTLPGVTSIGADPHKMGFATIPSGGLLFQDS
SYLHRLSVDTPYLTVNSQQTLSGTRSGASAASAYAVFKHLGREGYERIVRRCMKLTHELV
ARASEFGIEPLIDPVTNVLVLDVPDADSVRSAMKTRGWDVSITRDPRALRLVIMPHLSSE
NLNLFADDFADVVK
>A_M0L1G1
MRAEPQSFDRVLSSMCTEPHPAAREAAERFLATNPGDPGTYQTAAALEDRAVAMLGEIVG
LEIDTADAIGAADEDGSGGPTGYVTSGGTEANVQAVRIARERAARGTDRPSVVVPESAHF
SFRKAADLLQVDLEVVPTASDHRVDLDAVRAAVDEDTAAVVGVAGSTEYGRVDPIPELGE
IATSVDALLHVDAAWGGFALPFTDREWHFGHAAVDTMAIDPHKMGQAAVPAGGLLVRSAD
LLDELAVDTPYLESTSQATLTGTRSGAGVASAVAAMEELWPDGYRDQYVRSQHNAEWLAD
QLESRGYDVVEPVLPLVAASIPTWLFESLRAEGWRLSRTGDDERVRFVCMPHVTREMLES
FLADLDRLESRAARAVPSPGE
>A_C6A1N1
MAGESMIPNKGISEEELFTELEKRLKIDLTFDSGKILGSMCTYPHPLAQKIIQKYIDRNL
GDPGLHRGSKEIEEEAVQMLGELLHLKRAYGNIVSGGTEANVLAVRAFRNVSNVEKPELI
LPESAHFSFLKASDLLKVKLVWADLNRDYSVNVKDVESKITDNTIGIVGIAGTTGLGVVD
DIPALSDIAVDYGIPLHIDAAFGGFVIPFAKALGYELPDFDFKLKGVQSITIDPHKMGMA
PIPAGGIVFRKKKYMDAINVLAPYLAGGKIFQATITGTRLGANAIAVWALFKHLGFEGYK
NVVKEAMENALWFAEQIKRLDRVYLIREPMLNIVSFGSKRLKKIEAELKARGWGISAHRG
YIRIVMMPHVKREHLMNFLKDLREILRRV
>A_A0A2E4GBW7
MTGKKKTAQASIEAMYRVFTVPEAPDSTLSRIDQDISRNLAGFLQEHIVAVERDLAEVEK
DFADYNIPEKPIFVSEQAQFLLDKLVANSVHTASPAFIGHMTSALPYFMLPLSKIMIALN
QNLVKTETSKAFTPMERQVLGMIHRLVYQEDGAFYRKWMHDPRHTLGVMCSGGTIANLTA
LWVARNHAFPAEGSFRGLHQEGLYRALKYYGYEGAAILVSKRGHYSLRKAADVLGFGRDA
LVSVETDDDNRILPDALREKCLELQKQKIKVLAICGVAGTTETGNVDPLDAVADIAREFG
AHYHVDAAWGGPTLFSRSHKHLLRGIEKADSVTFDAHKQLYVPMGAGLVVFKDPALASSV
EHHAQYIIRKGSRDLGSTTLEGSRPGMSMLIHSGLRILGREGYEILIDQGIEKARTFAGM
IDAEPDFELVTRPELNILTYRYCPEPVQRALALADEFQAEKMNTCLNRITKFIQKTQRER
GKAFVSRTRLEPARYYHFPCIVFRVVLANPLTTRDILADILKEQRMLAQEEGIADEMSIL
HQMAEAVLKQRQPGARQA
>A_A0A2E4TVH8
MDTETFLDEVLSRVKIFLDSSQSDVRIRTEQTHDSLLRTSDLKLPMEGRGLESALDDIES
VLSHSVRTTAPGFMNPLWGGLSITSLAGELVTAATNTAMYTYEIAPIATLIESSILKRMA
ELADFGTSQGTLTTGGSNGNMLGLLCARQSKVPLSSQSGFDGTKMVAFVSEESHYSFNIA
SNVVGIGQSNLIKIRCNEQGQMRADSLEDEIERALENGQIPFAVLATSGTTVRGSFDPLR
EVAEIAHKYDLWMHVDAAWGGSCLFSTQYRSLMDGIELADSFCWDAHKMMGIPLICSAFI
VKDAEILRAVCSNGNTAHYLYLETGEDVDLGLYSLQCGRRNDALKLWLAWREIGDAGWAT
MLDGFMKLADYLERRVNQNESLEMMSNRMWTNVCFRYVGSSPEENLNHINTELRKRLIHD
GRFMVSRSTIDGNIVLRSVIANRSISEASLDSFLECVVSIGKDIERGLPPNQ
>A_D8JB87
MSADGLFLGTDRGDAAYRAAMEQATDAVLCAVAAREEPYSGASPDALAEYLDDPVVPEEG
RGLEATLDEVAERVLAHSVDPSNPRCGAHLQCPPMVPGLAAEVLLSATNQSLDSFDQAPA
ATLLEERVVGALCDLFGLPAGADGVFTSGGTQSNFQALLLARDQYCARQFDRDVQAEGLP
PEADSLRLLCSAEAHFTTNQSAHHLGLGEDAVVTVPADGDRRMDPGALEATLAALRKRGA
VPFALVGTAGTTDFGSIDPLGALADAAAEHGLWFHVDAAYGGALAVSDEYGDLLAGIERA
DSVAVDFHKLFYQPISCGALLVRDGDEFRWMARNAAYLNPEAHDDRGVPNLVSKSVQTTR
RFDALKPYVAFRALGRSGMAALVERTLELADEAASLLESADDFELLGEPTLNAVVFRYRP
CEGMTDEAASRLNADVRRELLADGRAVVARTEVDDATCLKLTLLNPTATLKDVGAILEAV
RECGSAVADPGEVIA
>A_A0A2R4X059
MLEATPQDFGRVLSSMCTEPCPPARAAADRFLATNPGDPGTYPEIARLEERAVDLLGEIT
ELADPAGYVATGGTEANIQAVRIARNRADTDDPVVVAPESAHFSFHKAAAMLDVEIRTTP
LVDYRADPDAIAEAIDSDTALVVAVAGSTEFGRVDPVPALAELAHDAGALCHVDAAWGGF
HLPFTDHDWSFSDAPIDTMTIDPHKAGQAPIPAGGLLARDESLLEVLEVETPYLESSTQV

```
TLTGTRSGAGVAGALAAMETLWPDGYRDNHERAMANATWLADELRERGYEVVDPELPLVA
ADVSQSIIDALRRRGWRISRTGAGDLRIVCMPHVTRSMLRSFVADLDWY
>A_N0BK04
MNSQNNLEYSKYSEHAETSETIRILKEFRSQDIPYNRVLSSMCTTPHPLALKAHEMFIET
NLGDPGIFQGTKKLEEKLIGMIGELLHGSEVAGYICSGGTEANIQGIRAGRNKMRDKIKD
SRNPNIVIPKSAHFSFEKIGDILGVEVRRAKLDEEYKVDVSEVEKLMDENTVCLVGIAGT
TELGQIDPIVELSKLAEENCVELHVDAAFGGLVIPFMNNPYPFDFQNDGVSSITIDPHKM
GMATIPSGGILFRNESYLRALEVETPYLTSRTQFTLTGTRPGTGVASAFAVLHGLGFDGM
KKIVMECLKNTRLLTEEMISLGFEPVIDPIMNVVSFKTEKAEKIRNELMKRRWIISSIKE
PRAIRFVIMPHVTEEVIKEFLSEFRKIINWV
>A_Q5JJ82
MFPERGASEEEVLRELEEKTREDLTFDSGKILGSMCTYPHPFAVKVVMKYIDRNLGDPGL
HIGSQKIEKEAVDMLANLLGLEKGYGHIVSGGTEANILAVRAMRNLAGIEKPELILPESA
HFSFIKAAEMLGVKLVWAELNDDYTVNVKDVEKKITDRTIGIVGIAGTTGLGVVDDIPAL
SDLALDYGLPLHVDAAFGGFVIPFAKALGYEIPDFDFRLKGVKSITIDPHKMGMVPIPAG
GIIFREKKFLDSISVLAPYLAGGKIWQATITGTRPGANALAVWAMIKHLGFDGYKEVVKE
KMELARWFASELKKIPGIYLIREPVLNIVSFGSEKLEELEKELKARGWGVSAHRGYIRIV
VMPHVKREHLEEFLRDLREIAKRL
>A_A0A1H3WVB1
MEQAVPQTFDRVLSSMCTRPHPVARAAAERFLATNPGDPETYRAVADLETEVVETLGELT
GLDQPRGYVASGGTEANIQAVRAARNRASTDDPNVVAPSSVHFSFQKAAEVLGVDLRIAP
VGSDRRADPEAMASLVDDDTALVVGVAGTTEYGRVDPIPELADLAAEADALCHVDAAWGG
FALPFTDHEWHFGHAAVDTMTIDPHKMGQAVVPAGGLLARDAATMDALAVETPYLESTSQ
ATLTGTRSGAGVASAAAALDELWPDGYRENYERAQANAEWVADALADRGFDVVDPVLPLV
AVDLPDSLFAAVQDRDWRLARTARGELRLVCMPHVTRDTLSAFLRDLDACRDEI
>A_H1YYJ3
MQKKGCPEEEIFSFLSFARDKDRKYDKVLSSMCTIPHPVAVRAHNMFIESNLGDPGLFAG
TAELESLLVREIGELMHIPDACGYATSGGTESNIQALRIAGKQARRKMPNVVVPESVHFS
FEKACDILSYELRTVPCDGNQKIDTSVLEDYIDKNTVCITGIAGSTEYGVVDPIEHLSDI
CSDREIFLHIDAAFGGFVLPFLKNAPKFDFELDGVSSISVDPHKMGMSTIPCGCLIARDP
SYFKSTEVETPYLTVQKECTLLGTRPGGPVAGALAVLRYLGRSGFEEIVGKCMNNNRRLI
DGMADLGYEVAVQPDVNVASFKCENSPKGWIVSRTREGHMRTVCMPHITEDIIDEFLKDV
SEINV
>A_Q8U1P6
MKFPRKGIPQEEVMRELEKYTSKDLSFSSGKILGSMCTLPHELAKEVFCMYMDRNLGDPG
LHPGTKKIEEEVIEMLSDLLHLERGYGHIVSGGTEANILAVRAFRNLADVENPELILPKS
AHFSFIKAGEMLGVKLIWADLNPDYTVDVKDVEAKISENTIGIVGIAGTTGLGVVDDIPA
LSDLARDYGIPLHVDAAFGGFVIPFAKELGYDLPDFDFKLKGVQSITIDPHKMGMAPIPA
GGIVFRHKKYLRAISVLAPYLAGGKIWQATITGTRPGASVLAVWALIKHLGFEGYMEIVD
RAMKLSRWFAEEIKKTPGAWLVREPMLNIVSFKTKNLRRVERELKSRGWGISAHRGYIRI
VSHASCDGGHD
>A_D1YVJ9
MREHGVDEDTIIRELKGACARNVPYERVLSSMCTTPHPIAIKAHKEFIVSNLGDPRLFPG
TASLEHACIGMLGELLHLPSAVGYITTGGTESNIQALRTARQLKHVDPGKANIVLPESAH
YSFDKAAQMLGVSLRRTPLDDEMKADMDAMAGLVDKNTIALVAVAGTTEFGQVDPIPAIS
KLALDENIFLHVDAAFGGFVIPFMKDPSKYRFDFELPGVMSIAIDPHKMGMSTIPSGGLL
YRDERHMKSLEISAQYLTSQVQSSLAGTRTGASAAATYAVMRHLGMDGYRRVVSECMDNT
MFLRDSLVDMDIELALEPIMNIVTAKLPDAQSTRKKLCDMGWFVSTTSRPEALRMVVMPH
VTRDVIEAFMADLKKIS
>A_B8D379
MYIGDINKAREWLEKAFSKTPNHLDGSILGSMTTMPHELGVEAFLRFIHINGNDPMVFPI
VKEAEEIIVKGIGGLFDVEHGMYTSGGTESNIMALYVGRRVNKGKENTVVVPSSIHRSID
KACLLMGCKLVKIPVDPLKPVDPAILEEYIRLYKPFAVVVTAGTTEAGVIDPVKEAGELA
EKYGVYLHVDAAYGGLLIPFLYRRGYITVDLRMFPGVSSLSVDMHKNGCAPIPSGLLFFS
NRGFLEQACFDMEYMPLGKSCGLLGTRPGGAVVASAAVFMAMGIKGYEENAVKMMENSYY
LYNGLKNIPELVVFKPILPINVFRSLRYSYIELFKVLAEKGVYVYKSPSLHALRVVVMPH
VSRQHLDKFINILKLIHGSG
>A_G7WKU8
MRDRGLSEEEVMRGLLKMRAKDLSYDRIFSSMCTPPHPIALKAHQLFQETNLGDPGLFPG
SAELEAEAVRMMAELLGHPEACGYLSTGGTESNIQAIRAARNSADFRDGNIVVPRSAHFS
FDKIGDLLSLEIRKADLDGDLKVEVGSVEELIDEKTVSLVGIAGTTEFGQVDPIDRLGRL
ALDWGIPLHVDAAFGGFVLPFLGGDWRWDFSVEGVTSITIDPHKMGMATIPGGGLLFRHP
EDLERLAAYAPYLTVARPKALTGTRSGAAAAAIWAVMSHLGMEGFKEVVNGCMALSRRMA
SGAKEIGIEPVIEPVMNVVTLRMEDPEGVRAALLGRRWRVSTTRSPKALRLIMMPHSTAE
NVDLFLGDLEDVLRKKA
>A_Q6M0Y7
MDEQDILNELREYRNQDLKYEEGYILGSMCTKPHPMARKISEMFFETNLGDPGLFKGTSK
LEKEVVSMIGGILHNKNAFGYLISGGTEANLTAMRAFKNISKSKGKPQNIIIPETAHFSF
DKAKDMMDLNVVRPPLTKYFTMDVKFIKDYIEDSKNEVSGIVGIAGCTELGSIDNICELS
KIAVENDILLHVDAAFGGFVIPFLDDKYKLDGYNYDFDFSLNGVSSITIDPHKMGLAPIS
AGGILFRDNMFKKYLDVDAPYLTEKQQATIIGTRSGVGVASTWGIMKLLGIDGYETLVNE
SMEKTMYLVKKAREYGFETAIDPVMNIVALNDENKHDTCMKLRDENWYVSVCRCVDALRI
VVMPHLEIEHIDGFLESLSNTKKY
>A_A2BJD5
MEWLRGGSWEEVARELGELRAGEPSPCRVAGSTVAEPLPVARRAYSLYADVNLNDPASWP
SVTKLLEGISRVLEELRLGHRWLVAVSGGSEAVLTGLYIAREYTRGRVVVASSAAHASVL
KAARVLGMEVKLVQVDSRLRIDLYALEKTLRGVQNVAAIVATAGVTDNGAVDPVRDVAKL
```

AWEHGAVVYVDAAFGGLPLLGLGSTETVLPRGGPALAGIDFHKHVAPPPSSILVSNTAEL
RDYIVFPAPYMPLGRQETLLWTRPASGLAAAYAALRALGASGVGELARYLYRLASKLASI
LEQRGVELLSPLDTPLVAFRPPSVEGALKRLRRRGWILYPSRLPGILRYVAKWCHEPGDV
EEIAEAVA
>A_A0A062V1G2
MSDPIMDGYFIHTISHFLDHVDSLKCIAPVLGIGYEKCMEGTRRNIDREKLAMLSRIKQG
EQSENAHNQNFPDEMSSIEKVTELLADYCLGMTIWAHPNAQANVVPPPTIPSITAFIAAA
IYNPNLIWDEYSCMFAEAELEAVAMLSDLVGYDSKKSSGIFTFGGTGTILYGCKLGVEKI
FSGRAMTEGVRDDVKIVASESSHYSRLNVAGWLGVGTKNVVTIPTTRENEMSLTDLEDYL
RHAFETGEKVAIILATLGTTDTFGIDDLASIVRLRDKLAAEYRLEHLPHIHADAVIGWAW
AVFKDYDFEINPLGFHARTLRSLQDSLQRISSLHMADSIGIDFHKTGYAPYISSAMLVKN
RQDLILLSRAPEQMPYLYQYGYYHPGIYTLECSRPGTGALAALANMRLLGKQGYRVLLGH
VVEMAEMLRDQLERHTFIKILNDYNYGPVTLFRVYPDGADAEEIFQRELTDPDYREQLEE
YNTYNLRIFKQIHERAMRGEGILLSWTDACRHANYPDGPPVAALKSFIMSPWTNLKAVDM
VARQVLEARMQAKK
>A_A0A1Q1FJ77
MSADELFLGSAAGNATYREALDRAADAVVEQFAAADRPYSGERPEALAARLEQPVLPESG
IGLRAAIDEVASEILPHSVGTSNPRCAAHLQCPPMIPGLAAEALLTATNQSLDSFDQAPA
ATVLEGRVVDALCDLFGYPDAADGVFTSGGTQSNFQALLLARDRHCQETFGTDVQASGLP
VEAGSLRVLCSEAAHFTAKQAAHHLGLGEDAVVTVETDDRRRLDPAALDARLDALEAAGA
EPFALVGTAGTTDFGSIDPLPALADRAAEHDLWFHVDAAYGGALALTDDHAAELEGVDRA
DSVAVDFHKLFYQPISCGALLLRDGGDFEWMARNAAYLNPAAHDETGVPNLVSKSVQTTR
RFDALKPYVAFRAVGRDRLGDLVAGTLDLAAETAALLEAADDFELVTDPTLNAVVFRYRP
EGGMADERVDALNAAVRRRVLEDGRAVVARTEVDGVTSLKVTLLNPTATLDDVAAMLDVV
RDCGADLRDGPEVSA
>A_A0A1Q1FH74
MQRVEPQDFSRVLSSMCTKPHPAAREAAEEFLATNPGDPGTYETVSELETEAVDTLGTIT
GLPNPAGYVTSGGTEANIQAVRIARNRAETDRPNMVAPESAHFSFNKAADILNVELRTTP
TDDDHRADPDAMAQVIDEDTVLVAGVAGSTEYGRVDPIPAIADLANEVDALCHVDAAWGG
FVLPFTDHRWHFGHADIDTMTIDPHKMGQAAIPSGGLVTHSKSLLDELAVETPYLESKSQ
ATLTGTRSGAGVASAVAAMDALWPVGYREQYHESMENAEWLADRLAARGHDVVDPELPIV
AADMSMPVTEDLRQRGWRVTKTGAGEMRVVCMPHVTRSMLRSFVAELDWF
>A_A0A328SBH2
MLDDGRNVDSIFRDLEEFKKMNMTYESGRILGSMCTKPDPIGLKAYKMFIETNLGDPGLF
EGSAKMESQVIDMLGKLLHLDDASGHIVTGGTEANFMAMTIAKYLFLENNSGVPEVILPC
NAHFSFKKICPMLSVKPVYVPLKDYRMDCSVIEEYITDNTMAIVALAGSTELGLVDDIAR
ISEIAQKNRLYLHVDAAFGGFIIPFMKNDKSHPLNFDFSCEGVDSITIDPHKMGLAPVPA
GGIIIRDKKHLEKLSVKTPYLTKEKQTTVVGTRTGASTAATWTLLNYYGKSGYKRIVNES
ISLTKYTYNQLKQIPQIKLTCKPDLNIISFKVENMDSKLLQRKLLKLGWRVSVSENPYAM
RLVLMPHVKKEHITEFIKTLKKVIEEK
>A_Q2NHY7
MFDKGRSKEDVFRDLNVFHNMDMKYSSGRILGSMCTKPDPVGLEAYKMFIETNLGDPGLF
KGTALMEQEVINSLGNLLHLKNPCGHIVTGGTEANIMAMCVAKYLYEEENEGTPELILPK
SAHFSFKKVLSMLSVKPVYVPLNNEYKIDVTKLPDLITDNTMAMVGIAGTTELGLVDDIP
EISKIAKSYGVYLHVDAALGGFIIPFLNYKNNNQLNFDFKCKGVSSITIDPHKMGLAPVP
SGGIIFRKKKYLEKLSIKTPYLTKDKQTTIVGTRTGASTAATWTLLNYHGMEGYKKIVEK
VINLTTYTYNKLNKNKHVTIIHKPELNIISFKVDNIDVDTLQKQLQAYGWIVSLAEYPHV
IRLVLMPHIKKEHIDEFLVDLDIIIQKNR
>A_A0A1D3L4E9
MENKGISHEEVLQSLRQFKKLDMTHKSGKILGSMCTCPHPVGVEAYKMFLESNLGDPGLF
KGTQKMEDEVITMLGELLGKRDVSGHIITGGTEANIMAMRAARNSAQHKRHIDEPEIIVP
KSAHFSFKKAADMLCLNLHEAELDDDYRMKMDSVRELITENTVAVVGVAGTTELGKIDPI
AELSELCLERDIYLHVDAAFGGFSIPFLKEAGHDFPEFDFGLEGVCSITIDPHKMGLAPI
PTGGILFREKKYLEAMSVKTPYLTKDRQATIVGTRTGASTAATWALMKYMGREGYTHVAE
RCMEVTSILAQGIEESDFKLVTQPQLNVVAFTSDEMSPDEISSKLKDKGWAVSIASYPRA
VRIIVMPHVKEEHVETFLKDLNELE
>A_A0A1D3KZK6
MEGLNTLSGKKNLEKMKVSEKEVTSTYGSRYFTESIPKYVMPEDEMPARAAYQLIHDELN
LDGNPALNLASFVTTWMEPEADKLIMESMDKNFVDNDEYPQTEKIQERVINMLARLFNAP
KECHSVGTGTIGSSEAIMLGLLAHKWTWKKRRQEEGKPFNKPNIVMGADVHTVWEKFALY
FDVELKLIPLERDTYTVTVDKVAEEIDENTICVGAVLGTTFTGQMDPIKEINDLLMDIKK
ENGWDIPIHVDGASGGFVAPFLYPELEWDFRLEQVRSINVSGHKYGLVYPGVGWLIFKDK
TDLPEELIFKVNYLGGLMPNYSLNFSKGSSTIIAQYYNLIRLGKSGYTDIMENMMSNSQY
LARKLEDSGKFEIINKEGMFPLVTVSLKDEEFTVFQLSEKLRQKGWIVPAYTLPENAEDV
AVMRMVVKENFGREMIDLLVDDVMEACESFEGEETVEKVEDQNPSLLY
>A_U1QCQ6
MMEPRQQQRPQDFDKVLSSMCTEPHPLAQEVAVECLAMNPGDPATYQTVAELETEAITRL
GEVAGIADPYGYIASGGTEANIQAIHAARNRASGEHLSGQPNVVAPESVHFSVNKAAEML
GVDLRVVPVDSDYRADLDAVRAAVDTDTVAVIGVAGSTEYGRVDPIPALTDIAHDVGAHM
HVDASWGGFVLPFTDYAWSFADAPIDSMAIDPHKFGRAPIPAGGLLAREQATVDALAVET
PYLETTSQATLTGTRSGAGVAGTVAVMEELWRAGYHQQYHTQMSNATFLAEQLTNRGIEV
APPTLPLVTAAVQSATIEALQAKGWRLARTTSGDLRIVCMPHVSRASLEAFLDDLDHVRS
>A_B6YUX2
MFPRKGASEEEVLAELEEKTAEDLTFDSGRILGSMCTYPHPFARKVISLYIDRNLGDPGL
HVGSQKIEEEAIQMLSNLLGLEKGYGNIVSGGTEANILAVRAFRNLADVEKPELILPRSA
HFSFLKASEMLSVKLVWAELKEDYSVDVNDVERKITDNTIGIVGIAGTTGLGVVDDIPAL
SDLAIDYGLPLHVDAAFGGFVIPFAKELGYDLPDFDFRLKGVQSITIDPHKMGMVPIPAG

GIIFRKKKFLEAISVPAPYLAGGKVWQATITGTRPGANALAVWAMIKHLGFEGYKEVVKG
AMELSRWFAGELKKIPGVYLIREPMLNIVSFGTTNLEEVEEKLKRRGWGISAHRGYIRIV
MMPHVRREHLEEFLRDLQEIITR
>A_A0A1I6LFT2
MQRAAPQDFDRVLSSMCTEPHPAAREAAERFLATNPGDPGTYEQVAELEREAVERLGAIT
GLSDPAGYVTSGGTEANIQAVRVARNRASDSTEDPNVVAPEHVHFSFRKAAELLGVELRT
APTTGHRADVDAMAELVDDDTVALVGVAGTTEYGFVDPIPAIADLAADAGALCHVDAAWG
GFYLPFTDHEWGFDHAAVDTLTIDPHKVGQAAVPAGGLLARSADLLDELAIDTPYLESTS
QVTLTGTRSGAGVASAVAAMDALWSEGYREQYERSMANAEWLADQLDARGHDVVGPELPL
VAADLSVPMTAELRERGWRVSKTGAGEMRVVCMPHVTRSMLRSFVADLDWY
>A_A0A0U5H3X9
MPRAEPTPAPQDFDRVLSSMCTEPHPAAREAAVAFLADNPGDPATYPAVADLETEAIDAL
GEVVGLDDPHGYVGSGGTEANIQAVRAARNLADGDVNVVAPESAHFSFQKAADVLGVELL
LAPTDDDHRADVDAVADLADDDTALVAGVAGTTEYGRVDPIPALADVAADVDARLHVDAA
WGGFVLPFTDHDWSFADAPVDTMTIDPHKMGQAPIPAGGFLARDAETLDALAIDTPYLES
DTQPTLGGTRSGAGVAGTHAALDALWPEGYRDQYERSQANADYLAAELREHGYDVVDPVL
PLVAADLPDAEFEALREDGWRISRTATGELRVVCMPHVTREMLDDFLASLSGVGNV
>A_W0I930
MTTFPEKGMSEEEVLNELEKRLSEDLTFDSGKILGSMCTYPHPLAQKIISLYIDRNLGDP
GLHVGSRKIEEETVQMLGNLLHLNKAYGNIVSGGTEANILAVRAFRNIADVENPELILPE
SAHFSFLKASEMLKVKLVWAELNDDYSVNVRDVENKITDNTIGIVGIAGTTGLGVVDDIP
ALSDLAQDYGLPLHVDAAFGGFVIPFAKALGYDLPDFDFKLKGVQSITIDPHKMGMAPIP
AGGIIFRKKKFIDAISVPAPYLAGGQIFQATITGTRPGANALAVWTLLKHLGFEGYKKIV
KEAMELSRWFAGQIKTLNGAYLIREPMLNIVSFGTKELEKVEKELKMRGWGISAHRGYIR
IVMMPHVKKIHLEEFLRDLKEILKVV
>A_Q46DU3
MNEQGLSEKEIFSYLEDVKSEDTDYYRVLSSMCTHPHRIAVEAHRLFIEANLGDLGLFAG
AHRLEKEVIRMLGELLHAQSVEIPSGEACESSVCGYLTTGGTESNIQAIRGMKNLVTEDG
KKSGEILNIVVPESAHFSFDKVANMMGIEVKRASLDPEFRVDIASAESLIDANTIGLVGI
AGNTEFGQVDPIEELSKLALENELFLHVDAAFGGFVIPFLEKPYSFDFKVPGVTSIAIDP
HKMGLSTIPSGALLFRSPFFMDSLKVNTPYLTTKSQFTLTGTRSGASAAATYAVMKYLGR
EGYRKNVQYCMQLTTKLVKEARKFGFEPLIEPVMNVVDLRVPNPDIVREQLLKKFGWNVS
ITRNPRSLRLVLMPHNTARDIEEFLQDLRKVTTEL
>A_O28275
MDIIEELRAYREKDIPYSRVLSSMCTVPHPVAVEAHRMFIETNLGDPGIFRGTVELEAKL
MRLIGDILHCETPAGYICSGGTEANIQGIRAARNVQKKENPNIVIPKTAHFSFEKIGDIL
GVKIKRAGVDEEYKVDVGQVEDLMDENTVAIVGIAGTTELGQIDPIVELSKLAEERQVEL
HVDAAFGGLVIPFMDNPYPFDFQNRGVSSITIDPHKMGMATIPAGGIIFRNESYLRALEV
ETPYLTSKTQFTLTGTRPGTGVASAYAVLKSLGFEGMREVVKNCLKNTRILVEEMRDLGF
EPVIEPVMNVVSFRTDEAERIKEELYRMRWVISTIREPKAIRFVVMPHVTEEVIKNFISD
FRKVLRR
>A_A0A0X3BK81
MREYGCPEEELFSFLSLSRQEDLGYQNILSSMCTLPHPVAARAHAMFLETNLGDPGLFPG
TAALERLLVRRLGALMHLPEAGGYATSGGTESNIQAFRIAKKRKRTRSPNVVVPESGHFS
FQKACDILGLEIRTVPLDAEFRMDVEAIDGLVDNNTIALVGVAGTTEYGVVDPIARLSEI
ALDQEVFLHVDAAFGGMVVPFLDRPIPFDFRLPGVNSISIDPHKMGMSTIPAGCLLVRDP
EYFSSLNVDTPYLTVKQEYTLAGTRPGASVAAAVAVLEYLGMDGMRAVVAGCMENARRLI
EGMETLGYQRAVTPDVNVATFSCDRAPAGWRVSRTRAGDMRIICMPHVTRDVVEAFLGDM
SDLDA
>A_A0A0X8V205
MSATDPVLLSDSKEIQERFSRMIRETLSAIFNSFSDDSAFSGIGPYDLREKINALGFLPE
QGKGFEKVLEETEKEILPHLLRTWSTKYMPHLHSPALTETICSELIIACFNDSMDSWDQG
PAATELEESMIHGLLALYGFPVDKGDGCITSGGSQSNISAIIAARDWYCARKFNWDVKMN
GLPPEYTKLRIYTSEISHFSMDKASHILGMGYQAVRKIPVDSKCRIDVKAFAKMLEDDVA
AGLYPYCAVATFGTTDFGSIDDAVGMRELCDRYGMHLHADAAYGSGLIMSRQFRDRIKAI
SICDSITVDFHKMFLLPISCSAILVKDRELLKCFELHADYLNREEDEEDGYINLVGKSMQ
TTRRFDALKVFMAFQTRGVDGYGKIIDTAVGNAMYFYKRISEDPAFMAPVEPELSSVVFA
LKSGDEVNKKIRRLLLSEGTVIGQTVMDGRVMLKFTLLNPNLEHTQIDAIIKRIKELESS
LL
>A_A0A328SMU3
MFNKGLAKDEVFRRLSVFQDMDLDYDSGKILGSMCTKPDPVAMEAYKMFIETNLGDPGLF
KGTALMENEVISSLGRLLHLSDASGHIVTGGTEANLMAMCVAKYLFELENDGVPEVILPR
SAHFSFKKIASMLSLKPVYVSLDDGYKMDVSMVEELITDNTMAIVAVAGTTELGMVDDIE
RISKIAYSKGIYLHVDAALGGFIIPFLENENNARLNFDFSCEGVCSITIDPHKMGLAPVP
AGGIIFRHKEHLNKLAVETPYLTHDKQTTIVGTRTGAATAATWTLLNHYGMEGYRKTVKK
VMELTRYTYERLSKINTVRVICKPELNLISFTPTNMEVNALKKELLAYGWHVSVAEHPHA
IRLVLMPHVKREHMDSFLKDLEKIMKK
>A_Q8PXA5
MNEQGLSEREIFSYLENAKSEDTDYYRVFSSMCTRPHKIAIEANRLFIEANLGDLGLFAG
AHKLEQEVVRMLGNLLHASSIDVPSGGLCQSSVCGYLTTGGTESNIQAVRGMKNLVTAGK
KEFKGTPNIVIPASAHFSFDKVADMMGIEVRRASLDSEFRVDMASVEKLINENTIGLVGI
AGNTEFGQIDPIDKLSEVALENELFLHVDAAFGGFVIPFLEKPQPFDFKVPGVTSIAIDP
HKMGLSTIPSGALLFRSPSFLDSLKVSTPYLTTKSQFTLTGTRSGASAAATCAVMKYLGY
EGYRKNVQYCMELTSKIVEEARKLGFEPLIEPVMNVVALKVPNPDLVRERLLKKFGWNVS
ITRTPRALRLVLMPHNSPEDIELFLEDLKKVTAEIKSP
>A_A0A1N6W5S0
MNKPPEERPASSAKSSAESTDSADGEDSSDSMPSEESLFLGSEKGAETYRTTMEQTTDAI

LDAFVENADPYSGTSPESLREEFAEMEMIPDSGEGLESALESAEPVLRNSVGVSDRQCLA
HLHCPPMISGLAAEAMLSATNQSMDSWDQSPAATHLETRMVEELCDLFGYGDSGDGVFTS
GGTQSNFMGLLLARERFAKERFGTNVQRSGLPHRAKAMRILCSEEAHFTAEQAAAHLGLG
ENAVVTVESNDDREMCPDALDQTLAELDERELLPFALVGTAGTTDFGSIDPLDELAERAE
EHDLWFHVDAAYGGALALSDRHRDLLSGIDRADSLSVDFHKLFYQPISCGAFLLRDGSQY
EHIARNASYLNPEGASVPNLVAKSAQTTRRFDALKPFLSFRALGRDGFGMLVDETIALAE
EVAELLASGSSFELVAEPTINAVVFRYRPTSDMADERLSWLNEAIRESLLREGDAVVART
EVDGVTALKFTLLNPRTTLTDVADILDAIERRGSSLRAVSPEVKR
>A_A0A1N6VH17
MQVAAPQSFRRVLSSMCTEPHPAARDAAERFLASNPGDPTTYPTVSALEDEAVGILGEMT
GLENPHGYIASGGTEANIQAVRAARNLAEAESGARTDSPNIVAPESAHFSFQKAADVLGV
ELRLAEVDANRRAEPDAVAELVDDDTVLVVGIAGTTEYGRVDPIPTLSEIAHDAGALLHV
DAAWGGFVLPFTDYEWNFDHAEVDTMGIDPHKMGQAAVPAGGFLAREKRVLDALAVETPY
LESTSQATLTGTRSGAGVASAWAAMDELWPAGYREQYERSQANAEWIAERFEACGYDVVD
PVLPLVAADVPRKTVEELQSLGWRVSPTGSGELRIVCMPHVTRSMLESFAADLESL
>A_A6UVR4
MDERAVLEELKKYRKMDLKYEDGAILGSMCTKPHPITKKISDMFFETNLGDPGLFRGTKK
LEDEVINNIGKFLNNPNPFGYIISGGTEANITAMRAINNIAKAKRKNHKTTVIMPETAHF
SFEKAREMMDLNLITPPLTKYYTMDLKYINDFIEDRNNKNDISVDGIVGIAGCTELGAID
NIKELSKIAEQNNIFLHVDAAFGGFVIPFLDDKYKLDNYCYEFDFSLNGVKSMTVDPHKM
GLAPIPAGGILFRDKSFKKYLDVEAPYLTDIHQATIIGTRSGVGVASTWGVMKLFGEEGY
KNLASECMDKTHYLVKEAKKLGFKPVIDPVLNIVALEDDNPEETSLKLRKMGWFISICKC
VKALRIIVMPHVEKEHIDKFLGALTEVKKN
>A_F4BYV2
MIMKETDVMPGKSAEESLDPEDWESMRMLGHRILDDMMDYLETLRDRPAWQHAPVDVKAH
FAGSPPVIPQTPEEIYQEFTQYILPYQIGNSHPRFWGWVAGTGTVMGMFAELISAATDAV
SGSFSYLSNNYVEMQVLDWCKTMLGYPATASGLITSGCSASNLIGLAVARNAKAQFDLRS
KGMRAAPQLMTLYCSEEAHSSIQKAVELLGFGSRALRRVPVNESMQIDLESLKEAIKTDR
EGGYHPICVVGVAGTTNTGAIDDLEALAEICSKEGLWLHVDGAFGAWAAIAPRSKHLVAG
IERADSLAFDLHKWMYLSYPIGCVFIRDADEHRRTFSLTPTYLAHGEGERGLTGIDVPWL
SDYGFELSRGFQALKAWMTIKEQGTEKYGRLIQQNIDQAHYLASLVEKSPQLEMALPVSL
NVVCFRYIRSNMSDSMLDLLNKQIEVELQEKGIAVPSIVTIKGKKYLHAAITNHRSLQSD
FDLLAREVVRIGDELG
>A_F4C0R3
MKHRDQIRQIAKKVNKSGQRSEMREKALPADEVMAILERTRERDYSYDRFLSTMCTRPHP
IAIKAHDMFLETNLGDPGLFPGVAGLEEEVVRMLGELLGCPLARGYISTGGTESNIQAIR
AAKNESGKCGGNIVVPASAHFSFDKIGDLLSLEVRKAELDSQLRVDLSSVESLIDEHTAA
LVGIAGTTEFGQVDPIEELSDLALEWGVHLHVDAAFGGFVLPFLDRSFAWDFSLPGVKSI
TIDPHKMGLATIPAGGLLFRNQECMNALETETHYLTKARQASLTGTRSGAAAAATYAVMM
HLGREGFREMVGYCMDLTDHLVRGAKEIGVEPLIEPVMNVVALRVPEPSKVRERLMDRDW
HVSITREPNRALRLILMGHMSHENVDLFLKDLKEVLSEFV
>A_D3RZ59
MVFKELLNFRLKDISYKRVLSSMCTVPHPLAVKAHIMFLETNLGDPGIFVGTWELERELI
KMLGKLLHNEKAAGYICSGGTEANIQGIRAARNLKRAKKPNIVIPKSAHFSFEKIGDLLA
VEIRRVGLDEEYRVDVGEVEKAIDENTVAIVGIAGTTELGQVDPIDELSKIAIEKDVPLH
VDAAFGGLVLPFLERKIPFDFELEGVTSITLDPHKMGMATIPAGGILFRDESFLKLLEVE
TPYLTTKYQFTLTGTRPGTGVASSYAVLKGLGFEGMKRIVKKCMENTNYLVEKMGEIGYE
PVIEPIMNVVAFKTERAEKIKEELYKRGWVISTIREPKAIRMVVMPHVTKEMIDEFVEEL
KKI
>A_L0JPS5
MTGGDLTGRHRAATDGPTPPDTASAFLGDPDGNAAYAAAIDRARDCLLESFATVDGPYAG
TDHETLRARIDELTVVPETGDSLAAVLETVAEEVLTDSVRVHDPDCVAHLHCPPAIPALA
AELLLSGTNQSLDSFDQAPAASVLEERVVDACCDLFDYPADADGVFTGGGTESNLLGLLL
ARDWYCQTRFDRDVQSAGLPPEAADLRLLCSDAAHFTADQAAHHLGLGEDAVVTVPTDDD
RRMNLAALDETLEALAADGRRPFAIVATAGTTDFGSIDPLAPLADRAAEHDLWLHVDAAY
GGACAISDSLRPKLAGIDRADSIAVDFHKLFYQPISCGAFLLRDGDRYRFLERNAAYLNP
ERDDAAGVPNLVSKSPRTTRRFDALKPFVTFNALGRAGVADCVEYVCELADAVADEIRAE
PALELCCDPELSAVVFRYRPSDRGARAGESDDRPGSASRPALDRLNRGIRDELFADGEAI
LARTEVDGTAALKLTLLNPKTTLSDLQAVLAAVVDRGEALEAETDST
>A_L0JRU9
MQIEPQAFDRVLSSMCTEPHPAARDAAERFLATNPGDPGTYPGVSELEEDAIALLSEIAG
LQEPAGYITSGGTEANIQAVRIARERADSRNPNVVMPESGHFSFQKAADLLGVDLRIVPT
DDDYRADLEAVRAAVDEDTAAVIGVAGTTEYGRVDPIPELGEIARSVDATMHVDAAWGGF
VLPFTDYEWNFDHAAVDTMAIDPHKMGQAAVPAGGLLVRDSALLDELAVDTPYLESTSQA
TLTGTRSGAGVASAVAAMEELWPTGYRRQYVRSQNNAEWLADALEKRGYEVADPTLPLVA
ADVPRSTFDALRAKGWRISRTATDELRVVCMPHVTREMLASFIGDLDRLEVRASVPITSD
D
>A_A0A1I0QDE1
MTGGLGQASRSRADEPTPPAAASAFLGDPEGNAAYADAIELAREVLVESFATAEGPYAGT
DHETLRERLADLPVVPDEGESLDAVLETVADEVLDDSVRVHDPDCVAHLHCPPTIPALAA
EVVLSGTNQSLDSFDQAPAASVLEERVVDACCDLFDYPAGADGVFTGGGTESNLLGLLLA
RDWYCERRFDRDVQAAGLPPEAADLRLLCSEAAHFTADQAAHHLGLGEDAVVSVPTDGDR
RMDVSALDETLDRLAAEGRHPFAIVGTAGTTDFGSIDPVDALADRAAERDLWLHVDAAYG
GACAISDRLRPKLAGIDRADSIAVDFHKLFYQPIGCGAFLLRDGDRYRLLERNAAYLNPE
RDDAAGVPNLVSKSPRTTRRFDALKPFVTFNALGRTGLADCVEYVCDLADAAADEIRAEP
ALELCCEPELSAVVFRYRPDRTDGCDRTADRADRADRTDCSSGEVVGRVNRAIRDELFAD
GEALLARTTVDGTPALKFTLLNPRTTLSDLRDTLAAVVDRGEALEREVIDSA

>A_A0A1I0Q4I5
MQAEPQAFDRVLSSMCTDPHPAARDAAERFLATNPGDPGTYPNVAALEDDAIELMGEIAG
LANPSGYITSGGTEANIQAVRIARERADARTPNVVMPESGHFSFQKAADLLGVELRIVPT
DDRYRADLDAVRAAVDDDTAAVIGVAGTTEYGRVDPIPELGEIARSVDATLHVDAAWGGF
VLPFTDYEWNFEHAPVDTMAIDPHKMGQAAVPAGGLLVRSADLLDELAVDTPYLESTSQA
TLTGTRSGAGVASAVAAMEELWPGGYRRQYVRSQNNAEWLADALEKRGYQVAEPTLPLVA
ADVPRSTFDALRAKGWRISRTATDELRIVCMPHVTREMLASFVGDLDRLEVRASVPVASD
D
>A_A0A1J1ACR1
MTHVSASSEPQSFDRVLSSMCTEPAPAARRAAMAFLGSNPGDPATFQTIAARERETVSML
GELVGLADPHGYVTAGGSEANIQAVRAARNRATVAEPNVVAPTSAHFSLRKAASLLDVEL
RLVETGPDHRADPKAMADAVDERTALVFGVAGSTEYGRVDPIPALVDLADSVDAMLHVDA
AFGGFFLPFTDREWHFGHAGIDSMTIDPHKAGRAAIPAGGFLARDRSVLDALSIQTPYLE
SESQVSLGGTRSGAGVASAHAALETLWPDGYRAAFERTMELARWLATELDSRGFDVIEPE
LPLVAWEASQSLFERLRDAGWRIARTQQGAIRIVVMPHVDRNMLDAFLTAVDRHRP
>A_A0A328SS11
MFRKRHDKRYIFEKLEAFHQMDMTYDSGRILSSMCTKPDDIALEAFRMFIETNLGDGGLF
KGTSMMEEEVIASLAKLLHSDDACGHIVTGGTEANIMAMTVAKYLFQEDNDNTPELILPR
TAHFSFKKACSMLSLNTVEVPLNDKHKIDIERLEDCITDNTMAIVAIAGSTEFGLVDDIG
EISKIAKSNDVYLHVDAALGGFIIPFLNYRNNTQLNFDFKCKGVSSITLDPHKMGLAPVP
AGGILFRHKKYLDKLAIDAPYLTKNVQTTIVGTRTGATTAAAWALINYYGMDGYADIVEE
SINLTRYTYNRLSQMNHVQVIVKPELNVIAFNVSDMKVNELKEKLFKKGWRVSNTVNPYA
VRLVLMPHVKKQHIDRFLVDLEEIIREYYS
>A_D3SYW6
MQREPQAFDRVLSSMCTTPHPVAREAAERFLATNPGDPGTYPTISALEDEAIELLGEVAG
LDDPAGYVASGGTEANIQAVRIARERARSTAATAETPTVVMPQSGHFSFQKAANVLGVDL
ELVPTDDEHRVDLEAVRACVDETTAMVVGVAGTTEYGRVDPIPELAEIAQSVDALLHVDA
AWGGFVLPFTDHAWHFDHAPVDTMAIDPHKMGQAAVPAGGLLVRDETLLDELAVDTPYLE
STSQATLTGTRSGAGVASAVAAMEELWPDGYRDQYVRSQNNAEWLAAELATRGYDVVDPE
LPLVAANVPEATFEALRDAGWRISSTGSGELRVVCMPHVTRTQLESFVAALDALESESET
PTATT
>A_A0A0E3P0J0
MNEQGLSEKEVLSLLKKAKSEDTDYYRVLSSMCTHPHAIAAEAHMLFIEANLGDLGLFPG
TYGFEKEVISMLGELLHAPSLRNSGKFSDGEFCGYLTTGGTESNIQAIRAMKNLSAHRKP
QITNPNIVIPDSAHFSFDKIANMLGIEIRRAGLDPDFRVDLASVEKLTDKNTIGLVGIAG
NTEFGQVDPIEGLSELVLEKDLFLHIDAAFGGFVIPFLEKSYPFDFEVPGVTSVAIDPHK
MGLSTIPSGALLFRSHSFLDSLQVKTPYLTTKSQFTLTGTRSGASVAATYAVMKHLGREG
YRKNVEYCMELTEKLVKGARKLGFYPLLTPVMNVVALGVPEPDLVREQLHEKFGWNVSIV
RSPRALRLVLMPHTTTQDIEEFLKALEKVVAEL
>A_Q8TUQ9
MNEQGLSEKEIFSYLENAKSEDTDYYRVLSSMCTHPHKIAVEANRLFIEANLGDLGLFAG
ASRLEQEVVGMLGELLHAPSIDVPFGGSCESSACGYLTTGGTESNIQAVRGMKNLVTTGK
KELKGAPNIVIPESAHFSFDKVADMMGIEVRRASLDSEFRVDMASIESLIDANTIGLIGI
AGNTEFGQIDPIDKLSEIALENELFLHIDAAFGGFVIPFLEKPQPFDFKLPGVTSIAVDP
HKMGLSTIPSGALLFRSASFLDSLKVNTPYLTTKAQFTLTGTRSGASAAATCAVMKYLGN
EGYRKNVQYCMQLTEKLVIEARKIGFEPLLEPVMNVVALKVPNPDFVREQMLERFGWNVS
ITRTPRALRLVLMPHNTLEDIEIFVQDLKEVTVEI
>A_R9SLT3
MDGKPKSKEEIFEELEQYQKKDMKYSDGRILGSMCTEAEPIAKEVFYKFINSNLGDPGLF
PGTKAIEDKAIKMIGSLVSIDNPYGHIVTGGTEANLMAMRAARNYARRYKNITEPEMIVP
KSAHFSFKKAADMFGMKVVEADMDGYLIDVNSLENKINKNTTVIVAIAGTTELGLIDNVE
EIAKIAKKHDIYLHVDAAFGGFVIPFLREEGYDFPKFDFSLDAVCSMTIDPHKMGLSVIP
SGCILFRDKKYLDVMAVKAPYLTKKEQSTIVGTRSGASSAATLAVMESLGREGYRKLALD
VMDKTMMLKEGLEDIGYDVVVEPQLNIVAFYHKDIDTDYLADLLEQRGWRVSTSSYPKAI
RVIVMKHISRDNIRDLLVDLKAISSNI
>A_A0A2Z4L867
MFKKTHDQKYIFEKLESFHQMDMTYDSGRILGSMCTKPDPIALEAFRMFTETNLGDGGLF
KGTSMMEDEVISSLSRLLHSSEACGHIVTGGTEANIMAMCAAKYIFQEDNEDTPEVILPR
TAHFSFKKACSMLSLKTVEVPLNDEYKIDTTALEDCITDNTMAIVAIAGSTEFGLVDDIH
EISKIAHSNDVYLHVDAALGGFIIPFLNYRNKTRLNFDFKCKGVSSITLDPHKMGLAPVP
AGGILFREKKYLDKLAIDAPYLTKNVQTTIVGTRTGATTAATWALINYYGMDGYADIVEG
SINLTGYTYNKLRQMKNVNVIVKPELNVIAFNVDDMKVNVLKEELFKKGWRVSNTVKPYA
IRLVLMPHVKREHIDEFLSDLEEIIGGS
>A_G2MMT9
MNRSNTPQAAFIDPSGGNAEAVRDFAEDVLDQLLGQLGTADDRSPLPEVSVIPSVMFPDS
SRPHGDLLTDLETIVTGSMNPAHPGYIGHMDTMPTTVSVLGDLVTSAVNNNMLSVEMSPV
FSELEVQLVERIADEFGLGPGAGGVLCSGGSLANLHALSVARNQAFTVHKEGLASVDRTP
VLFASEVAHTSLQKAAMLLGLGADAVIPVETDDDSRMAPTALAQAIETAEREDQAPFCVV
ATAGTTTTGNIDPLPAVRDVADAHDLWFHVDAAYGGALVFSEAERGRLDGIEAADSVTFN
PQKWCYVAKTCAMALFADGDVLQEDFRIGAPYMRGDDAIPNLGELSVQGTRRADILKLWL
TFQHLGRNGLEQLIDESYRLTAVIHEHVVEHDALECASRPEMNLLCFRAVPEWCPPEGRD
ELNNRLQQTLLSEHDIFVSLPTYRDSRWLRVVLLNPFTGEETIQRLFSGIDAFLTAGKTG
SN
>A_Q9HHV3
MEAACDAVLDVIGGAHGPYSGASYDRLRDRLADTTALPAEGHPLEDVLAAVREDVLANAV
HPSDETCVAHLQCPPMVPALAAEALLTASNQSLDSFDQAPAATVVEERLIADLTALYELG
PAADGVVTGGGTESNHQALLLARDDYVETVFGQSVREHGLPPAAQDLRILCSADAHFTAA

QSAALLGLGEDAVVTIPTDGRHRMDATALRETVERLNDAGKHPFAVVGTAGTTDFGSIDP
LSAVADVAATHDLWVHVDAAFGGALAVSDQHSERLSGIDRADSVAVDFHKLFFQPIACGA
LLVADGESFELMSRNAAYLNPAGDAVPNLVAKSTRTTRRFDALKPYVAFRTLGRDGLGAL
VDRTLALADDVAGLLRADPAFELACEPTLNAVTFRYRPVREHAHCDPGPWADHVTEAARE
QLFDAGTAVVARTTVDDRAHVKFTLMNPRTTVADIQSVLVAFKGHAATIEAADRTAPITT
AAGGDQ
>A_Q9HSA3
MTRGEARRPPQEFDRVLSSMCTTPHPAAREAAQAFLATNPGDPETYPAVAERERDAVALL
GEIVGLSSPHGYIAAGGTEANLQAVRAARNRADADAVNVVAPASAHFSFQKAADVLGVEL
RLAPTDGDHRADVAAVADLVDGDTAVVVGVAGTTEYGRVDPIPALADIAAGVDANLHVDA
AWGGFVLPFTDHDWSFADAPVNTMAIDPHKMGQAPVPAGGFLARDPETLDALAIETPYLE
SDTQPTLGGTRSGAGVAGALASLRALWPDGYREQYERTQGNAEYLAAELAARGYDVVDPE
LPLVAADMPDAEFQALREEGWRISRTASDALRVVCMPHVTREMLAAFLDDVDALA
>A_A0A2V3JDE0
MSMSREELEHLLAEKKRRDLSYSRILSSMCTYPHEVAVYAHKLFIESNLGDSGLFQGTKE
MEDEVIRTIGGLLGDENAYGYITTGGTESNIQAIRAVRNRKRKEGLRVSDMNIIVPETAH
FSFDKIADILGIEVRKAGLDQQLRVNPDSVQEFMDDGTICLVGIAGTTEFGQVDPIRELA
EIAKEKNVFLHVDAAFGGFVIPFLPDRARYEFDFSLEGVSSISIDPHKMGMSTIPAGCLL
FREESYLEELAVPTPYLTTKEQYSLTGTRSGASAAATFAVLKYLGSEGMKSIVDECMRLT
RFLVDGAREMGIEPVIEPVMNVVTLQLGDADRIASALRAKGWEVSTTRSPKSLRLVIMPH
VTEDMLRRFMEDLSEVVN
>A_Q6L2R7
MLKRFPERGIPLDEINSILDGYGKNDIKNSRGRLFTYFYDPGLKDLDDLSSILLKFYNRN
GMDYHAFPSTLKIENDLISMMSDLMHGNDDTSGTFTTGGTESILLAMKAARDLFLEKKEY
VPEIVAPVTAHPAFSKAAKYLGMKITRVPVNEDYIADDTINEYINDRTAAVIASAPSFPY
GGIDNIKDISEIALDKNTWFHVDACVGGMILPFLKGLGLNIKDFDFKLPGVSSMSIDLHK
YGFTPKGSSVVLYKNHDLRKRQIYVNADWPGYPMSNMGMQATKSAGPLAGSWATLNYLGL
DGYKKLAEKTLKAYRMIRSGITDLGYKIIGRPDATIFAFTHNDKDIIDLGIKMIENGWYP
QIQPGNVFIDMPDSVHLNISPVHLDVADEFLEFFNELDKNVKPRDKNSYNVNSVEKALEM
IKMEKSMLFRIIRYSRPEVSEKIFLEMTDEDFTYS
>A_A0A1H9J813
MTSNELAGQVRSADEPTPPAAASAFLGSADGNAAYADAIDLARECLLESFATVEGPYAGT
DHERLRERIDDLAVFPAEGEALSDTLETVADEVLADSVRVHDPGCVAHLHCPPAIPALAA
AVLLSGTNQSMDSFDQAPAASVLEERVVDACCELFEYPAGADGVFTGGGTESNFLGLLLA
RDWYCETRFDRSVQTAGLGPAAGDLRVCCSEAAHFTAEQAAHHLGLGEDAVVTVPTDDDR
RIDLAALDDTLERLEAAGRHPFAIVGTAGTTDFGSIDPLEGLADRAADRGLWLHVDAAYG
GACAISDRLRPKLAGIDRADSIAVDFHKLFYQPISCGAFLLRDGDRYRFLERNAAYLNPE
RDDAAGVPNLVSKSTRTTRRFDALKPFVTFNALGRTGVADCVEYVCELADAVAAAIRAEP
ALELCCEPELSTVVFRYRPDRPAHSDDRTEPLPAAAIDRVNRAVRDELLADGEVLLARTT
VDGAAALKFTLLNPRTTRSDLDAALAAVVDRGEALEREVIDSV
>A_A0A1H9Q7I0
MQSEPQAFDRVLSSMCTDPHPAARDAAERFLATNPGDPGTYPTVATLEDDAIEQLGEIAG
LDDPAGYITSGGTEANIQAVRIARERAGTRTPNVVMPESGHFSFRKAADLLGVELRVVPT
DDDHRTALEAVRASVDDDTAAVIGVAGTTEYGRVDPIPELGEIARSVDALLHVDAAWGGF
VLPFTDYDWQFAHAPVDTMTIDPHKMGQAAVPSGGLLVRSPDLLDALAVDTPYLESTAQA
TLTGTRSGAGVASAVAAMDELWPDGYRRQYVRSQNNADWLADALEKRGYEVADPTLPLVA
ADVPRSTFDALRAKGWRISRTGTGELRIVCMPHVTREMLASFVGDLDRLEVRASVPVASD
D
>A_E1RFW3
MLEDGLREDELFRHLSSIKEKDRSYRKVLSSMCSIPHPVAVRAHNIFIESNLGDPGLFMG
TASLEAELIERLGSLMSLPEACGYATSGGTESNIQALRIARENAGKKSPNVIIPESAHFS
FEKACDILSIEMRQAPSTEKYIVDTERMEDLIDGNTIGMVGVAGTTEYGTVDPIEHLSDI
ALDRDLFLHVDAAFGGLVLPFIKGSPPFDFRLDGVSSISVDPHKMGMSTIPCGCIMVRNP
DFFRSTEVDTPYLTVKKECTLCGTRPGGPVAGALAVLDHLGRKGMIEVVERCMENTRFLI
RGMEELGHPVAVQPSVNVASFSCDETPDGWIVSRTRHGHMRTVCMPHITRETLEEFLKDV
GEM
>A_A0A328S773
MLDKGMDKEKLFEKLFDYKKMDLDYASGKILGSMCTKPDPIALEAYKLFIETNLGDPGLF
TGTALMEREVIGVLGELLHLFNPSGHIVTGGTEANLMAMAVAKKLFLENHEGVPEVILPE
SAHFSFKKITSMLSLKPVYVPLNDEYKTDVSVIESLINDNTMAIVAIAGTTELGLVDDIE
KISEIAYKNNIYLHIDAAFGGFIIPFLEYDNEHALNFDFKCKGVSSITIDPHKMGLAPVP
AGGIIFRYSEHLEKLAVETPYLTKDKQTTIVGTRSGAGTAAIWTLLNYYGKEGYKKAVTQ
AIKLTNYTYEKLLDMENINVTCKPELNIISFTSNIKSPKQVKQELYEYGWRISVSANPYA
MRLVLMPHVKQEHIDNFLKDLNEILKRE
>A_A0A0F7IJ10
MDIIDELRIYRERDIPYSRVLSSMCTTPHPVALEAHRMFVETNLGDPGIFVGTTELERKV
IEMLGSLLNHPKAAGYISSGGTEANIQGIRAARNLKRVKNPNIVIPKSAHFSFEKIGDIL
GVEIRRAALDESYRVDTAEVERMVDENTVAIVGIAGTTELGQVDDIEELGRIARDMNVFL
HVDAAFGGLVLPFMDERIPFDFGVEGVSSITVDPHKMGMATIPAGGILFRGEEFLRALEV
ETPYLTSKYQYTLTGTRPGTGVASTYAVLKHLGFEGMKEIVERCLRNTRILVEEMEGIGF
EPVIRPIMNVVSFHAENAEKIKDELYRRRWVISTIRVPKAVRMVIMPHVTEEIIKEFISE
LRQVVRVV
>A_A0A165Z3S9
MEDKPVSRKEILKELEEIQKLDCKYSDGRILGSMCTEPHPFAKEVFCKFLDSNLGDPGLF
KGSKYIENKVIQSLGKLLSLDKSYGNIVTGGTEANIMAMRAARNYAMKYRGIVDGEIIIP
ESAHFSFKKAADMLNLKIIEVNLDDNFKIDVESLKNLISSKTVAIVAIAGTTELGLVDPI
EKIAKIANDNHIYFHVDAAFGGFSIPFLKELGYDFPDFDFSLPGVCSITVDPHKMGLAPI

PAGGIIFRKKEFLDVMAVDSPYLTVKTQSTIVGTRLGASSLAAYAIMKYFGKEGYCKIAS
ETMDKTKFLKEGLEEIGYDVVCEPELNLVAFNHPNKNAHELANELEELNWKVSVAKCPIS
IRVVLMNHIKKNHLKEFLEDLKEIY
>A_D3DZR8
MNKEPISEEEIFKELDFYQSQDCKYSDGRILGSMCTQAHPIAQKAFIQFLESNLGDPGLF
KGTKAIEDKVLKMIGSFLSIENPVGHIVTGGTEANIMAIRAARNIARDEKGISQGEIIVP
QSAHFSFKKASDILNLKLREIVLDDSYQLDASFVEDEINENTVAIVGVAGTTELGMIDPI
EELSNIALENNIHLHVDAAFGGFSIPFLKEIGYGLPEFDFSLKGVKSITVDPHKMGLAPI
PAGGILFRNEEYLDSISVNSPYLTIKHQSTIVGTRMGATSAATFAVMKYLGKDGYARLAK
ESLDNAIFLAESVKQLGYELVVEPKLNIVAFNHPKLETDDLAQLIEKRDWKVSCSSCPKA
IRVILMNHIKREHIVELISDLKDISESI
>A_A0A1M5JPP3
MGEPRPGDDTVADAEDGEGDDEVHRSADGDGSAGRGRALAEGLFLGSDAGNEAYAEAMTT
ARDAVLEVVGDPDGPYSGATYADLHERLDGATLPETGAPLDDVLADVRDGVLADAVYPSD
EACVAHLQCPPTVPSLAAEAMLAAVNQSLDSFDQAPAATVVEERLVADLTELFDLGPDAD
GVLTGGGTESNHQGLLLARDRYVAETFDRSVREAGLPPEAGDMRVLCSADAHFTAAQSAA
LLGLGEDAVVEVPTDDRRRMDPDALRAELDRLEEAGERPFALVATAGTTDFGSVDPLAEL
ADVADARDLWLHVDAAFGGALAVSDDHRDVLAGIGRADSLAVDFHKLFFQPIACGAFLLG
DGADFRYMSRNAAYLNPEADEVPNLVGKSTRTTRRFDALKPYVAFRTLGREGLAALVDRT
VALADRVAGEIRADPAFELACEPTINAVTFRYTPLRDRPDRDPGAWADRVNREARDRLFE
AGRGVVARTEVDGRAHLKFTLMNPRTTVDDVRDLLVALKGHAAAVEAEGAEQTDGAGLTD
REGTEATRSDRRRDGDAPPGATADATGVDR
>A_A0A1G9UVH9
MTVDDTFLGSRGGDEAYRAAMQRTTEAVLAQVADRERPYSGLSPAELADAVAEPVLPEEG
VGLPDAIDDVTAAVLPHSVATSHPRCAAHLQCPPMVPGLAAEALLTATNQSLDSFDQAPA
ATVLEERVVDALCDLFGLPEEGDGVFTSGGTQSNLQALLLARDRYCARVFGRNVQSAGLP
AEADALRVVCSEAAHFTAQQAAHHLGLGEDAVVTVPTDDRQRMDPDALAATLADLEAMSR
QPFAVVGTAGTTDFGSIDPLDALAERAAAHDCWFHVDAAYGGAVALTDEYASLLDGIEWA
DSIAVDFHKLFYQPISCGAFLLRDGDDFEWMARNAAYLNPEAHDDGGVPNLVAKSTQTTR
RFDALKPYLAFRALGREGMATLVGRTLELADEAAALLDAAEDFERLHDPTLNALVFRYRP
REGMADAAVGRLNAAVRERLLEDGRALVARTEVDGVQSLKWTLLNPTCTVEDLAATLDVL
RECGADALAGEVSA
>A_I7CDV7
MTSDGLVGRFRSAPDEPTPPAAADAFLGDPAGNAAYEDAIDRARDCLVDSFATADGPYAG
TDHETLRERIDELTVFPDEGDPLAAVLETVGEDVLADSVRVHDPGCVAHLHCPPAIPALA
AELLLSGTNQSLDSFDQAPAASVLEERVVDACCDLFEYPAGADGVFTGGGTESNFLGLLL
ARDWYCQTRFDRDVQAAGLPPEATDLRLCCSAAAHFTAEQAAHHLGLGEDAVVTVPTDDE
RRLDLEALDETLARLESEGRHPFALVGTAGTTDFGSIDPLEELADRAAERDLWFHVDAAY
GGACAISDRLRPKLAGIDRADSIGVDFHKLFYQPIGCGAFLLRDGDRYRFLERNATYLNP
ERDDAAGVPNLVSKSTRTTRRFDALKPFVTFNALGRTGVADCIEYVCDLADAVAADIRAE
PALELCCEPALSAVVFRYRPARSDAAARAAGSDERPESASTAAIDRVNRAIRDELLADGE
VVLARTEVDGTAALKLTLLNPKTTRWDLRAALQAVIDYGETIERDREAIDST
>A_D4GP47
MNGVGDVDGERRDPRPDAAKWFLSGDDDDRARYRDAMRRACDLVLDEFAAEATPYSGATP
EEVDDALARFEMLPHEGDGVAAALDRTEPILRNSVGVSDPTCIAHLQCPPTIPALAAEAL
LTATNQSMDSWDQSPAATQLERRFVGELCDLFGYEDGDGVFTSGGTQSNFVGLLLARNKV
VLEEYGVDVQHEGLPPEARDLRILCSADAHFTATQAASHLGLGENAVVTVPTDDDRRLSM
AAFDEAVADLRERGKRPFAIVATAGKTDFGSIDPLGPLAERAAELDCWYHVDAAWGGALA
LSDDHADKLAGIEAADSVAVDFHKLFYQPISCGAVLVRDASAYDLIDRNAAYLNPVRDDD
AGVPNLVSKSVQTTRRFDALKPFVTMQALGREGLASMVEYTIDLAADAARLVEADPDLRL
VHDSDLNVVVFRYVPDHGRDGGGPADGPVNGPSADDLNEAIRDSLLDDGEAVVARTTVDG
ETCLKLTLLNPRTTREDLRDLLREISDRGTALEPTTDDRHA
>A_A0A0F7PCV2
MTALGSAARPEPQSFERVLSSMCTEPHPAAREAATHFLATNPGDPDTYPAVAALEREVVS
MLGDVVDHQDPTGYVATGGTEANIQAVRAARNLAATDDPNVVGPESLHFSFQKAADVLDV
DLRLAPVDEDYRADVDAIADRIDDDTVLVVGIAGTTEYGRVDPIPALSDVARDADARFHV
DAAWGGFALPFTDEQWNFAHADIDTMTIDPHKLGQAAIPAGGFIATDERTLDALAVDTPY
LESTGQATLTGTRSGAGVASAHAALSAQWPDGYRENYETGMNLATWFAEEMRVRGYDVVS
PHHPIVAVDIPTGDFEALTEAGWRLARTSAGELRVVMMPHVTRDSLAGFLADVDDLASSG
EER
>A_F8D376
MQTEPQAFDRVLSSMCTEPHPVAREAAERFLATNPGDPGTYPTVSALEDEAIAMLGEIAG
LEEPSGYIAGGGTEANIQAVRIARERADATRPNVVMPESAHFSFRKAADLLGVDLRVVPT
DDRYRADLGAVRAAVDDDTAAVIGVAGSTEYGRVDPIPELGEIARSVDATLHVDAAWGGF
VLPFTDYEWNFSHAPVDTMAIDPHKMGQAAVPAGGLLVRDSALLDELAVDTPYLESTSQA
TLTGTRSGAGVASAVAAMEELWPSGYRSQYVRSRNNAEWLADALEKRGYDVVDPTLPLVA
ADVPRSTFDALRAKGWRISRTATDELRIVCMPHVTREMLASFVGDLDRLEVRASVPVACD
D
>A_Q3IT46
MQRAEPQEFSRVLSSMCTEPHPAAREAAERFLATNPGDPGTYETVSKLEREAVDMLGEVA
GLPDAAGYIASGGTEANIQAVRIARNRADTRTPNFVAPASAHFSFRKAADILGVELRTAP
LEDYRANLDGVAELIDSDTALVVGVAGTTEYGRVDPIPALADMAADAGALCHVDAAWGGF
VLPFTEHAWDFDDADIHTMTIDPHKMGQAAVPAGGLLARGPELLDELAIDTPYLESTSQV
TLTGTRSGAGVASAAAVMDELWRDGYRQQYETAQTNAHWLAAEVESRGFDVVDPVLPIVA
MDLPYDLVADLRERGWRLSRTEADEARIVCMPHVTRSMLEEFLTDLDRLA
>A_S0AU28
MLKQFPENGMDIQKIHETLDELGKNDIKNSRGRLFTYFYDPGIDELNKLQDIFLKFSNRN

GMDYHAFPSTLKLENDVIAMMASLLHGKEGSAGTFTTGGTESIILAMKAARDRFFEKHHG
VPEVILPVTAHPSFSKAVEYLGLKEIRLPVDEHYLADPELMRKAITENTAMIVGSAPSFP
YGTIDPVKELSDIALENNLWLHVDACVGGMILPFLKRLGHNVQDFDFTLPGVSSISVDLH
KYGFTPKGSSVIMYKNEELRKHQIYVNAKWPGYPMSNAGMQATKSAGPLAGTWSIMNYLG
YKGYTDLASKTLSAYKTLTKGIENIGYEITGKPDATIFAFQDNNNSIFTTGVNMIEKGWY
PQIQPSNLELGLPSTIHLNVCPVHVEVADEFLSDLENIHKNAGKDSGSRELAVESRDKAV
DYLVNLIEQNPEKKTLFFHMIYNLDPEKGEEIFRKITDMDFHAAEE
>A_F6BEM5
MEEKGISEREVLEALKKYREMDLKYENGRILGSMCTKPHPISKKIVEMFLETNLGDPGLF
KGTKKLEEEVIGMIGELLHNKNAFGYIITGGTEANLTAMRAIKNMKNKNAKIIIPETAHF
SFDKARDMMDLEFIKAPITKDYTIDVDFVRDYVEDYKVDGIVGIAGSTELGTIDNIEELS
KIAVENDIYLHVDAAFGGFVIPFLDERYKKKNINYKFDFSLEGVCSITIDPHKMGLSPIP
AGGILFRDKSFKKYLNIEAPYLTETQQATIVGTRAGFSVACTWGIMKLLGKEGYKKIVSE
CMENTIYLTKKAKEYGIESVIEPVMNIVALKDENPKETCSKLKKHGWYVSICKCVNALRI
VVMPHVKKEHIDEFIEVLVSLKS
>A_Q60358
MRNMQEKGVSEKEILEELKKYRSLDLKYEDGNIFGSMCSNVLPITRKIVDIFLETNLGDP
GLFKGTKLLEEKAVALLGSLLNNKDAYGHIVSGGTEANLMALRCIKNIWREKRRKGLSKN
EHPKIIVPITAHFSFEKGREMMDLEYIYAPIKEDYTIDEKFVKDAVEDYDVDGIIGIAGT
TELGTIDNIEELSKIAKENNIYIHVDAAFGGLVIPFLDDKYKKKGVNYKFDFSLGVDSIT
IDPHKMGHCPIPSGGILFKDIGYKRYLDVDAPYLTETRQATILGTRVGFGGACTYAVLRY
LGREGQRKIVNECMENTLYLYKKLKENNFKPVIEPILNIVAIEDEDYKEVCKKLRDRGIY
VSVCNCVKALRIVVMPHIKREHIDNFIEILNSIKRD
>A_A0A0U2V5X1
MNDLPIDKDEILKELDSIQNEDLKYSSGRILGSMCTEAHPFAKEVYTKFLDSNLGDPGLF
KGTKATEDKTIKIIGKLLNLDNAYGNIVTGGTEANIMAVRAARNHARKYKGIKNGEIILP
RSAHFSFKKAADMMNLKIIEADLDENYKIDVKSVKNKITDNTVAIVAIAGTTELGLIDPI
EEISQIAYENNIYFHVDAAFGGFSIPFLKDIGYDFPEFDFKLPGVCSITVDPHKMGLAPI
PAGGIIFRKKEYLEVMAVDSPYLTVKTQSTIVGTRLGASTVATYALLKYFGRSGYAELAD
QLMQNTLFLKESLEKIGYDVIVEPELNIVAFNHPKKSPDELSKELEKINGWKVSVANCPK
AIRIVLMNHVTKTHLEEFLNDLEELF
>A_W0JRZ3
MTEQYSPASDDRPSNGRALADRLFLGSDDGNRAYLGAVEQAAEAVVTTVGEADDPYTGRG
YRALRDHLDGETIPETGAPLSVVLDEVATDVLANSVVPSDEACVAHLQCPPMVPGLAAEM
LLTAVNQSMDSFDQAPAATVIEERVIDDLADLFSLGDAADGVMTSGGTQSNFQGLLLARN
RYVADRFDRSARANGLPPAATDMRVLCSEHAHFTAAQGAAHLGLGEDAVVSVPTDRTYRM
DPEALRTQLERMKRNGERPFALFATAGTTDFGSIDPLEELADIAAEHDLWYHVDAAYGGA
LAVSDEHRSAIAGIERADSLSVDFHKLFYQPISCGAFLLREGDDFDLMARHAAYLNPDGD
DAPHRVEKSTLTTRRFDALKPYIAFRTVGRKGLEALVDRSLSVATRTAELVRADDAYELV
CDPTLNVVTFRYQPSNDHPELADGEWSDRLNREIREFLFDAGEGIVARTTVEDRVTLKLT
LLNPRTTVDDIRTLLERGQRHAATVEAAELGTAPVETERGNAGGSTDGVER
>A_W0JV70
MNRSNSPEAGFIHPDGANAEAVRDLAEDVLDQLLEQLGAAEERSPLPDESTVPTVRIPAS
PRSQNDLLGDLETIVAGSMNPAHPGYIGHMDTMPTTVSVLGDLVASAVNNNMLSVEMSPV
FSELEVQLIETIASEFGLGPDAGGILASGGSLANLHALAVARNHAFDVHKGGLTGLDRKP
VLFASEVAHTSLQKAAMVLGLGTETVVAVETDADSRLKPSALKRAVERAERGGCVPFCVV
ATAGTTTTGNIDPLPAVRDIADEHDLWLHVDAAYGGALVFSEAERGRLDGIEAADSVTFN
PQKWCYVAKTCAMVLFADADVLQEDFRIGAPYMRGDDAIPNLGELSVQGTRRAEVLKLWL
TFQHLGREGLGQLIDESYRLTAVIRNRVAEHDALELASEPELNLVCFRAAPDWCPPDEQD
ALNGRLQRRLLSEQDIFVSLPTYRDNRWLRVVLLNPFTDETTLDRLFNGIDVFLDAERP
>A_W0JJ99
MQVEPQTFDRVLSSMCTEPHPDARTAAERFLATNPGDPGTYPTVTELEEEAVSMLGEIAG
LDSPAGYVASGGTEANIQAVRIARDRFESRTSNRVGDHTPNRNERSTPNVVMPESGHFSF
QKAANVLGVELRIVPTDDDHRADLEAVRHSVDDRTALVVGVAGTTEYGRVDPIPELGEIA
RSVDALFHVDAAWGGFVLPFTDYEWNFSHAPVDTMAIDPHKMGQAAVPAGGLLAREETLL
DELAIDTPYLESTSQATLTGTRSGAGVASAVAAMEALWPGGYQRQYARSQHNADWLADAL
VARGFEVVEPTLPLVAADVPVSTFDSLRREGWRISRTATGEMRIVCMPHVTRSMLEAFVA
DLDRLQCRASVPVVSDD
>A_I3TCG1
MTALEKNRVAGLLRDLYVLSSKTPRHELSILGSMTTPPDPLALYAFSVFSHTNLADIELF
PPLKDMYRDVLEFTATLYGSRKGYVTAGATESNIVALLVAREVHGRESSVVLAPDTVHLS
VEKGCWLLGCKLVKVPTGNKPVDPGLLEDYVRAHRPFAIVVTAGTTELGLVDPLREVARI
ASEYGIYLHVDAAYGGLIVPFLYEEGLLRDNVYFYPGVSSIAVDFHKFAAAPPPAGLILF
SSDEYLDKSCIEYSYTLSGRTCGILGTRPGGSLAGIWAVVKAVGAARLRERALWAYRVAA
DLYERISSLRGFEVVKPQTTIVAFRHRRVDSLALLRYLAERGLFVYKAPSIRGLRVVVMP
HFNEHLIGRFLDALDEVARNTPDA
>A_A0A346PMZ5
MTTVLDGDMQAEPQAFDRVLSSMCTKPHPVAREAAERFLATNPGDPGSYPTISALEDEAI
AVLGEIAGLEESAGYVASGGTEANIQAVRIARDRAETSQPNVVMPESGHFSFRKAADVLG
VELRIVPTDDDHRADVEAVRAAVDDETAMVVGVAGTTEYGRVDPIPELGEVASTVDALLH
VDAAWGGFVLPFTRYEWNFSHAPVDTMAIDPHKMGQAAVPAGGLLVRSSALLDELAVDTP
YLESTSQATLTGTRSGAGVASAVAAMDELWPTGYRRQYERAQHNADWLAEALEKRGYEVV
EPTLPLVAADLPRSTFEALRGRGWRISRTATDETRIVCMPHVTRKMLAGFVADLDRLEVR
ASVPVVADD
>A_D2RH62
MLEEFKSKDIPYSRVLSSMCTIPHPIAVKAHVEFINANLGDPAVFRGSAELEKEVVRMIG
ELLHHPNAKGYIASGGTEANIQAIRAFRNLKRVKKPNVVVPESAHFSFDKAGEILRVEIR

KAKLDGEFRVDVGDVERLIDDNTVGIVGIAGTTALGQIDPIEELSELALERDVFLHVDSA
FGGFVIPFLDLNVKFDFELEGVSSMTIDPHKMGLATIPAGCILFRDESFLKALAVKTPYL
ITEKQYSLTGTRPATGVASTYAVMKYLGFEGFRKVVRRCMEVTRYLVERMGELGFEPVIE
PIMNIVCFKCEKAFEIRNELYKRGWVVSAIRNPRALRFVVMPHVDFEVIDKFVEEMKNVL
RKV
>A_E3GX95
MMEKGLSEKQVLKELKNYKKLDCSYSSGKILGSMCTEPHPFAKKVYYEFLTTNLGDPGLF
RGTSILEKETIQMLSSLLNAEKAYGNIVTGGTEANLMAMRAARNISNIEKPEIIVPASAH
FSFNKASEILNLKLKIAKLDEEYKVNVESVKDKITSNTVAIVGIAGTTELGKVDPIPELS
KLCEDENIYLHVDAAFGGFVIPFLKDIGYKLPDFDFKLGGVSSITIDPHKMGLVPVPAGG
ILFRKKEYIDVQSVYTPYLTEERQSTIVGTRTGASVAATWAMLKYMGREGYRKVVRECME
TTKFLAKKISKIGLDLITKPELNIVAFDPGDTYEVAKKLENLGWLVSVSKNLDAIRIVVM
PHITKDHVKKFIEDLEDVI
>A_W0K0Y4
MTRAEPRPAPQNFDRVLSSMCTEPHPAAREAAVEFLADNPGDPATYPAVSELEAEAVGML
GDVVGLDDPHGYVGSGGTEANLQAVRAARNLADGDVNVVAPESAHFSFQKAAEVLDVELR
LAPLDDDHRADVGAVTDLADDDTALVVGVAGTTEFGRVDPIPALGEVATDVGANLHVDAA
WGGFVLPFTDHDWSFADAPVDTMTIDPHKMGQAPIPSGGFLARDPETLDALSIRTPYLES
ETQPTLGGTRSGAGVAGAHAALEALWPAGYREQYERSMANAEFLAAELEGRGYDVVDPVL
PLVAADLPDDEFAALRERGWRISRMAGGELRVVCMPHVTRGMLDRFLADIANIQ
>A_F0T911
MEDKGRSETEIFDELHQFKTRDMTHRSGKILGSMCTCPHPIGLNAFKMFLESNLGDPGLF
KGTQAMEDEVISSLGELLGERDVYGHIITGGTEANIMAMRAARNTFKHNYPDCEDVNIVV
PKSAHFSFKKAADMLCLDLLEAELDENYRVDINSLDELINENTAAVVAIAGTTELGKIDP
VEKISELCLKRGVYLHVDAAFGGYSIPFLNEMGYDLPNFDFSLPGVCSITIDPHKMGLAP
IPTGGILFRKKTFLESISIETPYLTEDRQSTIVGTRTGASTAATWALMNYLGKEGYRKVS
KECMEITELLHRGVVEAGFNPVTEPELNIVAFTSDEMTVDEIADGLERSGWAVSISSYPR
AIRIIVMPHVKEEHVELLLDDLRELRSLHSKE
>A_F7PNC6
MQRAEPQDFGRVLSSMCTEPHPAAREAAERFLATNPGDPGTYQTVSNLERQAVELLGEMT
GLGDPAGYVTSGGTEANIQAVRIARNRAETADPNVVVPDSAHFSFSKAAEMLDVELRRVP
TVDYRADVEAMADAIDDDTVAVVGVAGTTEYGHVDPIPALADLAQSADALMHVDAAFGGF
YLPFTDFAWHFGHAEIDTMTIDPHKVGQAAVPAGGFLARSSDLLDELAIDTPYLESRSQV
TLTGTRSGAGVASAVAAMEALWPDGYRQQYHTSMDNAEWLADALEDRGYTVVGPELPLLA
ADVSLSLIEQLRERGWRVTKTGAGEMRVVCMPHVTRSMLRSFVADLDWY
>A_A0A1H7UJE7
MSHESLDAPPESRSASDAESVDSLFLGSEVGAESYRDAIEQSVDIVANAFGEADSPYTGA
SPEAVEAQLASFDCLPEEGAGLDETLDHVSEEILAETIRVAHPACGAHLHCPPVIPGLAA
EVLLTAANQSLDSWDQSGAATILEQRMVETLAETFGYDDEADGVFTSGGTQSNFMGLLLA
RNKVADERFDHWVTEDGLPPEADRLRILCSADAHFTAKQSAAQLGLGENAVVSIETDDEH
RMSVDALDETLDTLDDEDLVPFALVGTAGTTDFGSIDPLPKLAARAHEHDLWFHIDAAYG
GALAFSDTHHEKLAAIECADSIAVDFHKLFYQPISCGAFLLQDGTNYQYIQRHAAYLNPE
SDDEAGVPNLVSKSVQTTRRFDALKPYVTFRTLGRKRLAKLVDSTLELASETAAYLQDND
RFACLHTPTLNAVVFRYVPEHVPSDTDTDQYVGTLNERIRDTLLNEGRAVVARTEVDGVE
SLKFTLLNPRATWADVESLLDDIERIGSDLEATTDASR
>A_A0A2Z2HQZ6
MRRTNASTKPLMIFPRGMQAEPQAFDRVLSSMCTEPHPVAREAAERFLATNPGDPGTYPT
ISALEDDAIAILGEIAGLEEPAGYLASGGTEANIQAMRIARERADTDRPNVVMPESGHFS
FRKAADLLEIELRIVPTDEDHRADLEAVRACVDDDTAAVVGVAGTTEYGRVDPIPELAEI
AHEVGATCHVDAAWGGFVLPFTDYEWNFAHAPIDTMAIDPHKMGQAAVPAGGLLVRSSDL
LDELAVDTPYLESTSQATLTGTRSGAGVASAVAAMEELWPGGYRTQYVRSQNNAEWLADA
LEKRGYEVVDPTLPLVAADVPRSTFDALRGRGWRISRTATGELRVVCMPHVTRKMLAAFV
ADLDRLEVRASVPVVADD
>A_F7XL45
MKNSGLTESQLFDILKEIKKKDTNYSRVLSAMCTHPHRIAVEAHMMFIESNMGDSGLFPG
TNEMEHCVIDMLSDLMHGQGVHGHMTTGGTESNIQALRSMRNFSESSRPNVVVPESAHFS
FDKIADVLRIEIRKASMDQEFKVDIESFESLIDENTVGLVGVAGSTEFGQIDPIEDISGL
AVENSLPLHVDAAFGGFVIPFLKKDYSFDFSLDGVTSIALDPHKMGLGTIPAGVLLFRGE
EYLSNLQTDTPYLTTQTQHSLTGTRSGGAVAATYAVMNYLGKDGYIEVVDYCMDLTEKLV
EGSYRIGIEPLIEPVMNVVALRIPDADLVRKILREKYGWMVSITRDPRCLRLVMMPHLTI
ANLELFLQDLEKAVKDVDTGMDS
>A_M1XLA6
MQRAEPQEFSRVLSSMCTEPHPTAREAAEQFLASNPGDPGTYGTVSTLEREAVDRLGTVA
ELADPAGYIASGGTESNVQAIRLARNRADTRTPNFVAPESAHFSFRKAAGVLGVELRTAP
LSDYRANLDAVAELIDSDTVCVVGVAGTTEYGRVDPIPALADMAADAGALCHVDAAWGGF
VLPFTEHAWSFADADIHTMTIDPHKMGRAAVPAGGLLARGPELLDELAIDTPYLESTSQM
TLTGTRSGAGVASAAAVMDELWRDGYGRQYRRARSNADWLAAELDDREFEVVEPALPIVT
VDLPARLIDDLRDAGWRLSRTEAGEARIVCMPHVTRSMLAEFLGDVDRLAE
>A_A0A328RY55
MHDKGRSKEDIFKDLNKFHSMDMCYDSGKILGSMCTKPDPIGVKAYEMFLETNLGDSGLF
KGTSMMEIDVINSLGRLLHLDNPYGHIVTGGTEANIMAMTVAKYLFEEEHTGVPELILPK
SAHFSFKKVLSMLSIKPVFVPLNDEYKIDVSKLDDLITENTMAIVAIAGTTELGLIDDIK
NISKIAYSNQVYLHVDAALGGFMIPFINMSRKPKIKFDFECKGVSSMTIDPHKMGLAPVP
AGGIIFRKKEYLEKLAVKTPYLTRDHQTTLVGTRTGASTAATWALINYYGKEGYKKIVDS
VLDLTKYTYTILKDMKHVHIVHKPELNLLSFISDNMEVDVLQERLLEYGWRVSVSEYPHA
IRLVLMPHIKKEHINQFLIDLENILNE
>A_L0KZR1

MQEYGQNWEDIESALQQAKSKDISYEKVLSSMCTYPHPVAVEAHRIFIESNMGDYGLFMG
TYELEKSVLTMLGDLLHNSHPYGYLTTGGTESNIQAVRAMRNACTSIKDPNIIVSGSAHF
SFDKIADILKINVRKARILPDLVVDTEDVLSLIDKNTVGLVGIAGSTEFGQVDPISELSK
IAIDNDLPLHIDAAFGGFLLPFLPNHVPFDFSLPGVTSIAIDPHKMGLSTIPSGALLFRE
EKMMELLKVDTPYLTISSQCTLTGTRSGASVASTYAVMKHLGKEGYQQVVNKCMKLTNLL
LDETKNIGVKPVIDPVMNIVALSVEEPKKIRTELATQFGWQVSVTKQPSSIRLVVMPHMT
EENILAFVRDLKTVIQQLQSSEINMQE
>A_B5ICZ4
MDEKEIEDLLQKYYLKDMHYEDGKILGSMYTKPPEIALKAFFKFYQANLGNPGLYKGTVE
IEREVVKFLLRLTSGKDDFFGHVVSGGTEANVIALWAAREMGYKRVLATQDAHFSIRKAA
NLLKLSLENVEIIKGRMSIEDLERKIKGGDIIVATAGTTPLGFIDPIEEIGKICEMHNCF
LHVDAAFGGYVIPFLRELGYTNKKFGFDISAVRTITIDPHKMGMAPYPAGGIVSKENIFE
KIEIEAPYLMVGKNEGLLGTRQSGSVAAAYAAQLYFGWDGYREIVKKCMENTNYLVKRAR
EENFEILEMPEMNIVNIKIKNVGKVKKELYARGWGISTNPKYSSLRIVVMPHVTKEIIDE
FLGELKNIKKSYGL
>A_C7P526
MSGGAPDATERLDQPPVDDAFLGSDEGNRAYAAAADAATRAVVTATDRATPYSGADPDTL
RDRFAGRRVLPERGQSVEETLGEVTDEVLSSVVGVFDPDCVAHLQCPPTIPGLAAETLVA
GTNQSMDSFDQAPAPSVCEERVVDALCDLLSFPAGADGVFTSGGTQSNLQGLLLAREWYC
RERLDCDVQTEGLPADADDLRVVTSEAAHFTAAQATAQLGLGEDAVVEVPTDDGYRMDPD
ALDATLADLTAAGCRPFALLGTAGTTDHGAVDPLPALADRAAEHDLWFHVDAAYGGALLL
SERERSTLDGIDRADSVAVDFHKLFYQPISCGAFLLGDGSQFRLQDRNAAYLNPEADDEA
GVPNLVGKSLQTTRRFDALKPYVTFRTLGRERLADWVEYVVDLATAVGDDVRDHPELELV
CEPQLSTVLFRYRPDEGDPDEINPAIRDRLLRAGRAVIARTEVGGTATLKFTLLNPRATR
SSLRALLESVVEHGTEIERERST
>A_C7P1L0
MQRAEPQSFERVLSSMCTEPHPTARKAAERFLATNPGDPGTYETVADLEREAVERLGTIA
GLGDPAGYVTSGGTEANVQAIRIARNRGDTDDPNVVAPEHAHFSFTKAAELLGVELRTAP
ATDYRADMDAMTHLADDDTVAVVGVAGTTEYGYVDPIPAVADLADAVGALCHVDAAWGGF
YLPFTDHDWHFGHADVDTLTIDPHKVGQAAVPAGGLLARSPDLLDELAIDTPYLESRSQV
TLTGTRSGAGVASAVAAMDALWRDGYRETYERAMGNAEWLAEQLDVRGHDVIGPELPLVA
ADLSIPMTTELRDRGWRVSKTGSGEMRVVCMPHVTRSMLRSFVADLDWY
>A_A2SSB4
MTNTEKTNTSPCAAEDALKVKALFLGPKSENREYFKNMLNFLMDEHMHWRSDFHPEDRMV
STAKEMRSEEYLTTLDRTSDVLMQLSNKLKETSMPWFSSRYLGHMNSDTLMVANLAYMAT
ILYNPNNVAYEASTATTPMEIEAGKDMATMLGFDPEQSWGHITTDGTIANYEGMWMARNL
KSFPPAVKKIRPDLVPGLDDWQIMNMSTTQVLDLIGKVKEAGCFDEVRNESARGIGAGDG
SLGVVLVPQSKHYSWVKAADVLGIGNKNLIQVQVNDHYHMDIDTLKSIIDDHIARKIPIM
AVVAVVGTTEEGAIDEVDRIVELRAEYEKQGINFYFHIDAAYGGYSRALYLDEQNRFMEY
DEVKERLSADGIFIHETEYPQREIYEAYKAIPAADSITIDPHKMGYIPYSAGGITIKDRR
ILDLISYFAAYVFESGEDSPTLLGSYIMEGSKAGATAAAVWATHRLIPLNVTGYGRIIGR
SIEGAQMLSNALTTTKYITVNDRKFQVETLAGKPDFNIVCMAFNEVGNTDLDAMNALNEK
IYNESSYVSGPVYKNDWITSKTALAREDYGDAPRNFVKRLGVPTAEWDRVGSVYVLRVCL
LNPFVSHNVHFDVLWDGLLAILKEKLAAAIAGDGKC
>A_A2STQ3
MEEKGCKREEVISLLSAYRAEDLHHDHILSSMCTIPHEMAVFVHGMFSATNLGDPGLFPG
TTKIEDRLVHSLGELMHHPGAGGYATSGGTESNLQAIRIAKKLKPEIKNPNIVVPASAHF
SFDKTCDILGLEMRTVPYGKNYTVDCDKMAEMVDKNTISVSAIAGTTEYGMIDDVERIAK
IALENDLFFHVDAAFGGMVIPFLPNPAPFDFEVPGVSSISLDPHKMGMSTIPCGCLLLRE
PEQFGTLNVDTPYLTVKKECTLAGTRPGADVAGAYAVIKLLGREGFRAVVAGCMENTRRL
IEGMEAFGYTRAVDPVMNVATFEAGPVPKGWIVSHTRAGHLRFVVMPHVTRDVIENFLAD
VAKIN
>A_A0A1I4CXR5
MDTSPPGVGTFLGTGAGSATYREAMDAAVDAVLDASASGPYSGETYDELEDEFDVDTLPD
TGSDLTHVVEWVGEHVLANSVVTSDPTCVAHLQCPTAIPGLAAEAMLTALNQSMDSWDQS
PAATVVEESMVEELCSLFGLPNAADGVFTSGGTASNLLGLLLARDRYVTETFGQRVQDEG
LPPEASDLRILCSDAGHFTAQQSAAVLGLGEDAVVSVPTDDDYRMDPEALDAELARLDDE
GKRPFALVATAGTTDFGSVDPLSALADRAAEHDLWFHVDAAYGGALAVSDRHREMLAGVS
RADSVSLDFHKLLYQPISCGAFLLRDGSHYDLIDRNASYLNPEADEEAGVSNLVGKSLAT
TRRFDALKPFVTFQTLGREGVAELVDYTLQLADDVAELVERDPALELAADPTLNAVVFRY
RPSQQHSDRETSEWVDTVNRRIRDHLLERGMGVVARTEVDGDAYLKFTLLNPQTTVDDVA
DLLVSVKQYGAAIESEVNE
>A_D7DV28
MDKSKEILSKLKEYRDLDLKYEKGNIFGSMCTKPHPITLEIIKMFYETNLGDPGLFIGTK
KLEEESIQMIGKLLHNPNAFGYIISGGTEANITAMRLFNNISKANFKNKKYGNKKNREDS
SKIIIPETAHFSFDKSKDMMNLDLIRPPLTEYYTSNVKWVKDYVEDTISKNGENSISGIV
GIAGCTELGTIDNIKELSKIAYTNDIPLHVDAAFGGFVIPFLEEKYKLKNYNYEFDFSLD
GVKTITIDPHKMGLSPISAGGIIFRNREYKKYLDIEAPYLTETLQATILGTRTGVGAATT
WGLLKLLCKDGYAKITHECMEKTTYLTNKLRENGFETVIEPVLNIIAIKDDNAKETCKKL
KEKGLYVSVCRCTNALRIVIMPHLEFEHLDNLVNTLCKINKK
>A_A0A1D2R9X5
MDDEKILNKLRELRAKDPDFSKGRVFSAVSSPPLEVALDAFKIFADTNALDEHLFSATGE
LERECISWLGNLLHNPKAAGYITTGGTEANMFALWAARESSKNKNEIIVPESAHYSIEKI
ARVMNLKINYTGLDENFRADVSEIESKINDKTLAVIATAGTPSLEMIDPVAKINELCDNI
FLHVDAAFGGFVIPFLDRKIPFDFKLSNVSSITIDPHKMGLAPFPSGAVILRSSKLLKNL
KILPPYLPIETYTLLGSRSGGAIASTWATFKSLGFAGYKKIVSGCMENTKFFCRELKVRG
FELLTEPDLNCIGIKIDNEKIVKKLEYVGWKISMNHKPKSLRIVVMPHVNKEDILAFIDD

MEKIAGVNK
>A_L0ALZ0
MRAEPQSFDRVLSSMCTEPHPAAREAAERFLATNPGDPGTYPTVADLEDDAVSLLGEIAG
LDEPAGYVASGGTEANVQAVRIARERAENGRPTVVLPESAHFSFQKAADLLDVELRVVPT
TDDGRADLEAVRACVNEDTAAVVGVAGSTEYGRVDPIPELGEIADSVDALLHVDAAWGGF
VLPFTDYEWHFGHAPVDTMAIDPHKMGQAAVPAGGLLARSSDLLDELAVETPYLESTSQA
TLTGTRSGAGVASAVAAIEELWPDGYRDQYVRSQNNAKWLASKLESRGYDVVEPALPLVA
ASVPTPLFEALRDEGWRLSRTGDGELRVVCMPHVTRDGLESFVADVDRLETRVGLEVAGA
SE
>T_K4HXK6
MAMLYGKHTHETDETLKPIFGASAERHDLPKYKLAKHALEPREADRLVRDQLSDEGNSRL
NLATFCQTYMEPEAVELMKDTLEKNAIDKSEYPRTAEIENRCVNIIANLWHAPEAESFTG
TSTIGSSEACMLAGLAMKFAWRKRAKANGLDLTAHQPNIVISAGYQVCWEKFCVYWDIDM
HVVPMDDDHMSLNVDHVLDYVDDYTIGIVGIMGITYTGQYDDLARLDAVVERYNRTTKFP
VYIHVDAASGGFYTPFIEPELKWDFRLNNVISINASGHKYGLVYPGVGWVIWRDQQYLPK
ELVFKVSYLGGELPTMAINFSHSASQLIGQYYNFIRFGFDGYREIQEKTHDVARYLAKSL
TKLGGFSLINDGHELPLICYELTADSDREWTLYDLSDRLLMKGWQVPTYPLPKNMTDRVI
QRIVVRADFGMSMAHDFIDDLTQAIHDLDQAHIVFHSDPQPKKYGFTH
>T_A0A454A8X6
MDQKLLTDFRSELLDSRFGAKAISTIAESKRFPLHEMRDDVAFQIINDELYLDGNARQNL
ATFCQTWDDDNVHKLMDLSINKNWIDKEEYPQSAAIDLRCVNMVADLWHAPAPKNGQAVG
TNTIGSSEACMLGGMAMKWRWRKRMEAAGKPTDKPNLVCGPVQICWHKFARYWDVELREI
PMRPGQLFMDPKRMIEACDENTIGVVPTFGVTYTGNYEFPQPLHDALDKFQADTGIDIDM
HIDAASGGFLAPFVAPDIVWDFRLPRVKSISASGHKFGLAPLGCGWVIWRDEEALPQELV
FNVDYLGGQIGTFAINFSRPAGQVIAQYYEFLRLGREGYTKVQNASYQVAAYLADEIAKL
GPYEFICTGRPDEGIPAVCFKLKDGEDPGYTLYDLSERLRLRGWQVPAFTLGGEATDIVV
MRIMCRRGFEMDFAELLLEDYKASLKYLSDHPKLQGIAQQNSFKHT
>T_A0A454A444
MDKKQVTDLRSELLDSRFGAKSISTIAESKRFPLHEMRDDVAFQIINDELYLDGNARQNL
ATFCQTWDDDNVHKLMDLSINKNWIDKEEYPQSAAIDLRCVNMVADLWHAPAPKNGQAVG
TNTIGSSEACMLGGMAMKWRWRKRMEAAGKPTDKPNLVCGPVQICWHKFARYWDVELREI
PMRPGQLFMDPKRMIEACDENTIGVVPTFGVTYTGNYEFPQPLHDALDKFQADTGIDIDM
HIDAASGGFLAPFVAPDIVWDFRLPRVKSISASGHKFGLAPLGCGWVIWRDEEALPQELV
FNVDYLGGQIGTFAINFSRPAGQVIAQYYEFLRLGREGYTKVQNASYQVAAYLADEIAKL
GPYEFICTGRPDEGIPAVCFKLKDGEDPGYTLYDLSERLRLRGWQVPAFTLGGEATDIVV
MRIMCRRGFEMDFAELLLEDYKASLKYLSDHPKLQGIAQQNSFKHT
>T_Q42472
MVLTKTATNDESVCTMFGSRYVRTTLPKYEIGENSIPKDAAYQIIKDELMLDGNPRLNLA
SFVTTWMEPECDKLIMDSINKNYVDMDEYPVTTELQNRCVNIIARLFNAPLEESETAVGV
GTVGSSEAIMLAGLAFKRKWQNKRKAEGKPYDKPNIVTGANVQVCWEKFARYFEVELKEV
NLSEGYYVMDPDKAAEMVDENTICVAAILGSTLNGEFEDVKRLNDLLVKKNEETGWNTPI
HVDAASGGFIAPFIYPELEWDFRLPLVKSINVSGHKYGLVYAGIGWVVWRAAEDLPEELI
FHINYLGADQPTFTLNFSKGSSQIIAQYYQLIRLGFEGYKNVMENCIENMVVLKEGIEKT
ERFNIVSKDQGVPVVAFSLKDHSFHNEFEISEMLRRFGWIVPAYTMPADAQHITVLRVVI
REDFSRTLAERLVADISKVLHELDTLPSKISKKMGIEGIAENVKEKKMEKEILMEVIVGW
RKFVKERKKMNGVC
>T_A0EJ89
MALSSATDSDGSIHSTFASRYVQESLPRFQIPSRSIPKDAAYQIISDELMLDGNPRLNLA
SFVTTWMEPECDKLIMQAINKNYVDMDEYPVTTELQNRCVNIIANLFNAPLGDGEEAVGV
GTVGSSEAIMLAGLAFKRKWQNKRKAEGKPYDKPNIVTGANVQVCWEKFARYFEVELKEV
KLKKDYYIMDPVKAVEMVDENTICVAAILGSTYNGEFEDVKLVNDLLIQKNKETGWDTPI
HVDAASGGFIAPFLYPELEWDFRLPLVKSINVSGHKYGLVYAGIGWVIWRAKQDLPEELI
FHINYLGADQPTFTLNFSKGASQIIAQYYQLIRLGFEGYRNIMGNCAANAKALSDGLVRT
GRFNILSKEIGVPLVAFSLKDSSRHDEYEISDHLRRFGWIVPAYTMAPDAQEVKLLRVVV
REDFNRSLAERLVHDIEKVLHELDTLPSKIAREVVASLVDGHPELKEVKDLGIDVTQFKS
SAVFNEIVNSQKAVKAWKKFVAQKANRVC
>T_Q04792
MLHRHGSKQKNFENIAGKVVHDLAGLQLLSNDVQKSAVQSGHQGSNNMRDTSSQGMANKY
SVPKKGLPADLSYQLIHNELTLDGNPHLNLASFVNTFTTDQARKLIDENLTKNLADNDEY
PQLIELTQRCISMLAQLWHANPDEEPIGCATTGSSEAIMLGGLAMKKRWEHRMKNAGKDA
SKPNIIMSSACQVALEKFTRYFEVECRLVPVSHRSHHMLDPESLWDYVDENTIGCFVILG
TTYTGHLENVEKVADVLSQIEAKHPDWSNTDIPIHADGASGGFIIPFGFEKEHMKAYGME
RWGFNHPRVVSMNTSGHKFGLTTPGLGWVLWRDESLLADELRFKLKYLGGVEETFGLNFS
RPGFQVVHQYFNFVSLGHSGYRTQFQNSLFVARAFSFELLNSSKLPGCFEIVSSIHESIE
NDSAPKSVKDYWEHPQAYKPGVPLVAFKLSKKFHEEYPEVPQAILSSLLRGRGWIIPNYP
LPKATDGSDEKEVLRVVFRSEMKLDLAQLLIVDIESILTKLIHSYEKVCHHIELASEQTP
ERKSSFIYEMLLALASPQDDIPTPDEIEKKNKLKETTTRNYRGTC
>T_Q99259
MASSTPSSSATSSNAGADPNTTNLRPTTYDTWCGVAHGCTRKLGLKICGFLQRTNSLEEK
SRLVSAFKERQSSKNLLSCENSDRDARFRRTETDFSNLFARDLLPAKNGEEQTVQFLLEV
VDILLNYVRKTFDRSTKVLDFHHPHQLLEGMEGFNLELSDHPESLEQILVDCRDTLKYGV
RTGHPRFFNQLSTGLDIIGLAGEWLTSTANTNMFTYEIAPVFVLMEQITLKKMREIVGWS
SKDGDGIFSPGGAISNMYSIMAARYKYFPEVKTKGMAAVPKLVLFTSEQSHYSIKKAGAA
LGFGTDNVILIKCNERGKIIPADFEAKILEAKQKGYVPFYVNATAGTTVYGAFDPIQEIA
DICEKYNLWLHVDAAWGGGLLMSRKHRHKLNGIERANSVTWNPHKMMGVLLQCSAILVKE
KGILQGCNQMCAGYLFQPDKQYDVSYDTGDKAIQCGRHVDIFKFWLMWKAKGTVGFENQI
NKCLELAEYLYAKIKNREEFEMVFNGEPEHTNVCFWYIPQSLRGVPDSPQRREKLHKVAP

KIKALMMESGTTMVGYQPQGDKANFFRMVISNPAATQSDIDFLIEEIERLGQDL
>T_Q548L6
MASSTPSPATSSNAGADPNTTNLRPTTYDTWCGVAHGCTRKLGLKICGFLQRTNSLEEKS
RLVSAFRERQSSKNLLSCENSDQGARFRRTETDFSNLFAQDLLPAKNGEEQTAQFLLEVV
DILLNYVRKTFDRSTKVLDFHHPHQLLEGMEGFNLELSDHPESLEQILVDCRDTLKYGVR
TGHPRFFNQLSTGLDIIGLAGEWLTSTANTNMFTYEIAPVFVLMEQITLKKMREIVGWSN
KDGDGIFSPGGAISNMYSIMAARYKYFPEVKTKGMAAVPKLVLFTSEHSHYSIKKAGAAL
GFGTDNVILIKCNERGKIIPADLEAKILDAKQKGYVPLYVNATAGTTVYGAFDPIQEIAD
ICEKYNLWLHVDAAWGGGLLMSRKHRHKLSGIERANSVTWNPHKMMGVLLQCSAILVKEK
GILQGCNQMCAGYLFQPDKQYDVSYDTGDKAIQCGRHVDIFKFWLMWKAKGTVGFENQIN
KCLELADYLYAKIKNREEFEMVFDGEPEHTNVCFWYIPQSLRGVPDSPERREKLHRVAPK
IKALMMESGTTMVGYQPQGDKANFFRMVISNPAATQSDIDFLIEEIERLGQDL
>T_Q05329
MASPGSGFWSFGSEDGSGDSENPGTARAWCQVAQKFTGGIGNKLCALLYGDAEKPAESGG
SQPPRAAARKAACACDQKPCSCSKVDVNYAFLHATDLLPACDGERPTLAFLQDVMNILLQ
YVVKSFDRSTKVIDFHYPNELLQEYNWELADQPQNLEEILMHCQTTLKYAIKTGHPRYFN
QLSTGLDMVGLAADWLTSTANTNMFTYEIAPVFVLLEYVTLKKMREIIGWPGGSGDGIFS
PGGAISNMYAMMIARFKMFPEVKEKGMAALPRLIAFTSEHSHFSLKKGAAALGIGTDSVI
LIKCDERGKMIPSDLERRILEAKQKGFVPFLVSATAGTTVYGAFDPLLAVADICKKYKIW
MHVDAAWGGGLLMSRKHKWKLSGVERANSVTWNPHKMMGVPLQCSALLVREEGLMQNCNQ
MHASYLFQQDKHYDLSYDTGDKALQCGRHVDVFKLWLMWRAKGTTGFEAHVDKCLELAEY
LYNIIKNREGYEMVFDGKPQHTNVCFWYIPPSLRTLEDNEERMSRLSKVAPVIKARMMEY
GTTMVSYQPLGDKVNFFRMVISNPAATHQDIDFLIEEIERLGQDL
>T_P48320
MASPGSGFWSFGSEDGSADPENPGTARAWCQVAQKFTGGIGNKLCALLYGDSGKPAEGGG
SVTSRAATGKVACTCDQKPCNCPKGDVNYAFLHATDLLPACDGERPTLAFLQDVMNILLQ
YVVKSFDRSTKVIDFHYPNELLQEYNWELADQPQNLEEILTHCQTTLKYAIKTGHPRYFN
QLSTGLDMVGLAADWLTSTANTNMFTYEIAPVFVLLEYVTLKKMREIIGWPGGSGDGIFS
PGGAISNMYAMLIARYKMFPEVKEKGMAAVPRLIAFTSEHSHFSLKKGAAALGIGTDSVI
LIKCDERGKMIPSDLERRILEVKQKGFVPFLVSATAGTTVYGAFDPLLAVADICKKYKIW
MHVDAAWGGGLLMSRKHKWKLSGVERANSVTWNPHKMMGVPLQCSALLVREEGLMQSCNQ
MHASYLFQQDKHYDLSYDTGDKALQCGRHVDVFKLWLMWRAKGTTGFEAHIDKCLELAEY
LYTIIKNREGYEMVFDGKPQHTNVCFWFVPPSLRTLEDNEERMSRLSKVAPVIKARMMEY
GTTMVSYQPLGDKVNFFRMVISNPAATHQDIDFLIEEIERLGQDL
>T_K7WYY0
MTSNSCNYHLTEKTAKLLATDLLPYKESAGPQTKEFLQKVIDVLMDFVRATNDRNEKVLD
FHHPEEMFRILDLDIPEKGLPLQQLIKDCETTLKYQVKTGHPRFFNQLSCGLDLVSMAGE
WLTATANTNMFTYEIAPVFILMERVVLTHMRELIGWNGGDSILAPGGSISNLYAFLAARH
KMFPGYKEKGSSVIPGELVMFTSDQSHYSVKSCASVGGLGTDNCVMVPSDLNGKMVPREL
ERLIIERKSKGQIPFFVTATAGTTVLGAFDPINEIADICEKYNCWLHVDAAWGGGLLLSK
KYRHPRLSGIERAKSVTWNPHKLMGALLQCSTIHFKEDGLLISCNQMSAEYLFMTDKLYD
VQYDTGDKVIQCGRHNDVFKLWLQWRAKGSEGFEKHMDRLMELTEYMVKRLRTMPDKYYL
IMEPECVNVSFWYIPRRLRGIIHDAKREAELGKICPILKARMMQSGTLMVGYQPDDRRPN
FFRSIISSAAVTEGDVDFMLAEFDRLGQDL
>T_K7XPX5
MAAQRKTEDVSKLRYTDLYPFRDDSAPTREFLQRVFDILWAYVDQQQDRSSKILDFHMPE
ELMQILDLELPDEPQPLQRVLGDCAEALKHQVRTGHPHFFNQLSSGLDIVSLAGEWLSAT
ANTNMFTYEIAPVFILMENVVMKKMRDLIGYTNGDSILAPGGSVSNLYAVMAARHKMFPS
YKTLGLKALPQLVMYTSEDSHYSVKGAGASIGLGTDNVVSIPVDKCGRMKVDLLEKEIQA
SKARGHVPFFVNCTAGSTVIGAFDPIHPIADICQRHGLWLHVDAAWGGGCLLSKKHRHLL
DGVERSDSVTWNPHKLMGTHLQCSTIHLKEDGLLLSCNQMCAEYLFQQDKHYDVSYDTGD
KVPQCGRHNDIFKLWLMWRAKGTVGFERQIDHLFDMSNYLVQKIKDRPDMFHLLLEPELV
NVCFWYIPKRLRGKPHSKEKEQELGVVTAQLKARMMNTGTLMITYQPIWDKPNFFRNIVS
NAGVRREDIDFLVDELDRLGHDL
>T_Q171S0
MPANGMFDVALQVIDDSNVSSGSDSAGVSEDEDVQLFCSKGNTIVPKPLKKSISKIKDEEFSKTAKANEKRYASLPSREHHQQFLTDFLSEVLN
NAVFNATERANKVLNWVDPEQLKRTLDLELKDEPDSHEKLLELTRATIKHSVKTGHPYFMNQLFSSVDPYGFAGQILTDALNPSVYTFEVSPVF
VLMEEVVLKEMRTIVGYPDGTGDGIFCPGGSMANGYSISCARFKHMPDVKTKGLHSLPRLVIFTSEDAHYSVKKLASFMGIGSDNVYPIHTDAI
GKIRVDHLESEILRAKSEGAVPFMVSATAGTTVIGAFDPLEQIADLCKKYNLWMHVDAAWGGGALMSKKYRSLLKGIERSDSVTWNPHKLLAAP
QQCSTFLTRHEGILSECHSTNATYLFQKDKFYDTQYDTGDKHIQCGRRADVLKFWFMWRAKGTSGLEQHIDKVFENAEHFTSSIKSREGFEMVV
ENPECTNVCFWYVPPGLRNVPRDSAEFTERLHKVAPKVKERMMREGSMMITYQPIHDKPNFFRLVLQNSALDKSDMNYIIDEIERLAADL
>T_A7U8C7
MPATGEDQDLVQDLIEEPATFSDAVLSSDEELFHQKCPKPAPIYSPVSKPVSFESLPNRRLHEEFLRSSVDVLLQEAVFEGTNRKNRVLQWREP
EELRRLMDFGVRSAPSTHEELLEVLKKVVTYSVKTGHPYFVNQLFSAVDPYGLVAQWATDALNPSVYTYEVSPVFVLMEEVVLREMRAIVGFEG
GKGDGIFCPGGSIANGYAISCARYRFMPDIKKKGLHSLPRLVLFTSEDAHYSIKKLASFQGIGTDNVYLIRTDARGRMDVSHLVEEIERSLREG
AAPFMVSATAGTTVIGAFDPIEKIADVCQKYKLWLHVDAAWGGGALVSAKHRHLLKGIERADSVTWNPHKLLTAPQQCSTLLLRHEGVLAEAHS
TNAAYLFQKDKFYDTKYDTGDKHIQCGRRADVLKFWFMWKAKGTSGLEKHVDKVFENARFFTDCIKNREGFEMVIAEPEYTNICFWYVPKSLRG
RKDEADYKDKLHKVAPRIKERMMKEGSMMVTYQAQKGHPNFFRIVFQNSGLDKADMVHLVEEIERLGSDL
>T_Q6L2R7
MLKRFPERGIPLDEINSILDGYGKNDIKNSRGRLFTYFYDPGLKDLDDLSSILLKFYNRNGMDYHAFPSTLKIENDLISMMSDLMHGNDDTSGT
FTTGGTESILLAMKAARDLFLEKKEYVPEIVAPVTAHPAFSKAAKYLGMKITRVPVNEDYIADDTINEYINDRTAAVIASAPSFPYGGIDNIKD
ISEIALDKNTWFHVDACVGGMILPFLKGLGLNIKDFDFKLPGVSSMSIDLHKYGFTPKGSSVVLYKNHDLRKRQIYVNADWPGYPMSNMGMQAT
KSAGPLAGSWATLNYLGLDGYKKLAEKTLKAYRMIRSGITDLGYKIIGRPDATIFAFTHNDKDIIDLGIKMIENGWYPQIQPGNVFIDMPDSVH
LNISPVHLDVADEFLEFFNELDKNVKPRDKNSYNVNSVEKALEMIKMEKSMLFRIIRYSRPEVSEKIFLEMTDEDFTYS
>T_O28946
MDSSSVDIVLNKGGHNKSFEVKIYRIHTTKIMSFPNGSDAEGVLKRLEDYAKNDFEPHSRRMWGHIYYAGLKDVVELARKAYLMYMDKTMLDFT
CFPSLLRMEREVVRMASSLLNGDEEVVGNFTYGGTESIMLALKAAREKFRKEEGGNVVPEIVLPATAHPAFWKSAEYLGMRCLRAKLDDELRAD

VETVKELVGDKTAMIVGSAPNYPFGVVDDIKALSDIAVDGKLWLHVDACLGGFHLPFFRELGEKIPDFDFSVEGVHSISADFHKYGLSPRGASV
ILYRNAKLREGQIFVMASWPGYPLVNTAVLSTRSAGTLAAAWAVMSYLGFDGYLKLAKKTLYAKKRLIDGLTELGLELLGSPEGAVLAFTSERH
NLFKVSTLMAEKGWYVQSQPGSKKLGFPRSLHFSVIPGHAEVVDEFLEDMREVLPECECSYPEMSSFDVSKLKFGEDGLPEDSELISELIHSMP
PEIVESVFKQFINELIFR
>T_Q4J9W7
MKAFPDKPLSKQDIYEIAKTYSRNDNEPLSGRMWGHIYSLGLPEDVIEVSMTLYNQFINKTMLDFTVYPSVLRFENDIIAMASSLLGGNEETVG
NFTFGGTESIMVATKSARDYFLKRHSSVIPEILLPVTAHPAFNKASDYLGMKVTPVKIDPERTTVDLEDLKSKLKENTAMIVASAPNYPFGTID
DVKALSEIAQDKKLWLHVDSCIGGFLLPFLRDLGEPIPPFDLSLEGVTSISADLHKYGYAPRGASVVLFRNSSYREGSIFVMSRWPGYPIVNTS
VLSTRSAGPLAAAWGIIHGLGKDGYRKLANRILNTRIKLTNELPKMGYRILGKPLGGIVSFTSEEFNLAELPTLMKGWFIQYQPGSRILGFPKS
IHLTIAPGHDKVVDEFLRDLERATIELRGKRLTLPPLQDFNDLQSIARALGIEEGKLPSNPTLINELMHEMPPELVENVLKLVINEYVFRPSRS
>T_Q9HSA3
MTRGEARRPPQEFDRVLSSMCTTPHPAAREAAQAFLATNPGDPETYPAVAERERDAVALLGEIVGLSSPHGYIAAGGTEANLQAVRAARNRADA
DAVNVVAPASAHFSFQKAADVLGVELRLAPTDGDHRADVAAVADLVDGDTAVVVGVAGTTEYGRVDPIPALADIAAGVDANLHVDAAWGGFVLP
FTDHDWSFADAPVNTMAIDPHKMGQAPVPAGGFLARDPETLDALAIETPYLESDTQPTLGGTRSGAGVAGALASLRALWPDGYREQYERTQGNA
EYLAAELAARGYDVVDPELPLVAADMPDAEFQALREEGWRISRTASDALRVVCMPHVTREMLAAFLDDVDALA
>T_Q2FSD2
MDAEGLSTDELFCFLQAKRNEDFSYSHILSSMCTTPHPVAVQAHNLFMETNLGDPGLFPGTATLEDRLIRWFADLYHEPSAGGCTTSGGTESNI
QVLRFCKKTKNVKEPNIIVPASAHFSFEKACGMMDIEMRVVPVDEQYRMKTDAAGELIDKNTCCIVGVAGTTEYGMTDPIPALGKLAEQEGVHL
HVDAAFGGYVLPFLDDAPPFDFSVPGVGSIAVDPHKMGLSTIPSGVLMVRDERVFCNLLVETPYLTTKQAYSLTGTRPGASVAAAYAVMAYLGR
KGMKALVTGCMENTRRMIEGMEAFGVHRKVTPDVNVATFEHVSVPSPWVVSYTRKGDLRIVCMPHVTRDVVEAFLSDFGESYVSHIS
>T_A0B9M9
MYTFPEKGLSEDMVTDLLKEMRSRDCPYDRLLSTMCTRPHPVAVRAYSMFLETNLGDPGLFPGTAEIERRVVGILGSLLGCSDATGYVSTGGTE
SNIQAVRAARNSSGRRDGNIVVPRSAHFSFDKIADLLNLEVRKAELDESLRVDVGDVERLIDDRTVCLVGIAGTTEFGQVDPIGDLSELAIENG
IPLHVDAAFGGFVLPFLEKDCMWDFRAEGVQSITIDPHKMGMSPIPAGGLIFRSSDPLRRLETETYYLTVSRQASLTGTRSGAAAAATYAVIMH
LGIDGYRKVVRRCMDMTEHLVSEARAMGIEPVIEPVMNVVALRVDDPPGVRRALLERGWHVSMTREPKALRLILMPHMTDENLDLFLSDLEDVL
ISLRRGG
>T_Q8PXA5
MNEQGLSEREIFSYLENAKSEDTDYYRVFSSMCTRPHKIAIEANRLFIEANLGDLGLFAGAHKLEQEVVRMLGNLLHASSIDVPSGGLCQSSVC
GYLTTGGTESNIQAVRGMKNLVTAGKKEFKGTPNIVIPASAHFSFDKVADMMGIEVRRASLDSEFRVDMASVEKLINENTIGLVGIAGNTEFGQ
IDPIDKLSEVALENELFLHVDAAFGGFVIPFLEKPQPFDFKVPGVTSIAIDPHKMGLSTIPSGALLFRSPSFLDSLKVSTPYLTTKSQFTLTGT
RSGASAAATCAVMKYLGYEGYRKNVQYCMELTSKIVEEARKLGFEPLIEPVMNVVALKVPNPDLVRERLLKKFGWNVSITRTPRALRLVLMPHN
SPEDIELFLEDLKKVTAEIKSP
>T_Q12VA2
MEENGKTKEEILLFLKKAKSADASYERVLSSMCTYPHEIAVLAHTQFIESNMGDPGLFPGTFNLEKQVLAMFGKMLHHKNSPEKAGYLTTGGTE
SNIQAIRSMHNFRHDISRPNIVMPESAHFSFDKVANLSGIEIRKASLDKLLKVDLDSVRSLIDKNTIGLVGIAGTTEFGQLDPINELSKIAIEK
GIFLHIDAAFGGFVIPFMDIDYTYDFRLEGVTSMTIDPHKMALSTIPSGGLLFKEPEYFECLEIHTPYLSVNKQYSLTGTRSGAGVASTYAVMK
HLGRKGYKKVVSDCMSVTKKLVDGAEKLGINTVIDPVLNIVALDVPEADLVRKKLLDEYGWHVSITRNPRALRIVIMPHIKNETIELFLKDLAK
VIK
>T_O27188
MTYTSGRILGSMCTSSHPLARRVYCDFLESNLGDPGLFRGTRELESGVIGMLGELLSEPDAAGHIITGGTEANLMAMRAARNMAGAEKPEIIVP
KSAHFSFRKAADILGLRLREAELDQDYRVDVESVRKLISENTVAVVGVAGTTELGRIDPVEELSEICLDEDIHLHIDAAFGGFIIPFLRETGAE
LPEFDFKLQGVSSITVDPHKMGLAPIPSGCILFRDASYLDAMSIETPYLTEKQQSTIVGTRTGASAAATWAIMKHMGREGYRKLALRVMGVTRR
LRDGLVELDYQLVVEPELNIVAFNHPAMGPHELADRLEELGWAVSVSSCPPAIRVVLMPHIMEEHIELLLRDLEGIRLREE
>T_Q60358
MRNMQEKGVSEKEILEELKKYRSLDLKYEDGNIFGSMCSNVLPITRKIVDIFLETNLGDPGLFKGTKLLEEKAVALLGSLLNNKDAYGHIVSGG
TEANLMALRCIKNIWREKRRKGLSKNEHPKIIVPITAHFSFEKGREMMDLEYIYAPIKEDYTIDEKFVKDAVEDYDVDGIIGIAGTTELGTIDN
IEELSKIAKENNIYIHVDAAFGGLVIPFLDDKYKKKGVNYKFDFSLGVDSITIDPHKMGHCPIPSGGILFKDIGYKRYLDVDAPYLTETRQATI
LGTRVGFGGACTYAVLRYLGREGQRKIVNECMENTLYLYKKLKENNFKPVIEPILNIVAIEDEDYKEVCKKLRDRGIYVSVCNCVKALRIVVMP
HIKREHIDNFIEILNSIKRD
>T_Q6M0Y7
MDEQDILNELREYRNQDLKYEEGYILGSMCTKPHPMARKISEMFFETNLGDPGLFKGTSKLEKEVVSMIGGILHNKNAFGYLISGGTEANLTAM
RAFKNISKSKGKPQNIIIPETAHFSFDKAKDMMDLNVVRPPLTKYFTMDVKFIKDYIEDSKNEVSGIVGIAGCTELGSIDNICELSKIAVENDI
LLHVDAAFGGFVIPFLDDKYKLDGYNYDFDFSLNGVSSITIDPHKMGLAPISAGGILFRDNMFKKYLDVDAPYLTEKQQATIIGTRSGVGVAST
WGIMKLLGIDGYETLVNESMEKTMYLVKKAREYGFETAIDPVMNIVALNDENKHDTCMKLRDENWYVSVCRCVDALRIVVMPHLEIEHIDGFLE
SLSNTKKY
>T_Q8TV92
MILQRDSDYSDGTVLGSMCTEPHPVAAEAFVAGLHVNLGDPYLFPNAYRAERECIGWLAETLLDHPAPEEAEGSIVSGGTEANILAAYAAREVT
GGREIIVPATRHFSFEKAARMLRMKLVEAPLRSDYTVDVDAVQDLISRDTALIVGIVGTTETGSVDDIEALSDVAEDHGVPLHVDAAFGGFTAP
FLREEYPLPRFGFDLEAVVSVTVDPHKMGLVPPPAGGIVFRDDEFPKAIEVYAPYLSGGGASQYTITGTRPGAPVLALYANILELGEEGYRRIA
FRCYEETLKVAEKARELGLELAVDPPHLNLVNIRLPDRGTAERLLRESEREGWKISVSTKPLGVRIVMMPHLDAETVSRFLELVARVLGG
>T_Q5JJ82
MFPERGASEEEVLRELEEKTREDLTFDSGKILGSMCTYPHPFAVKVVMKYIDRNLGDPGLHIGSQKIEKEAVDMLANLLGLEKGYGHIVSGGTE
ANILAVRAMRNLAGIEKPELILPESAHFSFIKAAEMLGVKLVWAELNDDYTVNVKDVEKKITDRTIGIVGIAGTTGLGVVDDIPALSDLALDYG
LPLHVDAAFGGFVIPFAKALGYEIPDFDFRLKGVKSITIDPHKMGMVPIPAGGIIFREKKFLDSISVLAPYLAGGKIWQATITGTRPGANALAV
WAMIKHLGFDGYKEVVKEKMELARWFASELKKIPGIYLIREPVLNIVSFGSEKLEELEKELKARGWGVSAHRGYIRIVVMPHVKREHLEEFLRD
LREIAKRL
>T_Q8U1P6
MKFPRKGIPQEEVMRELEKYTSKDLSFSSGKILGSMCTLPHELAKEVFCMYMDRNLGDPGLHPGTKKIEEEVIEMLSDLLHLERGYGHIVSGGT
EANILAVRAFRNLADVENPELILPKSAHFSFIKAGEMLGVKLIWADLNPDYTVDVKDVEAKISENTIGIVGIAGTTGLGVVDDIPALSDLARDY
GIPLHVDAAFGGFVIPFAKELGYDLPDFDFKLKGVQSITIDPHKMGMAPIPAGGIVFRHKKYLRAISVLAPYLAGGKIWQATITGTRPGASVLA
VWALIKHLGFEGYMEIVDRAMKLSRWFAEEIKKTPGAWLVREPMLNIVSFKTKNLRRVERELKSRGWGISAHRGYIRIVSHASCDGGHD
>T_Q9UZD5
MSKFPEKGLPREEVLNLLEDKTKVDLTFSSGKILGSMCTMPHELAIEVFARYIDRNLGDPGLHPGTRKIEEEVIEMLSDLLHLEKGYGHIVSGG
TEANILAVRAFRNISDAERPELILPKSAHFSFIKAGEMLGVKLVWAELKQDYAVDVKDVEAKISDNTIGIVGIAGTTGLGVVDDIPALSDLARE
YGIPLHVDAAFGGFVIPFAKSLGYDLPDFDFKLKGVESITIDPHKMGMAPIPAGGIIFRRKKYLKAISVLAPYLAGGKVWQATITGTRPGASVL
AVWALIKHLGFEGYREIVRKAMELSRWFAEEIKKLNNAWLVREPMLNIVSFQTKNLRKVERELKRRGWGISAHRGYIRIVFMPHVTKEMVEEFL
RDLREVLK

>T_O58679
MKFPRIGLPKEKVIELINEKTKKDLTFSSGKILGSMCTMPHDLAIEVYTKYIDRNLGDPGLHPGTRKIEEEVIEMISDLLHLEKGHGHIVSGGT
EANILAVRAFRNLSDVEKPELILPKSAHFSFIKAGEMLGVKLVWAELNPDYTVDVRDVEAKISDNTIGIVGIAGTTGLGVVDDIPALSDLARDY
GIPLHVDAAFGGFVIPFAKELGYELPDFDFKLKGVQSITIDPHKMGMAPIPAGGIVFRRKKYLKAISVLAPYLAGGKVWQATITGTRPGASVIA
VWALIKHLGFEGYMRIVERAMKLSRWFAEEIKKINNAWLVREPMLNIVSFQTKNLKKVERELKSRGWGISAHRGYIRIVFMPHVTREMIEEFLK
DLKEVLS
>T_Q0ICY8
MLTPLPKHHFPGEGREGNSTVQLLREELLLDGNSKQNLATFCQTYQAQSAMELMTLGVDKNLIDKDEYPQTAELEGRCVSMMADLWNAPGAAVG
CSTIGSSEAAMLGGMAAKWRWRKRREAAGLPTDKPNMVCGSVQICWKKFARYWDIEMRELEMLTGELCVSPERVLEAVDENTIFVVPTLGVTYH
GLYEDIESISKALDDLQARTGLDVPIHVDAASGGFLAPFCAPDLPLWDFRLERVKSINASGHKFGLAPLGVGWVLWRSQEDLPDELVFHVTYLG
GDMPTFQINFSRPAGQVIAQYHEFVRLGREGYRMLHMASHANAQYFAEKLREMDLFRIIHDGTPDKGIPTVVWTLDDNPKYGFNLYDFADRLRM
RGWQVPAYPFTGELESTAFQRILVKRDFTRDMADLLLEDIRQAIQHFQKHPITSNLAATEGASYNHL
>T_O06249
MSRSHPSVPAHSIAPAYTGRMFTAPVPALRMPDESMDPEAAYRFIHDELMLDGSSRLNLATFVTTWMDPEAEKLMAETFDKNMIDKDEYPATAA
IEARCVSMVADLFHAEGLRDHDPTSATGVSTIGSSEAVMLGGLALKWRWRQRVGSWKGRMPNLVMGSNVQVVWEKFCRYFDVEPRYLPMERGRY
VITPEQVLAAVDENTIGVVAILGTTYTGELEPIAEICAALDKLAAGGGVDVPVHVDAASGGFVVPFLHPDLVWDFRLPRVVSINVSGHKYGLTY
PGVGFVVWRGPEHLPEDLVFRVNYLGGDMPTFTLNFSRPGNQVVGQYYNFLRLGRDGYTKVMQALSHTARWLGDQLREVDHCEVISDGSAIPVV
SFRLAGDRGYTEFDVSHELRTFGWQVPAYTMPDNATDVAVLRIVVREGLSADLARALHDDAVTALAALDKVKPGGHFDAQHFAH
>T_Q737F8
MPQDRKAEVQKHAYERKEIMPDNSQSLPRHMQKELPHEFSVNPLFAREGESVVPRFHISDEGMLPETAYQIVHDEITLDGNARLNLATFVSTWM
EPAAEQLYAKSFDKNMIDKDEYPQTAEIEERCVRILANLWHSPSPLTTMGVSTTGSSEACMLGGLALKRRWQNARKSEGKPLDRPNIVFSSAVQ
VVWEKFANYWEVEPRYVKVSPEHPQLDPQGVLAAVDENTIGVVPILGETYTGLYEPVAEIAKALDDLQARTGLDIPMHVDAASGGFIAPFLQPD
LVWDFQLPRVKSINVSGHKYGLVYPGLGWIIWREAEDLPEDLIFRVSYLGGNMPTFALNFSRPGAQVLLQYYNYLRLGKSGYYDIQRASQKVAL
FLSKAIQKMEPFELLSDGSDIPVFAWRLKEGYTSNWNLYDLSRQLRVFGWQVPAYPLPPDMESVTIMRVVVRNGFSMDLAHLFLRNLKQTVAFL
DSLDGPMPHDTKCNNGFHH
>T_Q82HA9
MTKRDVAALFGNRFLTEPAPSQTFPEEGMTATDTMRLLDEDLVMEGDPQRNLATFVTTWMEPEAQRIIAENLHRNFIDHAEYPISAEIEQRCVR
MLADLFHAPGKTTGCRTQGSSEAIMLGALSLKWKWRERRQAANLPADRPNLVFGGDVHVVWEKFCRYFDVEPRIVPLAEDKYTIGPEDVEPHID
ENTIGVVAVVGTTFTGHKDDVVGIDKLLRDVRKERDLDIPIHVDGASGGFVWPFLYPDSKWDFRLEQVRSINVSGHKYGLVYPGIGWLVFREES
DLAKDLVFYENYLGKTDATFTLNFSTGASMVLAQYYNFVRLGRQGYTYVMETMQKNAHALADNLRSSGRFEVIGSDLEQLPLVAFRLAGEHAYD
ESDIAWQLSAERGWMVPAYTLPPNAERVKILRALVKETLSREQIERLTQDIADACATLDHKGGTTEVERAQIKRGTGY
>T_Q9X8J5
MSLHQGPRPSRDSDRDRRRLAVNPFHAAANPLGGMTEAPPAHRLPDSPLPPESAYRLVHDELMLDGNARLNLATFVTTWMEPQAGVLMSECRDK
NMIDKDEYPRTAELERRCVAMLADLWHAPDPSTAVGCSTTGSSEACMLAGMALKRRWALRNADRYPAKDVRPNLVMGVNVQVCWDKFCNFWEVE
ARQVPMEGDRFHLDPQAAAELCDENTIGVVGILGSTFDGSYEPVADLCAALDALQERTGLDVPVHVDGASGAMVAPFLDEDLVWDFRLPRVASI
NTSGHKYGLVYPGVGWALWRDAEALPEELVFRVNYLGGDMPTFALNFSRPGAQVVAQYYNFLRLGREGYRAVQQSARDIAGSLAERVAALGDFR
LLTRGDQLPVFAFTTADDVTAYDVFDVSRRLREGGWLVPAYTFPPHREDLSVLRVVCRNGFSADMADLLLADLERLLPELRRQPGPLTRDKGAA
TGFHH
>T_Q82EG0
MPLHKGPVKSDERPMSFNPFFGEANPVSGMTEAPPKHRLADGPLPPSTAYQLVHDELMLDGNARLNLATFVTTWMEPQAGVLMAECQDKNMIDK
DEYPRTAELEKRCVAMLADLWNAPDAGAAVGCSTTGSSEACMLAGLALKRRWARRNADRYPARDVRPNLVMGINVQVCWEKFCNFWEVEMRQVP
LEGERYHLDPGAAAELCDENTIGVVGILGSTFDGSYEPIAELCAALDALQERTGLDIPVHVDGASGAMIAPFLDEDLVWDFRLPRVASINTSGH
KYGLVSPGVGWALWRDSEALPEELVFRVNYLGGNMPTFALNFSRPGAQVVAQYYTFLRLGREGFRAVQQTTRNVARSLAERVAALGDFHLLTRG
DELPVFAFTTAPEVASYDVFDVSRRMREHGWLVPAYTFPPNREDLSVLRVVCRNGFSTDLAELFVEDLSRLLPDLRRQARPQTHDKDAATGFHH
>T_Q5AWQ2
MVTLSRVSQESSSHRQRREIRDISVQTTEYAGVYGTKYAAEELPLYVMNDNGMPPDVAEQMIRDELSLDGNPLLNMASFVTTYMEPQVETLMAA
AMRKNFIDFEQYPQSARMQTRCVNMIADLFNAPTNQESKEGTEHGESEGAEGAMGTSTVGSSEAIMLALLAMKKTWHKKRSDAGKDTSHPNIIM
NSAVQVCWEKAARYFDVEERYCYCTDDRYVIDPVQAVELVDENTIGICAIMGTTYTGHYEDVKAINDLLVQRNIDCPIHVDAASGGFVAPFICP
ELVWDFRLEKVVSINVSGHKYGLVYPGVGWIFWRSPEYLPRELVFNINYLGSEQATFTLNFSKGASHIIGQYYQLIRLGRNGYKAIMQNLVQVS
QNLARGLSDLGLLILSDNTGNGSGGVPLVAFRLPDDESRLFDEFAVSAVLRRRGWVVPAYTMAPRSNNLKLMRIVVREDFTAHRCGILIQDIKM
AIEWLEEMDETTIQRYTTYLAQHGTRLPNAHSFYKDEHSLHGKTGKTHAVC
>T_Q5B1Y3
MVHLASIKKDDDFEPVVNRVDSIKLDTIEEDDYSATVYGTRFATQQLPHAEMPDREMPREVAYRMIKDELSLDGNPMLNLASFVTTYMEDEAEK
LMAESFSKNFIDYEEYPQSAEIQNRCVNMIARLFNAPTDSDTDHPMGTSTVGSSEAIMLGTLAMKKRWQNKRKAEGKDYSRPNIVMNSAVQVCW
EKAARYFDVEERYVYCTEERYVIDPQQAVDLVDENTIGICAILGTTYTGEYEDVKAINDLLVERGLDCPIHVDAASGGFVAPFIHPTLQWDFRL
EKVVSINVSGHKYGLVYPGVGWVVWRSPEFLPKELIFNINYLGAEQASFTLNFSKGASHVIGQYYQMIRLGKRGYRSVMVNITRIADYLADQLE
QLGFIIMSQRRGRGLPLVAFRLPADRADETFDEFAIAHQLRERGWIVPAYTMAPHSNNLKLMRVVVREDFSMSRCDQLLSDIKLALKSLREMDQ
AMLERYTQHVRSHTTKSHRAKHTHPHYKNETHSLQGRTGKTHGVC
>T_Q7SCH4
MVHLTIIPKEDEIRDGLDIPLTGALKAVHLQLANDEDRFTTSVYGSKFAAADLPRHEMPDEEMPKEVAYRMIKDELSLDGNPLLNLASFVTTYM
EEEAEKLMTESLPKNFIDYEEYPQTADIQNRCVSMIGRLFNAPVKDAEASSAVGTSSVGSSEAIMLGVLAMKKRWKNKRIAEGKPADKPNLIMS
SAVQVCWEKATRYFEVEEKFVYCTPDRYVIDPKETVDLVDENTIGICCILGTTYTGEYEDVKAVNDLLVERGLDTPIHVDAASGGFVAPFVVPD
LEWDFRLKNVVSINVSGHKYGLVYPGVGWVVWRSAEYLPQELVFNINYLGADQASFTLNFSKGASQVIGQYYQLIRLGKHGYRAIMSNLTRTAD
YLAESLAALGFIIMSQKSGQGLPLVAFRLKEDPDRTYDEFALAHQLRVRGWIVPAYTMAPKTEGLKMLRIVVREDFSRNRCDGLISDIRSCQGI
LEQMDKETVKKQQEFIHKHHVVSGKASHNHPKYHKEKHSLQGKTGKTHSIC
>T_Q0TLK2
MIRTMLQGKLHRVKVTHADLHYEGSCAIDQDFLDAAGILENEAIDIWNVTNGKRFSTYAIAAERGSRIISVNGAAAHCASVGDIVIIASFVNMP
DEEARTWRPNVAYFEGDNEMKRTAKAIPVQVA
>T_Q9HV68
MHAIMLKAKLHRAEVTHAVLDYEGSCAIDGDWLDLSGIREYEQIQIYNIDNGERFTTYAIRAENGSKMISVNGAAAHKAKVGDRVIICAYAHYS
EAELASHKPRMLYMAPGNQLSHTSEAIPIQVA
>T_P52999
MYRTMMSGKLHRATVTEANLNYVGSITIDEDLIDAVGMLPNEKVQIVNNNNGARLETYIIPGKRGSGVICLNGAAARLVQEGDKVIIISYKMMS
DQEAASHEPKVAVLNDQNKIEQMLGNEPARTIL
>T_P9WIL2
MLRTMLKSKIHRATVTCADLHYVGSVTIDADLMDAADLLEGEQVTIVDIDNGARLVTYAITGERGSGVIGINGAAAHLVHPGDLVILIAYATMD
DARARTYQPRIVFVDAYNKPIDMGHDPAFVPENAGELLDPRLGVG

>T_Q72L22
MKRVMFHAKIHRATVTQADLHYVGSVTVDQDLLDAAGILPFEQVDIYDITNGARLTTYALPGERGSGVIGINGAAAHLVKPGDLVILVAYGVFD
EEEARNLKPTVVLVDERNRILEVRKG
>T_P58286
MLRTLIKSKIHRATVTQADLHYVGSVTIDADLLDAADLLPGELVHIVDVTNGARLETYVIEGERGSGVIGINGAAAHLVHPGDLVILISYAQVT
DAEARSLRPRVVHVDGDNRIVGLGADASEPVPGSDQERSPQAVSA
>T_Q18HQ3
MRRWLLKSKLHRARVTGTEKDYEGSISIDAALLSEADIAVGEQVQVVNVTNGERFETYTIEGESRQMELNGAAARLAETGDVIIVISYGLYVKD
EQPEPTVLLLDEENRISERE
>T_O66773
MLREMLKSKIHRLTVTDADLHYEGSLSLDEYLMELADLKPFEKIDVYNINNGARFQTYVIPAPRYSGEVKLNGAAARLGHKGDLIIIASYAQYT
EEELENYAPKLIFVNEKNQPVEVKESTEVK
>T_G4RJM3
MPELLKSKAHGLVVTGKDLHYEGSLTLGRDIMEATGLLPLERVEVYNVTNGARFTTYVIPGEDGQVVLNGAAARLGEVGDILIVASYECAADPT
GHVATVAIFQNNRLKEIKRVSPRDLR
>T_A1RUA9
MPVLLRAKAHGLVVTGKNLHYEGSLTLGRDIIEAAGFYPLEKVEVYNVTNGARFTTYVIPGRPGEVVLNGAAARLGEVGDVIIVAAYECVANPA
SHIATIAIFEGNKLKEVRKISLSDMY
>T_B1YBS3
MPVLLRAKAHGLVVTGKNLWYEGSLTLGRDIMEAAGFYPLERVEVYNVTNGARFSTYVIPGAAGEVVLNGAAARLGEVGDVLIVAAYDCVEDPT
AHVATVAIFQGNRLKEVRKIPTGVP
>T_Q6ZQY3
MSSDSDRQCPVDGDIDQQEMIPSKKNAVLVDGVVLNGPTTDAKAGEKFVEEACRLIMEEVVLKATDVNEKVCEWRPPEQLKQLLDLEMRDSGEP
PHKLLELCRDVIHYSVKTNHPRFFNQLYAGLDYYSLVARFMTEALNPSVYTYEVSPVFLLVEEAVLKKMIEFIGWKEGDGIFNPGGSVSNMYAM
NLARYKYCPDIKEKGLSGSPRLILFTSAECHYSMKKAASFLGIGTENVCFVETDGRGKMIPEELEKQVWQARKEGAAPFLVCATSGTTVLGAFD
PLDEIADICERHSLWLHVDASWGGSALMSRKHRKLLHGIHRADSVAWNPHKMLMAGIQCCALLVKDKSDLLKKCYSAKASYLFQQDKFYDVSYD
TGDKSIQCSRRPDAFKFWMTWKALGTLGLEERVNRALALSRYLVDEIKKREGFKLLMEPEYANICFWYIPPSLREMEEGPEFWAKLNLVAPAIK
ERMMKKGSLMLGYQPHRGKVNFFRQVVISPQVSREDMDFLLDEIDLLGKDM
>N_A0A1Y1VIL9
MVDKENTVTSEDNTKENVLKFFKDYEIREENVQQYIDSIIKDFYKPSPSIYASNNDSVKEIQSKFGDSSIPEGIICDKNNINKYFSEIKENVID
KATRVSSPNMVGHMTTALPFFHRPLSKLVTSMNQNVVKVETSATTTFLERETLSKLHRAFFNQSPEFYEKYLSITEGTFGCITSGGTIANLTAM
WVARNKAFPKTDSFDGIEKEGVISALRYYGYNDMAIVGSELMHYSFKKAADVLGMGLKNIYTIPVDSEYGIRIDLLIEKLEELKRKKIKVIAIV
GIACTTEVGSIDNLEVLADLAKKYEALFHVDGAWGGSFIMSEKYRYLLKGIELADTVTIDGHKLLYTPMGCGIVLFKSPLLPTTTIRKVASYVI
RAGSMDHGKTTLEGSRPANVLYLHASLNLLGHRGLSSLLDIAVENTNYMAELVKSYPQFEIISKPITNIFLYRYIPSWLRKDNEQFEVRYPVSP
NSKTIEIEEVSTLYSNSTQDQTFDISDLTDKEENSFVEEEPSQYTNEENQVIDYFTTELQAKQKREGKGFVSKTTVKTTKYPNYEKGIDVLRVV
IANPLVTHENIKAVVEEQIRLSQELEKEYKMKTQSI
>N_A0A1Y1WT12
MDNYNNEIINDIEEYVKQDINKNDSSHDWHHILRVKNLSIAITKGELENGKKVNLFNVTAISLLHDSIDSKYCQNIEDKIKEIRKFLISKQIKE
EDIDQILNGINNISYRKEIGRSPEERKVPLEIAIVQDADKLDAMGAIGISRCFAYSGAKGRPFYDPEIQPKINMSQEEYVAQSNSNQGTAINHF
YEKLFNLKYMMKTDYGKIMAIERDRYMREFVDRFIKEYNVNNSNNSIYDDDSNHNNNKHLKQKKRQSILEYDYCKEHLLNFDKRNKTEVVSLAY
GSRFATENLPKYIIPEKSSEPKVISQLIEDQIKIEANPSQNMATFVSTWMEPECEKLMNQAWSKNFADQDCYPMIQNIHKRCIEMLGHLFHAPQ
SHPFPQSSSSQNNHQVIGTATTGSSEAVMLGGLALKWRWKKIRKEKGLDDSKPNIIFGSNAQVALEKFARYFDVEMRMVPVDESTNFCLSPQRA
IKYVDENTIGIMVILGSTYTGHFEPVEEMCKVLDDYQEKTGIDIPIHVDAAGGGFIAPFAFPDLKWSFELNRVHSINVSGHKYGLVYPGIGWIV
WKSKDYLPKELIFQLNYLGSVEYTFTLNFSRTSTTVLGQYYNFLRLGYEGYTSIINNCLINARRLAYALNDLDYFNILSDVNKKYKNSYFSEQD
PNHDDKDNNQEYFQPCLPVVAFEIKQNYKEYQPHVTEANLSKLLKIHGWIVPCYELPPNEQNRTILRIVIRESHSEELINYLFKNIHQSIEDLI
DGKDYNLEKRKKRTSSMNYINEQNSLNLEKENKEMFDTKTKWGVC
>N_A0A1Y1XDI3
MIVLKFFKDCEIQEENVQQYIDNIIKEFYKPSPSIYAANTDTDEELKSKFGDSSIPQGIICDKNNINKYFTEIKETVIDKATRVSSPNMVGHMT
SALPFFHRPLSKLITSLNQNVVKVETSSTMTFLERETLSKLHRAFYNESDEFYEKYLSTTDGSFGCVTSGGTIANLTAMWVARNNAFPKTNDFE
GIEKEGVISALKYYGYNDMAIVGSELMHYSFKKAADILGMGLKNIYTIPVDSEYRIRIDLLIKKLEELRNKKIKVIAIVGIACTTEVGSIDNLE
ILSDLAKKYGALFHVDGAWGGSFIMSDKYQHLLKGIEKADTITIDGHKLLYTPMGCGIVLFKSPLLPTTTIRKVASYVIRADSMDHGKTTLEGS
RPANVLYLHASLNLLGHDGLSSILDIAVENTKYMADLVKTYPQFEIISYPVTNIFLYRYIPSWLRKKPINEHIEQMVEIKSNATARTFVANSED
NNEGTNDLSTLCSDTIQDLSDISDKEECSISYDESLYNEEENKIIDYFTTELQAKQKREGKGFVSKTTVKTTKYPNYRNGVAVFRVVIANPLVT
SENIKGVIEEQIKLGEIIEEDYKINHKTLL
>N_A0A1Y1ZL74
MVDKETIKTEENNTKENVLKFFKDCEINEENVQQYIDSIIKEFYKPSPSIYAANTDTVKELQSKFGDSSIPKGIVCDKSNIKNYFSEIKTNVID
KSTRVSSPNMVGHMTSALPFFHRPLSKLITSLNQNVVKVETSATMTFLERETLSKLHRAFYNNTDEFYEKYLSTTEGTFGCVTSGGTIANLTAM
WVARNNAFPKTEDFEGIEKEGVISALSYYGYNNMAIVGSELMHYSFKKAADILGMGLKNIYTIPVDSDYKMRIDLLIEKLEELKQKKIKVIAIV
GIACTTEVGSIDKLDILADIAKKYGALFHVDGAWGGSFIMSNKYHHLLKGIEKADTITIDGHKLLYTPMGCGIVLFKSPLLPTTTIRKVASYVI
RAGSMDHGKTTLEGSRPANVLYLHASLNLLGENGLSQLLDIGVNNAKYMAELIKSYPQFELISKPVTNIFLYRYIPSWLRKKSYEIEQIEVKSI
SSPISTVSTIPSPTQATTALPIIESEEENHNDNDNSITFCNDTLKDQISDISDFTDKEGSLIYEESHYNDEENAIIDYFTTELQARQKREGEGF
VSKTTVKTTKYPTYSKGVDVLRVVIANPLVTEDNIKAVVEEQIKLGEIIEEEYKMNHKLIK
>N_A0A1Y2AA19
MIVEEKLIQRLIIVNLMKRVFLNLFENDIQEVQMYIDKIIKDFYKPSPSVYTINSNSEKELMLKFGNSTIPKGISYDKKDIINYFFKIKTNVVE
NATRVSSPYMMGHMTSALPFFHKHISKLISSLNQNVVKVETSSTMTFLERETLSKLHRAFYNMEDNYYKELLTKNDGCFGCITSGGTIANLTAL
WISRNNAFPKTDNFNGIEKEGIISALKYYNYNDIAIIGSELMHYSFKKAADILGIGLNKIYTIPVDSEYKIRIDLLTEKLQELKQKRIKVITIV
GIASTTEVGSIDDLEKLADLAEEYGAMFHVDGAWGGSFILSDKYRHLLKGIQRADTITIDGHKLLYTPIGCGVVLFKSPFLPTTTIRKVANYII
KTDSMDHGKTTLEGSRPANALYIHASLNILGQHGLRYLLETSIENTKYMYNLINLYP
>N_A0A1Y2ADH1
MVDKDNIKSKENNAEEDVLKFFKDYGIQEENVQQYIDNIIKEFYKPSRSIYAANNDTIQELQSKFGNSSMPQGIICDKKNINKYFTELKTNVIN
KSTRVSSPNMVGHMTTALPFFHRPLSKLITSMNQNVVKVETSATMTFLERETLSKLHRAFYDNTNDFYEKYLSSTEGTFGCITSGGTIANLTAM
WVARNNAFPKTDDFEGIEKEGVISALRYYGYNDMAIVGSDLMHYSFKKAADILGMGLKNIYTIPVDSDYKMRIDLLIEKLEELKHKKIKVIAIV
GIACTTEVGSIDRLDILADLAKKYGALFHVDGAWGGSFIMSNKYRHLLKGIEKADTITIDGHKLLYTPIGCGIVLFKSPLLPTTTIRKVASYVI
RAESLDHGKTTLEGSRPANVLYLHASLNLLGKDGLSQLLDIAVNNTKYMADLVKSYPQFEIISKPVTNIFLYRYIPIWLRKKSYDIEQIEVKSL
SSAVFTNSSIPSPTPYNTTHPPLTFEENNNDKSDNKNENSITLCAGFVSKTTVKTTKYPTYTKGVAVLRVVIANPLVTQDNIKAIIEEQIKLGQ
ILEEEYKINHKVI
>N_A0A1Y2AZ59

MKQKRQSILDNDFYKDQILDIDKNNTEIVSLAYGSRFATENLPKNVIPEKSSEPKVISQLIEDQIKIEANPSQNMATFVSTWMEPECEKLMSQA
WSKNFADQDCYPMIQNIHKRCIGMLGHLFHAPDDQCVIGTATTGSSEAVMLGGLALKWRWRKKREEKGLDCSKPNIVFGSNAQVALEKFARYFD
VEVRMVPVDEETHFCLNPQRAIKYIDENTIGIMVIILGSTYTGHFEPVEEMCKILDDYQRKTGIDIPIHVDAASGGFIAPFAFPELKWSFELDR
VHSINVSGHKFGLVYPGIGWILWKSQEFLPKDLIFRLNYLGSVEYTFTLNFSRTSTTVIGQYYNFLRLGYEGYTSIINNCLINARRLALALEDL
DYFNILCDVNKKINMDNNSNSLIKGFKPCLPVIAFEIKKKYKKNKPYVTEANLSKLLKIHGWIVPCYDLPPNEQNRTILRIVVSNTNDFFIKKF
FLNFFFFFFFL
>N_A0A1Y2B2H7
MLKFGNSTIPKGISYDKKDIINYFFKIKTNVVENATRVSSPYMMGHMTSALPFFHKHISKLISSLNQNVVKVETSSTMTFLERETLSKLHRAFY
NMEDNYYKELLTKNDGCFGCITSGGTIANLTALWISRNNAFPKTDNFNGIEKEGIISALKYYNYNDIAIIGSELMHYSFKKAADILGIGLNKIY
TIPVDSEYKIRIDLLTEKLQELKQKRIKVITIVGIASTTEVGSIDDLEKLADLAEEYGAMFHVDGAWGGSFILSDKYRHLLKGIQRADTITIDG
HKLLYTPIGCGVVLFKSPFLPTTTIRKVANYIIKTDSMDHGKTTLEGSRPANALYIHASLNILGQHGLRYLLETSIENTKYMYNLINLYPQFEI
ISKPITNIFTYRYIPTWLQNKNIQIKNKNNDINSSFYTTNNNNNNNNNSNNNNNNKIYLEKYNNSDNAYERINDVTFNNEESEIIDDERLRKGFV
SKTIIKTTKYPNCTNGVAVLRVVIANPLITKESIKTVIEEQIEFGLKLEKEYCN
>N_A0A1Y3MYE4
MYCHGNTSALPFFQRPLSKLVTSLNQNVVKIETSSTMTFLEKEVLIKLHKLFFNEINAFYEKCITNNNKTFGCITSGGTVANLTAMWVARNNAF
PKTEDFEGIEKEGVISALKYYEYNDMAIVGSELMHYSFRKAADILGLGLNNIYSIPTDSEYKMRTDLLIKKLEELKLKRIKVIALIGIACTTEV
GSIDNLEVLAEIAKKYNIFFHVDGAWGGSFIISDKYKYLLKGIEKADSITIDGHKLLYTPMGCGVILFKSPYIASKTIKKVASYIIRENSMDHG
KTTLEGSRPANILYLHASLTLLGQHGLSTLLDMAIYNTKYMASLKCNYKIDKQSDPIFKDNLNKNINYYNLSNENNSKLNNSLYSKEENQIISY
FTNELQFRQKKYGKGFVSKTTIKSAKYSNCKDGISVFRVVIANPLVTKENIKFVIEEQIKFGLDLEKEKSNFIN
>N_A0A1Y3NC87
MNTILNFSSKYNSKKHPKLNSTTKKKRQSILNNDFYKDEFSNSQKNGSEIVSLAYGSRFSTENLPKYVIPQKSSEPKVISQLIEDQIKIEANPS
QNMATFVSTWMEPECENLMTQTWSKNFADQDCYPMIQNIHKRCIGMLGHLFHAPDDQEVIGTATTGSSEAVMLGGLALKWRWRKARMEKNLDCS
KPNIVFGSNAQVALEKFARYFDVEMRMVPVNEKTNFCLSPKRALKYIDENTIGVMVKLKK
>N_A0A1Y3NIR5
MWVARNNAFPKTDDFEGIEKEGVVSALRYYGYNDMAIVGSELMHYSFKKAADILGMGLKNIYTIPVDSEYGISGSIDNLEVLADLAKKYEALFH
VDGAWGGSFIMSDKYHHLLKGIEKADTITIDGN

**Supplementary File 2: Phylogenetic tree.** Provided in Newick format and used to draw Fig. 2 in the main text.

(((N_A0A1Y2ADH1:0.116131,N_A0A1Y1ZL74:0.064668)68:0.025074,(((E_A0A1S8W5A4:0.232462,E_F4NWP2:
0.204214)100:0.226855,E_A0A0L0HIP1:0.358706)100:0.221427,((((((B_Q2LV45:0.637131,((B_W0V410:0
.163894,B_A0A1I4S397:0.165029)100:0.166615,B_A0A254TI57:0.259658)100:0.291103)73:0.103885,((((
((((B_A0A1H2SAR8:0.363236,B_A0A1I2AY29:0.362688)32:0.054236,((B_K9U459:0.440050,B_K9Z2H8:0.33
3704)37:0.061388,B_A0A1I4JUD5:0.331214)20:0.061272)8:0.031513,(B_A0A0H4P273:0.423551,B_A0A0V8J
BT8:0.326687)93:0.119154)23:0.044369,(A_A0A1H7UJE7:0.286706,(((A_D4GP47:0.172082,A_I3R9H1:0.15
8133)100:0.200654,(((A_A0A1I4CXR5:0.254228,((A_E4NVT4:0.320556,(A_A0A1M5JPP3:0.124009,A_Q9HHV3
:0.252312)98:0.091098)67:0.045724,A_W0JRZ3:0.289616)100:0.115054)83:0.075483,((A_A0A1I0QDE1:0.
093358,(A_D2RU46:0.109511,(A_L0JPS5:0.111283,(A_I7CDV7:0.088035,A_A0A1H9J813:0.099306)41:0.022
164)21:0.030921)38:0.032729)100:0.205044,A_C7P526:0.265166)100:0.115022)26:0.036071,((A_A0A1G9
UVH9:0.192976,A_A0A1Q1FJ77:0.153183)67:0.043561,(A_D8JB87:0.163395,A_A0A1H1C830:0.089004)62:0.
034585)100:0.132556)68:0.074253)43:0.046219,A_A0A1N6W5S0:0.241239)76:0.071668)100:0.206724)90:
0.149403,A_A0A0X8V205:0.611248)99:0.243211,(((E_A0A016U7N4:0.785907,(((E_A0A138ZZL4:0.479704,(
(E_U4UGN6:0.433484,((E_G1N6G5:0.014767,(E_F1NXM1:0.000001,E_A0A1D5PSZ5:0.013288)99:0.016890)90
:0.012548,(((E_A0A0A0AJ64:0.023496,((((E_H0ZB55:0.010524,E_U3K9Z6:0.014238)90:0.005813,E_A0A0
91F3Y5:0.011966)94:0.031773,(E_A0A2I0MD22:0.005688,E_A0A1V4JEJ7:0.011245)70:0.016488)23:0.0036
82,(E_A0A0Q3PWB0:0.042340,E_A0A093GDE1:0.027837)32:0.004317)2:0.001471,((E_A0A099Z5N3:0.057235
,E_A0A093HSM0:0.020137)91:0.027774,(E_A0A087QWQ6:0.011618,(E_A0A091VHQ0:0.012677,(E_A0A091JKB2
:0.017176,E_A0A091FVI7:0.023667)44:0.001437)33:0.001618)32:0.001613)23:0.002839)4:0.001411)12:
0.001715,E_A0A091WHQ1:0.025943)45:0.012481,E_U3I574:0.027607)47:0.015341)100:0.462497)77:0.096
661,((E_A0A0V1H7S9:0.004551,(E_A0A0V1MFM8:0.008014,E_A0A0V0W3Q4:0.032654)41:0.001506)100:0.370
684,E_F6ZTZ7:0.477059)69:0.081217)93:0.104131)44:0.076342,((((E_A0A1U8A7E6:0.000001,E_A0A1U8A
5K9:0.000001)100:0.092475,(E_A0A200PVP3:0.121648,E_A0A2H5NKN4:0.119016)45:0.026770)100:0.13462
5,(E_D8R3Z6:0.145207,E_D8SSC2:0.349613)100:0.114878)18:0.043660,(E_A9SPU0:0.002127,E_A9SPU5:0.
005410)100:0.336773)38:0.060252,E_A0A251V0E5:0.472211)100:0.198436)39:0.052574,(((E_A0A370TBI7
:0.072246,(((E_A0A0C3I054:0.126266,E_A0A1L7XGT1:0.077302)44:0.024938,((E_K1X4T2:0.041449,E_A0A
218Z6N7:0.082549)95:0.028060,E_A0A2V1BNR7:0.079907)80:0.024232)33:0.019305,E_A0A2T3AUC5:0.0893
41)67:0.039151)100:0.203003,E_G1X4X9:0.333727)100:0.301565,E_A0A137PEN3:0.586318)37:0.066265)8
5:0.139533)100:0.367929,((((B_A0A1H4BYW3:0.141101,B_A0A1G9K0B9:0.151777)86:0.106159,B_A0A1M4ZH
C6:0.230758)100:0.458565,A_I3R7R5:0.611544)49:0.065969,A_F4BYV2:0.668824)86:0.111277)41:0.0701
91,E_C3Y5S8:0.991473)57:0.108322)9:0.044143,((((A_W0JV70:0.055545,A_I3R6X8:0.075417)68:0.04137
3,A_G2MMT9:0.136829)100:0.391984,B_G2PQR2:0.408077)100:0.289452,(((((((E_G1TK30:0.078909,E_H0V
9C5:0.078886)100:0.187421,T_Q6ZQY3:0.293026)99:0.099502,((((E_Q24062:0.006586,(E_B4Q567:0.0016
09,E_B4HXA1:0.000001)100:0.006439)100:0.339112,(T_A7U8C7:0.223068,T_Q171S0:0.211210)91:0.07730
6)69:0.059858,(E_A0A0N0BFM6:0.063123,E_A0A0L7RAZ6:0.103715)100:0.218273)80:0.064680,E_U4TX00:0
.493901)100:0.125970)75:0.056912,(E_A0A151WUY7:0.601863,E_T1KV43:0.475835)63:0.139837)39:0.057
969,(E_A0A369RT79:0.443588,(((T_K7XPX5:0.245564,T_K7WYY0:0.257451)100:0.125856,(E_A0A0V0YNL2:0
.000001,E_A0A0V0YMZ9:0.000001)100:0.313934)85:0.063932,((((E_I3KQZ2:0.011935,((E_H3CR31:0.0254
98,E_A0A2U9BKG5:0.023856)61:0.007414,E_A0A087YBQ9:0.029143)77:0.007446)99:0.057078,(E_F6NX32:0
.001626,E_E7FDZ2:0.000001)100:0.106772)99:0.040227,((((T_Q548L6:0.012911,E_H0V2I4:0.017140)67:
0.001927,T_Q99259:0.010638)99:0.032106,E_H9GNM5:0.040818)95:0.016471,E_H3AXV6:0.054001)96:0.03
3439)100:0.121760,(T_Q05329:0.019843,T_P48320:0.022178)100:0.266705)100:0.120518)98:0.122962)8
5:0.067308)87:0.186635,E_A0A090MC16:0.649276)96:0.151038,A_A0A2E4TVH8:0.640099)100:0.245186)61
:0.100568)29:0.051668,((B_F4KV58:0.314909,B_A0A1M6AT35:0.358907)49:0.063337,B_F0RIL5:0.375451)
100:0.451017)32:0.083145,((E_A0A179G349:0.505006,E_I1S8G3:0.619538)100:0.525424,((((E_A0A0D2IJ
V6:0.239542,E_A0A0D2EPM7:0.219966)81:0.084024,E_A0A0D1XU64:0.252593)99:0.156933,(((E_A0A1J9PGB
1:0.090108,(E_A0A2B7Z7I8:0.042027,(E_A0A1J9RHD0:0.055230,E_A0A2B7X0N5:0.055408)99:0.062440)64:
0.031742)100:0.234057,((E_A0A1L9V1T0:0.024737,((E_G7XA98:0.025918,(E_A0A1L9N0S7:0.009449,E_A0A
124BXG1:0.003815)74:0.003054)100:0.025994,E_A0A319APA8:0.043429)45:0.013861)100:0.058773,(E_A0
A319ELX8:0.132570,(E_A0A319ETC9:0.025217,E_A0A1R3RWV0:0.038984)82:0.021194)60:0.031978)100:0.2
32410)89:0.095673,(((((E_B2W8T8:0.096948,E_A0A177DW43:0.117259)40:0.013233,(((E_W6Z9E5:0.02285
0,E_W6Y6T7:0.014968)62:0.001771,E_M2TJG6:0.017009)100:0.051736,(E_R0IYN0:0.075684,E_A0A364MZ99
:0.070386)28:0.006005)80:0.023996)99:0.091023,E_Q0U7M3:0.168103)100:0.138103,E_R7Z469:0.325912
)99:0.100045,((E_F7VTW6:0.081933,E_F5HAE7:0.048642)100:0.377845,E_A0A218ZBM9:0.290542)100:0.22
1839)54:0.051691)87:0.117041)100:0.815651,(((((A_A0A1D2R9X5:0.501354,A_B5ICZ4:0.522270)46:0.0
69465,((((T_Q6L2R7:0.000001,A_Q6L2R7:0.000001)100:0.287897,A_S0AU28:0.243404)100:0.311981,(T_Q
4J9W7:0.379090,T_O28946:0.270306)97:0.241500)98:0.621198,(A_A2BJD5:1.078863,(A_B8D379:0.571118
,A_I3TCG1:0.513881)58:0.208923)87:0.191613)41:0.133817)15:0.057538,((((A_D5VUB3:0.242753,(T_Q6
0358:0.000001,A_Q60358:0.000001)100:0.160320)100:0.102935,(A_F6BEM5:0.085712,(A_D7DV28:0.25608
1,((T_Q6M0Y7:0.000001,A_Q6M0Y7:0.000001)100:0.180705,A_A6UVR4:0.188909)59:0.049936)96:0.085900
)72:0.055689)100:0.237567,(((A_A0A328S773:0.182577,(((A_A0A328SS11:0.072180,A_A0A2Z4L867:0.067
961)100:0.184823,(A_A0A328RY55:0.180790,A_Q2NHY7:0.111131)90:0.058571)71:0.059478,A_A0A328SMU3
:0.171226)60:0.067284)37:0.035334,A_A0A328SBH2:0.193921)100:0.309050,(((((T_Q9UZD5:0.062150,T_
O58679:0.052888)72:0.029371,(T_Q8U1P6:0.000001,A_Q8U1P6:0.000001)100:0.089436)91:0.069628,((A_
B6YUX2:0.063825,(T_Q5JJ82:0.000001,A_Q5JJ82:0.000001)100:0.106518)79:0.030612,(A_W0I930:0.0630
61,(A_A0A075LW21:0.118179,A_C6A1N1:0.088772)97:0.061105)90:0.052188)91:0.063453)99:0.321515,A_
E3GX95:0.240130)34:0.057319,(((A_A0A1D3L4E9:0.139158,A_F0T911:0.203025)100:0.070544,A_U6EEL6:0
.162662)100:0.112218,((A_D3DZR8:0.225281,((A_A0A165Z3S9:0.146338,A_A0A0U2V5X1:0.150585)99:0.09
4402,A_R9SLT3:0.242882)60:0.060365)100:0.113250,T_O27188:0.234840)48:0.056565)68:0.062157)24:0
.045298)44:0.057481)24:0.059040,((A_D2RH62:0.193554,((A_A0A0F7IJ10:0.152044,((A_N0BK04:0.16039

4,A_O28275:0.125227)99:0.068951,A_F2KP03:0.171430)52:0.039787)76:0.069690,A_D3RZ59:0.117659)96
:0.074912)98:0.115782,((((A_A0A2V3JDE0:0.225949,A_A0A1F2P723:0.322607)90:0.096621,(((A_G7WKU8:
0.256364,T_A0B9M9:0.206481)49:0.037859,A_F4C0R3:0.208662)100:0.144282,(((((((T_Q8PXA5:0.00000
1,A_Q8PXA5:0.000001)100:0.061392,A_Q8TUQ9:0.064441)100:0.063423,A_Q46DU3:0.073941)100:0.096569
,A_A0A0E3P0J0:0.120240)100:0.135550,(A_F7XL45:0.241928,A_D7EBV8:0.231017)52:0.075802)37:0.0504
88,T_Q12VA2:0.286461)9:0.028904,(A_K4MEZ9:0.283961,A_L0KZR1:0.291289)47:0.079670)67:0.067568,A
_A0A2V3JKE4:0.327464)52:0.060233)53:0.054290)87:0.057732,(A_Q0W498:0.192318,A_D1YVJ9:0.212418)
100:0.249004)33:0.041124,((A_A0A0F7PCV2:0.230735,((A_A0A1J1ACR1:0.353392,((T_Q9HSA3:0.000001,A
_Q9HSA3:0.000001)100:0.118970,(A_A0A0U5H3X9:0.076194,A_W0K0Y4:0.098164)98:0.050914)98:0.062461
)29:0.035386,((A_A0A1H3WVB1:0.180708,A_U1QCQ6:0.354701)15:0.048638,(A_A0A1N6VH17:0.189059,((((
A_L0ALZ0:0.069505,A_M0L1G1:0.124790)99:0.066173,((A_W0JJ99:0.120419,(A_D2RX89:0.052269,(((A_A0
A346PMZ5:0.097462,A_A0A2Z2HQZ6:0.064733)43:0.035726,(A_L0JRU9:0.053617,(A_A0A1H9Q7I0:0.070486,
A_A0A1I0Q4I5:0.032672)45:0.012360)49:0.021351)8:0.007346,A_F8D376:0.047074)16:0.025407)32:0.03
5081)41:0.020606,A_L0JVZ3:0.093515)48:0.040690)51:0.037448,A_D3SYW6:0.131487)81:0.087660,((A_Q
3IT46:0.041940,A_M1XLA6:0.122606)99:0.152741,((A_A0A1Q1FH74:0.154017,(A_F7PNC6:0.125687,((A_Q5
V1B4:0.106043,A_C7P1L0:0.086453)71:0.032680,(A_A0A1I6LFT2:0.044899,A_A0A1G8UBX3:0.143265)51:0.
027631)49:0.037662)53:0.035896)64:0.048387,A_A0A2R4X059:0.216134)61:0.032532)39:0.048577)19:0.
026629)10:0.025575)29:0.050210)27:0.043756)96:0.497193,((A_H1YYJ3:0.162570,A_E1RFW3:0.217228)1
00:0.107523,((A_A0A0X3BK81:0.004208,A_I7LLL5:0.017746)100:0.195939,((T_Q2FSD2:0.000001,A_Q2FSD
2:0.000001)100:0.363903,A_A2STQ3:0.281329)27:0.043855)46:0.065383)100:0.190889)47:0.081214)46:
0.036422)82:0.142788)16:0.056635)29:0.091846,T_Q8TV92:0.552644)78:0.149915,((T_Q04792:0.660169
,((N_A0A1Y3NC87:0.135952,N_A0A1Y2AZ59:0.059655)89:0.068704,N_A0A1Y1WT12:0.103333)100:0.350839)
95:0.128898,((((T_Q737F8:0.343392,(T_Q82EG0:0.093301,T_Q9X8J5:0.080394)99:0.312388)77:0.094090
,((T_Q0ICY8:0.308534,(T_A0A454A444:0.000594,T_A0A454A8X6:0.009589)100:0.325256)100:0.147204,T_
K4HXK6:0.536042)63:0.061426)41:0.046481,T_O06249:0.359522)85:0.103082,(((T_Q7SCH4:0.175913,(T_
Q5AWQ2:0.312036,T_Q5B1Y3:0.139514)75:0.069304)100:0.238610,(T_A0EJ89:0.129702,T_Q42472:0.14119
7)100:0.263298)78:0.099652,(T_Q82HA9:0.465540,(A_A0A1D3KZK6:0.122948,A_U6EDB6:0.188878)100:0.1
75414)72:0.091026)67:0.071225)62:0.076104)97:0.613128)82:0.287829,(((T_Q9HV68:0.256668,(T_Q0TL
K2:0.232583,((T_Q18HQ3:0.508524,(T_O66773:0.303966,(T_G4RJM3:0.022657,(T_A1RUA9:0.115330,T_B1Y
BS3:0.094351)98:0.173054)99:0.475189)63:0.108551)26:0.115569,(T_Q72L22:0.192948,((T_P9WIL2:0.2
66265,T_P58286:0.208491)74:0.079113,T_P52999:0.450309)25:0.086933)48:0.115216)58:0.256185)40:0
.135804)100:0.998020,A_A0A062V1G2:0.782861)52:0.367466,(A_A2SSB4:0.627176,E_A0A2B4SYD9:0.63616
9)68:0.275294)71:0.208411)72:0.068010)40:0.088597)10:0.051386)98:0.231457)62:0.049416,((B_A8ZV
T2:0.320795,(((B_A0A1W2CRD4:0.183595,B_A0A1W1H5C0:0.199570)54:0.047813,B_C0QAM8:0.201920)95:0.
071972,(B_K0NEB9:0.183367,B_I5B6M9:0.221602)46:0.054319)35:0.048539)41:0.064833,B_A0A1G5GHE0:0
.314358)98:0.328012)39:0.042292,(((((B_Q2SN93:0.208394,(B_A0A1Y0IE65:0.179683,((B_A0A1I6JRS3:0
.076710,(B_A0A1H2Q464:0.071504,B_N6WX51:0.119317)38:0.023462)40:0.018185,(B_A0A1I3QNC8:0.08023
8,A_A0A2E4GBW7:0.090085)53:0.021831)100:0.117105)69:0.039436)100:0.098538,((B_A0A0S2KBH9:0.214
051,B_A0A0F7JX37:0.242252)57:0.057567,((B_A0A1G6Z257:0.140083,B_Q1K434:0.125976)96:0.054035,(B
_A0A0B5FTX7:0.145619,(((B_B3E5L1:0.107493,((B_Q74CG6:0.045748,B_Q39V49:0.035150)99:0.042120,(B
_A1AMJ0:0.118661,B_B9M3A1:0.072893)86:0.021419)96:0.041045)93:0.046181,B_A0A0M4DHR6:0.118978)9
9:0.064848,B_A0A1H3YWG2:0.195962)80:0.040571)52:0.020224)100:0.118577)69:0.036020)97:0.059267,
((((((B_A3QG03:0.095965,(B_D4ZAE7:0.057537,(B_A8H648:0.088847,B_A8FX14:0.075010)40:0.022009)1
00:0.083689)94:0.045506,(((B_Q12LG2:0.097941,B_Q07ZT1:0.105265)81:0.030593,B_Q8EG41:0.081586)8
8:0.043000,(B_A1S4V1:0.178963,B_A0A1S2TUI8:0.119172)100:0.067558)84:0.037877)100:0.063606,(B_A
0A1H3XGS2:0.091555,B_A0A1E7Q4B7:0.123574)97:0.071802)97:0.050056,((B_A0A0F4QH65:0.075005,(B_A0
A0S2K4U2:0.072954,B_A0A0F4P522:0.055865)67:0.025776)97:0.048076,(B_A0A1I1P8H3:0.109200,B_A0A24
4CNG4:0.064856)86:0.032895)100:0.119642)90:0.055114,((B_A0A1G6DFW1:0.108732,B_A0A1Y6G133:0.100
781)74:0.043775,B_A3WL97:0.120353)100:0.202397)26:0.023247,(((B_A0A0C5WRF6:0.112139,B_Q6LS17:0
.107179)100:0.073360,((B_U4K3X4:0.121406,((B_A0A1G7ZM35:0.140887,(B_Q87QB3:0.095964,B_Q9KSV7:0
.111760)34:0.022970)30:0.018045,(B_B7VPR7:0.083135,B_A0A1E5CWM7:0.107115)60:0.034537)89:0.0306
45)93:0.052865,B_Q5E6F9:0.158662)93:0.064994)97:0.071326,((B_Q487K9:0.137615,B_A0A0D8CQ70:0.10
8062)100:0.078824,B_A0A1I0GM92:0.234390)68:0.043376)37:0.035367)10:0.024468,B_A0A0J8JNG0:0.275
108)25:0.018890)34:0.057197,(((((B_A0A075P054:0.131304,B_A0A1E7Z8S9:0.112356)73:0.036863,(B_A0
A1E8FJP1:0.069062,B_A0A1M5I3T2:0.081314)97:0.045928)91:0.050729,(B_G4QDQ4:0.039743,B_K6ZXH3:0.
040230)100:0.165301)95:0.056908,B_Q15NV7:0.153533)95:0.070704,B_A0A0U2ZFL7:0.230225)100:0.1579
99)94:0.096005,(((B_Q1LR80:0.103175,B_A0A0C4Y9E7:0.085945)100:0.250877,(B_A0A068QSK8:0.044479,
B_A0A068QSK3:0.045066)100:0.286963)99:0.110031,B_U4K047:0.528417)90:0.075563)100:0.150837)97:0
.101247,(((B_A0A1K1QNC9:0.488167,B_A0A127VCS8:0.503612)100:0.207843,B_A0A0C5WU90:0.483918)95:0
.082762,B_Q4BVC3:0.475823)91:0.135511)57:0.058555,(((B_E3I3F0:0.361412,B_W0PGU5:0.322398)95:0.
112173,B_K8W5U2:0.428800)99:0.118238,B_A0KNW7:0.441893)100:0.166481)31:0.033142,((B_K9VN97:0.2
80275,(B_K9TK94:0.268529,((B_A0A0C1NBK7:0.172298,(B_A0A139WW23:0.072757,B_A0A0C2M9Q6:0.077029)
99:0.064972)100:0.107417,B_A0A1Z3HL75:0.244658)86:0.065379)66:0.048629)92:0.082507,B_B8I983:0.
334475)100:0.146119)100:0.223114)100:0.340045)38:0.033800,(N_A0A1Y3NIR5:0.031370,N_A0A1Y1VIL9:
0.086909)88:0.032441)21:0.030709,((N_A0A1Y2B2H7:0.000001,N_A0A1Y2AA19:0.000001)100:0.268806,N_
A0A1Y3MYE4:0.253520)66:0.097090,N_A0A1Y1XDI3:0.092851)21:0.0;

**Supplementary File 3: Multiple sequence alignment.** Selected L-Aspartate decarboxylases are compared against known bacterial PanDs to show very little sequence conservation.

```
CLUSTAL O(1.2.4) multiple sequence alignment

sp|Q0TLK2|PAND_ECOL5                    ------------------------------------------------------------      0
sp|P9WIL2|PAND_MYCTO                    ------------------------------------------------------------      0
tr|A7U8C7|A7U8C7_TRICA                  ------------------------------------------------------------      0
tr|B8I983|B8I983_CLOCE                  ------------------------------------------------------------      0
tr|K9TK94|K9TK94_9CYAN                  ------------------------------------------------------------      0
tr|A0A1Y1ZL74|A0A1Y1ZL74_9FUNG          ------------------------------------------------------------      0
tr|A0A1Y2AA19|A0A1Y2AA19_9FUNG          ------------------------------------------------------------      0
tr|F4NWP2|F4NWP2_BATDJ                  -------------------------------------------MQTTAHI--STAEASLH      15
tr|A0A1S8W5A4|A0A1S8W5A4_9FUNG          MDGPKCKQEPATNTDTTIVSTTTVNTTAVSTTAATTAATTTTSTDGTGRIVAATPMSRLA      60

sp|Q0TLK2|PAND_ECOL5                    ------------------------------------------------------------      0
sp|P9WIL2|PAND_MYCTO                    ------------------------------------------------------------      0
tr|A7U8C7|A7U8C7_TRICA                  -------------------------------MPATGEDQDLVQ----------DLIEEPA      19
tr|B8I983|B8I983_CLOCE                  ------------------------------------------------------------      0
tr|K9TK94|K9TK94_9CYAN                  ------------------------------------------------------------      0
tr|A0A1Y1ZL74|A0A1Y1ZL74_9FUNG          ------------------------------------------------------------      0
tr|A0A1Y2AA19|A0A1Y2AA19_9FUNG          ------------------------------------------------------------      0
tr|F4NWP2|F4NWP2_BATDJ                  INHKDTDEGF-EDDYPEEFPFWYSFPSGEQHIDRHQQ----------------KLLFESS      58
tr|A0A1S8W5A4|A0A1S8W5A4_9FUNG          MVDADIDEGFHDDDYPEEFPFWYGGASGEQHMIHYGSTDDAVDETSAASSVASKTTPLSA      120

sp|Q0TLK2|PAND_ECOL5                    ------------------------------------------------------------      0
sp|P9WIL2|PAND_MYCTO                    ------------------------------------------------------------      0
tr|A7U8C7|A7U8C7_TRICA                  TFSDAVLSSDEELFHQKCPKPAPIYSPVSKPVSFE------------------SL-----      56
tr|B8I983|B8I983_CLOCE                  ---------------------MELTFN---PQNFSPEETQSGETKKEILDLFSVTDSSLL      36
tr|K9TK94|K9TK94_9CYAN                  ----------------------MLQIH---PVNSSIE-TWNPSIEGTILSLFDPSPQA-E      33
tr|A0A1Y1ZL74|A0A1Y1ZL74_9FUNG          -------------------------MVDKETI-----KTEENNTKENVLKFFKDC----E      26
tr|A0A1Y2AA19|A0A1Y2AA19_9FUNG          -------------------------MIVEEKL-----IQRLIIVNLMKRVFLNLF----E      26
tr|F4NWP2|F4NWP2_BATDJ                  SASKAVFPLDL--FLKQRSS-VDLRIRNDSTVAFD-DNHQSFHNQDSVLHYFVPT----T      110
tr|A0A1S8W5A4|A0A1S8W5A4_9FUNG          SAHKAVFPLDS--FLQSSST-IDLRIRSNDAVG----HPTSFHDHDSVLHYFVPT----T      169

sp|Q0TLK2|PAND_ECOL5                    ------------------------------------------------------------      0
sp|P9WIL2|PAND_MYCTO                    ------------------------------------------------------------      0
tr|A7U8C7|A7U8C7_TRICA                  PN--------RRLHEEFLRSSVDVLLQEAVFEGTNRKNRVLQWREPEELRRLMDFGVRSA      108
tr|B8I983|B8I983_CLOCE                  KARNETDRRVSEVINEFIGGSEV---------ST----------DVRLNDLLTNFCKSEI      77
tr|K9TK94|K9TK94_9CYAN                  NWEKNVDQQILKIAYQFLRQIEA---------SS----------DIELPDLMAQFQEFHL      74
tr|A0A1Y1ZL74|A0A1Y1ZL74_9FUNG          INEENVQQYIDSIIKEFYKPSPSI------YAAN----------TDTVKELQSKFGDSSI      70
tr|A0A1Y2AA19|A0A1Y2AA19_9FUNG          NDIQEVQMYIDKIIKDFYKPSPSV------YTIN----------SNSEKELMLKFGNSTI      70
tr|F4NWP2|F4NWP2_BATDJ                  KSEKLLDQYISGVIEAFLDEPAPI------YATA----------VSRTHSLTGKFGGLSI      154
tr|A0A1S8W5A4|A0A1S8W5A4_9FUNG          QSEKLLDKYISGVIEAFLHEPSPI------YTTS----------THIPHDPQSAFGGLSI      213

sp|Q0TLK2|PAND_ECOL5                    ------------------------------------------------------------      0
sp|P9WIL2|PAND_MYCTO                    ------------------------------------------------------------      0
tr|A7U8C7|A7U8C7_TRICA                  P-------STHE-ELLEVLKKVVTYSVKTGHPYFVNQLFSAVDPYGLVAQWATDALNPSV      160
tr|B8I983|B8I983_CLOCE                  PDKP----LNVDEYIAFFNKNIIRHSTRTSSPRYIGHMTSALPYFIKPLAGLIVTLNQNV      133
tr|K9TK94|K9TK94_9CYAN                  PDDA----SSFDTYLESLTKTLIPHSIHTSSPRFIGHMTSALPHFMRPLAKLMTALNQNL      130
tr|A0A1Y1ZL74|A0A1Y1ZL74_9FUNG          PKGIVCDKSNIKNYFSEIKTNVIDKSTRVSSPNMVGHMTSALPFFHRPLSKLITSLNQNV      130
tr|A0A1Y2AA19|A0A1Y2AA19_9FUNG          PKGISYDKKDIINYFFKIKTNVVENATRVSSPYMMGHMTSALPFFHKHISKLISSLNQNV      130
tr|F4NWP2|F4NWP2_BATDJ                  PDGNRSSVADLESYLNHLKLNVIDKSTRTGSSKMIGHMTTALPFFHRPLARLLAALNQNV      214
tr|A0A1S8W5A4|A0A1S8W5A4_9FUNG          PDGGRPSGSDLEAYLNHLKTNVIDRSTRTASSRMIGHMTTALPFFHRPLARLLAALNQNV      273

sp|Q0TLK2|PAND_ECOL5                    ----------MIRTMLQGKLHRVKVTHADLHY----------------------------      22
sp|P9WIL2|PAND_MYCTO                    ----------MLRTMLKSKIHRATVTCADLHY----------------------------      22
tr|A7U8C7|A7U8C7_TRICA                  YTYEVSPVFVLMEEVVLREM-RAIVGFE----------GGKGDGIFCPGGSIANGYAISC      209
tr|B8I983|B8I983_CLOCE                  VKAETAKVLTPYERQVIAMLHRAIYQFDSDFYNHHIQNRESVLGILTSGGTIANLTALWC      193
tr|K9TK94|K9TK94_9CYAN                  VKTETAKAFTPYERQALGILHRSLYQFPDEFYDYHLQNSHSTLGMVVSGGTSANFTAIWC      190
tr|A0A1Y1ZL74|A0A1Y1ZL74_9FUNG          VKVETSATMTFLERETLSKLHRAFYNNTDEFYEKYLSTTEGTFGCVTSGGTIANLTAMWV      190
tr|A0A1Y2AA19|A0A1Y2AA19_9FUNG          VKVETSSTMTFLERETLSKLHRAFYNMEDNYYKELLTKNDGCFGCITSGGTIANLTALWI      190
tr|F4NWP2|F4NWP2_BATDJ                  VKIETASTFTNLERQTLAMLHKAFFGSSDEFYTRYAYAPEYALGVMASGGTIANITALWI      274
tr|A0A1S8W5A4|A0A1S8W5A4_9FUNG          VKIETASTFTNLERQTLGMLHKAFYDLPDTFYQRHAYAPDYSLGVMASGGTIANITALWI      333
                                                    .      : :

sp|Q0TLK2|PAND_ECOL5                    ------------------------------------EGSCAIDQDFLDAAGILENEAIDI      46
sp|P9WIL2|PAND_MYCTO                    ------------------------------------VGSVTIDADLMDAADLLEGEQVTI      46
tr|A7U8C7|A7U8C7_TRICA                  ARYRFMPD--------IKKKGLH------SLPRLVLFTSEDAHYSIKKLASFQGIGTDNV      255
tr|B8I983|B8I983_CLOCE                  ARNSALGP--CEGFNGVEKEGMDAALKFYGYKDAVIIGSAMMHYSFEKAADLLGIGSNNS      251
tr|K9TK94|K9TK94_9CYAN                  ARNSSLGP--KGDFKGIEKEGLAAALEFYGYQGQVIIGSSLMHYSFEKAADLLGIGSQGL      248
tr|A0A1Y1ZL74|A0A1Y1ZL74_9FUNG          ARNNAFPK--TEDFEGIEKEGVISALSYYGYNNMAIVGSELMHYSFKKAADILGMGLKNI      248
tr|A0A1Y2AA19|A0A1Y2AA19_9FUNG          SRNNAFPK--TDNFNGIEKEGIISALKYYNYNDIAIIGSELMHYSFKKAADILGIGLNKI      248
tr|F4NWP2|F4NWP2_BATDJ                  ARNKALAVNAANGCRGIDKEGYISAMLKYGYKRAVIIGSTLMHYSFKKAADLLGLGEEGL      334
tr|A0A1S8W5A4|A0A1S8W5A4_9FUNG          ARNKALAPNSSNGCRGIDKEGVVSAMGKYGYKRAVIIGSALMHYSFKKAADLLGLGEEGL      393
                                                                              *   . .: . *.:

sp|Q0TLK2|PAND_ECOL5                    WNVTNGKRF----STY----AIAAERGSRIISVNGAAAH--CASVGDIVIIASFVN---M      93
sp|P9WIL2|PAND_MYCTO                    VDIDNGARL----VTY----AITGERGSGVIGINGAAAH--LVHPGDLVILIAYAT---M      93
```

```
tr|A7U8C7|A7U8C7_TRICA              YLIRTDARGRMDVSHLVEEIERSLREGAAPFMVSATAGTTVIGAFDPIEKIADVCQKYKL      315
tr|B8I983|B8I983_CLOCE              IKIPVDKNNHIDLSELRKTVEECHRKRRLIIAIIANAGTTDCGAIDPIERVAEIAYKEGC      311
tr|K9TK94|K9TK94_9CYAN              IKIPASRHNRIDLDCLRETVAECRAQKQHIIAIVGIAGTTDSGGIDPIEEMAAIAQAAGV      308
tr|A0A1Y1ZL74|A0A1Y1ZL74_9FUNG      YTIPVDSDYKMRIDLLIEKLEELKQKKIKVIAIVGIACTTEVGSIDKLDILADIAKKYGA      308
tr|A0A1Y2AA19|A0A1Y2AA19_9FUNG      YTIPVDSEYKIRIDLLTEKLQELKQKRIKVITIVGIASTTEVGSIDDLEKLADLAEEYGA      308
tr|F4NWP2|F4NWP2_BATDJ              VLIPVDDAFRMRIDVLKAKVEKCVAENTLVIAIVGISGTTETGSIDPLLDIACIAHKYHI      394
tr|A0A1S8W5A4|A0A1S8W5A4_9FUNG      CLIPTDAHFCMRVDLLKSKVDELISEGALIIAIVGIAGTTETGSIDPLFDIYSIAHRHNI      453
                                      :  .                   .    : : . :        . :  :

sp|Q0TLK2|PAND_ECOL5                PDEEARTWRPNVAYFEGDNEMKRT---AKAIPVQVA------------------------      126
sp|P9WIL2|PAND_MYCTO                DDARARTYQPRIVFVDAYNKPIDMGHDPAFVPENAGELLDPRLGVG--------------      139
tr|A7U8C7|A7U8C7_TRICA              WLHVDAAWGGGALVSAKHRHLLKGIERADSVTWNPHKLLTAPQQCSTLLLRHEGVLAEAH      375
tr|B8I983|B8I983_CLOCE              HFHVDAAWGGPLLFSDKYRNRLKGIELADSVTIDGHKQLYLPMGLGMIFMRNPQAA-KSI      370
tr|K9TK94|K9TK94_9CYAN              HFHVDAAWGGPLIFSQQHRHKLAGIEQADSVTIDAHKQLYAPMGIGMVMFQNPQLA-KAI      367
tr|A0A1Y1ZL74|A0A1Y1ZL74_9FUNG      LFHVDGAWGGSFIMSNKYHHLLKGIEKADTITIDGHKLLYTPMGCGIVLFKSPLLPTTTI      368
tr|A0A1Y2AA19|A0A1Y2AA19_9FUNG      MFHVDGAWGGSFILSDKYRHLLKGIQRADTITIDGHKLLYTPIGCGVVLFKSPFLPTTTI      368
tr|F4NWP2|F4NWP2_BATDJ              HFHVDAAWGGPLIFSPEHSSKLAGISQADTITVDGHKQLYTPMGLGILLAKCPSLV-TFI      453
tr|A0A1S8W5A4|A0A1S8W5A4_9FUNG      HFHVDAAWGGPLIFSPEHRCKLNGISQADSITVDGHKQLYTPMGLGILLLRSPSLA-LYI      512
                                          ::                      :  :

sp|Q0TLK2|PAND_ECOL5                ------------------------------------------------------------      126
sp|P9WIL2|PAND_MYCTO                ------------------------------------------------------------      139
tr|A7U8C7|A7U8C7_TRICA              STNAAYLFQKDKFYDTKYDTGDKHIQCGRRADVLKFWFMWKAKGTSGLEKHVDKVFENAR      435
tr|B8I983|B8I983_CLOCE              EKNSNYIIRRGS-----IDLGRRSLEGSRPAMSIYLHAALNIIAKGGYEFLINEGIKKAE      425
tr|K9TK94|K9TK94_9CYAN              EKHACYTVREGS-----ADLGQRSLEGSRPAMSLFLHAGLHAIGLKGYEFLIDEGIRKTQ      422
tr|A0A1Y1ZL74|A0A1Y1ZL74_9FUNG      RKVASYVIRAGS-----MDHGKTTLEGSRPANVLYLHASLNLLGENGLSQLLDIGVNNAK      423
tr|A0A1Y2AA19|A0A1Y2AA19_9FUNG      RKVANYIIKTDS-----MDHGKTTLEGSRPANALYIHASLNILGQHGLRYLLETSIENTK      423
tr|F4NWP2|F4NWP2_BATDJ              RKTAGYVIRHDS-----PDLGKFTLEGSRPANVLYLHASLNLLGKQGLGTLMTRSVTVVK      508
tr|A0A1S8W5A4|A0A1S8W5A4_9FUNG      RKTASYVIRNDS-----PDLGKYTLEGSRPANALYLHASLSLLGKHGLGILVTRSVTIVR      567

sp|Q0TLK2|PAND_ECOL5                ------------------------------------------------------------      126
sp|P9WIL2|PAND_MYCTO                ------------------------------------------------------------      139
tr|A7U8C7|A7U8C7_TRICA              FFTDCIKNRE--GFEMVIAEPEYTNICFWYVPKSLRGRKDEADYKDKLHKVAPRIKERMM      493
tr|B8I983|B8I983_CLOCE              YMAGLIKSMR--EFE-LLTEPDMNMLLYRFIPERLRGKAADHMLDDA-------------      469
tr|K9TK94|K9TK94_9CYAN              YMAEQVRLSP--EFE-LLAEPEINLLIYRYIPEQLREFVAKGELTET-------------      466
tr|A0A1Y1ZL74|A0A1Y1ZL74_9FUNG      YMAELIKSYP--QFE-LISKPVTNIFLYRYIPSWLRKKSYEIEQIEVKSISSPISTVSTI      480
tr|A0A1Y2AA19|A0A1Y2AA19_9FUNG      YMYNLINLYP--------------------------------------------------      433
tr|F4NWP2|F4NWP2_BATDJ              QMAVRLNLHPSQSFQ-TLHEPMSNVLLYRYIPSDLRESIADGTYVPNPEDEDRVSEVT--      565
tr|A0A1S8W5A4|A0A1S8W5A4_9FUNG      QTAVRLDSHPSRCFQ-ILHQPMSNLLLYRYVPSALRDAIADGSYISTKDDEAWISEAT--      624

sp|Q0TLK2|PAND_ECOL5                ------------------------------------------------------------      126
sp|P9WIL2|PAND_MYCTO                ------------------------------------------------------------      139
tr|A7U8C7|A7U8C7_TRICA              ----------------------------------------------KEGSMM--------      499
tr|B8I983|B8I983_CLOCE              ------------------------------------------------------------      469
tr|K9TK94|K9TK94_9CYAN              ------------------------------------------------------------      466
tr|A0A1Y1ZL74|A0A1Y1ZL74_9FUNG      PSPTQATTALPIIESEEENHNDNDNSITFCNDTLKDQISDISDFTDKEGSLIYEESHYND      540
tr|A0A1Y2AA19|A0A1Y2AA19_9FUNG      ------------------------------------------------------------      433
tr|F4NWP2|F4NWP2_BATDJ              -------KRLQIYQA-------------SCIP---------------SDAV-FPEVSTHA      589
tr|A0A1S8W5A4|A0A1S8W5A4_9FUNG      -------RRIQIRQA-------------SCVS---------------VGSV-HPHSLTSP      648

sp|Q0TLK2|PAND_ECOL5                ------------------------------------------------------------      126
sp|P9WIL2|PAND_MYCTO                ------------------------------------------------------------      139
tr|A7U8C7|A7U8C7_TRICA              --------------------------------VTYQAQKGHPNFFRIVFQNSGLDKADMV      527
tr|B8I983|B8I983_CLOCE              -DNEVINKYNEQLQKLQRNGGYSFVSRTSFSSELYQN--KSLVALRAVLANPLTCEANIN      526
tr|K9TK94|K9TK94_9CYAN              -QNQAIDLVNEQLQKAQRHAGKTFIARTITQTTRYGP--MAIVALRAALANPLTTEADID      523
tr|A0A1Y1ZL74|A0A1Y1ZL74_9FUNG      EENAIIDYFTTELQARQKREGEGFVSKTTVKTTKYPTYSKGVDVLRVVIANPLVTEDNIK      600
tr|A0A1Y2AA19|A0A1Y2AA19_9FUNG      ------------------------------------------------------------      433
tr|F4NWP2|F4NWP2_BATDJ              TDES--TNGEAGLHSQQNQLQGGFVSRTRVLF-----KGFHVNALRVVVANPLTTWRDVE      642
tr|A0A1S8W5A4|A0A1S8W5A4_9FUNG      NPTG--GIDPPAPFGQQQQPLPGFVSRTRVWF-----RGHHVDALRVVVANPLTTWDDIE      701

sp|Q0TLK2|PAND_ECOL5                ------------------------------------------------------------      126
sp|P9WIL2|PAND_MYCTO                ------------------------------------------------------------      139
tr|A7U8C7|A7U8C7_TRICA              HLVEEIERLGSDL-----------------------------------------------      540
tr|B8I983|B8I983_CLOCE              EVINDQLNIAKRLNL---------------------------------------------      541
tr|K9TK94|K9TK94_9CYAN              AVLDDQIALATQLNLGGSQF----------------------------------------      543
tr|A0A1Y1ZL74|A0A1Y1ZL74_9FUNG      AVVEEQIKLGEIIEEEYKMNHKLIK-----------------------------------      625
tr|A0A1Y2AA19|A0A1Y2AA19_9FUNG      ------------------------------------------------------------      433
tr|F4NWP2|F4NWP2_BATDJ              GVISDQLKMGAMIEEEMKREQMVKRVRDMYPMHRPNA-----VVPDSAKSCTALAACDGT      697
tr|A0A1S8W5A4|A0A1S8W5A4_9FUNG      GVISDQLHIGAEIEAEMQREQMVKRLQSWISTPCSSVSDTMLSEPEAAKATLSFVSAT-D      760

sp|Q0TLK2|PAND_ECOL5                -------------------     126
sp|P9WIL2|PAND_MYCTO                -------------------     139
tr|A7U8C7|A7U8C7_TRICA              -------------------     540
tr|B8I983|B8I983_CLOCE              -------------------     541
tr|K9TK94|K9TK94_9CYAN              -------------------     543
tr|A0A1Y1ZL74|A0A1Y1ZL74_9FUNG      -------------------     625
tr|A0A1Y2AA19|A0A1Y2AA19_9FUNG      -------------------     433
tr|F4NWP2|F4NWP2_BATDJ              DTCKPDDKIEWWPGWPFDL     716
tr|A0A1S8W5A4|A0A1S8W5A4_9FUNG      LHNGSHGDPGWWPGWPFDL     779
```

**Supplementary File 4: Variant calling results.** Provided as tab-delimited text file showing non-synonymous (NSY) and synonymous (SYN) mutations found after whole-genome sequencing analysis of strains IMX2300 and IMX2300-1. First column shows GenBank identifiers for the corresponding *Saccharomyces cerevisiae* CEN.PK113-7D chromosome. Links are provided to the amino acid sequence deposited in GenBank and to the *Saccharomyces* Genome Database for CEN.PK113-7D proteins with an *S. cerevisiae* S288C.

```
#ID	Position	Strain	Description	Links
CP046083.1	343428	IMX2300,IMX2300-1	CDS,gene-SCEN_C01950,rna-
gnl|wl|mrna.SCEN_C01950,SCEN_C01950,trans_orient:+,loc_in_cds:703,codon_pos:1,codon:Act-Gct,pep:T->A,Thr-235-Ala,(NSY)
	https://www.ncbi.nlm.nih.gov/protein/QHB07239
CP046083.1	344005	IMX2300,IMX2300-1	CDS,gene-SCEN_C01960,rna-
gnl|wl|mrna.SCEN_C01960,SCEN_C01960,trans_orient:+,loc_in_cds:324,codon_pos:3,codon:atA-atG,pep:I->M,Ile-108-Met,(NSY)
	https://www.ncbi.nlm.nih.gov/protein/QHB07240
CP046084.1	2165	IMX2300-1	CDS,gene-SCEN_D00100,rna-
gnl|wl|mrna.SCEN_D00100,SCEN_D00100,trans_orient:+,loc_in_cds:467,codon_pos:2,codon:tTg-tCg,pep:L->S,Leu-156-Ser,(NSY)
	https://www.ncbi.nlm.nih.gov/protein/QHB07243
CP046084.1	2171	IMX2300-1	CDS,gene-SCEN_D00100,rna-
gnl|wl|mrna.SCEN_D00100,SCEN_D00100,trans_orient:+,loc_in_cds:473,codon_pos:2,codon:cAt-cCt,pep:H->P,His-158-Pro,(NSY)
	https://www.ncbi.nlm.nih.gov/protein/QHB07243
CP046084.1	2174	IMX2300-1	CDS,gene-SCEN_D00100,rna-
gnl|wl|mrna.SCEN_D00100,SCEN_D00100,trans_orient:+,loc_in_cds:476,codon_pos:2,codon:tTt-tCt,pep:F->S,Phe-159-Ser,(NSY)
	https://www.ncbi.nlm.nih.gov/protein/QHB07243
CP046087.1	1032619	IMX2300,IMX2300-1	CDS,gene-SCEN_G05290,rna-gnl|wl|mrna.SCEN_G05290,SCEN0G05290,trans_orient:-
,loc_in_cds:943,codon_pos:1,codon:Ttg-Atg,pep:L->M,Leu-315-Met,(NSY)	https://www.ncbi.nlm.nih.gov/protein/QHB08902	Mtm1
	https://www.yeastgenome.org/locus/S000003489
CP046088.1	1995	IMX2300	CDS,gene-SCEN_H00100,rna-gnl|wl|mrna.SCEN_H00100,SCEN_H00100,trans_orient:-,loc_in_cds:2129,codon_pos:2,codon:gGa-
gCa,pep:G->A,Gly-710-Ala,(NSY)	https://www.ncbi.nlm.nih.gov/protein/QHB08943
CP046091.1	673935	IMX2300,IMX2300-1	CDS,gene-SCEN_K03430,rna-
gnl|wl|mrna.SCEN_K03430,SCEN_K03430,trans_orient:+,loc_in_cds:853,codon_pos:1,codon:Agt-Ggt,pep:S->G,Ser-285-Gly,(NSY)
	https://www.ncbi.nlm.nih.gov/protein/QHB10105
CP046083.1	344449	IMX2300,IMX2300-1	CDS,gene-SCEN_C01960,rna-
gnl|wl|mrna.SCEN_C01960,SCEN_C01960,trans_orient:+,loc_in_cds:768,codon_pos:3,codon:gcG-gcA,pep:A->A,Ala-256-Ala,(SYN)
	https://www.ncbi.nlm.nih.gov/protein/QHB07240
CP046084.1	1190735	IMX2300-1	CDS,gene-SCEN_D06170,rna-
gnl|wl|mrna.SCEN_D06170,SCEN_D06170,trans_orient:+,loc_in_cds:42,codon_pos:3,codon:gaT-gaC,pep:D->D,Asp-14-Asp,(SYN)
	https://www.ncbi.nlm.nih.gov/protein/QHB07833
CP046084.1	1190738	IMX2300-1	CDS,gene-SCEN_D06170,rna-
gnl|wl|mrna.SCEN_D06170,SCEN_D06170,trans_orient:+,loc_in_cds:45,codon_pos:3,codon:acA-acC,pep:T->T,Thr-15-Thr,(SYN)
	https://www.ncbi.nlm.nih.gov/protein/QHB07833
CP046084.1	390159	IMX2300,IMX2300-1	CDS,gene-SCEN_D02160,rna-gnl|wl|mrna.SCEN_D02160,SCEN0D02160,trans_orient:-
,loc_in_cds:1569,codon_pos:3,codon:aaC-aaT,pep:N->N,Asn-523-Asn,(SYN)	https://www.ncbi.nlm.nih.gov/protein/QHB07444	Gpr1
	https://www.yeastgenome.org/locus/S000002193
CP046084.1	856084	IMX2300,IMX2300-1	CDS,gene-SCEN_D04500,rna-
gnl|wl|mrna.SCEN_D04500,SCEN_D04500,trans_orient:+,loc_in_cds:423,codon_pos:3,codon:gaT-gaC,pep:D->D,Asp-141-Asp,(SYN)
	https://www.ncbi.nlm.nih.gov/protein/QHB07671
```

CP046084.1	856087	IMX2300,IMX2300-1	CDS,gene-SCEN_D04500,rna-gnl|wl|mrna.SCEN_D04500,SCEN_D04500,trans_orient:+,loc_in_cds:426,codon_pos:3,codon:gaT-gaC,pep:D->D,Asp-142-Asp,(SYN)	https://www.ncbi.nlm.nih.gov/protein/QHB07671

CP046084.1	856099	IMX2300-1	CDS,gene-SCEN_D04500,rna-gnl|wl|mrna.SCEN_D04500,SCEN_D04500,trans_orient:+,loc_in_cds:438,codon_pos:3,codon:ttC-ttT,pep:F->F,Phe-146-Phe,(SYN)	https://www.ncbi.nlm.nih.gov/protein/QHB07671

CP046085.1	35976	IMX2300,IMX2300-1	CDS,gene-SCEN_E00250,rna-gnl|wl|mrna.SCEN_E00250,SCEN0E00250,trans_orient:+,loc_in_cds:615,codon_pos:3,codon:tcC-tcT,pep:S->S,Ser-205-Ser,(SYN)	https://www.ncbi.nlm.nih.gov/protein/QHB08008	Npr2	https://www.yeastgenome.org/locus/S000000788
